# Iterative Gene Enrichment Analysis: interpretable networks for human and AI-assisted biological insights

**DOI:** 10.64898/2026.09.06.749570

**Authors:** Maxim V. Kuleshov, Yair Benita

## Abstract

**Background:** Over-representation analysis (ORA) is widely used to interpret gene lists from high-throughput biological experiments. However, ORA often produces long and fragmented lists of enriched terms that are difficult to translate into coherent, testable biological hypotheses. This challenge is amplified by the rapid growth of gene set collections and the need to integrate results across multiple gene-set collections. Large language models (LLMs) can assist in summarizing enrichment outputs, but their effectiveness is limited by the lack of reproducibility and statistically supported representations.

**Results:** We introduce iterative Gene Enrichment Analysis (iGEA), a software framework that transforms ORA-based enrichment outputs into structured, interpretable networks, supporting human interpretation and AI-assisted exploration. iGEA iteratively selects the most significant enriched term, removes its overlapping genes from the input list, and repeats enrichment until no significant terms remain. Applied independently across gene-set collections, this procedure yields a compact set of non-overlapping enriched terms within each collection. Integration of collection-specific results yields a cross-collection gene-term network in which genes connect terms from different collections. Using a published set of HIV dependency factors, iGEA identified five compact modules spanning secretory trafficking, nuclear transport, transcription elongation, proteostasis, and innate immune signaling, enabling rapid hypothesis generation and interactive exploration. The resulting network structure supports standardized prompting and provides a structured representation for LLM-assisted summarization and exploration of enrichment results.

**Conclusionsn:** iGEA provides a software framework for gene enrichment analysis that addresses the interpretation challenge of ORA by generating compact, empirically benchmarked cross-collection gene-term networks. This network representation reduces within-collection redundancy, exposes relationships across gene-set collections, and provides a structure that makes modules and hub genes easier to identify through visualization, network-based analysis, and LLM-assisted exploration.

**Availability:** Source code is available at: https://github.com/aion-labs/Gene-Enrichment-Analysis

A web-based version of the application is available at: https://iterative-gene-enrichment.streamlit.app

## Background

High-throughput molecular profiling studies frequently generate gene lists of interest, most commonly from differential expression analysis, genome-wide association studies, or large-scale perturbation screens. A standard approach to interpret gene lists is gene set enrichment, where the input is compared to curated gene sets to identify biologically informative overlaps. Two broad classes of methods are commonly used: over-representation analysis (ORA), based on the hypergeometric test [1, 2] and Gene Set Enrichment Analysis (GSEA), which analyzes ranked gene lists [3]. In this study, we focus on ORA-based methods. The goal of enrichment analysis is to formulate testable biological hypotheses. However, translating enrichment outputs into coherent and reliable biological interpretations remains challenging, particularly as gene set collections expand and results must be integrated across many collections.

For the past two decades, the statistical foundations of over-representation analysis have changed little. Progress has instead come from expansion of gene set databases, such as MSigDB [4] and Enrichr [2], which now contain 30 and 218 collections, respectively (as of August 2025), covering transcriptional regulation, pathways, disease signatures, and functional ontologies. As a result, enrichment analysis has increasingly become an interpretation bottleneck: users must synthesize many partially overlapping terms across multiple collections into a coherent, testable hypothesis. This synthesis step is difficult to perform reliably and is not well supported by existing enrichment outputs.

Many enrichment tools attempt to improve interpretability by organizing, clustering, or simplifying long lists of terms. These methods can be broadly categorized into three approaches: clustering of terms by partial or full gene overlap, clustering based on ontology structure (as in Gene Ontology), and network-based clustering of gene-term networks. Examples include DAVID [5] and ClusterProfiler [6] for gene overlap clustering, Revigo [7] for Gene Ontology based clustering and EnrichmentMap for network-based clustering and visualization [8]. Related approaches have also addressed non-redundant GO summarization using semantic similarity, ontology structure, or term relatedness [9, 10]. While these methods help organize enrichment results, the challenge of synthesizing a coherent and testable biological hypothesis remains significant, particularly when integrating results across multiple collections.

Large language models (LLMs) have recently been explored as a complementary strategy. For example, GPT-based systems such as ChatGPT can generate plausible narratives about the functions of enriched gene sets [11, 12]. In practice, LLMs are most effective when they are given structured, constrained inputs that reflect statistical evidence and expose interpretable relationships. Unstructured term lists provide little topology and may encourage narrative drift.

Here, we present iterative Gene Enrichment Analysis (iGEA), a framework designed to produce interpretable, cross-collection representations of enrichment results that support hypothesis generation and structured LLM-assisted exploration. iGEA begins with a standard ORA within a given collection, selects the most significant term, assigns the overlapping genes to that term, and removes them from the input list before repeating the analysis on the remaining genes. This iterative procedure reduces the dominance of highly overlapping terms associated with the strong enrichment signals (e.g., cell cycle), allowing additional enriched terms to be prioritized in subsequent iterations. As a result, iGEA can identify enriched terms that are not retained by FDR-filtered ORA and generates a compact within-collection representation of enrichment results. These collection-specific outputs are subsequently integrated into a cross-collection gene-term network.

The central design goal of iGEA is to make enrichment results interpretable as a network rather than as a list of terms. After iGEA is applied separately to each gene-set collection, the collection-specific outputs are integrated through their shared genes. In this cross-collection gene-term network, genes act as links between biological annotations derived from different sources. Because within-collection redundancy is reduced before integration, modules and hub genes in the resulting network are no longer dominated by repeated terms within the same collection and more likely to reflect convergence across complementary annotation sources. This makes the iGEA output suitable for network-based exploration, including identification of connected components, modules, bridge genes, and hubs, as well as structured prompting for conversational AI tools.

We demonstrate iGEA by analyzing a list of 275 genes required for HIV infection that were identified using a genome-wide siRNA knockdown screen [13]. This dataset provides a useful benchmark because several related HIV dependency-factor screens produced poorly overlapping gene lists [14, 15], whereas subsequent cross-study meta-analyses showed that reproducible biological signals could be identified at the pathway and process level [16, 17]. Meta-analysis therefore provides an approximate pathway-level reference for distinguishing robust biological signal from study-specific noise. We use this benchmark to ask whether iGEA, applied to a single HIV screen, can recover pathway-level biology supported by prior cross-study analyses. We show that iGEA supports biological interpretation by producing a compact cross-collection network that can be inspected directly by human experts or used as a structured input for standardized LLM-assisted exploration.

## Implementation

### Gene Enrichment Analysis

To compute gene set enrichment, we performed over-representation analysis (ORA) using a hypergeometric test implemented with SciPy’s hypergeom.sf() function, which returns the probability of observing at least the given overlap by chance. For each gene set collection, we first filtered gene sets by size to include only those within a user-defined range (10-600 genes per gene set by default). This filtering improves interpretability and stability by excluding extremely small gene sets, where minimal overlap can yield disproportionately low p-values, and very large gene sets that are often biologically broad. In an iterative framework, such broad categories could remove many genes from the input list while providing limited biological insight. The selected default range captures the majority of curated pathway terms across the collections used in this study while excluding statistical extremes.

For each collection, the input gene list and the background gene list were intersected with the genes represented in that collection after filtering. The hypergeometric test was then computed using the parameters (n_overlap - 1, M, n_term, N), where n_overlap is the number of overlapping genes between the input list and the term, M is the collection-specific background size, n_term is the number of genes in the tested term, and N is the collection-specific input size. For example, if the input list contains 400 genes but only 350 are represented in the selected collection, then N = 350. If the background contains 10,000 genes but only 4,500 are represented in the selected collection, then M = 4,500. If a tested term contains 50 genes and overlaps with 15 genes from the collection-specific input list, the p-value is computed as hypergeom.sf(15 -1, 4500, 50, 350).

### Iterative Gene Enrichment Analysis

iGEA is performed separately for each gene-set collection. The process begins with an ORA on the full input gene list. The most significant enriched term is selected and recorded. The genes overlapping with this term are then assigned to that term and removed from the input gene list. ORA is repeated on the remaining genes, and the next most significant term is selected. This iterative selection and assignment procedure continues until no additional terms meet the user-defined significance, by default p<0.01. This process is also described by a pseudo-algorithm in Supplementary Table 1.

### Gene-Set Collection

Gene-set collections were obtained from the MSigDB (version 2025.1) [4]. For this study, we focused on canonical pathway collections and Gene Ontology collections. In addition, we constructed a protein interaction-derived gene-set collection using STRING-DB [18]. Human protein-protein interaction data (STRING v12.0) were filtered to retain high-confidence interactions with score ≥ 0.9. Each protein was defined as a “term,” and its interacting partners were used to define the gene set. The resulting collection included 12,323 gene sets covering 12,320 genes with an average of 16.4 genes per set. The collection was exported in GMT format for compatibility with the MSigDB-based pipeline. Including this interaction-derived collection provides an orthogonal perspective that can complement pathway and ontology annotations in the cross-collection iGEA network.

### Cross-collection iGEA Network

The primary output of iGEA is a cross-collection gene-term network designed for biological interpretation by both human experts and conversational AI tools. Results from individual collections are integrated into a gene-term network. An edge is created between a gene and a term if the gene was assigned to that term by iGEA in that collection. No edge weights are applied. Nodes are colored by collection to facilitate cross-collection interpretation by visualization. Networks were exported using the DOT language (Graphviz) and visualized interactively using Plotly. Importantly, the non-overlap constraint is applied only within each gene-set collection. The cross-collection network is generated through genes associated with enriched terms in different gene-set collections.

### Statistical Analysis of iGEA Networks

Statistical significance was assessed at the cross-collection network level rather than at the individual term level. Conventional FDR correction for enrichment analysis assumes that the input gene list is fixed. In iGEA, the input gene list is adaptively modified after each selected term, thus later tests depend on earlier selections. Dependency-aware procedures such as Benjamini-Yekutieli FDR address correlated tests in fixed hypothesis settings, but they do not directly resolve this adaptive gene-removal process [19]. We therefore use raw p-values for iterative term selection and evaluate statistical support at the level of the final cross-collection network using random-gene-list benchmarks. To establish empirical null distributions, iGEA was applied to random gene lists spanning sizes from 50 to 1000 genes (step size 50). For each gene list size, 1000 random lists were generated, resulting in 20,000 random iGEA runs. Null distributions were generated for 11 collections: eight pathway collections (Hallmark, Reactome, KEGG Legacy, KEGG Medicus, BioCarta, Pathway Interaction Database, Canonical Pathways, and WikiPathways) and three Gene Ontology collections (Biological Process, Molecular Function, and Cellular Component).

For each iGEA run, we constructed the cross-collection gene-term network and extracted the largest connected cluster. The following cluster-level metrics were computed: total cluster size (genes + terms), number of genes, number of terms, number of edges, average edges per gene, and the number of distinct collections represented in the cluster. Each observed iGEA result was benchmarked against the null distribution from the closest matching gene list size bin (e.g., an input list of 280 genes was benchmarked against the 300-gene null distribution). For each metric, significance was summarized as a z-score relative to the corresponding null distribution, computed on the largest connected cluster only. This network-level benchmarking provides an empirical measure of whether the observed network structure exceeds random expectation and can be used to guide downstream interpretation.

For benchmarking in each iGEA run, the user-selected raw p-value threshold was matched provided it was ≤ 0.01, and the maximum number of iGEA iterations was capped at 30. If the user selected a subset of collections, benchmarking was performed using the corresponding subset (up to 11 collections). Because generating 1000 random iGEA networks for every possible parameter combination is computationally intensive, some parameters were not matched during benchmarking, including minimum overlap and maximum gene set size. Random iGEA runs were generated using default settings for these parameters. Regardless of benchmarking constraints, the full iGEA results for the user-selected parameters are always reported.

### Interpretation by Artificial Intelligence

To demonstrate how iGEA supports LLM-assisted exploration, we provided the cross-collection iGEA network to a large language model (GPT-5, OpenAI) using a standardized prompt generated by the iGEA application (Supplementary Table 2). All 11 collections were provided to the model, including the Human Phenotype Ontology, so that phenotype associations were available as additional context. In this workflow, iGEA provides an empirically benchmarked network structure (genes, terms, and network topology), and the LLM operates as an exploratory interpretation layer to summarize modules, identify hubs, and propose hypotheses. Standardized prompting provides a consistent starting point for user-driven conversational exploration of specific clusters, biological themes, or gene-level relationships in the network.

## Results

To evaluate whether iGEA provides a more compact and structured representation of enrichment results, we applied iGEA and standard ORA to a published set of 275 HIV dependency factors identified by a genome-wide siRNA screen [13] (272 mapped to valid human gene symbols). iGEA was applied across 11 gene-set collections. The Human Phenotype Ontology returned terms that were phenotype descriptors unrelated to the experimental context (for example HP: BLUE SCLERAE, HP: CUPPED EAR, and HP: FOAMY URINE) and was not carried forward into the cross-collection network. The network analysis therefore includes 7 pathway collections and 3 Gene Ontology collections (10 collections total); per-collection enrichment results for all 11 collections are reported in Supplementary Tables 3 and 4.

Across the 10 collections, ORA identified 252 terms with p < 0.01, of which 87 (just over one third) remained significant after FDR correction (q < 0.05) (Supplementary Table 3). iGEA identified 62 significant terms (p < 0.01) across the same collections (Supplementary Table 4). For five collections, iGEA and FDR-adjusted ORA produced identical results. In four additional collections, all FDR-significant ORA terms were captured by iGEA, which also identified additional terms. In the remaining two collections (Reactome and GO Biological Process), overlap was limited and ORA produced a large number of redundant terms (Supplementary Figure 1).

To illustrate the conceptual differences between ORA and iGEA, and for simplicity, we focus on a single collection of Gene Ontology Biological Process (GOBP) as an example. For the HIV input gene list ORA identified 14 significant terms (FDR < 0.05) associated with 57 unique genes, forming four clusters dominated by Golgi-related functions (5 terms) and nuclear export (6 terms) (Figure 1A). In contrast, iGEA identified 10 significant terms (p < 0.01) associated with 96 unique genes. By design, each iGEA term was linked to a distinct subset of genes. Four terms were shared between methods (Golgi transport, nuclear export, glycolipid biosynthetic process, and protein-containing complex disassembly), each representing a cluster within ORA, while six additional terms were uniquely identified by iGEA (Figure 1B).

**Fig. 1.**
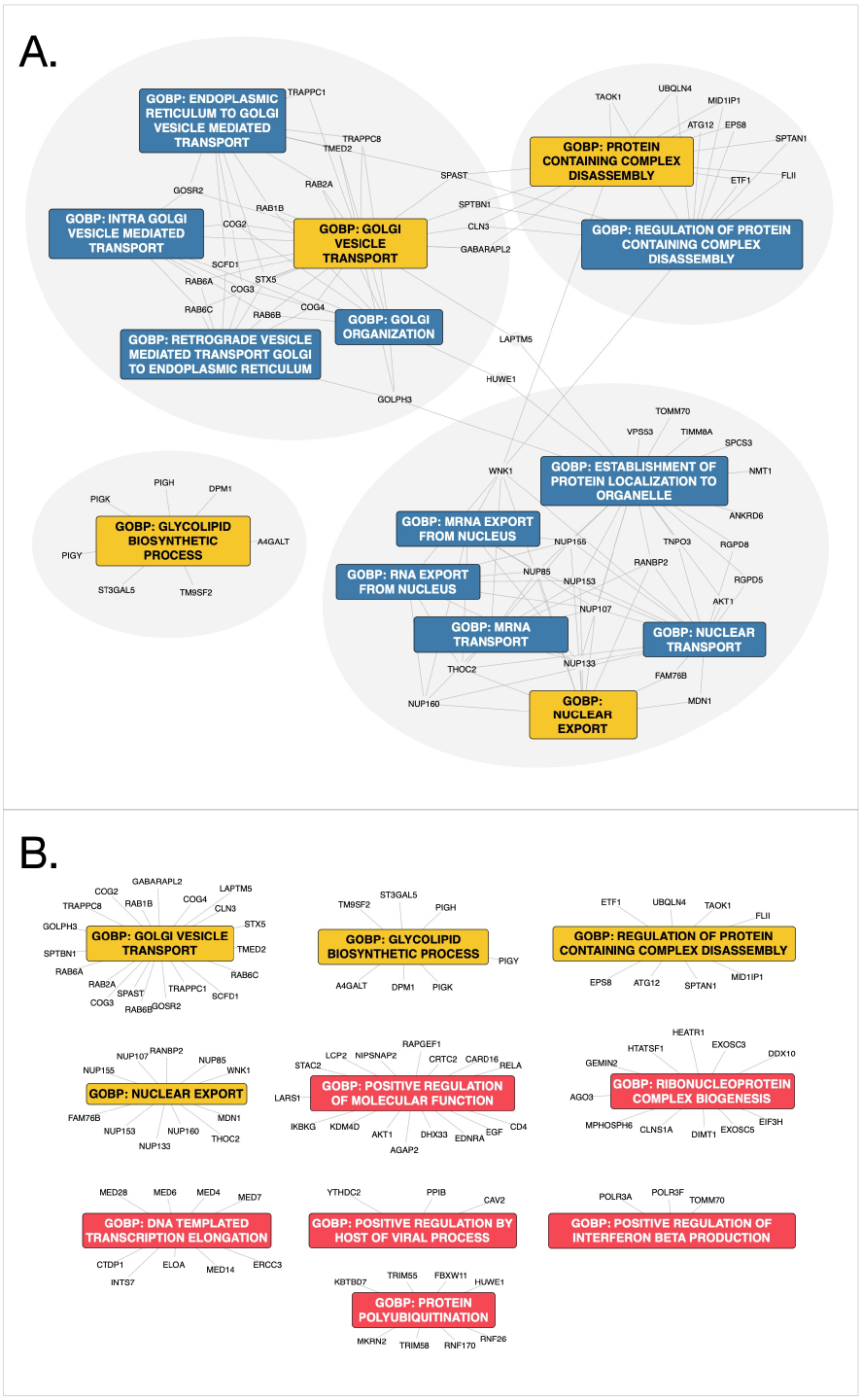
ORA and iGEA. Gene Ontology Biological Process results of ORA (A) and iGEA (B). Results are shown as a network of term-gene associations. Terms in blue were unique to ORA, terms in red unique to iGEA and terms in yellow were captured in both. ORA terms shown after FDR correction at q < 0.05 and iGEA terms are shown using the iterative raw p-value threshold p < 0.01.

Because iGEA performs iterative enrichment on adaptively reduced gene lists, conventional term-level multiple testing p-value correction is not directly applicable as the null hypothesis is violated. Instead, we assessed significance at the cross-collection network level (see Methods). Following iGEA, a gene-term network is generated for all tested collections. Clusters can only emerge when genes are linked to terms from different collections and statistics were computed on the largest connected cluster and compared to 1000 random gene lists of similar size, in this case 300. The largest connected cluster of the HIV iGEA network contained 179 genes and 62 terms, with an average of 2.3 edges per gene. Statistical benchmarking was performed using the 10 gene-set collections included in the network, for all of which permutation data were available. Compared to 1,000 random gene lists of size 300, these values corresponded to z-scores of 7.39 (number of genes), 8.44 (number of terms), and 3.00 (average edges per gene). The statistical report generated by iGEA is shown in Supplementary Table 5.

For visualization clarity, Figure 2 shows a reduced Gene Ontology (GO) only network generated from the three GO collections. This visualization contains 156 genes connected with 29 terms which is a subset of the larger network (Supplementary Figure 2). The same GO network with gene names is available in Supplementary Figure 3. Even in this simplified representation of 3 GO collections, interpretation based solely on visual inspection of the network is challenging. To assist with interpretation, we developed a standardized ChatGPT prompt (Supplementary Table 2). ChatGPT summarized the GO-based iGEA network as follows: “This network resolves into five tightly connected modules: ER-Golgi trafficking & glycosylation; nuclear pore/export & transcription/RNA processing; ubiquitin-proteasome/Cullin-RING ligases; small-GTPase/vesicle adaptors & endocytosis; and an innate-immune axis centered on POLR3A/F→TOMM70→IFN-β. Hub genes bridging these modules (e.g., NUPs, MED subunits, RAB1B/RAB2A, EXOSC3/5, FBXW11, SPAST, DPM1/STT3A/PIGY) suggest a coordinated state of transcriptional reprogramming, nucleocytoplasmic transport, vesicle remodeling, proteostasis, and antiviral signaling” (Supplementary Table 6). The ChatGPT response for the 11-collection network is shown in Supplementary Table 7 and can be summarized as dominated by an endomembrane/secretory-trafficking axis (Golgi/ER transport, clathrin endocytosis, Rab/ARF machinery) tightly linked to membrane protein processing (N-linked glycosylation + GPI-anchor synthesis) and a second axis of information flow control (nuclear pore/export + RNA decay). Layered on top is a growth-factor / inflammatory signaling core (EGF-EGFR-PI3K/AKT-NFκB; IL-12/IL-17-JAK/NFκB), suggesting a coordinated program coupling receptor trafficking/processing to signaling outputs.

**Fig. 2.**
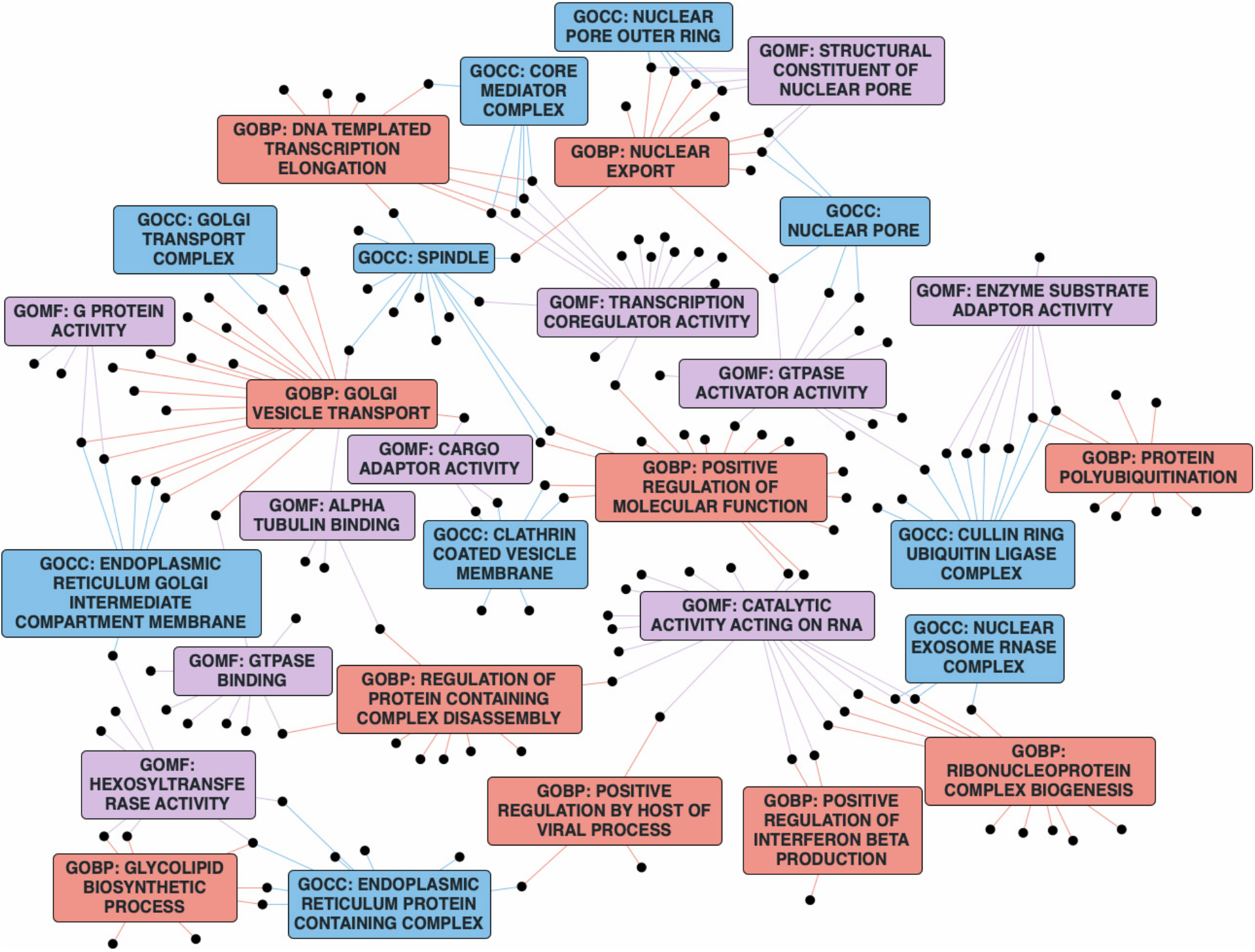
Iterative GEA network. Iterative GEA results shown as a network limited to Gene Ontology collections (Biological Process (GOBP, red); Molecular Function (GOMF, purple) and Cellular Component (GOCC, blue)). Genes from the input list overlapping with a term are shown as black dots. The network with full gene names is shown in Supplementary Figure 3. iGEA network generated using p < 0.01.

We next asked whether the iGEA network contained sufficient biological signal to support context-blinded interpretation by ChatGPT. In the initial prompt, the model was given only the network representation, including gene names, enriched term names, collection labels, and network topology; the experimental context and the way the gene list was generated were not provided. Based on this information ChatGPT proposed the following experimental context: “A perturbation/screen tied to cell-surface receptor trafficking and inflammatory signaling (EGFR/NFκB/JAK) with a strong contribution from secretory pathway capacity and glycoprotein maturation.” (Supplementary Table 7). It further proposed several possible contexts: “(i) A receptor-dependent signaling readout (e.g., EGFR pathway activation, NFκB reporter activity); (ii) A host-pathogen entry/replication phenotype (viral/bacterial toxin entry); (iii) A secretion-based assay (secreted reporter/cytokine output).” We next asked ChatGPT to map the iGEA network into modules corresponding to the HIV life cycle. The model generated eight modules mapped to five life cycle stages (Supplementary Figure 4).

We next attempted to interpret ORA results using the same workflow limited to the 3 Gene Ontology collections. ORA-derived gene-term networks were difficult to provide directly to ChatGPT due to their size. The most practical format was a JSON list of enriched terms and their overlapping genes. ChatGPT summarized the ORA results as follows: “Your ORA reads like two tightly connected modules: (i) a dominant early secretory/Golgi trafficking axis (intra-Golgi, ER<->Golgi, and retrograde Golgi->ER) and (ii) a nucleocytoplasmic transport/mRNA-export axis centered on the NUP107-160 outer-ring complex. A third, supporting layer involves protein-complex disassembly/autophagy and glyco(lipid) biosynthesis, consistent with organelle remodeling and membrane reconfiguration under secretory stress.” When asked to propose the experimental context, ChatGPT concluded: “The data best support a Golgi/early secretory pathway remodeling program tightly coupled to boosted nuclear export of mRNA, with autophagy-mediated cleanup and membrane/glycolipid reconfiguration as enabling processes.” The full response is shown in Supplementary Table 8.

### Random gene lists and LLM interpretation

To illustrate the need to separate statistical evidence from LLM-generated narratives, we generated three random gene lists of size 300 and analyzed them with iGEA using the same 10 collections. The largest clusters were extracted from these networks and are visualized in Supplementary Figure 5A-C. All random networks were significantly smaller and less connected than the HIV largest cluster (Supplementary Figure 5D). Despite the relatively small size and low density of these random networks, ChatGPT generated plausible hypotheses and experimental context. For example, for Supplementary Figure 5A it proposed: “Your 300-gene list likely reflects a population (or condition) undergoing cytokine-driven immune activation (IL-2/STAT5 and broader interleukin signaling).” (Supplementary Table 12). We further tested random enrichment with respect to gene list size and found that iGEA clusters increase in size as the gene list size grows but decrease in connectivity (Supplementary Figure 6).

For comparison, we performed functional clustering using DAVID across six collections (KEGG, Reactome, WikiPathways, and the three Gene Ontology collections) (Supplementary Table 9). This analysis identified 94 clusters comprising 834 terms. Applying FDR < 0.05 reduced the output to 108 terms. The resulting report spanned 38 pages and remained difficult to synthesize into a coherent hypothesis.

We further evaluated iGEA combined with ChatGPT summarization on additional datasets: (1) a 75-gene hypoxia response signature [20], and (2) a 426-gene transcriptional signature associated with antibody titers following influenza vaccination[21]. In both cases, iGEA networks showed significant enrichment (z-scores > 3 across metrics of genes and edges). ChatGPT proposed experimental contexts that were consistent with the underlying studies. For hypoxia, it stated: “Your gene list reflects a hypoxia-driven adaptive remodeling state in which HIF signaling induces a glycolytic shift and simultaneously activates an extracellular matrix maturation/remodeling program.” (Supplementary Table 10). For the vaccination dataset, ChatGPT identified the signature as most likely reflecting bulk RNA-seq from an inflamed, immune-cell-rich tissue driven by activated myeloid/microglial cells and/or neutrophil infiltration, as seen in infections, IBD-like inflammation, neuroinflammation, immune-complex inflammation, or strong bacterial/cytokine stimulation (Supplementary Table 11). Follow-up queries further illustrated how structured conversational exploration can help users examine network-derived hypotheses. When asked what was inconsistent between the vaccination dataset and the iGEA network, ChatGPT noted that neutrophil degranulation terms may be unexpected in PBMC samples, but suggested that similar mechanisms in monocytes could explain the observation.

## Discussion

For more than two decades, gene set over-representation analysis (ORA) has been a central tool for interpreting molecular datasets. Its purpose is twofold: to recover known biology as a positive control and to generate new, testable hypotheses. We revisited a well-characterized HIV siRNA screen [13] because the HIV dependency-factor literature provides a useful pathway-level benchmark: several independent screens produced poorly overlapping gene lists, whereas subsequent meta-analyses showed that reproducible biological signals could be recovered at the pathway and process level [16, 17]. iGEA is not intended to solve cross-study reproducibility directly. Rather, we used a single HIV screen to ask whether iGEA could recover pathway-level themes supported by multiple studies and meta-analyses.

In common practice, ORA results are interpreted by assembling biological hypotheses from enriched term labels. This approach is feasible when the number of terms is limited and when deep domain expertise is available. However, as gene-set collections have expanded, interpretation has increasingly become a challenge. Redundancy reduction methods can help organize enriched terms, but building hypotheses by browsing clustered labels remains challenging. In the HIV dataset presented here, ER-Golgi trafficking emerged as the dominant signal across collections and methods. This is expected: HIV induces broad reconfiguration of host protein maturation, trafficking, and membrane composition [22]. The challenge is not identifying this primary signal but uncovering the subtler processes beneath it that may reveal new biology.

iGEA addresses this challenge by enforcing a one-gene one-term constraint within each gene-set collection. The collection-specific outputs are then integrated into a cross-collection gene-term network, where genes may connect to multiple terms originating from different collections. Because gene-set collections are curated from different biological perspectives, cross-collection connectivity captures multiple aspects of the same biological system. The resulting network provides a structured overview of the processes represented in the gene list. The value of iGEA is greatest when multiple complementary gene-set collections are integrated, because cross-collection connections provide the network structure needed for module and hub-based interpretation. However, as networks become denser, interpreting their topology visually becomes cognitively demanding.

Large language models, however, are not statistical engines. When applied to random gene lists, ChatGPT generated biological narratives despite weaker network structure. As expected, larger random gene lists are more likely to produce apparent enrichment by chance. These observations underscore the importance of network-level statistical benchmarking prior to LLM-assisted exploration. They also suggest a useful practice: in the initial prompt, users should provide the network structure but withhold experimental context, allowing the model to first interpret the network without being guided by prior assumptions. Network statistics establish whether the observed structure exceeds random expectation, while context-blinded prompting helps assess whether the network itself contains recognizable biological signal. LLMs should therefore be used to explore and summarize statistically supported network structure, rather than as independent evidence of biological correctness.

Conversational AI tools can be useful at this stage as exploratory interfaces. In our initial prompt, ChatGPT was given the structured iGEA network, but not the experimental context or how the gene list was generated. The prompt asked the model to infer a possible context and to support its interpretation by summarizing modules and hub genes, allowing users to evaluate the response against specific network features. Together with network-level statistics showing that the observed structure exceeded random expectation, this context-blinded analysis helps anchor the LLM response in network-derived biological signal, although it does not guarantee the correctness of the LLM-generated responses.

Using only the three Gene Ontology collections for visualization clarity, ChatGPT proposed a viral infection context despite the dominance of ER-Golgi-related terms and the absence of explicit HIV annotations. It also highlighted immune antiviral components driven by NF-κB, which were not emphasized in FDR-filtered ORA outputs. When expanded to all 11 collections, the model generated higher-resolution modules and mapped them to stages of the HIV life cycle. This illustrates how iGEA networks can support structured exploration of known biology.

An alternative strategy is to construct protein interaction networks and annotate dense clusters using ORA. This approach was used in prior HIV meta-analyses [16, 17], which concluded that pathway-level overlap across screens was robust despite limited gene-level reproducibility. Notably, the five clusters identified by ChatGPT from the iGEA network corresponded closely to themes identified in those meta-analyses, suggesting that iGEA can recover HIV pathway-level themes from a single screen.

We did not attempt to benchmark the correctness, consistency, or accuracy of LLM-generated interpretations. The examples of HIV, hypoxia and vaccination datasets demonstrate how iGEA networks can be used as structured inputs for conversational AI systems, not that LLM interpretation is superior to alternative approaches. The studies presented here were previously reported in the scientific literature. Therefore, memorization of biological context by modern LLMs cannot be excluded as a contributing factor. Future studies using blinded datasets and formal benchmarking frameworks will be required to assess the correctness, consistency, and utility of LLM-assisted interpretation.

Because empirical benchmarking is computationally intensive, iGEA uses precomputed random gene-list distributions for the default benchmarking collections and parameter settings. These benchmarks allow users to assess whether the largest cross-collection network component exceeds random expectation before exploring alternative parameter settings. For users requiring parameter-specific benchmarking, the provided code allows re-computation of random gene-list statistics under customized settings.

Beyond the HIV example, iGEA generated significant network structures across gene lists of different sizes, including a compact hypoxia signature and a larger vaccination-associated transcriptional signature. In both cases, network-level statistics supported non-random organization, and the resulting networks could be summarized into biologically coherent hypotheses. However, because the likelihood of random enrichment increases with gene-list size, we recommend limiting input lists to fewer than ∼800 genes to maintain robust statistical discrimination.

Several limitations remain. iGEA is designed for unranked gene lists and is not intended for ranked gene lists or other quantitative gene-level inputs such as fold changes, expression values, effect sizes, or perturbation scores. Other methods such as GSEA can address ranked gene lists and are not directly comparable [3]. The one-term per-gene per-collection constraint simplifies interpretation but may partition genes that legitimately participate in multiple processes. As with all enrichment analyses, results depend on collection selection, parameter thresholds, and background definition. Computational cost is also higher than standard ORA, as each iteration requires a separate enrichment calculation.

iGEA also depends on cross-collection integration for much of its interpretive value. With only one or a few collections, the method primarily reduces within-collection redundancy and may provide limited network-level context. As more collections are added, genes assigned to terms in different collections create a more informative network. This was illustrated in the HIV analysis: the GO-only network contained 156 genes and 29 terms and supported a module-level summary, whereas the 10-collection network expanded the representation to 179 genes and 62 terms and added pathway and signaling context, including receptor trafficking, glycosylation, RNA decay, and inflammatory signaling. This contrasts with conventional ORA, where adding collections often increases interpretive burden. We therefore recommend applying iGEA to multiple complementary collections when the goal is cross-collection interpretation, while recognizing that collection choice can influence the resulting network.

## Conclusions

For years, enrichment analysis has left researchers with the most difficult step: assembling overlapping term labels into a coherent biological hypothesis. Redundancy-reduction tools improved organization but did not resolve the interpretive burden, and network visualizations alone are often too dense for direct synthesis. LLMs are powerful summarizers, yet without structured and statistically supported inputs they may generate plausible but weakly supported narratives. iGEA bridges this gap by coupling a cross-collection network representation with empirical network-level statistics, providing a structured foundation for LLM-assisted exploration. In this framework, network-level statistics establish whether non-random biological organization is present, while conversational AI can assist users in exploring and summarizing the resulting network structure.

## Supporting information

Supplementary File - Peer Review

Supplementary File - Previous manuscript versions

## Acknowledgements

We thank Code Ocean for their support. The original Streamlit code was adopted from Code Ocean and written by MK while at Code Ocean. The iGEA method was added with support of AION Labs. Features and UI improvements and tests were added by YB using Cursor (www.cursor.com). Code Ocean also supported this study by providing computing infrastructure and by hosting the application on codeocean.com.

## Availability and requirements

- Project name: iterative Gene Enrichment Analysis
- Project source code: https://github.com/aion-labs/Gene-Enrichment-Analysis
- Version used in this manuscript: v1.0.0, https://github.com/aion-labs/Gene-Enrichment-Analysis/releases/tag/v1.0.0
- Project web server: https://iterative-gene-enrichment.streamlit.app
- Web server app source code: https://github.com/ybenita/iterative-gene-enrichment-app
- Operating system(s): Platform independent
- Programming language: Python
- Other requirements: Python 3.12+
- License: GPL-3.0 license
- Restrictions for non-academics: None

The public web application is deployed using Streamlit Community Cloud, which imposes memory and runtime constraints. To ensure stable hosting and responsive performance, we generated a streamlined version of iGEA optimized for web deployment. The hosted application therefore includes a curated subset of 12 collections and does not expose certain advanced functionalities (e.g., adding custom collections or recomputing empirical null distributions from new random gene lists). The full-featured version of iGEA, including support for additional GMT collections and regeneration of random benchmarking distributions under custom parameters, is available in the official GitHub repository.

## Declarations

- Consent to Publish: Not applicable.
- Consent to Participate: Not applicable.
- Data Availability: All data generated or analyzed during this study are included in this published article and its supplementary information files. Source code is openly available at https://github.com/aion-labs/Gene-Enrichment-Analysis. A web server implementation is available at https://iterative-gene-enrichment.streamlit.app.
- Author Contributions: MK implemented the initial code and wrote the software. YB developed the iterative GEA method, added features and tests, and conducted the analyses. Both authors contributed to the manuscript and approved the final version.
- Ethics Approval: Not applicable.
- Competing Interests: Y.B. is an employee of AION Labs. M.K. is a paid software developer. AION Labs provided funding for this work. The authors declare no additional competing interests.

