## Supplementary Figures 1-6 for "Iterative Gene Enrichment Analysis: interpretable networks for human and AI-assisted biological insights"

Maxim V. Kuleshov, Yair Benita — AION Labs, Rehovot, Israel

#### Contents

**Supplementary Figures 1–6** — legends below, figures following.

**Supplementary Tables** included in this document:

| | |
|---|---|
| Supplementary Table 1 | iGEA pseudocode |
| Supplementary Table 2 | Standardized LLM prompt |
| Supplementary Table 5 | Network statistical report |
| Supplementary Table 6 | iGEA AI response, Gene Ontology network |
| Supplementary Table 7 | iGEA AI response, full network |
| Supplementary Table 8 | ORA AI response, Gene Ontology network |
| Supplementary Table 10 | Hypoxia signature AI response |
| Supplementary Table 11 | Vaccination signature AI response |
| Supplementary Table 12 | Random gene list AI response |

**Provided as separate files**, because they are machine-readable data or already paginated:

| | |
|---|---|
| Supplementary Table 3 | ORA results (data table) |
| Supplementary Table 4 | iGEA results (data table) |
| Supplementary Table 9 | DAVID functional annotation cluster report |

### Supplementary figure legends

**Supplementary Figure 1. Terms identified by iGEA and ORA in each gene-set collection.** Number of unique enriched terms identified for the HIV dependency factor gene list across the 10 gene-set collections, shown as stacked bars. Blue, terms identified by both methods; teal, terms identified by iGEA only; orange, terms identified by ORA only. **(A)** Both methods evaluated at a raw p-value threshold of $p < 0.01$. **(B)** ORA terms filtered at FDR $q < 0.05$ and iGEA terms at $p < 0.01$.

**Supplementary Figure 2. Complete cross-collection iGEA network for the HIV dependency factor gene list.** The full iGEA gene–term network generated from the 272 mapped HIV dependency factors across all 10 gene-set collections. Term nodes are coloured by source collection, as keyed in the figure; genes are shown as labelled nodes. Edges connect each gene to the term it was assigned to by iGEA within a given collection. Network generated using $p < 0.01$. The reduced Gene Ontology view shown in Figure 2 of the main text is a subset of this network.

**Supplementary Figure 3. Gene Ontology iGEA network with full gene names.** The Gene Ontology-only iGEA network shown in Figure 2 of the main text, displayed with all gene names visible. Terms are coloured by collection: GO Biological Process (red), GO Molecular Function (purple), GO Cellular Component (blue). Genes from the input list overlapping with a term are shown as labelled nodes. Network generated using $p < 0.01$.

**Supplementary Figure 4. LLM-generated mapping of the iGEA network onto the HIV replication cycle.** Output of a follow-up query in which ChatGPT was asked to organise the cross-collection iGEA network into modules corresponding to stages of the HIV life cycle. The model generated eight modules (A–H), each listing its supporting enriched terms and member genes, and assigned them to five stages: (1) entry/fusion, (2) cytoplasmic trafficking and uncoating, (3) nuclear import and integration readiness, (4) replication and gene expression, and (5) assembly and secretion. Dashed arrows indicate progression through the replication cycle; solid coloured lines indicate the module-to-stage assignments proposed by the model.

**Supplementary Figure 5. Largest connected clusters from random gene lists compared with the HIV dependency factor network. (A–C)** Largest connected clusters obtained by applying iGEA to three independent random gene lists of 300 genes each, using the same 10 gene-set collections and parameters as the HIV analysis. **(D)** Largest connected cluster obtained from the 272 mapped HIV dependency factors under identical settings — the network shown in Supplementary Figure 2, reproduced here at matched scale so that the random and HIV networks can be compared directly. Term nodes are coloured by source collection, as keyed in the figure. The full response for the network in panel A is provided in Supplementary Table 12.

**Supplementary Figure 6. Effect of input gene list size on iGEA cluster size and density.** Distribution of largest-cluster size (upper row) and largest-cluster density (lower row) across random gene lists of increasing size, at three iterative significance thresholds (columns: $p < 0.001$, $p < 0.01$, $p < 0.05$). Each box shows the interquartile range with median; whiskers extend to the furthest values within range; individual horizontal marks show single random runs. A representative subset of the gene list sizes used in the full benchmark is shown here for legibility.

**Supplementary Figure 1.** Terms identified by iGEA and ORA in each gene-set collection

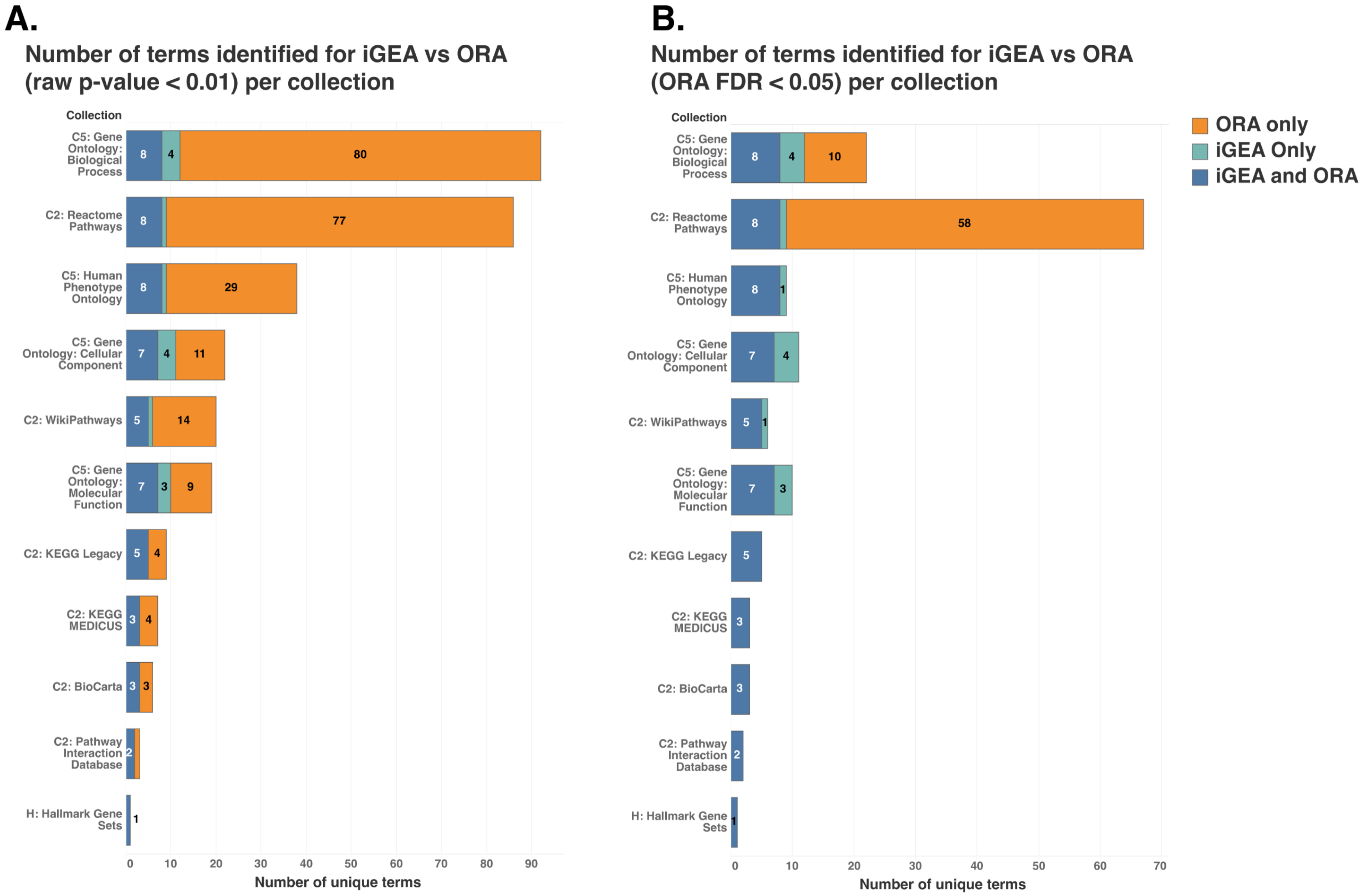

**Supplementary Figure 2.** Complete cross-collection iGEA network for the HIV dependency factor gene list

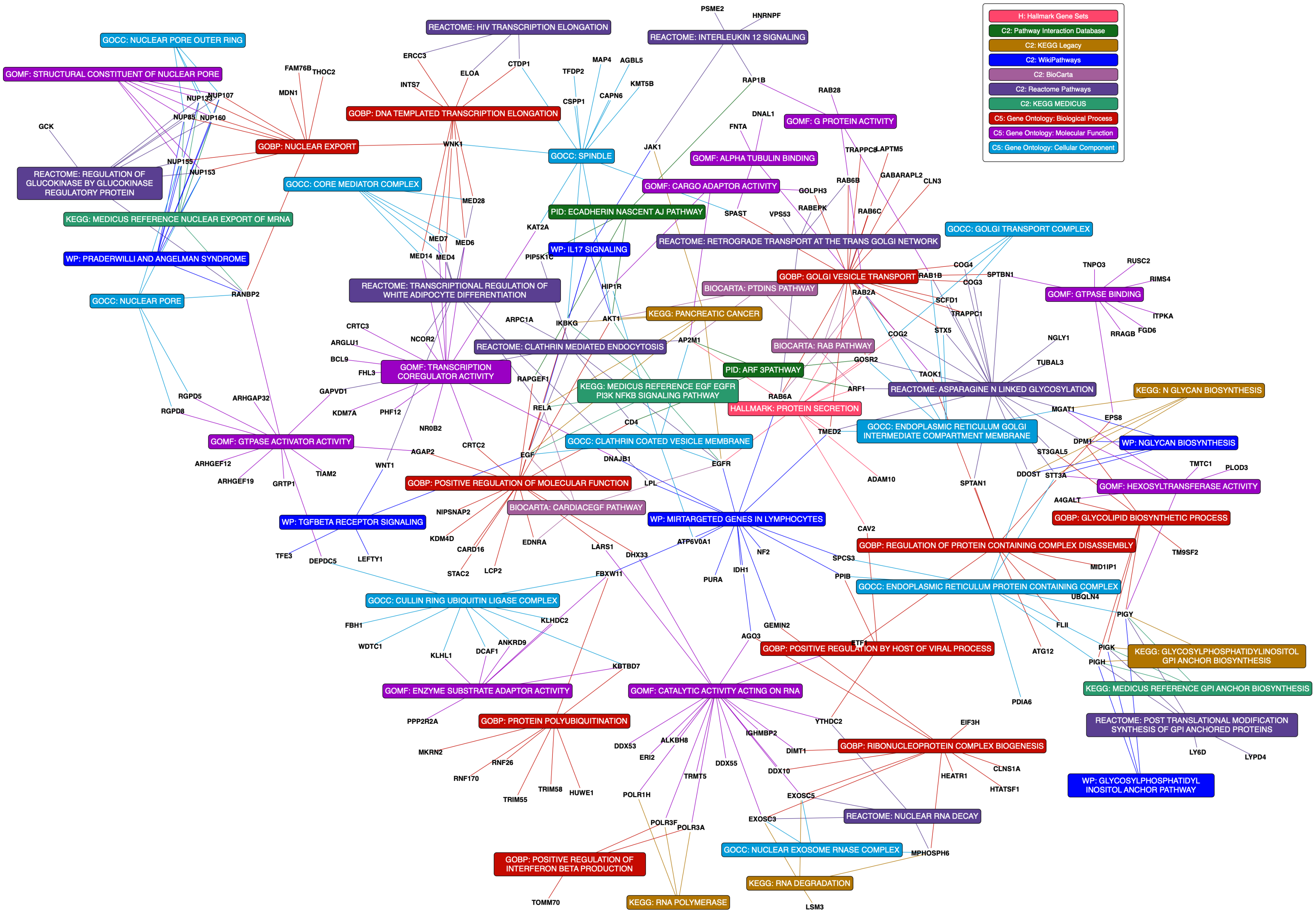

**Supplementary Figure 3.** Gene Ontology iGEA network with full gene names

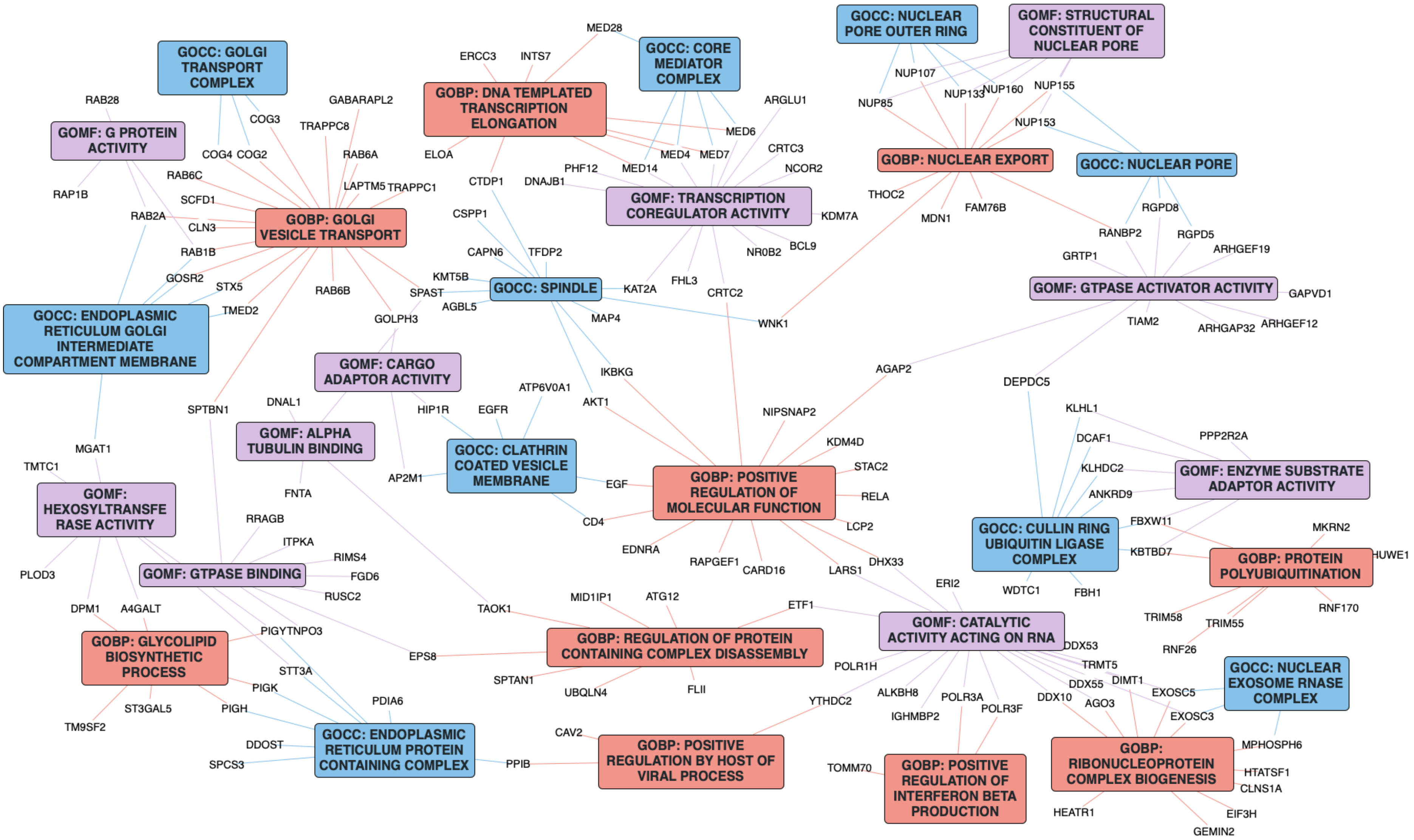

**Supplementary Figure 4.** LLM-generated mapping of the iGEA network onto the HIV replication cycle

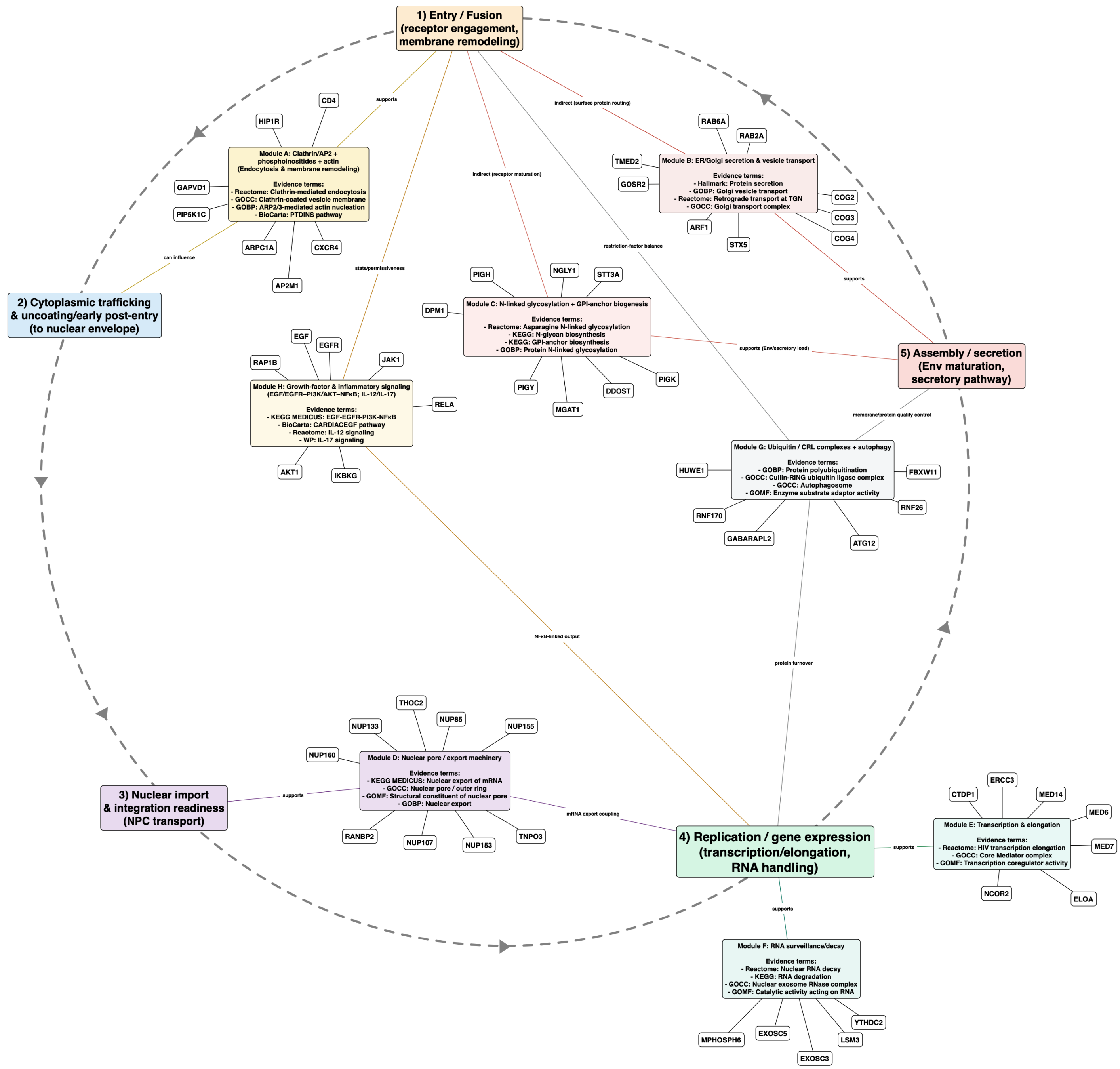

**Supplementary Figure 5.** Largest connected clusters from random gene lists compared with the HIV dependency factor network

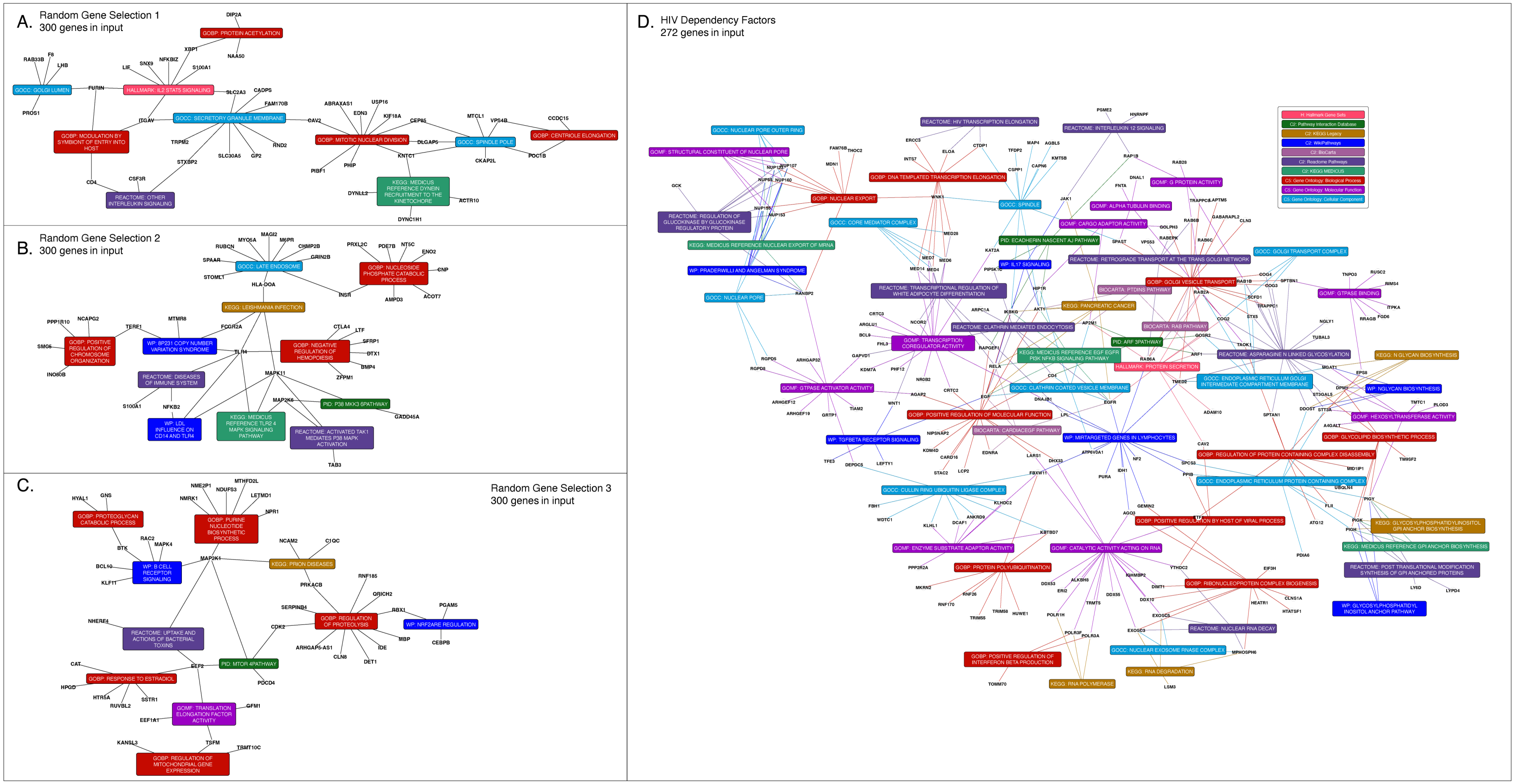

**Supplementary Figure 6.** Effect of input gene list size on iGEA cluster size and density

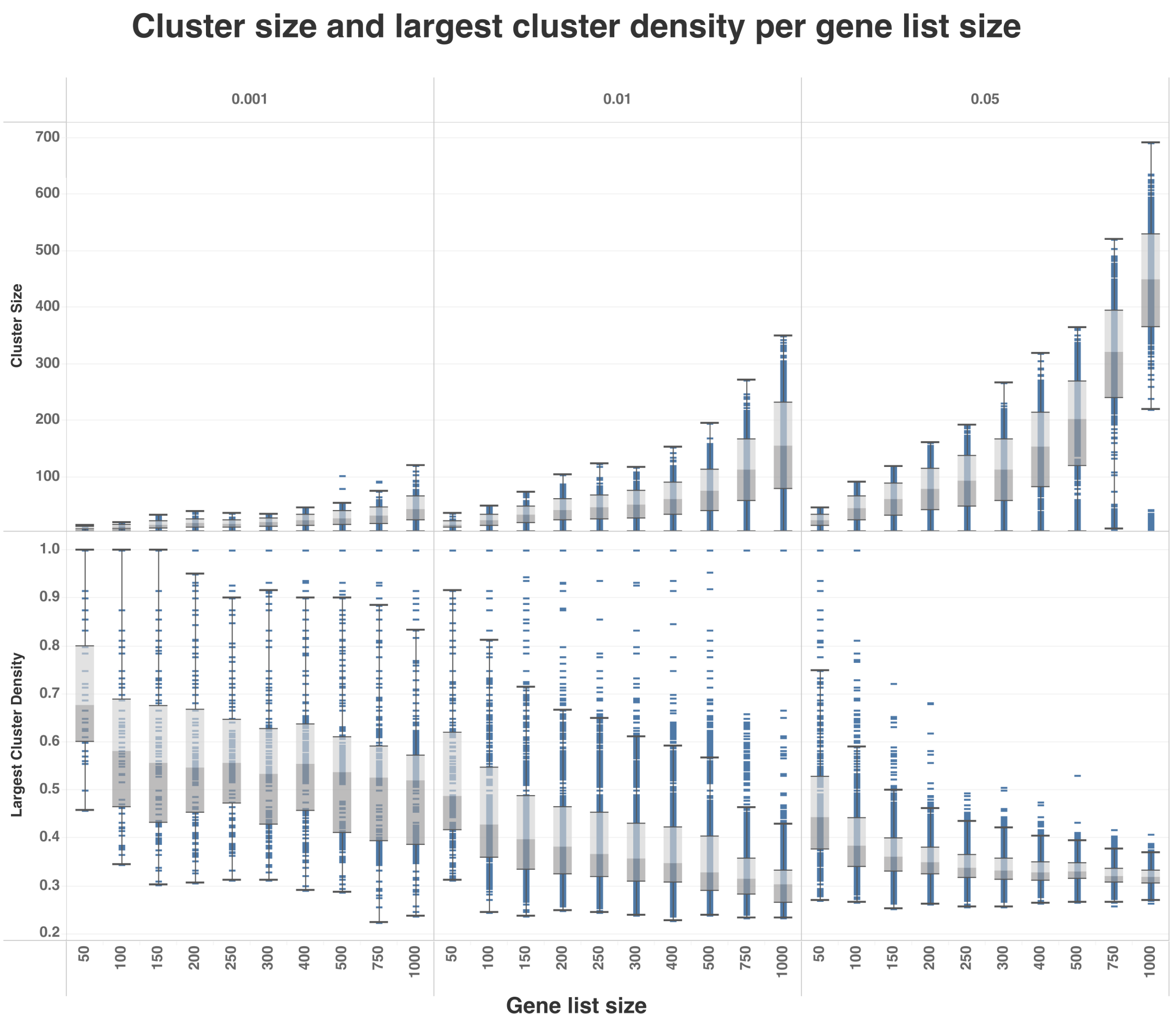

**Supplementary Table 1.** iGEA pseudocode

~~~
Iterative Gene Enrichment Analysis (iGEA) Algorithm

Algorithm Overview
The iterative enrichment algorithm repeatedly identifies the most significant enriched term, removes its
overlapping genes from the input gene set, and repeats the enrichment analysis on the remaining genes. This
process continues until termination criteria are met.

Algorithm Pseudocode
```
ALGORITHM: Iterative Gene Enrichment Analysis

INPUT:
    - Background_gene_set is selected (B)
    - input genes (G)
    - Libraries: list of libraries (L)
    - p_threshold: P-value cutoff to be used as stopping criteria (default: 0.01)
    - min_overlap: Minimum gene overlap between input (G) and gene set term (T)(default: 3)
    - max_gene_set_size: The maximum number of genes allowed within a term (default:600)
    - Max_iterations: Maximum iterations allowed (default: 10)


INITIALIZE:
  results ← []

For Library in L:
    WHILE (genes remaining > 0 and iteration <= max_iteration):
        remaining_genes ← gene_set.genes
        iteration ← 1

      // Create gene set from remaining genes
      current_gene_set ← GeneSet(remaining_genes, background_gene_set)

      // Perform enrichment analysis
      enrichment ← Enrichment(
        gene_set=current_gene_set,
        gene_set_library=gene_set_library,
        background_gene_set=background_gene_set
      )

      // Get enrichment results (sorted by p-value, ascending)
      results_list ← enrichment.results

      // Filter results by minimum overlap requirement
      filtered_results ← FILTER results_list WHERE overlap_size >= min_overlap and max_gene_set_size <
term_gene_size

      // Select top-ranked term (lowest p-value)
      top_term ← filtered_results[0]

      IF top_pvalue >= p_threshold:
        BREAK (exit loop)

      // Record iteration result
      record ← {
        Iteration: iteration,
        Term: top_term["term"],
        "iteration p-value": top_pvalue,
        "iteration overlapping genes": top_term["overlap_size"],
        "Genes removed for next iteration": top_overlap_genes
      }
      results.APPEND(record)

      // Remove overlapping genes from remaining set
      remaining_genes ← remaining_genes - top_overlap_genes

      // Increment iteration counter
      iteration ← iteration + 1

    END WHILE

END For
~~~

**Supplementary Table 2.** Standardized LLM prompt

```
You are a computational biologist analyzing an iterative gene set enrichment network. Iterative means that for
each library source, the top enriched gene set is found, saved and the genes are removed from the initial gene
list. The remaining genes are then tested for enrichment again. Thus, each gene appears only once per library
source tested but can be linked to multiple terms that originate from different libraries. The results are
provided as a DOT network represents the relationships between genes and gene sets(=terms).

**NETWORK STRUCTURE:**
[Insert DOT Network]

**NETWORK INTERPRETATION GUIDE:**

**Node Types:**
- **Gene nodes** (type="gene"): Individual genes from the input gene set
- **Term nodes** (type="term"): Enriched biological pathways/processes/gene sets
- **Library Source**: Each term node includes a "library" attribute indicating the source database

**Library Sources and Their Biological Context:**
- **H: Hallmark gene sets**: Curated gene sets representing well-defined biological states or processes
- **C2: BioCarta**: Canonical pathways from BioCarta database
- **C2: KEGG MEDICUS**: Metabolic and signaling pathways from KEGG
- **C2: Pathway Interaction Database**: Curated human signaling pathways
- **C2: Reactome Pathways**: Expert-curated biological pathways
- **C2: WikiPathways**: Community-curated biological pathways
- **C5: Gene Ontology: Biological Process**: Biological processes from Gene Ontology
- **C5: Gene Ontology: Cellular Component**: Cellular components from Gene Ontology
- **C5: Gene Ontology: Molecular Function**: Molecular functions from Gene Ontology
- **C5: Human Phenotype Ontology**: Human phenotypes and diseases
- **Protein Interaction**: Protein-protein interaction networks

**Edge Connections:**
- Each edge (gene -- term) represents a gene's membership in that biological term
- More connections indicate genes that are part of multiple enriched processes
- The network topology shows which genes are central to the biological response

**Iteration Information:**
- Term nodes are labeled with iteration numbers (term_1_, term_2_, etc.)
- Earlier iterations (lower numbers) represent the most statistically significant enrichments
- Ignore iteration number for biological interpretation, instead focus on network hubs densely connected for
interesting biological insights

**ANALYSIS REQUEST:**

Please analyze this network and provide:

1. **Key Biological Insights:**
   - What are the most significant biological processes/pathways identified?
   - Which genes appear to be central to the biological response?

2. **Network Topology Analysis:**
   - Which genes are "hub" genes (connected to multiple terms)?
   - Are there distinct clusters or modules in the network?
   - What does the connectivity pattern suggest about biological organization?

3. **Biological Hypothesis:**
   - Based on the network structure, what biological hypothesis can you generate?
   - How do the different library sources support or complement each other?
   - Try to formulate one hypothesis based on the most central terms and how they possibly related to one
another.

4. **Estimated Experimental Context**
   - Can you hypothesize on what was the experiment that generated this result?

**RESPONSE STRUCTURE:**
- **Executive Summary** (2-3 sentences)
- **Key Biological Processes Identified** (by library source)
- **Network Topology Insights**
- **Proposed Biological Hypothesis**
- **Estimated Experimental Context**

**IMPORTANT NOTES:**
- Focus on biological interpretation, not statistical significance or iteration number
```

- Consider the functional relationships between connected genes and terms
- Look for unexpected connections that might reveal novel or unexpected biology
- Consider the broader biological context and literature
- Note that protein interaction library could shed light on physical interaction between genes while terms indicate functional interaction
- Different library sources may provide complementary biological insights

**Supplementary Table 5.** Network statistical report

```
==================================================================================================
CLUSTER-BY-CLUSTER STATISTICAL REPORT
==================================================================================================

Gene List: HIV_dependency_factors.symbols
Gene List Size: 272
Actual Size Used for Null Distribution: 300 (rounded from 272)
Total Clusters in Network: 1
Note: Only the largest cluster is benchmarked (null distribution built from largest clusters only)

==================================================================================================
IMPORTANT: Library Information for Statistical Analysis
==================================================================================================

Statistics were computed using 10 libraries with permutation data:
  H: Hallmark Gene Sets, C2: CP: BioCarta, C2: CP: KEGG MEDICUS, C2: CP: Pathway Interaction Database, C2: CP:
Reactome Pathways, C2: CP: WikiPathways, C2: CP: KEGG Legacy, C5: GO: Biological Process, C5: GO: Cellular
Component, C5: GO: Molecular Function

The full network visualization includes all selected libraries.
However, statistical benchmarks are only computed for libraries with available
permutation data to ensure accurate comparison against the null distribution.

==================================================================================================

This report shows the largest cluster with benchmark statistics comparing
against random gene lists of similar size.
Note: Only the largest cluster is benchmarked, as the null distribution
is built from the largest cluster in each permutation.

Interpretation:
  • Z-score > 2.0 and Percentile > 95%: Significantly better than random (top 5%)
  • Z-score > 1.0 and Percentile > 84%: Better than random (top 16%)
  • Z-score ~ 0.0 and Percentile ~ 50%: Similar to random
  • Z-score < -1.0 and Percentile < 16%: Worse than random

==================================================================================================

--------------------------------------------------------------------------------------------------
LARGEST CLUSTER
--------------------------------------------------------------------------------------------------

Basic Cluster Metrics:
  Cluster Size:       241 nodes (genes + terms)
  Number of Genes:    179
  Number of Terms:    62
  Number of Edges:    412
  Average Edges per Gene: 2.3017
  Number of Libraries: 10

Statistical Benchmarks vs Random Gene Lists:
  Metric                   Value     Null Mean   Null Std    Z-score   Percentile   Status
------------------------------------------------------------------------------------------------------------
  Cluster Size                241       37.95       26.22       7.74     100.0%   SIGNIFICANTLY BETTER
  Number of Genes             179       29.38       20.24       7.39     100.0%   SIGNIFICANTLY BETTER
  Number of Terms              62        8.57        6.33       8.44     100.0%   SIGNIFICANTLY BETTER
  Number of Edges             412       43.51       32.61      11.30     100.0%   SIGNIFICANTLY BETTER
  Average Edges per Gene   2.3017      1.4235      0.2929       3.00      99.9%   SIGNIFICANTLY BETTER
  Number of Libraries          10        4.56        2.25       2.42      99.2%   SIGNIFICANTLY BETTER

Terms in Cluster (62 total, ranked by centrality):
  Rank  Term                                                        Centrality (genes)
  ------------------------------------------------------------------------------------------
   1.   C5: Gene Ontology: Biological Process:Gene Ontology: Biological Process: GOBP: GOLGI VESICLE TRANSPORT
20
   2.   C2: Reactome Pathways:Reactome Pathways: REACTOME: ASPARAGINE N LINKED GLYCOSYLATION  19
   3.   C5: Gene Ontology: Molecular Function:Gene Ontology: Molecular Function: GOMF: CATALYTIC ACTIVITY
ACTING ON RNA  18
   4.   C5: Gene Ontology: Biological Process:Gene Ontology: Biological Process: GOBP: POSITIVE REGULATION OF
MOLECULAR FUNCTION  16
   5.   C5: Gene Ontology: Molecular Function:Gene Ontology: Molecular Function: GOMF: TRANSCRIPTION
```

COREGULATOR ACTIVITY 15

6. C2: WikiPathways:WikiPathways: WP: MIRTARGETED GENES IN LYMPHOCYTES 14
7. C5: Gene Ontology: Cellular Component:Gene Ontology: Cellular Component: GOCC: SPINDLE 12
8. C5: Gene Ontology: Biological Process:Gene Ontology: Biological Process: GOBP: NUCLEAR EXPORT 11
9. C5: Gene Ontology: Biological Process:Gene Ontology: Biological Process: GOBP: RIBONUCLEOPROTEIN COMPLEX BIOGENESIS 11
10. C5: Gene Ontology: Molecular Function:Gene Ontology: Molecular Function: GOMF: GTPASE ACTIVATOR ACTIVITY 11
11. H: Hallmark Gene Sets:Hallmark Gene Sets: HALLMARK: PROTEIN SECRETION 9
12. C2: Reactome Pathways:Reactome Pathways: REACTOME: TRANSCRIPTIONAL REGULATION OF WHITE ADIPOCYTE DIFFERENTIATION 9
13. C5: Gene Ontology: Cellular Component:Gene Ontology: Cellular Component: GOCC: CULLIN RING UBIQUITIN LIGASE COMPLEX 9
14. C5: Gene Ontology: Biological Process:Gene Ontology: Biological Process: GOBP: DNA TEMPLATED TRANSCRIPTION ELONGATION 9
15. C2: Reactome Pathways:Reactome Pathways: REACTOME: CLATHRIN MEDIATED ENDOCYTOSIS 8
16. C2: Reactome Pathways:Reactome Pathways: REACTOME: REGULATION OF GLUCOKINASE BY GLUCOKINASE REGULATORY PROTEIN 8
17. C5: Gene Ontology: Cellular Component:Gene Ontology: Cellular Component: GOCC: ENDOPLASMIC RETICULUM PROTEIN CONTAINING COMPLEX 8
18. C5: Gene Ontology: Biological Process:Gene Ontology: Biological Process: GOBP: REGULATION OF PROTEIN CONTAINING COMPLEX DISASSEMBLY 8
19. C5: Gene Ontology: Molecular Function:Gene Ontology: Molecular Function: GOMF: GTPASE BINDING 8
20. C5: Gene Ontology: Biological Process:Gene Ontology: Biological Process: GOBP: PROTEIN POLYUBIQUITINATION 8
21. C5: Gene Ontology: Molecular Function:Gene Ontology: Molecular Function: GOMF: HEXOSYLTRANSFERASE ACTIVITY 7
22. C5: Gene Ontology: Molecular Function:Gene Ontology: Molecular Function: GOMF: ENZYME SUBSTRATE ADAPTOR ACTIVITY 7
23. C5: Gene Ontology: Biological Process:Gene Ontology: Biological Process: GOBP: GLYCOLIPID BIOSYNTHETIC PROCESS 7
24. C2: WikiPathways:WikiPathways: WP: PRADERWILLI AND ANGELMAN SYNDROME 7
25. C2: KEGG Legacy:KEGG Legacy: KEGG: PANCREATIC CANCER 6
26. C5: Gene Ontology: Cellular Component:Gene Ontology: Cellular Component: GOCC: CLATHRIN COATED VESICLE MEMBRANE 6
27. C2: KEGG MEDICUS:KEGG MEDICUS: KEGG: MEDICUS REFERENCE NUCLEAR EXPORT OF MRNA 6
28. C5: Gene Ontology: Molecular Function:Gene Ontology: Molecular Function: GOMF: STRUCTURAL CONSTITUENT OF NUCLEAR PORE 6
29. C5: Gene Ontology: Cellular Component:Gene Ontology: Cellular Component: GOCC: ENDOPLASMIC RETICULUM GOLGI INTERMEDIATE COMPARTMENT MEMBRANE 6
30. C5: Gene Ontology: Cellular Component:Gene Ontology: Cellular Component: GOCC: AUTOPHAGOSOME 5

... and 32 more terms

====================================================================================================
END OF REPORT
====================================================================================================

## Supplementary Table 6

iGEA AI response, Gene Ontology network

Full ChatGPT response for the Gene Ontology-only iGEA network. The Executive Summary is quoted in the Results.

**Executive Summary**
This network resolves into five tightly connected modules: **ER–Golgi trafficking & glycosylation**, **nuclear pore/export & transcription/RNA processing**, **ubiquitin–proteasome/Cullin-RING ligases**, **small-GTPase/vesicle adaptors & endocytosis**, and an **innate-immune axis centered on POLR3A/F→TOMM70→IFN-β**. Hub genes bridging these modules (e.g., **NUPs, MED subunits, RAB1B/RAB2A, EXOSC3/5, FBXW11, SPAST, DPM1/STT3A/PIGY**) suggest a coordinated state of transcriptional reprogramming, nucleocytoplasmic transport, vesicle remodeling, proteostasis, and antiviral signaling.

**Key Biological Processes Identified (by library source)**
**C5: Gene Ontology – Biological Process (GOBP)**

- **Golgi vesicle transport**: COG2/3/4, GOSR2, STX5, TMED2, RAB1B/RAB2A/RAB6A/B/C, RAB6C, RAB6A/B, SCFD1, TRAPPC1/8, GOLPH3, LAPTM5, SPTBN1, SPAST, RAB6 family.
  ⇒ Core **ER–Golgi/ERGIC trafficking** machinery.
- **Nuclear export**: NUP107/133/153/155/160/85, RANBP2, WNK1, THOC2.
  ⇒ **Nuclear pore–mRNA export** axis.
- **DNA-templated transcription elongation**: MED14/4/6/7/28, ERCC3, CTDP1, ELOA, INTS7.
  ⇒ **Mediator/TFIIH/elongation** activities.
- **Glycolipid biosynthetic process**: A4GALT, DPM1, PIGH/PIGK/PIGY, ST3GAL5, TM9SF2, TMTC1.
  ⇒ **N-/O-glycosylation & GPI-anchor** biosynthesis.
- **Protein polyubiquitination**: HUWE1, FBXW11, MKRN2, RNF170/26, TRIM55/58.
  ⇒ **UPS/ERAD & signaling** regulation.
- **Ribonucleoprotein complex biogenesis**: EXOSC3/5, MPHOSPH6, GEMIN2, DDX10, DIMT1, EIF3H, HEATR1, HTATSF1, AGO3.
  ⇒ **RNP processing/exosome** functions.
- **Positive regulation of interferon-β production**: POLR3A, POLR3F, TOMM70.
  ⇒ **Pol III→RIG-I/MAVS** antiviral pathway.
- **Positive regulation by host of viral process**: CAV2, PPIB, YTHDC2.
  ⇒ Host factors that **viruses exploit** (entry/replication).
- **Regulation of protein-containing complex disassembly**: ATG12, FLII, ETF1, MID1IP1, SPTAN1, TAOK1, UBQLN4.
  ⇒ **Proteostasis & complex turnover**.
- **Positive regulation of molecular function** (broad signaling set): AKT1, IKBKG, RELA, RAPGEF1, CD4, EGF, EDNRA, etc.
  ⇒ **Signal transduction** tone.

**C5: Gene Ontology – Cellular Component (GOCC)**

- **Core Mediator complex**: MED14/4/6/7/28.
- **Nuclear pore / outer ring**: NUP107/133/160/85; **nuclear pore**: NUP153/155, RANBP2, RGPD5/8.
- **Nuclear exosome RNase complex**: EXOSC3/5, MPHOSPH6.
- **Clathrin-coated vesicle membrane**: AP2M1, HIP1R, EGFR, EGF, CD4, ATP6V0A1.
- **Golgi transport complex / ERGIC membrane**: COG2/3/4, STX5, GOSR2, TMED2, RAB1B/RAB2A, MGAT1.
- **ER protein-containing complex**: STT3A, DDOST, PDIA6, PIGH/PIGK/PIGY, PPIB, SPCS3.
- **Cullin-RING ubiquitin ligase complex**: FBXW11, KBTBD7, KLHDC2, KLHL1, DCAF1, DEPDC5, WDTC1.
- **Spindle**: AKT1, SPAST, MAP4, KMT5B, TFDP2, CSPP1, WNK1, CTDP1, AGBL5, CAPN6.

**C5: Gene Ontology – Molecular Function (GOMF)**

- **Transcription coregulator activity**: MED14/4/6/7/28, KAT2A, CRTC2/3, BCL9, NCOR2, NR0B2, PHF12.
- **Catalytic activity acting on RNA**: DDX/DHX helicases (DDX10/53/55, DHX33), TRMT5, ALKBH8, ERI2, LARS1, ETF1, POLR1H, POLR3A/F, IGHMBP2, AGO3.
- **Structural constituent of nuclear pore**: NUP107/133/153/155/160/85.
- **GTPase activator/binding & G-protein activity**:
  – **GAP/GEF**: AGAP2, ARHGAP32, ARHGEF12/19, DEPDC5, GAPVD1, GRTP1, RANBP2, RGPD5/8, TIAM2.
  – **Binding**: EPS8, FGD6, ITPKA, RIMS4, RRAGB, RUSC2, SPTBN1, TNPO3.
  – **Small GTPases**: RAB1B/RAB2A/RAB28, RAP1B.
- **Cargo adaptor activity**: AP2M1, HIP1R, GOLPH3.
- **Hexosyltransferase activity**: A4GALT, DPM1, MGAT1, PIGY, PLOD3, STT3A, TMTC1.

- **α-tubulin binding**: SPAST, TAOK1, FNTA, DNAL1.

---

**Network Topology Insights**

**Hub genes (multi-term, cross-library)**

- **Nuclear pore/export hubs**: **NUP107/133/160/85, NUP153/155, RANBP2** (BP: nuclear export; CC: pore/outer ring; MF: structural constituent/GAP).
- **Mediator/transcription hubs**: **MED14/MED4/MED6/MED7** (BP: elongation; CC: Mediator; MF: coregulator).
- **ER–Golgi trafficking hubs**: **RAB1B/RAB2A** (BP: Golgi transport; CC: ERGIC; MF: G-protein), **COG2/3/4**, **STX5**, **TMED2**, **GOSR2**.
- **RNA processing hubs**: **EXOSC3/EXOSC5/MPHOSPH6** (BP: RNP biogenesis; CC: exosome; MF: RNA catalysis).
- **UPS/CRL hubs**: **FBXW11, DCAF1, KBTBD7, KLHL1/KLHDC2** (BP: polyubiquitination; CC: CRL; MF: substrate adaptor).
- **Glycosylation/OST hubs**: **DPM1, STT3A, PIGY** (BP: glycolipid biosynthesis; CC: ER complex; MF: hexosyltransferase).
- **Bridging cytoskeleton–trafficking**: **SPAST** (BP: Golgi transport; CC: spindle; MF: α-tubulin binding).
- **Signaling/innate immunity bridges**: **POLR3A/POLR3F/TOMM70**, **IKBKG/RELA**, **EGF/EGFR** (with **AP2M1/HIP1R** for endocytosis).

**Modules/clusters**

1. **Secretory & glycosylation module**: COG complex, STX5/GOSR2/TMED2, RAB1B/2A, STT3A/DDOST/SPCS3, PIGH/PIGK/PIGY, MGAT1, DPM1, ST3GAL5/TMTC1 → **protein maturation, GPI-anchors, ERAD-linked quality control**.
2. **Nucleus/export & RNA biogenesis module**: NUPs/THOC2/RANBP2 ↔ Mediator/elongation ↔ EXOSC complex → **transcription–processing–export pipeline**.
3. **Ubiquitination/CRL module**: FBXW11/KBTBD7/KLHL1/KLHDC2/DCAF1/RNFs/TRIMs → **signal routing & proteostasis**.
4. **Small-GTPase & endocytosis module**: RABs, AGAP/ARHGAP/ARHGEF set, EPS8/FGD6/RUSC2, AP2M1/HIP1R/GOLPH3, clathrin membrane terms → **vesicle budding, cargo selection, receptor trafficking**.
5. **Antiviral/IFN module**: POLR3A/F & **TOMM70** (IFN-β), **IKBKG/RELA** (NF-κB), **CAV2/PPIB/YTHDC2** (host-proviral factors) → **tension between antiviral signaling and host-dependency factors**.

**Connectivity suggests** a coordinated response where **transcriptional upshift** (Mediator/elongation) feeds into **enhanced RNA processing/export**, while **secretory & glycosylation capacity** and **vesicle routing** scale to handle cytokines/receptors; **UPS/CRL** shapes signaling and clears misfolded load; **innate sensors** trigger IFN-β, and **endocytosis** turns over activated receptors or mediates pathogen entry.

---

**Proposed Biological Hypothesis**

**Hypothesis:** The gene set reflects **cells mounting an early antiviral/innate-immune response** (likely **Pol III–RIG-I/MAVS–IFN-β** activation) coupled to **extensive remodeling of the secretory pathway** and **proteostasis**, enabling cytokine production and surface receptor turnover.

- **Evidence lines:**
  - **IFN-β axis**: POLR3A/POLR3F (DNA→5′ppp-RNA sensing), TOMM70 (MAVS adaptor).
  - **NF-κB signaling tone**: IKBKG (NEMO) & RELA within broad "positive regulation of molecular function."
  - **Secretory scaling**: Strong **ER–Golgi/ERGIC** and **hexosyltransferase/OST** signatures (STT3A, DDOST, DPM1, PIGH/K/Y, MGAT1).
  - **Receptor/endocytic turnover**: AP2M1, HIP1R, EGFR/EGF, clathrin-coated vesicle membrane.
  - **Proteostasis & ERAD**: CRL adaptors (FBXW11, KBTBD7, KLHL1/KLHDC2, DCAF1), RNF/TRIM E3s, UBQLN4.
  - **Nucleus ↔ cytoplasm flow**: NUP outer ring/core + THO/TREX (THOC2) + exosome (EXOSC3/5, MPHOSPH6) + elongation (MEDs).
  - **Cross-module bridges**: **RANBP2** (nuclear pore & GAP activity), **SPAST** (microtubules–Golgi), **RAB1B/RAB2A** (ERGIC trafficking).

**How sources complement each other:**

- **GO:BP** contributes **process-level narratives** (transport, export, IFN-β, ubiquitination).
- **GO:CC** localizes those processes to **specific machines/compartments** (Mediator, nuclear pore outer ring, exosome, ERGIC, CRL, spindle).
- **GO:MF** ties genes to **biochemical roles** (coregulator, RNA helicase/methyltransferase, hexosyltransferase, GAP/GEF, cargo adaptor), clarifying **mechanistic levers** inside the processes.

**Estimated Experimental Context**

Most consistent with **transcriptomic (or hit-list) data from human cells exposed to a viral mimic or infection** that activates the **Pol III → RIG-I/MAVS → IFN-β pathway**—for example:

- **Cytosolic AT-rich DNA transfection (e.g., poly(dA:dT))** or **5′ppp-dsRNA** in epithelial or immune-derived cells,
  **or** an **RNA/DNA virus** that relies on clathrin entry and host glycosylation.

Expected timing: **early–mid response (∼4–12 h)** where **transcription/export and secretory capacity** are being re-tooled, **IFN-β induction** is evident, and **broad ISG saturation** is not yet dominant. The strong **ER–Golgi/glycosylation** and **CRL/UPS** signals also fit **heightened cytokine secretion and receptor remodeling** plus **quality-control/ERAD** during stress.

## Supplementary Table 7

iGEA AI response, full network

Full ChatGPT response for the complete cross-collection iGEA network.

[Response to initial prompt](#)
[Follow up on EGFR](#)
[Follow up to interpret in the context of HIV life cycle](#)
[Follow up on what is missing](#)

Structured initial prompt:
You are a computational biologist analyzing an iterative gene set enrichment network. Iterative means that for each library source, the top enriched gene set is found, saved and the genes are removed from the initial gene list. The remaining genes are then tested for enrichment again. Thus, each gene appears only once per library source tested but can be linked to multiple terms that originate from different libraries. The results are provided as a DOT network represents the relationships between genes and gene sets(=terms).

The original gene list included 272 genes and was tested across the following libraries: H: Hallmark Gene Sets, C2: BioCarta, C2: KEGG Legacy, C2: KEGG MEDICUS, C2: Pathway Interaction Database, C2: Reactome Pathways, C2: WikiPathways, C5: Gene Ontology: Biological Process, C5: Gene Ontology: Cellular Component, C5: Gene Ontology: Molecular Function, C5: Human Phenotype Ontology.

**NETWORK STRUCTURE:**
```
graph iterative_enrichment_all {
graph [layout=neato];
node [shape=ellipse];
subgraph cluster_legend_libs {
label="Libraries";
node [shape=box, style=filled, fontcolor="white"];
legend_lib_1 [label="H: Hallmark Gene Sets", fillcolor="#FF6B6B"];
legend_lib_2 [label="C2: BioCarta", fillcolor="#DDA0DD"];
legend_lib_3 [label="C2: KEGG Legacy", fillcolor="#F8C471"];
legend_lib_4 [label="C2: KEGG MEDICUS", fillcolor="#98D8C8"];
legend_lib_5 [label="C2: Pathway Interaction Database", fillcolor="#440154"];
legend_lib_6 [label="C2: Reactome Pathways", fillcolor="#BB8FCE"];
legend_lib_7 [label="C2: WikiPathways", fillcolor="#31688e"];
legend_lib_8 [label="C5: Gene Ontology: Biological Process", fillcolor="#F1948A"];
legend_lib_9 [label="C5: Gene Ontology: Cellular Component", fillcolor="#85C1E9"];
legend_lib_10 [label="C5: Gene Ontology: Molecular Function", fillcolor="#D7BDE2"];
legend_lib_11 [label="C5: Human Phenotype Ontology", fillcolor="#F7DC6F"];
}
"gene_ADAM10" [label="ADAM10", type="gene", width=0.7, height=0.7];
"gene_AP2M1" [label="AP2M1", type="gene", width=0.7, height=0.7];
"gene_ARF1" [label="ARF1", type="gene", width=0.7, height=0.7];
"gene_CAV2" [label="CAV2", type="gene", width=0.7, height=0.7];
"gene_COG2" [label="COG2", type="gene", width=0.7, height=0.7];
"gene_EGFR" [label="EGFR", type="gene", width=0.7, height=0.7];
"gene_GOSR2" [label="GOSR2", type="gene", width=0.7, height=0.7];
"gene_RAB2A" [label="RAB2A", type="gene", width=0.7, height=0.7];
"gene_TMED2" [label="TMED2", type="gene", width=0.7, height=0.7];
"term_1_HALLMARK_PROTEIN_SECRETION" [label="HALLMARK: PROTEIN SECRETION", style=filled, fillcolor="#FF6B6B", fontcolor="white", type="term", width=2, height=2, library="H: Hallmark Gene Sets"];
"gene_ADAM10" -- "term_1_HALLMARK_PROTEIN_SECRETION" [color="#FF6B6B"];
"gene_AP2M1" -- "term_1_HALLMARK_PROTEIN_SECRETION" [color="#FF6B6B"];
"gene_ARF1" -- "term_1_HALLMARK_PROTEIN_SECRETION" [color="#FF6B6B"];
"gene_CAV2" -- "term_1_HALLMARK_PROTEIN_SECRETION" [color="#FF6B6B"];
"gene_COG2" -- "term_1_HALLMARK_PROTEIN_SECRETION" [color="#FF6B6B"];
"gene_EGFR" -- "term_1_HALLMARK_PROTEIN_SECRETION" [color="#FF6B6B"];
"gene_GOSR2" -- "term_1_HALLMARK_PROTEIN_SECRETION" [color="#FF6B6B"];
"gene_RAB2A" -- "term_1_HALLMARK_PROTEIN_SECRETION" [color="#FF6B6B"];
"gene_TMED2" -- "term_1_HALLMARK_PROTEIN_SECRETION" [color="#FF6B6B"];
"gene_AKT1" [label="AKT1", type="gene", width=0.7, height=0.7];
"gene_EDNRA" [label="EDNRA", type="gene", width=0.7, height=0.7];
"gene_EGF" [label="EGF", type="gene", width=0.7, height=0.7];
"gene_RAB6A" [label="RAB6A", type="gene", width=0.7, height=0.7];
"gene_RAB6C" [label="RAB6C", type="gene", width=0.7, height=0.7];
"gene_RELA" [label="RELA", type="gene", width=0.7, height=0.7];
"term_1_BIOCARTA_CARDIACEGF_PATHWAY" [label="BIOCARTA: CARDIACEGF PATHWAY", style=filled, fillcolor="#DDA0DD", fontcolor="white", type="term", width=2, height=2, library="C2: BioCarta"];
"term_2_BIOCARTA_RAB_PATHWAY" [label="BIOCARTA: RAB PATHWAY", style=filled, fillcolor="#DDA0DD", fontcolor="white", type="term", width=2, height=2, library="C2: BioCarta"];
"term_3_BIOCARTA_PTDINS_PATHWAY" [label="BIOCARTA: PTDINS PATHWAY", style=filled, fillcolor="#DDA0DD", fontcolor="white", type="term", width=2, height=2, library="C2: BioCarta"];
"gene_AKT1" -- "term_3_BIOCARTA_PTDINS_PATHWAY" [color="#DDA0DD"];
"gene_AP2M1" -- "term_3_BIOCARTA_PTDINS_PATHWAY" [color="#DDA0DD"];
"gene_ARF1" -- "term_3_BIOCARTA_PTDINS_PATHWAY" [color="#DDA0DD"];
"gene_EDNRA" -- "term_1_BIOCARTA_CARDIACEGF_PATHWAY" [color="#DDA0DD"];
"gene_EGF" -- "term_1_BIOCARTA_CARDIACEGF_PATHWAY" [color="#DDA0DD"];
"gene_EGFR" -- "term_1_BIOCARTA_CARDIACEGF_PATHWAY" [color="#DDA0DD"];
"gene_RAB2A" -- "term_2_BIOCARTA_RAB_PATHWAY" [color="#DDA0DD"];
"gene_RAB6A" -- "term_2_BIOCARTA_RAB_PATHWAY" [color="#DDA0DD"];
"gene_RAB6C" -- "term_2_BIOCARTA_RAB_PATHWAY" [color="#DDA0DD"];
"gene_RELA" -- "term_1_BIOCARTA_CARDIACEGF_PATHWAY" [color="#DDA0DD"];
"gene_DDOST" [label="DDOST", type="gene", width=0.7, height=0.7];
"gene_DPM1" [label="DPM1", type="gene", width=0.7, height=0.7];
"gene_EXOSC3" [label="EXOSC3", type="gene", width=0.7, height=0.7];
"gene_EXOSC5" [label="EXOSC5", type="gene", width=0.7, height=0.7];
"gene_IKBKG" [label="IKBKG", type="gene", width=0.7, height=0.7];
"gene_JAK1" [label="JAK1", type="gene", width=0.7, height=0.7];
"gene_LSM3" [label="LSM3", type="gene", width=0.7, height=0.7];
"gene_MGAT1" [label="MGAT1", type="gene", width=0.7, height=0.7];
"gene_MPHOSPH6" [label="MPHOSPH6", type="gene", width=0.7, height=0.7];
"gene_PIGH" [label="PIGH", type="gene", width=0.7, height=0.7];
"gene_PIGK" [label="PIGK", type="gene", width=0.7, height=0.7];
"gene_PIGY" [label="PIGY", type="gene", width=0.7, height=0.7];
"gene_POLR1H" [label="POLR1H", type="gene", width=0.7, height=0.7];
"gene_POLR3A" [label="POLR3A", type="gene", width=0.7, height=0.7];
"gene_POLR3F" [label="POLR3F", type="gene", width=0.7, height=0.7];
"gene_STT3A" [label="STT3A", type="gene", width=0.7, height=0.7];
"term_1_KEGG_PANCREATIC_CANCER" [label="KEGG: PANCREATIC CANCER", style=filled, fillcolor="#F8C471", fontcolor="white", type="term", width=2, height=2, library="C2: KEGG Legacy"];
"term_2_KEGG_N_GLYCAN_BIOSYNTHESIS" [label="KEGG: N GLYCAN BIOSYNTHESIS", style=filled, fillcolor="#F8C471", fontcolor="white", type="term", width=2, height=2, library="C2: KEGG Legacy"];
"term_3_KEGG_GLYCOSYLPHOSPHATIDYLINOSITOL_GPI_ANCHOR_BIOSYNTHESIS" [label="KEGG: GLYCOSYLPHOSPHATIDYLINOSITOL GPI ANCHOR BIOSYNTHESIS", style=filled, fillcolor="#F8C471", fontcolor="white", type="term", width=2, height=2,
library="C2: KEGG Legacy"];
"term_4_KEGG_RNA_POLYMERASE" [label="KEGG: RNA POLYMERASE", style=filled, fillcolor="#F8C471", fontcolor="white", type="term", width=2, height=2, library="C2: KEGG Legacy"];
"term_5_KEGG_RNA_DEGRADATION" [label="KEGG: RNA DEGRADATION", style=filled, fillcolor="#F8C471", fontcolor="white", type="term", width=2, height=2, library="C2: KEGG Legacy"];
"gene_AKT1" -- "term_1_KEGG_PANCREATIC_CANCER" [color="#F8C471"];
"gene_DDOST" -- "term_2_KEGG_N_GLYCAN_BIOSYNTHESIS" [color="#F8C471"];
"gene_DPM1" -- "term_2_KEGG_N_GLYCAN_BIOSYNTHESIS" [color="#F8C471"];
"gene_EGF" -- "term_1_KEGG_PANCREATIC_CANCER" [color="#F8C471"];
"gene_EGFR" -- "term_1_KEGG_PANCREATIC_CANCER" [color="#F8C471"];
"gene_EXOSC3" -- "term_5_KEGG_RNA_DEGRADATION" [color="#F8C471"];
"gene_EXOSC5" -- "term_5_KEGG_RNA_DEGRADATION" [color="#F8C471"];
"gene_IKBKG" -- "term_1_KEGG_PANCREATIC_CANCER" [color="#F8C471"];
"gene_JAK1" -- "term_1_KEGG_PANCREATIC_CANCER" [color="#F8C471"];
"gene_LSM3" -- "term_5_KEGG_RNA_DEGRADATION" [color="#F8C471"];
"gene_MGAT1" -- "term_2_KEGG_N_GLYCAN_BIOSYNTHESIS" [color="#F8C471"];
"gene_MPHOSPH6" -- "term_5_KEGG_RNA_DEGRADATION" [color="#F8C471"];
"gene_PIGH" -- "term_3_KEGG_GLYCOSYLPHOSPHATIDYLINOSITOL_GPI_ANCHOR_BIOSYNTHESIS" [color="#F8C471"];
"gene_PIGK" -- "term_3_KEGG_GLYCOSYLPHOSPHATIDYLINOSITOL_GPI_ANCHOR_BIOSYNTHESIS" [color="#F8C471"];
"gene_PIGY" -- "term_3_KEGG_GLYCOSYLPHOSPHATIDYLINOSITOL_GPI_ANCHOR_BIOSYNTHESIS" [color="#F8C471"];
"gene_POLR1H" -- "term_4_KEGG_RNA_POLYMERASE" [color="#F8C471"];
"gene_POLR3A" -- "term_4_KEGG_RNA_POLYMERASE" [color="#F8C471"];
"gene_POLR3F" -- "term_4_KEGG_RNA_POLYMERASE" [color="#F8C471"];
"gene_RELA" -- "term_1_KEGG_PANCREATIC_CANCER" [color="#F8C471"];
"gene_STT3A" -- "term_2_KEGG_N_GLYCAN_BIOSYNTHESIS" [color="#F8C471"];
"gene_NUP107" [label="NUP107", type="gene", width=0.7, height=0.7];
"gene_NUP133" [label="NUP133", type="gene", width=0.7, height=0.7];
"gene_NUP155" [label="NUP155", type="gene", width=0.7, height=0.7];
"gene_NUP160" [label="NUP160", type="gene", width=0.7, height=0.7];
"gene_NUP85" [label="NUP85", type="gene", width=0.7, height=0.7];
"gene_RANBP2" [label="RANBP2", type="gene", width=0.7, height=0.7];
"term_1_KEGG_MEDICUS_REFERENCE_EGF_EGFR_PI3K_NFKB_SIGNALING_PATHWAY" [label="KEGG: MEDICUS REFERENCE EGF EGFR PI3K NFKB SIGNALING PATHWAY", style=filled, fillcolor="#98D8C8", fontcolor="white", type="term", width=2, height=2,
library="C2: KEGG MEDICUS"];
```

"term_2_KEGG_MEDICUS_REFERENCE_NUCLEAR_EXPORT_OF_MRNA" [label="KEGG: MEDICUS REFERENCE NUCLEAR EXPORT OF MRNA", style=filled, fillcolor="#98D8C8", fontcolor="white", type="term", width=2, height=2, library="C2: KEGG MEDICUS"];
"term_3_KEGG_MEDICUS_REFERENCE_GPI_ANCHOR_BIOSYNTHESIS" [label="KEGG: MEDICUS REFERENCE GPI ANCHOR BIOSYNTHESIS", style=filled, fillcolor="#98D8C8", fontcolor="white", type="term", width=2, height=2, library="C2: KEGG MEDICUS"];
"gene_AKT1" -- "term_1_KEGG_MEDICUS_REFERENCE_EGF_EGFR_PI3K_NFKB_SIGNALING_PATHWAY" [color="#98D8C8"];
"gene_EGF" -- "term_1_KEGG_MEDICUS_REFERENCE_EGF_EGFR_PI3K_NFKB_SIGNALING_PATHWAY" [color="#98D8C8"];
"gene_EGFR" -- "term_1_KEGG_MEDICUS_REFERENCE_EGF_EGFR_PI3K_NFKB_SIGNALING_PATHWAY" [color="#98D8C8"];
"gene_IKBKG" -- "term_1_KEGG_MEDICUS_REFERENCE_EGF_EGFR_PI3K_NFKB_SIGNALING_PATHWAY" [color="#98D8C8"];
"gene_NUP107" -- "term_2_KEGG_MEDICUS_REFERENCE_NUCLEAR_EXPORT_OF_MRNA" [color="#98D8C8"];
"gene_NUP133" -- "term_2_KEGG_MEDICUS_REFERENCE_NUCLEAR_EXPORT_OF_MRNA" [color="#98D8C8"];
"gene_NUP155" -- "term_2_KEGG_MEDICUS_REFERENCE_NUCLEAR_EXPORT_OF_MRNA" [color="#98D8C8"];
"gene_NUP160" -- "term_2_KEGG_MEDICUS_REFERENCE_NUCLEAR_EXPORT_OF_MRNA" [color="#98D8C8"];
"gene_NUP85" -- "term_2_KEGG_MEDICUS_REFERENCE_NUCLEAR_EXPORT_OF_MRNA" [color="#98D8C8"];
"gene_PIGH" -- "term_3_KEGG_MEDICUS_REFERENCE_GPI_ANCHOR_BIOSYNTHESIS" [color="#98D8C8"];
"gene_PIGK" -- "term_3_KEGG_MEDICUS_REFERENCE_GPI_ANCHOR_BIOSYNTHESIS" [color="#98D8C8"];
"gene_PIGY" -- "term_3_KEGG_MEDICUS_REFERENCE_GPI_ANCHOR_BIOSYNTHESIS" [color="#98D8C8"];
"gene_RANBP2" -- "term_2_KEGG_MEDICUS_REFERENCE_NUCLEAR_EXPORT_OF_MRNA" [color="#98D8C8"];
"gene_RELA" -- "term_1_KEGG_MEDICUS_REFERENCE_EGF_EGFR_PI3K_NFKB_SIGNALING_PATHWAY" [color="#98D8C8"];
"gene_CD4" [label="CD4", type="gene", width=0.7, height=0.7];
"gene_PIP5K1C" [label="PIP5K1C", type="gene", width=0.7, height=0.7];
"gene_RAP1B" [label="RAP1B", type="gene", width=0.7, height=0.7];
"gene_RAPGEF1" [label="RAPGEF1", type="gene", width=0.7, height=0.7];
"term_1_PID_ARF_3PATHWAY" [label="PID: ARF 3PATHWAY", style=filled, fillcolor="#440154", fontcolor="white", type="term", width=2, height=2, library="Unknown Library"];
"term_2_PID_ECADHERIN_NASCENT_AJ_PATHWAY" [label="PID: ECADHERIN NASCENT AJ PATHWAY", style=filled, fillcolor="#440154", fontcolor="white", type="term", width=2, height=2, library="Unknown Library"];
"gene_AKT1" -- "term_2_PID_ECADHERIN_NASCENT_AJ_PATHWAY" [color="#440154"];
"gene_AP2M1" -- "term_1_PID_ARF_3PATHWAY" [color="#440154"];
"gene_ARF1" -- "term_1_PID_ARF_3PATHWAY" [color="#440154"];
"gene_CD4" -- "term_1_PID_ARF_3PATHWAY" [color="#440154"];
"gene_GOSR2" -- "term_1_PID_ARF_3PATHWAY" [color="#440154"];
"gene_PIP5K1C" -- "term_2_PID_ECADHERIN_NASCENT_AJ_PATHWAY" [color="#440154"];
"gene_RAP1B" -- "term_2_PID_ECADHERIN_NASCENT_AJ_PATHWAY" [color="#440154"];
"gene_RAPGEF1" -- "term_2_PID_ECADHERIN_NASCENT_AJ_PATHWAY" [color="#440154"];
"gene_ARPC1A" [label="ARPC1A", type="gene", width=0.7, height=0.7];
"gene_COG3" [label="COG3", type="gene", width=0.7, height=0.7];
"gene_COG4" [label="COG4", type="gene", width=0.7, height=0.7];
"gene_CTDP1" [label="CTDP1", type="gene", width=0.7, height=0.7];
"gene_ELOA" [label="ELOA", type="gene", width=0.7, height=0.7];
"gene_ERCC3" [label="ERCC3", type="gene", width=0.7, height=0.7];
"gene_GAPVD1" [label="GAPVD1", type="gene", width=0.7, height=0.7];
"gene_GCK" [label="GCK", type="gene", width=0.7, height=0.7];
"gene_HIP1R" [label="HIP1R", type="gene", width=0.7, height=0.7];
"gene_HNRNPF" [label="HNRNPF", type="gene", width=0.7, height=0.7];
"gene_LPL" [label="LPL", type="gene", width=0.7, height=0.7];
"gene_LY6D" [label="LY6D", type="gene", width=0.7, height=0.7];
"gene_LYPD4" [label="LYPD4", type="gene", width=0.7, height=0.7];
"gene_MED14" [label="MED14", type="gene", width=0.7, height=0.7];
"gene_MED28" [label="MED28", type="gene", width=0.7, height=0.7];
"gene_MED4" [label="MED4", type="gene", width=0.7, height=0.7];
"gene_MED6" [label="MED6", type="gene", width=0.7, height=0.7];
"gene_MED7" [label="MED7", type="gene", width=0.7, height=0.7];
"gene_NCOR2" [label="NCOR2", type="gene", width=0.7, height=0.7];
"gene_NGLY1" [label="NGLY1", type="gene", width=0.7, height=0.7];
"gene_NUP153" [label="NUP153", type="gene", width=0.7, height=0.7];
"gene_PSME2" [label="PSME2", type="gene", width=0.7, height=0.7];
"gene_RAB1B" [label="RAB1B", type="gene", width=0.7, height=0.7];
"gene_RAB6B" [label="RAB6B", type="gene", width=0.7, height=0.7];
"gene_RABEPK" [label="RABEPK", type="gene", width=0.7, height=0.7];
"gene_SCFD1" [label="SCFD1", type="gene", width=0.7, height=0.7];
"gene_SPTAN1" [label="SPTAN1", type="gene", width=0.7, height=0.7];
"gene_SPTBN1" [label="SPTBN1", type="gene", width=0.7, height=0.7];
"gene_ST3GAL5" [label="ST3GAL5", type="gene", width=0.7, height=0.7];
"gene_STX5" [label="STX5", type="gene", width=0.7, height=0.7];
"gene_TRAPPC1" [label="TRAPPC1", type="gene", width=0.7, height=0.7];
"gene_TUBAL3" [label="TUBAL3", type="gene", width=0.7, height=0.7];
"gene_VPS53" [label="VPS53", type="gene", width=0.7, height=0.7];
"gene_WNT1" [label="WNT1", type="gene", width=0.7, height=0.7];
"gene_YTHDC2" [label="YTHDC2", type="gene", width=0.7, height=0.7];
"term_1_REACTOME_REGULATION_OF_GLUCOKINASE_BY_GLUCOKINASE_REGULATORY_PROTEIN" [label="REACTOME: REGULATION OF GLUCOKINASE BY GLUCOKINASE REGULATORY PROTEIN", style=filled, fillcolor="#BB8FCE", fontcolor="white", type="term", width=2, height=2, library="C2: Reactome Pathways"];
"term_2_REACTOME_ASPARAGINE_N_LINKED_GLYCOSYLATION" [label="REACTOME: ASPARAGINE N LINKED GLYCOSYLATION", style=filled, fillcolor="#BB8FCE", fontcolor="white", type="term", width=2, height=2, library="C2: Reactome Pathways"];
"term_3_REACTOME_TRANSCRIPTIONAL_REGULATION_OF_WHITE_ADIPOCYTE_DIFFERENTIATION" [label="REACTOME: TRANSCRIPTIONAL REGULATION OF WHITE ADIPOCYTE DIFFERENTIATION", style=filled, fillcolor="#BB8FCE", fontcolor="white", type="term", width=2, height=2, library="C2: Reactome Pathways"];
"term_4_REACTOME_CLATHRIN_MEDIATED_ENDOCYTOSIS" [label="REACTOME: CLATHRIN MEDIATED ENDOCYTOSIS", style=filled, fillcolor="#BB8FCE", fontcolor="white", type="term", width=2, height=2, library="C2: Reactome Pathways"];
"term_5_REACTOME_NUCLEAR_RNA_DECAY" [label="REACTOME: NUCLEAR RNA DECAY", style=filled, fillcolor="#BB8FCE", fontcolor="white", type="term", width=2, height=2, library="C2: Reactome Pathways"];
"term_6_REACTOME_INTERLEUKIN_12_SIGNALING" [label="REACTOME: INTERLEUKIN 12 SIGNALING", style=filled, fillcolor="#BB8FCE", fontcolor="white", type="term", width=2, height=2, library="C2: Reactome Pathways"];
"term_7_REACTOME_RETROGRADE_TRANSPORT_AT_THE_TRANS_GOLGI_NETWORK" [label="REACTOME: RETROGRADE TRANSPORT AT THE TRANS GOLGI NETWORK", style=filled, fillcolor="#BB8FCE", fontcolor="white", type="term", width=2, height=2, library="C2: Reactome Pathways"];
"term_8_REACTOME_POST_TRANSLATIONAL_MODIFICATION_SYNTHESIS_OF_GPI_ANCHORED_PROTEINS" [label="REACTOME: POST TRANSLATIONAL MODIFICATION SYNTHESIS OF GPI ANCHORED PROTEINS", style=filled, fillcolor="#BB8FCE", fontcolor="white", type="term", width=2, height=2, library="C2: Reactome Pathways"];
"term_9_REACTOME_HIV_TRANSCRIPTION_ELONGATION" [label="REACTOME: HIV TRANSCRIPTION ELONGATION", style=filled, fillcolor="#BB8FCE", fontcolor="white", type="term", width=2, height=2, library="C2: Reactome Pathways"];
"gene_AP2M1" -- "term_4_REACTOME_CLATHRIN_MEDIATED_ENDOCYTOSIS" [color="#BB8FCE"];
"gene_ARF1" -- "term_2_REACTOME_ASPARAGINE_N_LINKED_GLYCOSYLATION" [color="#BB8FCE"];
"gene_ARPC1A" -- "term_4_REACTOME_CLATHRIN_MEDIATED_ENDOCYTOSIS" [color="#BB8FCE"];
"gene_CD4" -- "term_4_REACTOME_CLATHRIN_MEDIATED_ENDOCYTOSIS" [color="#BB8FCE"];
"gene_COG2" -- "term_2_REACTOME_ASPARAGINE_N_LINKED_GLYCOSYLATION" [color="#BB8FCE"];
"gene_COG3" -- "term_2_REACTOME_ASPARAGINE_N_LINKED_GLYCOSYLATION" [color="#BB8FCE"];
"gene_COG4" -- "term_2_REACTOME_ASPARAGINE_N_LINKED_GLYCOSYLATION" [color="#BB8FCE"];
"gene_CTDP1" -- "term_9_REACTOME_HIV_TRANSCRIPTION_ELONGATION" [color="#BB8FCE"];
"gene_DDOST" -- "term_2_REACTOME_ASPARAGINE_N_LINKED_GLYCOSYLATION" [color="#BB8FCE"];
"gene_DPM1" -- "term_2_REACTOME_ASPARAGINE_N_LINKED_GLYCOSYLATION" [color="#BB8FCE"];
"gene_EGF" -- "term_4_REACTOME_CLATHRIN_MEDIATED_ENDOCYTOSIS" [color="#BB8FCE"];
"gene_EGFR" -- "term_4_REACTOME_CLATHRIN_MEDIATED_ENDOCYTOSIS" [color="#BB8FCE"];
"gene_ELOA" -- "term_9_REACTOME_HIV_TRANSCRIPTION_ELONGATION" [color="#BB8FCE"];
"gene_ERCC3" -- "term_9_REACTOME_HIV_TRANSCRIPTION_ELONGATION" [color="#BB8FCE"];
"gene_EXOSC3" -- "term_5_REACTOME_NUCLEAR_RNA_DECAY" [color="#BB8FCE"];
"gene_EXOSC5" -- "term_5_REACTOME_NUCLEAR_RNA_DECAY" [color="#BB8FCE"];
"gene_GAPVD1" -- "term_4_REACTOME_CLATHRIN_MEDIATED_ENDOCYTOSIS" [color="#BB8FCE"];
"gene_GCK" -- "term_1_REACTOME_REGULATION_OF_GLUCOKINASE_BY_GLUCOKINASE_REGULATORY_PROTEIN" [color="#BB8FCE"];
"gene_GOSR2" -- "term_2_REACTOME_ASPARAGINE_N_LINKED_GLYCOSYLATION" [color="#BB8FCE"];
"gene_HIP1R" -- "term_4_REACTOME_CLATHRIN_MEDIATED_ENDOCYTOSIS" [color="#BB8FCE"];
"gene_HNRNPF" -- "term_6_REACTOME_INTERLEUKIN_12_SIGNALING" [color="#BB8FCE"];
"gene_JAK1" -- "term_6_REACTOME_INTERLEUKIN_12_SIGNALING" [color="#BB8FCE"];
"gene_LPL" -- "term_3_REACTOME_TRANSCRIPTIONAL_REGULATION_OF_WHITE_ADIPOCYTE_DIFFERENTIATION" [color="#BB8FCE"];
"gene_LY6D" -- "term_8_REACTOME_POST_TRANSLATIONAL_MODIFICATION_SYNTHESIS_OF_GPI_ANCHORED_PROTEINS" [color="#BB8FCE"];
"gene_LYPD4" -- "term_8_REACTOME_POST_TRANSLATIONAL_MODIFICATION_SYNTHESIS_OF_GPI_ANCHORED_PROTEINS" [color="#BB8FCE"];
"gene_MED14" -- "term_3_REACTOME_TRANSCRIPTIONAL_REGULATION_OF_WHITE_ADIPOCYTE_DIFFERENTIATION" [color="#BB8FCE"];
"gene_MED28" -- "term_3_REACTOME_TRANSCRIPTIONAL_REGULATION_OF_WHITE_ADIPOCYTE_DIFFERENTIATION" [color="#BB8FCE"];
"gene_MED4" -- "term_3_REACTOME_TRANSCRIPTIONAL_REGULATION_OF_WHITE_ADIPOCYTE_DIFFERENTIATION" [color="#BB8FCE"];
"gene_MED6" -- "term_3_REACTOME_TRANSCRIPTIONAL_REGULATION_OF_WHITE_ADIPOCYTE_DIFFERENTIATION" [color="#BB8FCE"];
"gene_MED7" -- "term_3_REACTOME_TRANSCRIPTIONAL_REGULATION_OF_WHITE_ADIPOCYTE_DIFFERENTIATION" [color="#BB8FCE"];
"gene_MGAT1" -- "term_2_REACTOME_ASPARAGINE_N_LINKED_GLYCOSYLATION" [color="#BB8FCE"];
"gene_MPHOSPH6" -- "term_5_REACTOME_NUCLEAR_RNA_DECAY" [color="#BB8FCE"];
"gene_NCOR2" -- "term_3_REACTOME_TRANSCRIPTIONAL_REGULATION_OF_WHITE_ADIPOCYTE_DIFFERENTIATION" [color="#BB8FCE"];
"gene_NGLY1" -- "term_2_REACTOME_ASPARAGINE_N_LINKED_GLYCOSYLATION" [color="#BB8FCE"];
"gene_NUP107" -- "term_1_REACTOME_REGULATION_OF_GLUCOKINASE_BY_GLUCOKINASE_REGULATORY_PROTEIN" [color="#BB8FCE"];
"gene_NUP133" -- "term_1_REACTOME_REGULATION_OF_GLUCOKINASE_BY_GLUCOKINASE_REGULATORY_PROTEIN" [color="#BB8FCE"];
"gene_NUP153" -- "term_1_REACTOME_REGULATION_OF_GLUCOKINASE_BY_GLUCOKINASE_REGULATORY_PROTEIN" [color="#BB8FCE"];
"gene_NUP155" -- "term_1_REACTOME_REGULATION_OF_GLUCOKINASE_BY_GLUCOKINASE_REGULATORY_PROTEIN" [color="#BB8FCE"];
"gene_NUP160" -- "term_1_REACTOME_REGULATION_OF_GLUCOKINASE_BY_GLUCOKINASE_REGULATORY_PROTEIN" [color="#BB8FCE"];
"gene_NUP85" -- "term_1_REACTOME_REGULATION_OF_GLUCOKINASE_BY_GLUCOKINASE_REGULATORY_PROTEIN" [color="#BB8FCE"];
"gene_PIGH" -- "term_8_REACTOME_POST_TRANSLATIONAL_MODIFICATION_SYNTHESIS_OF_GPI_ANCHORED_PROTEINS" [color="#BB8FCE"];
"gene_PIGK" -- "term_8_REACTOME_POST_TRANSLATIONAL_MODIFICATION_SYNTHESIS_OF_GPI_ANCHORED_PROTEINS" [color="#BB8FCE"];
"gene_PIGY" -- "term_8_REACTOME_POST_TRANSLATIONAL_MODIFICATION_SYNTHESIS_OF_GPI_ANCHORED_PROTEINS" [color="#BB8FCE"];
"gene_PIP5K1C" -- "term_4_REACTOME_CLATHRIN_MEDIATED_ENDOCYTOSIS" [color="#BB8FCE"];
"gene_PSME2" -- "term_6_REACTOME_INTERLEUKIN_12_SIGNALING" [color="#BB8FCE"];
"gene_RAB1B" -- "term_2_REACTOME_ASPARAGINE_N_LINKED_GLYCOSYLATION" [color="#BB8FCE"];
"gene_RAB6A" -- "term_7_REACTOME_RETROGRADE_TRANSPORT_AT_THE_TRANS_GOLGI_NETWORK" [color="#BB8FCE"];
"gene_RAB6B" -- "term_7_REACTOME_RETROGRADE_TRANSPORT_AT_THE_TRANS_GOLGI_NETWORK" [color="#BB8FCE"];
"gene_RABEPK" -- "term_7_REACTOME_RETROGRADE_TRANSPORT_AT_THE_TRANS_GOLGI_NETWORK" [color="#BB8FCE"];
"gene_RANBP2" -- "term_1_REACTOME_REGULATION_OF_GLUCOKINASE_BY_GLUCOKINASE_REGULATORY_PROTEIN" [color="#BB8FCE"];
"gene_RAP1B" -- "term_6_REACTOME_INTERLEUKIN_12_SIGNALING" [color="#BB8FCE"];
"gene_RELA" -- "term_3_REACTOME_TRANSCRIPTIONAL_REGULATION_OF_WHITE_ADIPOCYTE_DIFFERENTIATION" [color="#BB8FCE"];
"gene_SCFD1" -- "term_2_REACTOME_ASPARAGINE_N_LINKED_GLYCOSYLATION" [color="#BB8FCE"];
"gene_SPTAN1" -- "term_2_REACTOME_ASPARAGINE_N_LINKED_GLYCOSYLATION" [color="#BB8FCE"];
"gene_SPTBN1" -- "term_2_REACTOME_ASPARAGINE_N_LINKED_GLYCOSYLATION" [color="#BB8FCE"];
"gene_ST3GAL5" -- "term_2_REACTOME_ASPARAGINE_N_LINKED_GLYCOSYLATION" [color="#BB8FCE"];
"gene_STT3A" -- "term_2_REACTOME_ASPARAGINE_N_LINKED_GLYCOSYLATION" [color="#BB8FCE"];
"gene_STX5" -- "term_2_REACTOME_ASPARAGINE_N_LINKED_GLYCOSYLATION" [color="#BB8FCE"];
"gene_TMED2" -- "term_2_REACTOME_ASPARAGINE_N_LINKED_GLYCOSYLATION" [color="#BB8FCE"];
"gene_TRAPPC1" -- "term_2_REACTOME_ASPARAGINE_N_LINKED_GLYCOSYLATION" [color="#BB8FCE"];
"gene_TUBAL3" -- "term_2_REACTOME_ASPARAGINE_N_LINKED_GLYCOSYLATION" [color="#BB8FCE"];
"gene_VPS53" -- "term_7_REACTOME_RETROGRADE_TRANSPORT_AT_THE_TRANS_GOLGI_NETWORK" [color="#BB8FCE"];

"gene_WNT1" -- "term_3_REACTOME_TRANSCRIPTIONAL_REGULATION_OF_WHITE_ADIPOCYTE_DIFFERENTIATION" [color="#BB8FCE"];
"gene_YTHDC2" -- "term_5_REACTOME_NUCLEAR_RNA_DECAY" [color="#BB8FCE"];
"gene_AGO3" [label="AGO3", type="gene", width=0.7, height=0.7];
"gene_ATP6V0A1" [label="ATP6V0A1", type="gene", width=0.7, height=0.7];
"gene_DNAJB1" [label="DNAJB1", type="gene", width=0.7, height=0.7];
"gene_FBXW11" [label="FBXW11", type="gene", width=0.7, height=0.7];
"gene_GEMIN2" [label="GEMIN2", type="gene", width=0.7, height=0.7];
"gene_IDH1" [label="IDH1", type="gene", width=0.7, height=0.7];
"gene_LEFTY1" [label="LEFTY1", type="gene", width=0.7, height=0.7];
"gene_NF2" [label="NF2", type="gene", width=0.7, height=0.7];
"gene_PPIB" [label="PPIB", type="gene", width=0.7, height=0.7];
"gene_PURA" [label="PURA", type="gene", width=0.7, height=0.7];
"gene_SPCS3" [label="SPCS3", type="gene", width=0.7, height=0.7];
"gene_TFE3" [label="TFE3", type="gene", width=0.7, height=0.7];
"term_1_WP_PRADERWILLI_AND_ANGELMAN_SYNDROME" [label="WP: PRADERWILLI AND ANGELMAN SYNDROME", style=filled, fillcolor="#31688e", fontcolor="white", type="term", width=2, height=2, library="Unknown Library"];
"term_2_WP_IL17_SIGNALING" [label="WP: IL17 SIGNALING", style=filled, fillcolor="#31688e", fontcolor="white", type="term", width=2, height=2, library="Unknown Library"];
"term_3_WP_GLYCOSYLPHOSPHATIDYL_INOSITOL_ANCHOR_PATHWAY" [label="WP: GLYCOSYLPHOSPHATIDYL INOSITOL ANCHOR PATHWAY", style=filled, fillcolor="#31688e", fontcolor="white", type="term", width=2, height=2, library="Unknown Library"];
"term_4_WP_TGFBETA_RECEPTOR_SIGNALING" [label="WP: TGFBETA RECEPTOR SIGNALING", style=filled, fillcolor="#31688e", fontcolor="white", type="term", width=2, height=2, library="Unknown Library"];
"term_5_WP_MIRTARGETED_GENES_IN_LYMPHOCYTES" [label="WP: MIRTARGETED GENES IN LYMPHOCYTES", style=filled, fillcolor="#31688e", fontcolor="white", type="term", width=2, height=2, library="Unknown Library"];
"term_6_WP_NGLYCAN_BIOSYNTHESIS" [label="WP: NGLYCAN BIOSYNTHESIS", style=filled, fillcolor="#31688e", fontcolor="white", type="term", width=2, height=2, library="Unknown Library"];
"gene_AGO3" -- "term_5_WP_MIRTARGETED_GENES_IN_LYMPHOCYTES" [color="#31688e"];
"gene_AKT1" -- "term_2_WP_IL17_SIGNALING" [color="#31688e"];
"gene_ATP6V0A1" -- "term_5_WP_MIRTARGETED_GENES_IN_LYMPHOCYTES" [color="#31688e"];
"gene_DDOST" -- "term_6_WP_NGLYCAN_BIOSYNTHESIS" [color="#31688e"];
"gene_DNAJB1" -- "term_5_WP_MIRTARGETED_GENES_IN_LYMPHOCYTES" [color="#31688e"];
"gene_DPM1" -- "term_6_WP_NGLYCAN_BIOSYNTHESIS" [color="#31688e"];
"gene_EGF" -- "term_4_WP_TGFBETA_RECEPTOR_SIGNALING" [color="#31688e"];
"gene_EGFR" -- "term_5_WP_MIRTARGETED_GENES_IN_LYMPHOCYTES" [color="#31688e"];
"gene_FBXW11" -- "term_5_WP_MIRTARGETED_GENES_IN_LYMPHOCYTES" [color="#31688e"];
"gene_GEMIN2" -- "term_5_WP_MIRTARGETED_GENES_IN_LYMPHOCYTES" [color="#31688e"];
"gene_IDH1" -- "term_5_WP_MIRTARGETED_GENES_IN_LYMPHOCYTES" [color="#31688e"];
"gene_IKBKG" -- "term_2_WP_IL17_SIGNALING" [color="#31688e"];
"gene_JAK1" -- "term_2_WP_IL17_SIGNALING" [color="#31688e"];
"gene_LEFTY1" -- "term_4_WP_TGFBETA_RECEPTOR_SIGNALING" [color="#31688e"];
"gene_LPL" -- "term_5_WP_MIRTARGETED_GENES_IN_LYMPHOCYTES" [color="#31688e"];
"gene_MGAT1" -- "term_6_WP_NGLYCAN_BIOSYNTHESIS" [color="#31688e"];
"gene_NF2" -- "term_5_WP_MIRTARGETED_GENES_IN_LYMPHOCYTES" [color="#31688e"];
"gene_NUP107" -- "term_1_WP_PRADERWILLI_AND_ANGELMAN_SYNDROME" [color="#31688e"];
"gene_NUP133" -- "term_1_WP_PRADERWILLI_AND_ANGELMAN_SYNDROME" [color="#31688e"];
"gene_NUP153" -- "term_1_WP_PRADERWILLI_AND_ANGELMAN_SYNDROME" [color="#31688e"];
"gene_NUP155" -- "term_1_WP_PRADERWILLI_AND_ANGELMAN_SYNDROME" [color="#31688e"];
"gene_NUP160" -- "term_1_WP_PRADERWILLI_AND_ANGELMAN_SYNDROME" [color="#31688e"];
"gene_NUP85" -- "term_1_WP_PRADERWILLI_AND_ANGELMAN_SYNDROME" [color="#31688e"];
"gene_PIGH" -- "term_3_WP_GLYCOSYLPHOSPHATIDYL_INOSITOL_ANCHOR_PATHWAY" [color="#31688e"];
"gene_PIGK" -- "term_3_WP_GLYCOSYLPHOSPHATIDYL_INOSITOL_ANCHOR_PATHWAY" [color="#31688e"];
"gene_PIGY" -- "term_3_WP_GLYCOSYLPHOSPHATIDYL_INOSITOL_ANCHOR_PATHWAY" [color="#31688e"];
"gene_PPIB" -- "term_5_WP_MIRTARGETED_GENES_IN_LYMPHOCYTES" [color="#31688e"];
"gene_PURA" -- "term_5_WP_MIRTARGETED_GENES_IN_LYMPHOCYTES" [color="#31688e"];
"gene_RAB6A" -- "term_5_WP_MIRTARGETED_GENES_IN_LYMPHOCYTES" [color="#31688e"];
"gene_RANBP2" -- "term_1_WP_PRADERWILLI_AND_ANGELMAN_SYNDROME" [color="#31688e"];
"gene_RELA" -- "term_2_WP_IL17_SIGNALING" [color="#31688e"];
"gene_SPCS3" -- "term_5_WP_MIRTARGETED_GENES_IN_LYMPHOCYTES" [color="#31688e"];
"gene_STT3A" -- "term_6_WP_NGLYCAN_BIOSYNTHESIS" [color="#31688e"];
"gene_TFE3" -- "term_4_WP_TGFBETA_RECEPTOR_SIGNALING" [color="#31688e"];
"gene_TMED2" -- "term_5_WP_MIRTARGETED_GENES_IN_LYMPHOCYTES" [color="#31688e"];
"gene_WNT1" -- "term_4_WP_TGFBETA_RECEPTOR_SIGNALING" [color="#31688e"];
"gene_A4GALT" [label="A4GALT", type="gene", width=0.7, height=0.7];
"gene_AGAP2" [label="AGAP2", type="gene", width=0.7, height=0.7];
"gene_ATG12" [label="ATG12", type="gene", width=0.7, height=0.7];
"gene_CARD16" [label="CARD16", type="gene", width=0.7, height=0.7];
"gene_CLN3" [label="CLN3", type="gene", width=0.7, height=0.7];
"gene_CLNS1A" [label="CLNS1A", type="gene", width=0.7, height=0.7];
"gene_CRTC2" [label="CRTC2", type="gene", width=0.7, height=0.7];
"gene_DDX10" [label="DDX10", type="gene", width=0.7, height=0.7];
"gene_DHX33" [label="DHX33", type="gene", width=0.7, height=0.7];
"gene_DIMT1" [label="DIMT1", type="gene", width=0.7, height=0.7];
"gene_EIF3H" [label="EIF3H", type="gene", width=0.7, height=0.7];
"gene_EPS8" [label="EPS8", type="gene", width=0.7, height=0.7];
"gene_ETF1" [label="ETF1", type="gene", width=0.7, height=0.7];
"gene_FAM76B" [label="FAM76B", type="gene", width=0.7, height=0.7];
"gene_FLII" [label="FLII", type="gene", width=0.7, height=0.7];
"gene_GABARAPL2" [label="GABARAPL2", type="gene", width=0.7, height=0.7];
"gene_GOLPH3" [label="GOLPH3", type="gene", width=0.7, height=0.7];
"gene_HEATR1" [label="HEATR1", type="gene", width=0.7, height=0.7];
"gene_HTATSF1" [label="HTATSF1", type="gene", width=0.7, height=0.7];
"gene_HUWE1" [label="HUWE1", type="gene", width=0.7, height=0.7];
"gene_INTS7" [label="INTS7", type="gene", width=0.7, height=0.7];
"gene_KBTBD7" [label="KBTBD7", type="gene", width=0.7, height=0.7];
"gene_KDM4D" [label="KDM4D", type="gene", width=0.7, height=0.7];
"gene_LAPTM5" [label="LAPTM5", type="gene", width=0.7, height=0.7];
"gene_LARS1" [label="LARS1", type="gene", width=0.7, height=0.7];
"gene_LCP2" [label="LCP2", type="gene", width=0.7, height=0.7];
"gene_MDN1" [label="MDN1", type="gene", width=0.7, height=0.7];
"gene_MID1IP1" [label="MID1IP1", type="gene", width=0.7, height=0.7];
"gene_MKRN2" [label="MKRN2", type="gene", width=0.7, height=0.7];
"gene_NIPSNAP2" [label="NIPSNAP2", type="gene", width=0.7, height=0.7];
"gene_RNF170" [label="RNF170", type="gene", width=0.7, height=0.7];
"gene_RNF26" [label="RNF26", type="gene", width=0.7, height=0.7];
"gene_SPAST" [label="SPAST", type="gene", width=0.7, height=0.7];
"gene_STAC2" [label="STAC2", type="gene", width=0.7, height=0.7];
"gene_TAOK1" [label="TAOK1", type="gene", width=0.7, height=0.7];
"gene_THOC2" [label="THOC2", type="gene", width=0.7, height=0.7];
"gene_TM9SF2" [label="TM9SF2", type="gene", width=0.7, height=0.7];
"gene_TOMM70" [label="TOMM70", type="gene", width=0.7, height=0.7];
"gene_TRAPPC8" [label="TRAPPC8", type="gene", width=0.7, height=0.7];
"gene_TRIM55" [label="TRIM55", type="gene", width=0.7, height=0.7];
"gene_TRIM58" [label="TRIM58", type="gene", width=0.7, height=0.7];
"gene_UBQLN4" [label="UBQLN4", type="gene", width=0.7, height=0.7];
"gene_WNK1" [label="WNK1", type="gene", width=0.7, height=0.7];
"term_10_GOBP_POSITIVE_REGULATION_OF_INTERFERON_BETA_PRODUCTION" [label="GOBP: POSITIVE REGULATION OF INTERFERON BETA PRODUCTION", style=filled, fillcolor="#F1948A", fontcolor="white", type="term", width=2, height=2, library="C5: Gene Ontology: Biological Process"];
"term_11_GOBP_PROTEIN_N_LINKED_GLYCOSYLATION_VIA_ASPARAGINE" [label="GOBP: PROTEIN N LINKED GLYCOSYLATION VIA ASPARAGINE", style=filled, fillcolor="#F1948A", fontcolor="white", type="term", width=2, height=2, library="C5: Gene Ontology: Biological Process"];
"term_12_GOBP_ARP2_3_COMPLEX_MEDIATED_ACTIN_NUCLEATION" [label="GOBP: ARP2 3 COMPLEX MEDIATED ACTIN NUCLEATION", style=filled, fillcolor="#F1948A", fontcolor="white", type="term", width=2, height=2, library="C5: Gene Ontology: Biological Process"];
"term_1_GOBP_GOLGI_VESICLE_TRANSPORT" [label="GOBP: GOLGI VESICLE TRANSPORT", style=filled, fillcolor="#F1948A", fontcolor="white", type="term", width=2, height=2, library="C5: Gene Ontology: Biological Process"];
"term_2_GOBP_NUCLEAR_EXPORT" [label="GOBP: NUCLEAR EXPORT", style=filled, fillcolor="#F1948A", fontcolor="white", type="term", width=2, height=2, library="C5: Gene Ontology: Biological Process"];
"term_3_GOBP_DNA_TEMPLATED_TRANSCRIPTION_ELONGATION" [label="GOBP: DNA TEMPLATED TRANSCRIPTION ELONGATION", style=filled, fillcolor="#F1948A", fontcolor="white", type="term", width=2, height=2, library="C5: Gene Ontology: Biological Process"];
"term_4_GOBP_GLYCOLIPID_BIOSYNTHETIC_PROCESS" [label="GOBP: GLYCOLIPID BIOSYNTHETIC PROCESS", style=filled, fillcolor="#F1948A", fontcolor="white", type="term", width=2, height=2, library="C5: Gene Ontology: Biological Process"];
"term_5_GOBP_REGULATION_OF_PROTEIN_CONTAINING_COMPLEX_DISASSEMBLY" [label="GOBP: REGULATION OF PROTEIN CONTAINING COMPLEX DISASSEMBLY", style=filled, fillcolor="#F1948A", fontcolor="white", type="term", width=2, height=2, library="C5: Gene Ontology: Biological Process"];
"term_6_GOBP_POSITIVE_REGULATION_OF_MOLECULAR_FUNCTION" [label="GOBP: POSITIVE REGULATION OF MOLECULAR FUNCTION", style=filled, fillcolor="#F1948A", fontcolor="white", type="term", width=2, height=2, library="C5: Gene Ontology: Biological Process"];
"term_7_GOBP_POSITIVE_REGULATION_BY_HOST_OF_VIRAL_PROCESS" [label="GOBP: POSITIVE REGULATION BY HOST OF VIRAL PROCESS", style=filled, fillcolor="#F1948A", fontcolor="white", type="term", width=2, height=2, library="C5: Gene Ontology: Biological Process"];
"term_8_GOBP_PROTEIN_POLYUBIQUITINATION" [label="GOBP: PROTEIN POLYUBIQUITINATION", style=filled, fillcolor="#F1948A", fontcolor="white", type="term", width=2, height=2, library="C5: Gene Ontology: Biological Process"];
"term_9_GOBP_RIBONUCLEOPROTEIN_COMPLEX_BIOGENESIS" [label="GOBP: RIBONUCLEOPROTEIN COMPLEX BIOGENESIS", style=filled, fillcolor="#F1948A", fontcolor="white", type="term", width=2, height=2, library="C5: Gene Ontology: Biological Process"];
"gene_A4GALT" -- "term_4_GOBP_GLYCOLIPID_BIOSYNTHETIC_PROCESS" [color="#F1948A"];
"gene_AGAP2" -- "term_6_GOBP_POSITIVE_REGULATION_OF_MOLECULAR_FUNCTION" [color="#F1948A"];
"gene_AGO3" -- "term_9_GOBP_RIBONUCLEOPROTEIN_COMPLEX_BIOGENESIS" [color="#F1948A"];
"gene_AKT1" -- "term_6_GOBP_POSITIVE_REGULATION_OF_MOLECULAR_FUNCTION" [color="#F1948A"];
"gene_ARF1" -- "term_12_GOBP_ARP2_3_COMPLEX_MEDIATED_ACTIN_NUCLEATION" [color="#F1948A"];
"gene_ARPC1A" -- "term_12_GOBP_ARP2_3_COMPLEX_MEDIATED_ACTIN_NUCLEATION" [color="#F1948A"];
"gene_ATG12" -- "term_5_GOBP_REGULATION_OF_PROTEIN_CONTAINING_COMPLEX_DISASSEMBLY" [color="#F1948A"];
"gene_CARD16" -- "term_6_GOBP_POSITIVE_REGULATION_OF_MOLECULAR_FUNCTION" [color="#F1948A"];
"gene_CAV2" -- "term_7_GOBP_POSITIVE_REGULATION_BY_HOST_OF_VIRAL_PROCESS" [color="#F1948A"];
"gene_CD4" -- "term_6_GOBP_POSITIVE_REGULATION_OF_MOLECULAR_FUNCTION" [color="#F1948A"];
"gene_CLN3" -- "term_1_GOBP_GOLGI_VESICLE_TRANSPORT" [color="#F1948A"];
"gene_CLNS1A" -- "term_9_GOBP_RIBONUCLEOPROTEIN_COMPLEX_BIOGENESIS" [color="#F1948A"];
"gene_COG2" -- "term_1_GOBP_GOLGI_VESICLE_TRANSPORT" [color="#F1948A"];
"gene_COG3" -- "term_1_GOBP_GOLGI_VESICLE_TRANSPORT" [color="#F1948A"];
"gene_COG4" -- "term_1_GOBP_GOLGI_VESICLE_TRANSPORT" [color="#F1948A"];
"gene_CRTC2" -- "term_6_GOBP_POSITIVE_REGULATION_OF_MOLECULAR_FUNCTION" [color="#F1948A"];
"gene_CTDP1" -- "term_3_GOBP_DNA_TEMPLATED_TRANSCRIPTION_ELONGATION" [color="#F1948A"];
"gene_DDOST" -- "term_11_GOBP_PROTEIN_N_LINKED_GLYCOSYLATION_VIA_ASPARAGINE" [color="#F1948A"];
"gene_DDX10" -- "term_9_GOBP_RIBONUCLEOPROTEIN_COMPLEX_BIOGENESIS" [color="#F1948A"];
"gene_DHX33" -- "term_6_GOBP_POSITIVE_REGULATION_OF_MOLECULAR_FUNCTION" [color="#F1948A"];
"gene_DIMT1" -- "term_9_GOBP_RIBONUCLEOPROTEIN_COMPLEX_BIOGENESIS" [color="#F1948A"];
"gene_DPM1" -- "term_4_GOBP_GLYCOLIPID_BIOSYNTHETIC_PROCESS" [color="#F1948A"];

```
"gene_EDNRA" -- "term_6_GOBP_POSITIVE_REGULATION_OF_MOLECULAR_FUNCTION" [color="#F1948A"];
"gene_EGF" -- "term_6_GOBP_POSITIVE_REGULATION_OF_MOLECULAR_FUNCTION" [color="#F1948A"];
"gene_EIF3H" -- "term_9_GOBP_RIBONUCLEOPROTEIN_COMPLEX_BIOGENESIS" [color="#F1948A"];
"gene_ELOA" -- "term_3_GOBP_DNA_TEMPLATED_TRANSCRIPTION_ELONGATION" [color="#F1948A"];
"gene_EPS8" -- "term_5_GOBP_REGULATION_OF_PROTEIN_CONTAINING_COMPLEX_DISASSEMBLY" [color="#F1948A"];
"gene_ERCC3" -- "term_3_GOBP_DNA_TEMPLATED_TRANSCRIPTION_ELONGATION" [color="#F1948A"];
"gene_ETF1" -- "term_5_GOBP_REGULATION_OF_PROTEIN_CONTAINING_COMPLEX_DISASSEMBLY" [color="#F1948A"];
"gene_EXOSC3" -- "term_9_GOBP_RIBONUCLEOPROTEIN_COMPLEX_BIOGENESIS" [color="#F1948A"];
"gene_EXOSC5" -- "term_9_GOBP_RIBONUCLEOPROTEIN_COMPLEX_BIOGENESIS" [color="#F1948A"];
"gene_FAM76B" -- "term_2_GOBP_NUCLEAR_EXPORT" [color="#F1948A"];
"gene_FBXW11" -- "term_8_GOBP_PROTEIN_POLYUBIQUITINATION" [color="#F1948A"];
"gene_FLII" -- "term_5_GOBP_REGULATION_OF_PROTEIN_CONTAINING_COMPLEX_DISASSEMBLY" [color="#F1948A"];
"gene_GABARAPL2" -- "term_1_GOBP_GOLGI_VESICLE_TRANSPORT" [color="#F1948A"];
"gene_GEMIN2" -- "term_9_GOBP_RIBONUCLEOPROTEIN_COMPLEX_BIOGENESIS" [color="#F1948A"];
"gene_GOLPH3" -- "term_1_GOBP_GOLGI_VESICLE_TRANSPORT" [color="#F1948A"];
"gene_GOSR2" -- "term_1_GOBP_GOLGI_VESICLE_TRANSPORT" [color="#F1948A"];
"gene_HEATR1" -- "term_9_GOBP_RIBONUCLEOPROTEIN_COMPLEX_BIOGENESIS" [color="#F1948A"];
"gene_HIP1R" -- "term_12_GOBP_ARP2_3_COMPLEX_MEDIATED_ACTIN_NUCLEATION" [color="#F1948A"];
"gene_HTATSF1" -- "term_9_GOBP_RIBONUCLEOPROTEIN_COMPLEX_BIOGENESIS" [color="#F1948A"];
"gene_HUWE1" -- "term_8_GOBP_PROTEIN_POLYUBIQUITINATION" [color="#F1948A"];
"gene_IKBKG" -- "term_6_GOBP_POSITIVE_REGULATION_OF_MOLECULAR_FUNCTION" [color="#F1948A"];
"gene_INTS7" -- "term_3_GOBP_DNA_TEMPLATED_TRANSCRIPTION_ELONGATION" [color="#F1948A"];
"gene_KBTBD7" -- "term_8_GOBP_PROTEIN_POLYUBIQUITINATION" [color="#F1948A"];
"gene_KDM4D" -- "term_6_GOBP_POSITIVE_REGULATION_OF_MOLECULAR_FUNCTION" [color="#F1948A"];
"gene_LAPTM5" -- "term_1_GOBP_GOLGI_VESICLE_TRANSPORT" [color="#F1948A"];
"gene_LARS1" -- "term_6_GOBP_POSITIVE_REGULATION_OF_MOLECULAR_FUNCTION" [color="#F1948A"];
"gene_LCP2" -- "term_6_GOBP_POSITIVE_REGULATION_OF_MOLECULAR_FUNCTION" [color="#F1948A"];
"gene_MDN1" -- "term_2_GOBP_NUCLEAR_EXPORT" [color="#F1948A"];
"gene_MED14" -- "term_3_GOBP_DNA_TEMPLATED_TRANSCRIPTION_ELONGATION" [color="#F1948A"];
"gene_MED28" -- "term_3_GOBP_DNA_TEMPLATED_TRANSCRIPTION_ELONGATION" [color="#F1948A"];
"gene_MED4" -- "term_3_GOBP_DNA_TEMPLATED_TRANSCRIPTION_ELONGATION" [color="#F1948A"];
"gene_MED6" -- "term_3_GOBP_DNA_TEMPLATED_TRANSCRIPTION_ELONGATION" [color="#F1948A"];
"gene_MED7" -- "term_3_GOBP_DNA_TEMPLATED_TRANSCRIPTION_ELONGATION" [color="#F1948A"];
"gene_MGAT1" -- "term_11_GOBP_PROTEIN_N_LINKED_GLYCOSYLATION_VIA_ASPARAGINE" [color="#F1948A"];
"gene_MID1IP1" -- "term_5_GOBP_REGULATION_OF_PROTEIN_CONTAINING_COMPLEX_DISASSEMBLY" [color="#F1948A"];
"gene_MKRN2" -- "term_8_GOBP_PROTEIN_POLYUBIQUITINATION" [color="#F1948A"];
"gene_MPHOSPH6" -- "term_9_GOBP_RIBONUCLEOPROTEIN_COMPLEX_BIOGENESIS" [color="#F1948A"];
"gene_NIPSNAP2" -- "term_6_GOBP_POSITIVE_REGULATION_OF_MOLECULAR_FUNCTION" [color="#F1948A"];
"gene_NUP107" -- "term_2_GOBP_NUCLEAR_EXPORT" [color="#F1948A"];
"gene_NUP133" -- "term_2_GOBP_NUCLEAR_EXPORT" [color="#F1948A"];
"gene_NUP153" -- "term_2_GOBP_NUCLEAR_EXPORT" [color="#F1948A"];
"gene_NUP155" -- "term_2_GOBP_NUCLEAR_EXPORT" [color="#F1948A"];
"gene_NUP160" -- "term_2_GOBP_NUCLEAR_EXPORT" [color="#F1948A"];
"gene_NUP85" -- "term_2_GOBP_NUCLEAR_EXPORT" [color="#F1948A"];
"gene_PIGH" -- "term_4_GOBP_GLYCOLIPID_BIOSYNTHETIC_PROCESS" [color="#F1948A"];
"gene_PIGK" -- "term_4_GOBP_GLYCOLIPID_BIOSYNTHETIC_PROCESS" [color="#F1948A"];
"gene_PIGY" -- "term_4_GOBP_GLYCOLIPID_BIOSYNTHETIC_PROCESS" [color="#F1948A"];
"gene_POLR3A" -- "term_10_GOBP_POSITIVE_REGULATION_OF_INTERFERON_BETA_PRODUCTION" [color="#F1948A"];
"gene_POLR3F" -- "term_10_GOBP_POSITIVE_REGULATION_OF_INTERFERON_BETA_PRODUCTION" [color="#F1948A"];
"gene_PPIB" -- "term_7_GOBP_POSITIVE_REGULATION_BY_HOST_OF_VIRAL_PROCESS" [color="#F1948A"];
"gene_RAB1B" -- "term_1_GOBP_GOLGI_VESICLE_TRANSPORT" [color="#F1948A"];
"gene_RAB2A" -- "term_1_GOBP_GOLGI_VESICLE_TRANSPORT" [color="#F1948A"];
"gene_RAB6A" -- "term_1_GOBP_GOLGI_VESICLE_TRANSPORT" [color="#F1948A"];
"gene_RAB6B" -- "term_1_GOBP_GOLGI_VESICLE_TRANSPORT" [color="#F1948A"];
"gene_RAB6C" -- "term_1_GOBP_GOLGI_VESICLE_TRANSPORT" [color="#F1948A"];
"gene_RANBP2" -- "term_2_GOBP_NUCLEAR_EXPORT" [color="#F1948A"];
"gene_RAPGEF1" -- "term_6_GOBP_POSITIVE_REGULATION_OF_MOLECULAR_FUNCTION" [color="#F1948A"];
"gene_RELA" -- "term_6_GOBP_POSITIVE_REGULATION_OF_MOLECULAR_FUNCTION" [color="#F1948A"];
"gene_RNF170" -- "term_8_GOBP_PROTEIN_POLYUBIQUITINATION" [color="#F1948A"];
"gene_RNF26" -- "term_8_GOBP_PROTEIN_POLYUBIQUITINATION" [color="#F1948A"];
"gene_SCFD1" -- "term_1_GOBP_GOLGI_VESICLE_TRANSPORT" [color="#F1948A"];
"gene_SPAST" -- "term_1_GOBP_GOLGI_VESICLE_TRANSPORT" [color="#F1948A"];
"gene_SPTAN1" -- "term_5_GOBP_REGULATION_OF_PROTEIN_CONTAINING_COMPLEX_DISASSEMBLY" [color="#F1948A"];
"gene_SPTBN1" -- "term_1_GOBP_GOLGI_VESICLE_TRANSPORT" [color="#F1948A"];
"gene_ST3GAL5" -- "term_4_GOBP_GLYCOLIPID_BIOSYNTHETIC_PROCESS" [color="#F1948A"];
"gene_STAC2" -- "term_6_GOBP_POSITIVE_REGULATION_OF_MOLECULAR_FUNCTION" [color="#F1948A"];
"gene_STT3A" -- "term_11_GOBP_PROTEIN_N_LINKED_GLYCOSYLATION_VIA_ASPARAGINE" [color="#F1948A"];
"gene_STX5" -- "term_1_GOBP_GOLGI_VESICLE_TRANSPORT" [color="#F1948A"];
"gene_TAOK1" -- "term_5_GOBP_REGULATION_OF_PROTEIN_CONTAINING_COMPLEX_DISASSEMBLY" [color="#F1948A"];
"gene_THOC2" -- "term_2_GOBP_NUCLEAR_EXPORT" [color="#F1948A"];
"gene_TM9SF2" -- "term_4_GOBP_GLYCOLIPID_BIOSYNTHETIC_PROCESS" [color="#F1948A"];
"gene_TMED2" -- "term_1_GOBP_GOLGI_VESICLE_TRANSPORT" [color="#F1948A"];
"gene_TOMM70" -- "term_10_GOBP_POSITIVE_REGULATION_OF_INTERFERON_BETA_PRODUCTION" [color="#F1948A"];
"gene_TRAPPC1" -- "term_1_GOBP_GOLGI_VESICLE_TRANSPORT" [color="#F1948A"];
"gene_TRAPPC8" -- "term_1_GOBP_GOLGI_VESICLE_TRANSPORT" [color="#F1948A"];
"gene_TRIM55" -- "term_8_GOBP_PROTEIN_POLYUBIQUITINATION" [color="#F1948A"];
"gene_TRIM58" -- "term_8_GOBP_PROTEIN_POLYUBIQUITINATION" [color="#F1948A"];
"gene_UBQLN4" -- "term_5_GOBP_REGULATION_OF_PROTEIN_CONTAINING_COMPLEX_DISASSEMBLY" [color="#F1948A"];
"gene_WNK1" -- "term_2_GOBP_NUCLEAR_EXPORT" [color="#F1948A"];
"gene_YTHDC2" -- "term_7_GOBP_POSITIVE_REGULATION_BY_HOST_OF_VIRAL_PROCESS" [color="#F1948A"];
"gene_AGBL5" [label="AGBL5", type="gene", width=0.7, height=0.7];
"gene_ANKRD9" [label="ANKRD9", type="gene", width=0.7, height=0.7];
"gene_ATG16L2" [label="ATG16L2", type="gene", width=0.7, height=0.7];
"gene_CAPN6" [label="CAPN6", type="gene", width=0.7, height=0.7];
"gene_CSPP1" [label="CSPP1", type="gene", width=0.7, height=0.7];
"gene_DCAF1" [label="DCAF1", type="gene", width=0.7, height=0.7];
"gene_DEPDC5" [label="DEPDC5", type="gene", width=0.7, height=0.7];
"gene_FBH1" [label="FBH1", type="gene", width=0.7, height=0.7];
"gene_KAT2A" [label="KAT2A", type="gene", width=0.7, height=0.7];
"gene_KLHDC2" [label="KLHDC2", type="gene", width=0.7, height=0.7];
"gene_KLHL1" [label="KLHL1", type="gene", width=0.7, height=0.7];
"gene_KMT5B" [label="KMT5B", type="gene", width=0.7, height=0.7];
"gene_MAP4" [label="MAP4", type="gene", width=0.7, height=0.7];
"gene_PDIA6" [label="PDIA6", type="gene", width=0.7, height=0.7];
"gene_RGPD5" [label="RGPD5", type="gene", width=0.7, height=0.7];
"gene_RGPD8" [label="RGPD8", type="gene", width=0.7, height=0.7];
"gene_TFDP2" [label="TFDP2", type="gene", width=0.7, height=0.7];
"gene_WDTC1" [label="WDTC1", type="gene", width=0.7, height=0.7];
"term_10_GOCC_SPINDLE" [label="GOCC: SPINDLE", style=filled, fillcolor="#85C1E9", fontcolor="white", type="term", width=2, height=2, library="C5: Gene Ontology: Cellular Component"];
"term_11_GOCC_AUTOPHAGOSOME" [label="GOCC: AUTOPHAGOSOME", style=filled, fillcolor="#85C1E9", fontcolor="white", type="term", width=2, height=2, library="C5: Gene Ontology: Cellular Component"];
"term_1_GOCC_NUCLEAR_PORE_OUTER_RING" [label="GOCC: NUCLEAR PORE OUTER RING", style=filled, fillcolor="#85C1E9", fontcolor="white", type="term", width=2, height=2, library="C5: Gene Ontology: Cellular Component"];
"term_2_GOCC_CORE_MEDIATOR_COMPLEX" [label="GOCC: CORE MEDIATOR COMPLEX", style=filled, fillcolor="#85C1E9", fontcolor="white", type="term", width=2, height=2, library="C5: Gene Ontology: Cellular Component"];
"term_3_GOCC_GOLGI_TRANSPORT_COMPLEX" [label="GOCC: GOLGI TRANSPORT COMPLEX", style=filled, fillcolor="#85C1E9", fontcolor="white", type="term", width=2, height=2, library="C5: Gene Ontology: Cellular Component"];
"term_4_GOCC_ENDOPLASMIC_RETICULUM_PROTEIN_CONTAINING_COMPLEX" [label="GOCC: ENDOPLASMIC RETICULUM PROTEIN CONTAINING COMPLEX", style=filled, fillcolor="#85C1E9", fontcolor="white", type="term", width=2, height=2, library="C5: Gene Ontology: Cellular Component"];
"term_5_GOCC_ENDOPLASMIC_RETICULUM_GOLGI_INTERMEDIATE_COMPARTMENT_MEMBRANE" [label="GOCC: ENDOPLASMIC RETICULUM GOLGI INTERMEDIATE COMPARTMENT MEMBRANE", style=filled, fillcolor="#85C1E9", fontcolor="white", type="term", width=2, height=2, library="C5: Gene Ontology: Cellular Component"];
"term_6_GOCC_NUCLEAR_EXOSOME_RNASE_COMPLEX" [label="GOCC: NUCLEAR EXOSOME RNASE COMPLEX", style=filled, fillcolor="#85C1E9", fontcolor="white", type="term", width=2, height=2, library="C5: Gene Ontology: Cellular Component"];
"term_7_GOCC_CULLIN_RING_UBIQUITIN_LIGASE_COMPLEX" [label="GOCC: CULLIN RING UBIQUITIN LIGASE COMPLEX", style=filled, fillcolor="#85C1E9", fontcolor="white", type="term", width=2, height=2, library="C5: Gene Ontology: Cellular Component"];
"term_8_GOCC_NUCLEAR_PORE" [label="GOCC: NUCLEAR PORE", style=filled, fillcolor="#85C1E9", fontcolor="white", type="term", width=2, height=2, library="C5: Gene Ontology: Cellular Component"];
"term_9_GOCC_CLATHRIN_COATED_VESICLE_MEMBRANE" [label="GOCC: CLATHRIN COATED VESICLE MEMBRANE", style=filled, fillcolor="#85C1E9", fontcolor="white", type="term", width=2, height=2, library="C5: Gene Ontology: Cellular Component"];
"gene_AGBL5" -- "term_10_GOCC_SPINDLE" [color="#85C1E9"];
"gene_AKT1" -- "term_10_GOCC_SPINDLE" [color="#85C1E9"];
"gene_ANKRD9" -- "term_7_GOCC_CULLIN_RING_UBIQUITIN_LIGASE_COMPLEX" [color="#85C1E9"];
"gene_AP2M1" -- "term_9_GOCC_CLATHRIN_COATED_VESICLE_MEMBRANE" [color="#85C1E9"];
"gene_ATG12" -- "term_11_GOCC_AUTOPHAGOSOME" [color="#85C1E9"];
"gene_ATG16L2" -- "term_11_GOCC_AUTOPHAGOSOME" [color="#85C1E9"];
"gene_ATP6V0A1" -- "term_9_GOCC_CLATHRIN_COATED_VESICLE_MEMBRANE" [color="#85C1E9"];
"gene_CAPN6" -- "term_10_GOCC_SPINDLE" [color="#85C1E9"];
"gene_CD4" -- "term_9_GOCC_CLATHRIN_COATED_VESICLE_MEMBRANE" [color="#85C1E9"];
"gene_CLN3" -- "term_11_GOCC_AUTOPHAGOSOME" [color="#85C1E9"];
"gene_COG2" -- "term_3_GOCC_GOLGI_TRANSPORT_COMPLEX" [color="#85C1E9"];
"gene_COG3" -- "term_3_GOCC_GOLGI_TRANSPORT_COMPLEX" [color="#85C1E9"];
"gene_COG4" -- "term_3_GOCC_GOLGI_TRANSPORT_COMPLEX" [color="#85C1E9"];
"gene_CSPP1" -- "term_10_GOCC_SPINDLE" [color="#85C1E9"];
"gene_CTDP1" -- "term_10_GOCC_SPINDLE" [color="#85C1E9"];
"gene_DCAF1" -- "term_7_GOCC_CULLIN_RING_UBIQUITIN_LIGASE_COMPLEX" [color="#85C1E9"];
"gene_DDOST" -- "term_4_GOCC_ENDOPLASMIC_RETICULUM_PROTEIN_CONTAINING_COMPLEX" [color="#85C1E9"];
"gene_DEPDC5" -- "term_7_GOCC_CULLIN_RING_UBIQUITIN_LIGASE_COMPLEX" [color="#85C1E9"];
"gene_EGF" -- "term_9_GOCC_CLATHRIN_COATED_VESICLE_MEMBRANE" [color="#85C1E9"];
"gene_EGFR" -- "term_9_GOCC_CLATHRIN_COATED_VESICLE_MEMBRANE" [color="#85C1E9"];
"gene_EXOSC3" -- "term_6_GOCC_NUCLEAR_EXOSOME_RNASE_COMPLEX" [color="#85C1E9"];
"gene_EXOSC5" -- "term_6_GOCC_NUCLEAR_EXOSOME_RNASE_COMPLEX" [color="#85C1E9"];
"gene_FBH1" -- "term_7_GOCC_CULLIN_RING_UBIQUITIN_LIGASE_COMPLEX" [color="#85C1E9"];
"gene_FBXW11" -- "term_7_GOCC_CULLIN_RING_UBIQUITIN_LIGASE_COMPLEX" [color="#85C1E9"];
"gene_GABARAPL2" -- "term_11_GOCC_AUTOPHAGOSOME" [color="#85C1E9"];
"gene_GOSR2" -- "term_5_GOCC_ENDOPLASMIC_RETICULUM_GOLGI_INTERMEDIATE_COMPARTMENT_MEMBRANE" [color="#85C1E9"];
"gene_HIP1R" -- "term_9_GOCC_CLATHRIN_COATED_VESICLE_MEMBRANE" [color="#85C1E9"];
"gene_IKBKG" -- "term_10_GOCC_SPINDLE" [color="#85C1E9"];
"gene_KAT2A" -- "term_10_GOCC_SPINDLE" [color="#85C1E9"];
```

```
"gene_KBTBD7" -- "term_7_GOCC_CULLIN_RING_UBIQUITIN_LIGASE_COMPLEX" [color="#85C1E9"];
"gene_KLHDC2" -- "term_7_GOCC_CULLIN_RING_UBIQUITIN_LIGASE_COMPLEX" [color="#85C1E9"];
"gene_KLHL1" -- "term_7_GOCC_CULLIN_RING_UBIQUITIN_LIGASE_COMPLEX" [color="#85C1E9"];
"gene_KMT5B" -- "term_10_GOCC_SPINDLE" [color="#85C1E9"];
"gene_MAP4" -- "term_10_GOCC_SPINDLE" [color="#85C1E9"];
"gene_MED14" -- "term_2_GOCC_CORE_MEDIATOR_COMPLEX" [color="#85C1E9"];
"gene_MED28" -- "term_2_GOCC_CORE_MEDIATOR_COMPLEX" [color="#85C1E9"];
"gene_MED4" -- "term_2_GOCC_CORE_MEDIATOR_COMPLEX" [color="#85C1E9"];
"gene_MED6" -- "term_2_GOCC_CORE_MEDIATOR_COMPLEX" [color="#85C1E9"];
"gene_MED7" -- "term_2_GOCC_CORE_MEDIATOR_COMPLEX" [color="#85C1E9"];
"gene_MGAT1" -- "term_5_GOCC_ENDOPLASMIC_RETICULUM_GOLGI_INTERMEDIATE_COMPARTMENT_MEMBRANE" [color="#85C1E9"];
"gene_MPHOSPH6" -- "term_6_GOCC_NUCLEAR_EXOSOME_RNASE_COMPLEX" [color="#85C1E9"];
"gene_NUP107" -- "term_1_GOCC_NUCLEAR_PORE_OUTER_RING" [color="#85C1E9"];
"gene_NUP133" -- "term_1_GOCC_NUCLEAR_PORE_OUTER_RING" [color="#85C1E9"];
"gene_NUP153" -- "term_8_GOCC_NUCLEAR_PORE" [color="#85C1E9"];
"gene_NUP155" -- "term_8_GOCC_NUCLEAR_PORE" [color="#85C1E9"];
"gene_NUP160" -- "term_1_GOCC_NUCLEAR_PORE_OUTER_RING" [color="#85C1E9"];
"gene_NUP85" -- "term_1_GOCC_NUCLEAR_PORE_OUTER_RING" [color="#85C1E9"];
"gene_PDIA6" -- "term_4_GOCC_ENDOPLASMIC_RETICULUM_PROTEIN_CONTAINING_COMPLEX" [color="#85C1E9"];
"gene_PIGH" -- "term_4_GOCC_ENDOPLASMIC_RETICULUM_PROTEIN_CONTAINING_COMPLEX" [color="#85C1E9"];
"gene_PIGK" -- "term_4_GOCC_ENDOPLASMIC_RETICULUM_PROTEIN_CONTAINING_COMPLEX" [color="#85C1E9"];
"gene_PIGY" -- "term_4_GOCC_ENDOPLASMIC_RETICULUM_PROTEIN_CONTAINING_COMPLEX" [color="#85C1E9"];
"gene_PPIB" -- "term_4_GOCC_ENDOPLASMIC_RETICULUM_PROTEIN_CONTAINING_COMPLEX" [color="#85C1E9"];
"gene_RAB1B" -- "term_5_GOCC_ENDOPLASMIC_RETICULUM_GOLGI_INTERMEDIATE_COMPARTMENT_MEMBRANE" [color="#85C1E9"];
"gene_RAB2A" -- "term_5_GOCC_ENDOPLASMIC_RETICULUM_GOLGI_INTERMEDIATE_COMPARTMENT_MEMBRANE" [color="#85C1E9"];
"gene_RANBP2" -- "term_8_GOCC_NUCLEAR_PORE" [color="#85C1E9"];
"gene_RGPD5" -- "term_8_GOCC_NUCLEAR_PORE" [color="#85C1E9"];
"gene_RGPD8" -- "term_8_GOCC_NUCLEAR_PORE" [color="#85C1E9"];
"gene_SPAST" -- "term_10_GOCC_SPINDLE" [color="#85C1E9"];
"gene_SPCS3" -- "term_4_GOCC_ENDOPLASMIC_RETICULUM_PROTEIN_CONTAINING_COMPLEX" [color="#85C1E9"];
"gene_STT3A" -- "term_4_GOCC_ENDOPLASMIC_RETICULUM_PROTEIN_CONTAINING_COMPLEX" [color="#85C1E9"];
"gene_STX5" -- "term_5_GOCC_ENDOPLASMIC_RETICULUM_GOLGI_INTERMEDIATE_COMPARTMENT_MEMBRANE" [color="#85C1E9"];
"gene_TFDP2" -- "term_10_GOCC_SPINDLE" [color="#85C1E9"];
"gene_TMED2" -- "term_5_GOCC_ENDOPLASMIC_RETICULUM_GOLGI_INTERMEDIATE_COMPARTMENT_MEMBRANE" [color="#85C1E9"];
"gene_UBQLN4" -- "term_11_GOCC_AUTOPHAGOSOME" [color="#85C1E9"];
"gene_WDTC1" -- "term_7_GOCC_CULLIN_RING_UBIQUITIN_LIGASE_COMPLEX" [color="#85C1E9"];
"gene_WNK1" -- "term_10_GOCC_SPINDLE" [color="#85C1E9"];
"gene_ALKBH8" [label="ALKBH8", type="gene", width=0.7, height=0.7];
"gene_ARGLU1" [label="ARGLU1", type="gene", width=0.7, height=0.7];
"gene_ARHGAP32" [label="ARHGAP32", type="gene", width=0.7, height=0.7];
"gene_ARHGEF12" [label="ARHGEF12", type="gene", width=0.7, height=0.7];
"gene_ARHGEF19" [label="ARHGEF19", type="gene", width=0.7, height=0.7];
"gene_BCL9" [label="BCL9", type="gene", width=0.7, height=0.7];
"gene_CRTC3" [label="CRTC3", type="gene", width=0.7, height=0.7];
"gene_DDX53" [label="DDX53", type="gene", width=0.7, height=0.7];
"gene_DDX55" [label="DDX55", type="gene", width=0.7, height=0.7];
"gene_DNAL1" [label="DNAL1", type="gene", width=0.7, height=0.7];
"gene_ERI2" [label="ERI2", type="gene", width=0.7, height=0.7];
"gene_FGD6" [label="FGD6", type="gene", width=0.7, height=0.7];
"gene_FHL3" [label="FHL3", type="gene", width=0.7, height=0.7];
"gene_FNTA" [label="FNTA", type="gene", width=0.7, height=0.7];
"gene_GRTP1" [label="GRTP1", type="gene", width=0.7, height=0.7];
"gene_IGHMBP2" [label="IGHMBP2", type="gene", width=0.7, height=0.7];
"gene_ITPKA" [label="ITPKA", type="gene", width=0.7, height=0.7];
"gene_KDM7A" [label="KDM7A", type="gene", width=0.7, height=0.7];
"gene_NR0B2" [label="NR0B2", type="gene", width=0.7, height=0.7];
"gene_PHF12" [label="PHF12", type="gene", width=0.7, height=0.7];
"gene_PLOD3" [label="PLOD3", type="gene", width=0.7, height=0.7];
"gene_PPP2R2A" [label="PPP2R2A", type="gene", width=0.7, height=0.7];
"gene_RAB28" [label="RAB28", type="gene", width=0.7, height=0.7];
"gene_RIMS4" [label="RIMS4", type="gene", width=0.7, height=0.7];
"gene_RRAGB" [label="RRAGB", type="gene", width=0.7, height=0.7];
"gene_RUSC2" [label="RUSC2", type="gene", width=0.7, height=0.7];
"gene_TIAM2" [label="TIAM2", type="gene", width=0.7, height=0.7];
"gene_TMTC1" [label="TMTC1", type="gene", width=0.7, height=0.7];
"gene_TNPO3" [label="TNPO3", type="gene", width=0.7, height=0.7];
"gene_TRMT5" [label="TRMT5", type="gene", width=0.7, height=0.7];
"term_10_GOMF_GTPASE_BINDING" [label="GOMF: GTPASE BINDING", style=filled, fillcolor="#D7BDE2", fontcolor="white", type="term", width=2, height=2, library="C5: Gene Ontology: Molecular Function"];
"term_1_GOMF_STRUCTURAL_CONSTITUENT_OF_NUCLEAR_PORE" [label="GOMF: STRUCTURAL CONSTITUENT OF NUCLEAR PORE", style=filled, fillcolor="#D7BDE2", fontcolor="white", type="term", width=2, height=2, library="C5: Gene Ontology: Molecular
Function"];
"term_2_GOMF_CATALYTIC_ACTIVITY_ACTING_ON_RNA" [label="GOMF: CATALYTIC ACTIVITY ACTING ON RNA", style=filled, fillcolor="#D7BDE2", fontcolor="white", type="term", width=2, height=2, library="C5: Gene Ontology: Molecular Function"];
"term_3_GOMF_GTPASE_ACTIVATOR_ACTIVITY" [label="GOMF: GTPASE ACTIVATOR ACTIVITY", style=filled, fillcolor="#D7BDE2", fontcolor="white", type="term", width=2, height=2, library="C5: Gene Ontology: Molecular Function"];
"term_4_GOMF_TRANSCRIPTION_COREGULATOR_ACTIVITY" [label="GOMF: TRANSCRIPTION COREGULATOR ACTIVITY", style=filled, fillcolor="#D7BDE2", fontcolor="white", type="term", width=2, height=2, library="C5: Gene Ontology: Molecular Function"];
"term_5_GOMF_ENZYME_SUBSTRATE_ADAPTOR_ACTIVITY" [label="GOMF: ENZYME SUBSTRATE ADAPTOR ACTIVITY", style=filled, fillcolor="#D7BDE2", fontcolor="white", type="term", width=2, height=2, library="C5: Gene Ontology: Molecular Function"];
"term_6_GOMF_ALPHA_TUBULIN_BINDING" [label="GOMF: ALPHA TUBULIN BINDING", style=filled, fillcolor="#D7BDE2", fontcolor="white", type="term", width=2, height=2, library="C5: Gene Ontology: Molecular Function"];
"term_7_GOMF_CARGO_ADAPTOR_ACTIVITY" [label="GOMF: CARGO ADAPTOR ACTIVITY", style=filled, fillcolor="#D7BDE2", fontcolor="white", type="term", width=2, height=2, library="C5: Gene Ontology: Molecular Function"];
"term_8_GOMF_HEXOSYLTRANSFERASE_ACTIVITY" [label="GOMF: HEXOSYLTRANSFERASE ACTIVITY", style=filled, fillcolor="#D7BDE2", fontcolor="white", type="term", width=2, height=2, library="C5: Gene Ontology: Molecular Function"];
"term_9_GOMF_G_PROTEIN_ACTIVITY" [label="GOMF: G PROTEIN ACTIVITY", style=filled, fillcolor="#D7BDE2", fontcolor="white", type="term", width=2, height=2, library="C5: Gene Ontology: Molecular Function"];
"gene_A4GALT" -- "term_8_GOMF_HEXOSYLTRANSFERASE_ACTIVITY" [color="#D7BDE2"];
"gene_AGAP2" -- "term_3_GOMF_GTPASE_ACTIVATOR_ACTIVITY" [color="#D7BDE2"];
"gene_AGO3" -- "term_2_GOMF_CATALYTIC_ACTIVITY_ACTING_ON_RNA" [color="#D7BDE2"];
"gene_ALKBH8" -- "term_2_GOMF_CATALYTIC_ACTIVITY_ACTING_ON_RNA" [color="#D7BDE2"];
"gene_ANKRD9" -- "term_5_GOMF_ENZYME_SUBSTRATE_ADAPTOR_ACTIVITY" [color="#D7BDE2"];
"gene_AP2M1" -- "term_7_GOMF_CARGO_ADAPTOR_ACTIVITY" [color="#D7BDE2"];
"gene_ARGLU1" -- "term_4_GOMF_TRANSCRIPTION_COREGULATOR_ACTIVITY" [color="#D7BDE2"];
"gene_ARHGAP32" -- "term_3_GOMF_GTPASE_ACTIVATOR_ACTIVITY" [color="#D7BDE2"];
"gene_ARHGEF12" -- "term_3_GOMF_GTPASE_ACTIVATOR_ACTIVITY" [color="#D7BDE2"];
"gene_ARHGEF19" -- "term_3_GOMF_GTPASE_ACTIVATOR_ACTIVITY" [color="#D7BDE2"];
"gene_BCL9" -- "term_4_GOMF_TRANSCRIPTION_COREGULATOR_ACTIVITY" [color="#D7BDE2"];
"gene_CRTC2" -- "term_4_GOMF_TRANSCRIPTION_COREGULATOR_ACTIVITY" [color="#D7BDE2"];
"gene_CRTC3" -- "term_4_GOMF_TRANSCRIPTION_COREGULATOR_ACTIVITY" [color="#D7BDE2"];
"gene_DCAF1" -- "term_5_GOMF_ENZYME_SUBSTRATE_ADAPTOR_ACTIVITY" [color="#D7BDE2"];
"gene_DDX10" -- "term_2_GOMF_CATALYTIC_ACTIVITY_ACTING_ON_RNA" [color="#D7BDE2"];
"gene_DDX53" -- "term_2_GOMF_CATALYTIC_ACTIVITY_ACTING_ON_RNA" [color="#D7BDE2"];
"gene_DDX55" -- "term_2_GOMF_CATALYTIC_ACTIVITY_ACTING_ON_RNA" [color="#D7BDE2"];
"gene_DEPDC5" -- "term_3_GOMF_GTPASE_ACTIVATOR_ACTIVITY" [color="#D7BDE2"];
"gene_DHX33" -- "term_2_GOMF_CATALYTIC_ACTIVITY_ACTING_ON_RNA" [color="#D7BDE2"];
"gene_DIMT1" -- "term_2_GOMF_CATALYTIC_ACTIVITY_ACTING_ON_RNA" [color="#D7BDE2"];
"gene_DNAJB1" -- "term_4_GOMF_TRANSCRIPTION_COREGULATOR_ACTIVITY" [color="#D7BDE2"];
"gene_DNAL1" -- "term_6_GOMF_ALPHA_TUBULIN_BINDING" [color="#D7BDE2"];
"gene_DPM1" -- "term_8_GOMF_HEXOSYLTRANSFERASE_ACTIVITY" [color="#D7BDE2"];
"gene_EPS8" -- "term_10_GOMF_GTPASE_BINDING" [color="#D7BDE2"];
"gene_ERI2" -- "term_2_GOMF_CATALYTIC_ACTIVITY_ACTING_ON_RNA" [color="#D7BDE2"];
"gene_ETF1" -- "term_2_GOMF_CATALYTIC_ACTIVITY_ACTING_ON_RNA" [color="#D7BDE2"];
"gene_EXOSC3" -- "term_2_GOMF_CATALYTIC_ACTIVITY_ACTING_ON_RNA" [color="#D7BDE2"];
"gene_EXOSC5" -- "term_2_GOMF_CATALYTIC_ACTIVITY_ACTING_ON_RNA" [color="#D7BDE2"];
"gene_FBXW11" -- "term_5_GOMF_ENZYME_SUBSTRATE_ADAPTOR_ACTIVITY" [color="#D7BDE2"];
"gene_FGD6" -- "term_10_GOMF_GTPASE_BINDING" [color="#D7BDE2"];
"gene_FHL3" -- "term_4_GOMF_TRANSCRIPTION_COREGULATOR_ACTIVITY" [color="#D7BDE2"];
"gene_FNTA" -- "term_6_GOMF_ALPHA_TUBULIN_BINDING" [color="#D7BDE2"];
"gene_GAPVD1" -- "term_3_GOMF_GTPASE_ACTIVATOR_ACTIVITY" [color="#D7BDE2"];
"gene_GOLPH3" -- "term_7_GOMF_CARGO_ADAPTOR_ACTIVITY" [color="#D7BDE2"];
"gene_GRTP1" -- "term_3_GOMF_GTPASE_ACTIVATOR_ACTIVITY" [color="#D7BDE2"];
"gene_HIP1R" -- "term_7_GOMF_CARGO_ADAPTOR_ACTIVITY" [color="#D7BDE2"];
"gene_IGHMBP2" -- "term_2_GOMF_CATALYTIC_ACTIVITY_ACTING_ON_RNA" [color="#D7BDE2"];
"gene_ITPKA" -- "term_10_GOMF_GTPASE_BINDING" [color="#D7BDE2"];
"gene_KAT2A" -- "term_4_GOMF_TRANSCRIPTION_COREGULATOR_ACTIVITY" [color="#D7BDE2"];
"gene_KBTBD7" -- "term_5_GOMF_ENZYME_SUBSTRATE_ADAPTOR_ACTIVITY" [color="#D7BDE2"];
"gene_KDM7A" -- "term_4_GOMF_TRANSCRIPTION_COREGULATOR_ACTIVITY" [color="#D7BDE2"];
"gene_KLHDC2" -- "term_5_GOMF_ENZYME_SUBSTRATE_ADAPTOR_ACTIVITY" [color="#D7BDE2"];
"gene_KLHL1" -- "term_5_GOMF_ENZYME_SUBSTRATE_ADAPTOR_ACTIVITY" [color="#D7BDE2"];
"gene_LARS1" -- "term_2_GOMF_CATALYTIC_ACTIVITY_ACTING_ON_RNA" [color="#D7BDE2"];
"gene_MED14" -- "term_4_GOMF_TRANSCRIPTION_COREGULATOR_ACTIVITY" [color="#D7BDE2"];
"gene_MED4" -- "term_4_GOMF_TRANSCRIPTION_COREGULATOR_ACTIVITY" [color="#D7BDE2"];
"gene_MED6" -- "term_4_GOMF_TRANSCRIPTION_COREGULATOR_ACTIVITY" [color="#D7BDE2"];
"gene_MED7" -- "term_4_GOMF_TRANSCRIPTION_COREGULATOR_ACTIVITY" [color="#D7BDE2"];
"gene_MGAT1" -- "term_8_GOMF_HEXOSYLTRANSFERASE_ACTIVITY" [color="#D7BDE2"];
"gene_NCOR2" -- "term_4_GOMF_TRANSCRIPTION_COREGULATOR_ACTIVITY" [color="#D7BDE2"];
"gene_NR0B2" -- "term_4_GOMF_TRANSCRIPTION_COREGULATOR_ACTIVITY" [color="#D7BDE2"];
"gene_NUP107" -- "term_1_GOMF_STRUCTURAL_CONSTITUENT_OF_NUCLEAR_PORE" [color="#D7BDE2"];
"gene_NUP133" -- "term_1_GOMF_STRUCTURAL_CONSTITUENT_OF_NUCLEAR_PORE" [color="#D7BDE2"];
"gene_NUP153" -- "term_1_GOMF_STRUCTURAL_CONSTITUENT_OF_NUCLEAR_PORE" [color="#D7BDE2"];
"gene_NUP155" -- "term_1_GOMF_STRUCTURAL_CONSTITUENT_OF_NUCLEAR_PORE" [color="#D7BDE2"];
"gene_NUP160" -- "term_1_GOMF_STRUCTURAL_CONSTITUENT_OF_NUCLEAR_PORE" [color="#D7BDE2"];
"gene_NUP85" -- "term_1_GOMF_STRUCTURAL_CONSTITUENT_OF_NUCLEAR_PORE" [color="#D7BDE2"];
"gene_PHF12" -- "term_4_GOMF_TRANSCRIPTION_COREGULATOR_ACTIVITY" [color="#D7BDE2"];
"gene_PIGY" -- "term_8_GOMF_HEXOSYLTRANSFERASE_ACTIVITY" [color="#D7BDE2"];
"gene_PLOD3" -- "term_8_GOMF_HEXOSYLTRANSFERASE_ACTIVITY" [color="#D7BDE2"];
"gene_POLR1H" -- "term_2_GOMF_CATALYTIC_ACTIVITY_ACTING_ON_RNA" [color="#D7BDE2"];
"gene_POLR3A" -- "term_2_GOMF_CATALYTIC_ACTIVITY_ACTING_ON_RNA" [color="#D7BDE2"];
```

"gene_POLR3F" -- "term_2_GOMF_CATALYTIC_ACTIVITY_ACTING_ON_RNA" [color="#D7BDE2"];
"gene_PPP2R2A" -- "term_5_GOMF_ENZYME_SUBSTRATE_ADAPTOR_ACTIVITY" [color="#D7BDE2"];
"gene_RAB1B" -- "term_9_GOMF_G_PROTEIN_ACTIVITY" [color="#D7BDE2"];
"gene_RAB28" -- "term_9_GOMF_G_PROTEIN_ACTIVITY" [color="#D7BDE2"];
"gene_RAB2A" -- "term_9_GOMF_G_PROTEIN_ACTIVITY" [color="#D7BDE2"];
"gene_RANBP2" -- "term_3_GOMF_GTPASE_ACTIVATOR_ACTIVITY" [color="#D7BDE2"];
"gene_RAP1B" -- "term_9_GOMF_G_PROTEIN_ACTIVITY" [color="#D7BDE2"];
"gene_RGPD5" -- "term_3_GOMF_GTPASE_ACTIVATOR_ACTIVITY" [color="#D7BDE2"];
"gene_RGPD8" -- "term_3_GOMF_GTPASE_ACTIVATOR_ACTIVITY" [color="#D7BDE2"];
"gene_RIMS4" -- "term_10_GOMF_GTPASE_BINDING" [color="#D7BDE2"];
"gene_RRAGB" -- "term_10_GOMF_GTPASE_BINDING" [color="#D7BDE2"];
"gene_RUSC2" -- "term_10_GOMF_GTPASE_BINDING" [color="#D7BDE2"];
"gene_SPAST" -- "term_6_GOMF_ALPHA_TUBULIN_BINDING" [color="#D7BDE2"];
"gene_SPTBN1" -- "term_10_GOMF_GTPASE_BINDING" [color="#D7BDE2"];
"gene_STT3A" -- "term_8_GOMF_HEXOSYLTRANSFERASE_ACTIVITY" [color="#D7BDE2"];
"gene_TAOK1" -- "term_6_GOMF_ALPHA_TUBULIN_BINDING" [color="#D7BDE2"];
"gene_TIAM2" -- "term_3_GOMF_GTPASE_ACTIVATOR_ACTIVITY" [color="#D7BDE2"];
"gene_TMTC1" -- "term_8_GOMF_HEXOSYLTRANSFERASE_ACTIVITY" [color="#D7BDE2"];
"gene_TNPO3" -- "term_10_GOMF_GTPASE_BINDING" [color="#D7BDE2"];
"gene_TRMT5" -- "term_2_GOMF_CATALYTIC_ACTIVITY_ACTING_ON_RNA" [color="#D7BDE2"];
"gene_YTHDC2" -- "term_2_GOMF_CATALYTIC_ACTIVITY_ACTING_ON_RNA" [color="#D7BDE2"];
"gene_ASXL2" [label="ASXL2", type="gene", width=0.7, height=0.7];
"gene_CXCR4" [label="CXCR4", type="gene", width=0.7, height=0.7];
"gene_H3_3A" [label="H3-3A", type="gene", width=0.7, height=0.7];
"gene_LRIF1" [label="LRIF1", type="gene", width=0.7, height=0.7];
"gene_POU1F1" [label="POU1F1", type="gene", width=0.7, height=0.7];
"gene_SP110" [label="SP110", type="gene", width=0.7, height=0.7];
"gene_TIMM8A" [label="TIMM8A", type="gene", width=0.7, height=0.7];
"gene_TMEM163" [label="TMEM163", type="gene", width=0.7, height=0.7];
"term_1_HP_FOAMY_URINE" [label="HP: FOAMY URINE", style=filled, fillcolor="#F7DC6F", fontcolor="white", type="term", width=2, height=2, library="C5: Human Phenotype Ontology"];
"term_2_HP_ABNORMAL_GLYCOSYLATION" [label="HP: ABNORMAL GLYCOSYLATION", style=filled, fillcolor="#F7DC6F", fontcolor="white", type="term", width=2, height=2, library="C5: Human Phenotype Ontology"];
"term_3_HP_DILATION_OF_VIRCHOW_ROBIN_SPACES" [label="HP: DILATION OF VIRCHOW ROBIN SPACES", style=filled, fillcolor="#F7DC6F", fontcolor="white", type="term", width=2, height=2, library="C5: Human Phenotype Ontology"];
"term_4_HP_OVERWEIGHT" [label="HP: OVERWEIGHT", style=filled, fillcolor="#F7DC6F", fontcolor="white", type="term", width=2, height=2, library="C5: Human Phenotype Ontology"];
"term_5_HP_PAPILLOMA" [label="HP: PAPILLOMA", style=filled, fillcolor="#F7DC6F", fontcolor="white", type="term", width=2, height=2, library="C5: Human Phenotype Ontology"];
"term_6_HP_ASPIRATION_PNEUMONIA" [label="HP: ASPIRATION PNEUMONIA", style=filled, fillcolor="#F7DC6F", fontcolor="white", type="term", width=2, height=2, library="C5: Human Phenotype Ontology"];
"term_7_HP_EEG_WITH_FOCAL_SPIKES" [label="HP: EEG WITH FOCAL SPIKES", style=filled, fillcolor="#F7DC6F", fontcolor="white", type="term", width=2, height=2, library="C5: Human Phenotype Ontology"];
"term_8_HP_BLUE_SCLERAE" [label="HP: BLUE SCLERAE", style=filled, fillcolor="#F7DC6F", fontcolor="white", type="term", width=2, height=2, library="C5: Human Phenotype Ontology"];
"term_9_HP_CUPPED_EAR" [label="HP: CUPPED EAR", style=filled, fillcolor="#F7DC6F", fontcolor="white", type="term", width=2, height=2, library="C5: Human Phenotype Ontology"];
"gene_AKT1" -- "term_5_HP_PAPILLOMA" [color="#F7DC6F"];
"gene_ASXL2" -- "term_3_HP_DILATION_OF_VIRCHOW_ROBIN_SPACES" [color="#F7DC6F"];
"gene_ATP6V0A1" -- "term_6_HP_ASPIRATION_PNEUMONIA" [color="#F7DC6F"];
"gene_CD4" -- "term_5_HP_PAPILLOMA" [color="#F7DC6F"];
"gene_COG2" -- "term_2_HP_ABNORMAL_GLYCOSYLATION" [color="#F7DC6F"];
"gene_COG3" -- "term_2_HP_ABNORMAL_GLYCOSYLATION" [color="#F7DC6F"];
"gene_COG4" -- "term_2_HP_ABNORMAL_GLYCOSYLATION" [color="#F7DC6F"];
"gene_CSPP1" -- "term_6_HP_ASPIRATION_PNEUMONIA" [color="#F7DC6F"];
"gene_CXCR4" -- "term_5_HP_PAPILLOMA" [color="#F7DC6F"];
"gene_DDOST" -- "term_2_HP_ABNORMAL_GLYCOSYLATION" [color="#F7DC6F"];
"gene_DEPDC5" -- "term_7_HP_EEG_WITH_FOCAL_SPIKES" [color="#F7DC6F"];
"gene_DPM1" -- "term_2_HP_ABNORMAL_GLYCOSYLATION" [color="#F7DC6F"];
"gene_EDNRA" -- "term_9_HP_CUPPED_EAR" [color="#F7DC6F"];
"gene_ERCC3" -- "term_5_HP_PAPILLOMA" [color="#F7DC6F"];
"gene_FBXW11" -- "term_9_HP_CUPPED_EAR" [color="#F7DC6F"];
"gene_GAPVD1" -- "term_1_HP_FOAMY_URINE" [color="#F7DC6F"];
"gene_GCK" -- "term_4_HP_OVERWEIGHT" [color="#F7DC6F"];
"gene_H3_3A" -- "term_3_HP_DILATION_OF_VIRCHOW_ROBIN_SPACES" [color="#F7DC6F"];
"gene_HUWE1" -- "term_9_HP_CUPPED_EAR" [color="#F7DC6F"];
"gene_IKBKG" -- "term_5_HP_PAPILLOMA" [color="#F7DC6F"];
"gene_KMT5B" -- "term_3_HP_DILATION_OF_VIRCHOW_ROBIN_SPACES" [color="#F7DC6F"];
"gene_LRIF1" -- "term_6_HP_ASPIRATION_PNEUMONIA" [color="#F7DC6F"];
"gene_NGLY1" -- "term_3_HP_DILATION_OF_VIRCHOW_ROBIN_SPACES" [color="#F7DC6F"];
"gene_NUP107" -- "term_1_HP_FOAMY_URINE" [color="#F7DC6F"];
"gene_NUP133" -- "term_1_HP_FOAMY_URINE" [color="#F7DC6F"];
"gene_NUP160" -- "term_1_HP_FOAMY_URINE" [color="#F7DC6F"];
"gene_NUP85" -- "term_1_HP_FOAMY_URINE" [color="#F7DC6F"];
"gene_PIGY" -- "term_9_HP_CUPPED_EAR" [color="#F7DC6F"];
"gene_POLR3A" -- "term_8_HP_BLUE_SCLERAE" [color="#F7DC6F"];
"gene_POU1F1" -- "term_4_HP_OVERWEIGHT" [color="#F7DC6F"];
"gene_PPIB" -- "term_8_HP_BLUE_SCLERAE" [color="#F7DC6F"];
"gene_PURA" -- "term_6_HP_ASPIRATION_PNEUMONIA" [color="#F7DC6F"];
"gene_SP110" -- "term_6_HP_ASPIRATION_PNEUMONIA" [color="#F7DC6F"];
"gene_SPTAN1" -- "term_7_HP_EEG_WITH_FOCAL_SPIKES" [color="#F7DC6F"];
"gene_SPTBN1" -- "term_4_HP_OVERWEIGHT" [color="#F7DC6F"];
"gene_STT3A" -- "term_2_HP_ABNORMAL_GLYCOSYLATION" [color="#F7DC6F"];
"gene_TAOK1" -- "term_4_HP_OVERWEIGHT" [color="#F7DC6F"];
"gene_TFE3" -- "term_7_HP_EEG_WITH_FOCAL_SPIKES" [color="#F7DC6F"];
"gene_THOC2" -- "term_4_HP_OVERWEIGHT" [color="#F7DC6F"];
"gene_TIMM8A" -- "term_6_HP_ASPIRATION_PNEUMONIA" [color="#F7DC6F"];
"gene_TMEM163" -- "term_8_HP_BLUE_SCLERAE" [color="#F7DC6F"];
"gene_TRMT5" -- "term_8_HP_BLUE_SCLERAE" [color="#F7DC6F"];
"gene_WNT1" -- "term_8_HP_BLUE_SCLERAE" [color="#F7DC6F"];
}

**NETWORK INTERPRETATION GUIDE:**

**Node Types:**
- **Gene nodes** (type="gene"): Individual genes from the input gene set
- **Term nodes** (type="term"): Enriched biological pathways/processes/gene sets
- **Library Source**: Each term node includes a "library" attribute indicating the source database

**Library Sources and Their Biological Context:**
- **H: Hallmark gene sets**: Curated gene sets representing well-defined biological states or processes
- **C2: BioCarta**: Canonical pathways from BioCarta database
- **C2: KEGG MEDICUS**: Metabolic and signaling pathways from KEGG
- **C2: Pathway Interaction Database**: Curated human signaling pathways
- **C2: Reactome Pathways**: Expert-curated biological pathways
- **C2: WikiPathways**: Community-curated biological pathways
- **C5: Gene Ontology: Biological Process**: Biological processes from Gene Ontology
- **C5: Gene Ontology: Cellular Component**: Cellular components from Gene Ontology
- **C5: Gene Ontology: Molecular Function**: Molecular functions from Gene Ontology
- **C5: Human Phenotype Ontology**: Human phenotypes and diseases
- **Protein Interaction**: Protein-protein interaction networks

**Edge Connections:**
- Each edge (gene -- term) represents a gene's membership in that biological term
- More connections indicate genes that are part of multiple enriched processes
- The network topology shows which genes are central to the biological response

**Iteration Information:**
- Term nodes are labeled with iteration numbers (term_1_, term_2_, etc.)
- Earlier iterations (lower numbers) represent the most statistically significant enrichments

- Ignore iteration number for biological interpretation, instead focus on network hubs densely connected for interesting biological insights

**ANALYSIS REQUEST:**

Please analyze this network and provide:

1. **Key Biological Insights:**
   - What are the most significant biological processes/pathways identified?
   - Which genes appear to be central to the biological response?

2. **Network Topology Analysis:**
   - Which genes are "hub" genes (connected to multiple terms)?
   - Are there distinct clusters or modules in the network?
   - What does the connectivity pattern suggest about biological organization?

3. **Biological Hypothesis:**
   - Based on the network structure, what biological hypothesis can you generate?
   - How do the different library sources support or complement each other?
   - Try to formulate one hypothesis based on the most central terms and how they possibly related to one another.

4. **Estimated Experimental Context**
   - Can you hypothesize on what was the experiment that generated this result?

**RESPONSE STRUCTURE:**
- **Executive Summary** (2-3 sentences)
- **Key Biological Processes Identified** (by library source)
- **Network Topology Insights**
- **Proposed Biological Hypothesis**
- **Estimated Experimental Context**

**IMPORTANT NOTES:**
- Focus on biological interpretation, not statistical significance or iteration number
- Consider the functional relationships between connected genes and terms
- Look for unexpected connections that might reveal novel or unexpected biology
- Consider the broader biological context and literature
- Note that protein interaction library could shed light on physical interaction between genes while terms indicate functional interaction
- Different library sources may provide complementary biological insights
--

### Executive Summary

This iterative enrichment network is dominated by an **endomembrane/secretory–trafficking axis** (Golgi/ER transport, clathrin endocytosis, Rab/ARF machinery) tightly linked to **membrane protein processing** (N-linked glycosylation + GPI-anchor synthesis) and a second axis of **information flow control** (nuclear pore/export + RNA decay). Layered on top is a **growth-factor / inflammatory signaling core** (EGF–EGFR–PI3K/AKT–NFκB; IL-12/IL-17–JAK/NFκB), suggesting a coordinated program coupling receptor trafficking/processing to signaling outputs.

---

### Key Biological Processes Identified (by library source)

#### H: Hallmark

- **Protein secretion** (HALLMARK: PROTEIN SECRETION)

Genes here (e.g., **ARF1, RAB2A, COG2, GOSR2, TMED2, AP2M1, EGFR**) point to **ER→Golgi→plasma membrane routing** and vesicle formation/fusion.

#### C2: BioCarta

- **EGF/EGFR signaling module** (CARDIACEGF PATHWAY): **EGF, EGFR, RELA, EDNRA**
- **Rab pathway**: **RAB2A, RAB6A, RAB6C**
- **Phosphoinositide signaling** (PTDINS): **AKT1, AP2M1, ARF1**

→ BioCarta is essentially highlighting the **signaling + membrane dynamics** coupling.

#### C2: KEGG Legacy

- **Pancreatic cancer** (a proxy for canonical oncogenic signaling wiring): **EGF/EGFR, AKT1, RELA, IKBKG, JAK1**
- **N-glycan biosynthesis**: **DDOST, DPM1, MGAT1, STT3A**
- **GPI-anchor biosynthesis**: **PIGH, PIGK, PIGY**
- **RNA polymerase / RNA degradation**: **POLR1H/POLR3A/POLR3F** and **EXOSC3/EXOSC5/LSM3/MPHOSPH6**

→ Strong split into **membrane protein maturation** + **RNA production/turnover**.

**C2: KEGG MEDICUS**

- **EGF–EGFR–PI3K–NFκB signaling reference**: **EGF, EGFR, AKT1, RELA, IKBKG**
- **Nuclear export of mRNA**: **NUP107/NUP133/NUP155/NUP160/NUP85, RANBP2**
- **GPI-anchor biosynthesis reference**: **PIGH/PIGK/PIGY**

→ Reinforces that "nuclear export" and "GPI" aren't incidental; they're a consistent second/third pillar.

**C2: Pathway Interaction Database (PID)**

- **ARF pathway / adhesion junction initiation**: **ARF1/AP2M1/CD4/GOSR2** and **AKT1/PIP5K1C/RAP1B/RAPGEF1**

→ Connects vesicle dynamics to **cell surface organization / adhesion machinery**.

**C2: Reactome**

- **Clathrin-mediated endocytosis**: **AP2M1, HIP1R, ARPC1A, CD4, EGFR, EGF, PIP5K1C, GAPVD1**
- **Asparagine N-linked glycosylation**: a broad ER/Golgi processing cohort (**DDOST, DPM1, STT3A, MGAT1, COG2/3/4, TMED2, STX5, TRAPPC1, SCFD1, RAB1B…**)
- **Nuclear RNA decay**: **EXOSC3/5, MPHOSPH6, YTHDC2**
- **IL-12 signaling**: **JAK1** plus immune-associated factors (**PSME2, RAP1B, HNRNPF**)

→ Reactome ties the story together mechanistically: **receptor internalization + glycoprotein biogenesis + RNA surveillance + cytokine signaling**.

**C5: GO Biological Process**

Prominent themes:

- **Golgi vesicle transport**
- **Protein N-linked glycosylation via asparagine**
- **Nuclear export**
- **ARP2/3-mediated actin nucleation**
- **Protein polyubiquitination**
- **Positive regulation of interferon-β production** (notably **POLR3A/POLR3F**, plus **TOMM70**)

→ This adds an innate immune / nucleic-acid sensing flavor on top of trafficking.

**C5: GO Cellular Component**

- **Nuclear pore / outer ring**
- **Core mediator complex**
- **Golgi transport complex**
- **ER protein-containing complex**
- **Clathrin-coated vesicle membrane**
- **Cullin-RING ubiquitin ligase complex**
- **Autophagosome**
- **Nuclear exosome RNase complex**

→ These are "where" the biology happens: **NPC**, **clathrin vesicles**, **ER/Golgi**, **ubiquitin ligase scaffolds**, **autophagy**, **RNA exosome**.

**C5: GO Molecular Function**

- **Catalytic activity acting on RNA** (RNA metabolism cluster)
- **Structural constituent of nuclear pore**
- **Cargo adaptor activity** (AP2M1, HIP1R, GOLPH3…)
- **GTPase activator activity / GTPase binding** (Rab/Arf/Rho regulators)
- **Hexosyltransferase activity** (glycosylation enzymes)
- **Transcription coregulator activity** (Mediator/NCOR2/BCL9/CRTCs…)

→ Functionally, it partitions into **RNA control**, **vesicle adaptor/GTPase regulation**, **glycan enzymology**, **transcriptional control**.

**C5: Human Phenotype Ontology**

- **Abnormal glycosylation** (COG2/3/4, DDOST, DPM1, STT3A)
- **Papilloma** (AKT1, CD4, CXCR4, IKBKG, ERCC3…)
- **Overweight** (GCK, POU1F1, SPTBN1, TAOK1, THOC2…)
- **Aspiration pneumonia** (ATP6V0A1, CSPP1, PURA, SP110, TIMM8A…)
- **Foamy urine** (GAPVD1 + NUPs)

→ These phenotype terms are *interpretive overlays* that often appear when gene lists include fundamental cellular machinery (glycosylation, immune signaling, etc.), but they do underscore the **glycosylation** and **immune/surface receptor** components.

---

**Network Topology Insights**

**Hub genes (multi-term connectors)**

A few genes repeatedly bridge multiple libraries/terms, making them "organizing hubs":

1. **EGF / EGFR / AKT1 / RELA / IKBKG / JAK1**
   - These connect cancer-like signaling (KEGG), canonical pathways (BioCarta/KEGG MEDICUS), and immune modules (Reactome IL-12; WikiPathways IL-17).
   - Interpretation: a **signaling backbone** that is sensitive to trafficking and membrane composition.
2. **AP2M1 / ARF1 / RAB2A (± RAB6A/B/C) / HIP1R / PIP5K1C / GAPVD1**
   - These sit at the intersection of **clathrin endocytosis**, **Golgi transport**, and **Rab/Arf regulation**.
   - Interpretation: a **membrane traffic backbone** (internalization, sorting, retrograde transport).
3. **COG2/3/4, STT3A, DDOST, DPM1, MGAT1, PIGH/PIGK/PIGY, TMED2, STX5, SCFD1, TRAPPC1/8**
   - Dense connections across KEGG/Reactome/GO/HP.
   - Interpretation: a cohesive **ER→Golgi glycoprotein biogenesis + GPI-anchor module**.
4. **NUP107/NUP133/NUP155/NUP160/NUP85 (+ NUP153, RANBP2)**
   - Co-annotated across KEGG MEDICUS + GO CC/MF + GO BP (nuclear export).
   - Interpretation: a distinct **nuclear pore / mRNA export module**.
5. **EXOSC3/EXOSC5/MPHOSPH6/LSM3/YTHDC2**
   - KEGG RNA degradation + Reactome nuclear RNA decay + GO MF "acting on RNA".
   - Interpretation: an **RNA surveillance/turnover module**.

**Distinct clusters/modules**

You can almost read the network as **modules connected by a few bridges**:

- **Module A: Secretory/vesicle trafficking & endocytosis**
- **Module B: Glycosylation + GPI-anchor biogenesis (ER/Golgi processing)**
- **Module C: Nuclear pore/export + RNA decay (information flow control)**
- **Module D: Growth factor + inflammatory signaling (EGFR/PI3K/NFκB; IL-12/IL-17)**

**Bridging logic:** membrane processing/trafficking (A/B) plausibly modulates receptor abundance and signaling intensity (D), while nuclear export/RNA decay (C) tunes the transcriptional/post-transcriptional state that supports the response.

**What the connectivity suggests about organization**

Because the enrichment is iterative *per library*, the persistence of the same themes across independent libraries is a strong topology signal: you're not seeing one "lucky" database hit—you're seeing **convergent annotation**. The graph suggests the gene list is enriched for **core cellular logistics (endomembrane + RNA handling)** with a **surface receptor/inflammatory signaling overlay**.

---

**Proposed Biological Hypothesis**

**Hypothesis:** The underlying perturbation induces (or selects for) a state where **membrane protein maturation and trafficking (ER/Golgi glycosylation + GPI anchoring + vesicle transport + clathrin endocytosis)** are coordinated with **EGFR-centered signaling and NFκB/JAK inflammatory pathways**, potentially to regulate **cell-surface receptor availability and downstream immune-like transcriptional programs**. In parallel, **nuclear export and nuclear RNA decay** appear as a coupled regulatory layer—consistent with cells reprogramming gene expression while also relying on RNA quality control during stress or immune activation.

How the libraries complement each other:

- **Hallmark** gives the high-level state ("protein secretion").
- **Reactome/GO** provide mechanistic granularity (clathrin endocytosis, ER→Golgi glycosylation, autophagy, RNA decay, nuclear pore).
- **KEGG/BioCarta/KEGG MEDICUS** anchor canonical signaling wiring (EGF–EGFR–PI3K–NFκB) and classic biosynthetic pathways (N-glycan, GPI anchor).
- **PID** bridges trafficking to adhesion/junction dynamics (often downstream of receptor signaling and membrane remodeling).
- **HPO** echoes the biological importance of glycosylation and immune/surface receptor pathways (even if not literal phenotypes of your experiment).

A concrete mechanistic story consistent with the topology:

Altered ER/Golgi processing (N-glycosylation/GPI) changes the folding/trafficking of membrane proteins, which shifts receptor internalization and signaling (EGFR), engaging NFκB/JAK pathways; simultaneously, the cell leans on nuclear export and RNA decay to rapidly reshape the transcriptome while maintaining RNA quality under stress/activation.

---

**Estimated Experimental Context**

A plausible origin for a 272-gene hit list with this architecture is a **functional genomic screen** (CRISPR knockout/activation or RNAi) or **proteomics/genetics-derived candidate set** where the measured phenotype is tightly tied to the cell surface and trafficking, for example:

- **A receptor-dependent signaling readout** (e.g., EGFR pathway activation, NFκB reporter activity) where hits include trafficking/glycosylation genes that control receptor abundance/turnover.
- **A host–pathogen entry/replication phenotype** (viral/bacterial toxin entry often depends on clathrin trafficking, Golgi/ER processing, and innate immune signaling—supported here by "positive regulation of interferon-β production" and multiple immune pathway terms).
- **A secretion-based assay** (secreted reporter/cytokine output), which would naturally enrich for ER/Golgi transport, glycosylation, and vesicle coat/adaptor machinery.

If I had to pick *one* best-fit: **a perturbation/screen tied to cell-surface receptor trafficking and inflammatory signaling** (EGFR/NFκB/JAK) with a strong contribution from **secretory pathway capacity and glycoprotein maturation**.

---

If you want, I can also extract a **minimal set of "module marker genes"** (e.g., top 5 per module) from this network to guide follow-up validation (qPCR/KO/rescue/trafficking assays).

---

**Prompt:** The experiment was an siRNA screen where cells were infected with HIV and genes were knocked down using siRNA. The hits from the screen are genes that when knocked down block HIV infection. EGFR has not been directly implicated in HIV infection. Explain the EGFR finding in the network.

**Response:**

EGFR showing up here doesn't necessarily mean "HIV uses EGFR as a receptor." In an siRNA HIV-infection screen, EGFR can appear as a **central network node for three pretty mundane reasons**, all consistent with *your* network (endocytosis/trafficking + PI3K/AKT/NFκB + secretion/glycosylation).

**1) Annotation / enrichment artifact: EGFR is a "pathway magnet"**

EGFR sits in tons of canonical pathway gene sets (EGF–EGFR–PI3K–AKT–NFκB, generic "cancer" maps, clathrin/endocytosis contexts, etc.). In your DOT, EGFR is *pulled into* multiple terms that are really about:

- **clathrin-mediated endocytosis** (AP2M1, HIP1R, PIP5K1C, ARPC1A, GAPVD1…)
- **PI3K/AKT and NFκB signaling** (AKT1, RELA, IKBKG, JAK1…)

So EGFR can become "central" because it's a highly annotated node that co-occurs with the *true* functional module genes, not because it's the causal bottleneck.

**Tell-tale sign:** EGFR's neighboring genes in the network are classic trafficking/signaling hubs (AP2M1/ARF1/RABs/AKT1/RELA/IKBKG), not an EGFR-specific signature.

**2) EGFR knockdown can indirectly break HIV by perturbing the endocytosis/trafficking machinery**

Even if HIV entry is via CD4/CXCR4 (or CCR5), the cell's ability to support infection depends heavily on:

- **membrane remodeling + actin dynamics**
- **clathrin/AP2-dependent internalization and sorting**
- **PI(4,5)P2 / phosphoinositide regulation** (your PTDINS/PIP5K1C links)

EGFR is one of the most intensively trafficked receptors in the cell and is tightly coupled to clathrin/AP2 machinery. Knocking down EGFR can shift the balance of adaptor availability, coated pit dynamics, and downstream lipid/actin regulation—*without* EGFR being "the" entry receptor.

Practical interpretation: EGFR may be acting as a **load-bearing client** of the endocytic system; removing it changes endocytosis "operating conditions," and HIV loses efficiency at entry or early post-entry steps.

**3) EGFR may be a proxy for a required signaling state (PI3K/AKT ↔ NFκB) rather than an HIV-specific factor**

Your network repeatedly hits **AKT1–RELA–IKBKG–JAK1** and cytokine signaling terms. HIV's productive infection strongly benefits from a permissive transcriptional/signaling environment (NFκB-related programs are the obvious one, but also broad survival/metabolic states).

So EGFR could be showing up because:

- EGFR-driven signaling often feeds into **PI3K/AKT** and can crosstalk with **NFκB**.
- Your enrichment terms cluster those components together, so EGFR becomes a "representative" of that module.

In this view, EGFR knockdown blocks infection because it **drops the cell out of a permissive signaling regime**, not because HIV "binds EGFR."

---

**How to decide which explanation is correct (fast, screen-friendly validation)**

**A. Rule out off-target / reagent artifacts**

- Require **≥2 independent siRNAs** against EGFR to reproduce the phenotype.
- Rescue with an **siRNA-resistant EGFR cDNA** (even partial rescue is informative).

**B. Map the blocked step (entry vs post-entry)**

Use matched viral tools:

- HIV Env pseudotype vs **VSV-G** pseudotype
  - If EGFR KD blocks **both**, it's likely a general trafficking/signaling/viability effect.
  - If it blocks **HIV Env only**, it implicates entry/co-receptor trafficking/membrane organization.
- **BlaM-Vpr fusion assay** (entry/fusion)
- Early vs late RT qPCR products (post-entry)

**C. Test the "EGFR is really endocytosis/trafficking" model**

- Flow cytometry for **surface CD4/CXCR4** after EGFR KD (do receptors drop? mislocalize?)
- Generic endocytosis readout (e.g., **transferrin uptake**) to see if clathrin/AP2 function is globally impaired.

**D. Test the "signaling state" model**

- Measure **p-AKT**, **NFκB nuclear translocation**, or an LTR reporter after EGFR KD.
- Compare with pharmacology: do EGFR inhibitors phenocopy the siRNA effect *at non-toxic doses*?

---

**Bottom line**

EGFR's presence in your network is most parsimoniously explained as a **hub annotation + module proxy** for the real biology your screen is screaming: **clathrin/vesicle trafficking + phosphoinositide/actin control + NFκB/AKT-linked permissiveness**. EGFR can land in that center because it's deeply entangled with those systems—even if it's not a canonical HIV dependency factor.

If you tell me what cell line and which infection readout you used (reporter virus? p24? luciferase?), I can predict which of the tests above will be most discriminating.

---

**Prompt**: Analyze my network in the context of the HIV lifecycle (viral entry, transition to nucleus, replication and viral assembly and secretion)

Below is your network "re-labeled" along the **HIV lifecycle**, using the modules that repeatedly appear across libraries (Reactome/GO/KEGG/BioCarta/WP/HPO). I'll call out the **most plausible stage(s)** each module can affect **in a target-cell infection siRNA screen** (i.e., you knock down genes in cells and ask whether infection is blocked).

---

**Executive Summary**

Your hits cluster into four lifecycle-relevant modules: **(1) entry/membrane remodeling & endocytosis**, **(2) cytoplasmic trafficking to the nucleus**, **(3) nuclear pore transport + transcription/elongation + RNA handling**, and **(4) secretory pathway/glycosylation** (which can impact either late steps *or* indirectly affect entry by altering surface protein maturation/availability). The strongest "early infection" signatures are **clathrin/AP2–phosphoinositide–actin** and **nuclear pore/import factors (NUPs, TNPO3)**—both classic bottlenecks for productive HIV infection.

---

**1) Viral Entry / Fusion / Early Post-fusion Events**

**A) Receptor/co-receptor axis (direct entry dependencies)**

- **CD4**, **CXCR4** appear in your network (CD4 in multiple terms; CXCR4 in HPO papilloma node).

*Interpretation:* these can be bona fide hits if your virus is CXCR4-tropic and the assay measures early infection.

**B) Membrane remodeling + clathrin/AP2 endocytosis module (very strong in your network)**

**Terms that scream this:**

- Reactome: **CLATHRIN MEDIATED ENDOCYTOSIS**
- GOCC: **CLATHRIN COATED VESICLE MEMBRANE**
- PID: ARF pathway; nascent adherens junction pathway (membrane/cytoskeleton coupling)
- GOMF: **CARGO ADAPTOR ACTIVITY**

**Key genes in that module:**

- **AP2M1** (AP2 adaptor)
- **HIP1R** (links clathrin structures to actin)
- **PIP5K1C** (makes PI(4,5)P2 → essential for AP2/clathrin recruitment)
- **ARPC1A** / GO BP **ARP2/3-mediated actin nucleation**
- **GAPVD1** (endocytic trafficking regulator)
- **CAV2** (membrane microdomains/caveolar context; can affect receptor organization)

**How this maps to HIV:**

- HIV entry/fusion is extremely sensitive to **local actin remodeling and membrane organization** at the contact site.
- Even when HIV fuses at the plasma membrane, productive entry often needs the cell's **phosphoinositide + actin + adaptor** machinery to cluster receptors, remodel cortical actin, and coordinate membrane dynamics.
- If your assay is reporter expression hours later, these genes can block infection **upstream of reverse transcription** (less fusion/entry) or **immediately downstream** (failed stabilization/trafficking of the incoming core).

**Practical read:** this is one of the most "screen-plausible" early-stage modules in your network.

---

**2) Cytoplasmic Trafficking and "Transition to the Nucleus"**

After fusion, the reverse transcription complex / pre-integration complex must traverse the cytoplasm and reach the nuclear envelope.

**A) Small GTPase trafficking backbone (routing, sorting, long-range movement)**

**Terms:**

- BioCarta: **RAB PATHWAY**
- GO BP: **GOLGI VESICLE TRANSPORT**
- Reactome: **RETROGRADE TRANSPORT AT THE TRANS-GOLGI NETWORK**

**Genes:**

- **ARF1**
- **RAB2A, RAB6A, RAB6B, RAB6C, RAB1B**
- **COG2/3/4**, **TRAPPC1/8**, **SCFD1**, **STX5**, **TMED2**, **GOSR2**, **VPS53**

**How this maps to HIV:**

- These are not "HIV-specific" factors, but they control **membrane traffic highways** and **organelle interface architecture** that can influence:
  - trafficking of receptors and restriction factors
  - availability of lipids/cofactors at the plasma membrane
  - endosomal routing if entry is endocytosis-involved in your system
- In some screens, disturbing trafficking networks also perturbs the cell's ability to support **capsid trafficking** indirectly (cytoskeleton coupling, membrane interactions, compartmentalization).

**B) Cytoskeleton/microtubule-adjacent signals (movement scaffolds)**

**GOCC spindle**, **GOMF alpha-tubulin binding**, and genes like **MAP4, SPAST, DNAL1, FNTA, TAOK1** suggest broader cytoskeletal reliance.

- HIV cores often engage **microtubule-based transport** en route to the nucleus; disrupting cytoskeletal regulators can reduce nuclear delivery.

**Practical read:** this module looks like "cellular logistics" needed to **physically deliver** the incoming complex to the nuclear periphery and to maintain proper membrane/cytoskeletal organization.

---

### 3) Nuclear Entry, Integration-Readiness, and Nuclear Gene Expression Control

This is the **clearest, cleanest lifecycle mapping** in your network.

#### A) Nuclear pore complex (NPC) / nuclear import module (classic HIV dependency)

**Terms:**

- KEGG MEDICUS: **NUCLEAR EXPORT OF mRNA** (really NPC-related)
- GOCC: **NUCLEAR PORE**, **NUCLEAR PORE OUTER RING**
- GOMF: **STRUCTURAL CONSTITUENT OF NUCLEAR PORE**
- GO BP: **NUCLEAR EXPORT**

**Genes:**

- **NUP107, NUP133, NUP155, NUP160, NUP85, NUP153**
- **RANBP2** (NPC-associated)
- **TNPO3** (transportin-3; shows up in your gene list)

**How this maps to HIV:**

- HIV (especially lentiviruses) can infect non-dividing cells because it can access the nucleus through the NPC.
- Many NUPs and transport factors are known "host dependency factors" for the **nuclear import / nuclear entry** phase of HIV infection.
- In a target-cell infection screen, **NUPs/TNPO3** are exactly the kind of hits that reduce infection without affecting viability dramatically.

**Practical read:** if you see strong NUP/TNPO3 signal, the screen is likely capturing a bottleneck around **nuclear access / nuclear transport**.

#### B) Transcriptional control & elongation (post-integration expression)

You have a striking term:

- Reactome: **HIV TRANSCRIPTION ELONGATION**

**Genes:**

- **CTDP1, ELOA, ERCC3** (plus mediator-related factors elsewhere)

You also have:

- GOCC: **CORE MEDIATOR COMPLEX**
- Many **MED** subunits: **MED4, MED6, MED7, MED14, MED28**
- Transcription coregulator MF terms: **NCOR2, BCL9, CRTC2/3, KAT2A, PHF12, NR0B2**, etc.

**How this maps to HIV:**

- After integration, productive infection requires host transcription machinery and elongation control.
- If your assay readout is a reporter dependent on LTR-driven transcription, this module can be a *major* determinant of "infection" as measured.

**Practical read:** the presence of the explicit **HIV transcription elongation** term plus mediator/coregulator modules strongly suggests your readout includes a **gene-expression step** (not just entry).

---

### 4) Viral RNA Processing, Stability, Export, and Translation

Even after transcription, HIV relies on host RNA processing and export.

#### A) Nuclear RNA decay / exosome complex (RNA quality control)

**Terms:**

- Reactome: **NUCLEAR RNA DECAY**

- KEGG: **RNA DEGRADATION**
- GOCC: **NUCLEAR EXOSOME RNASE COMPLEX**
- GOMF: **CATALYTIC ACTIVITY ACTING ON RNA**

**Genes:**
- **EXOSC3, EXOSC5**, **MPHOSPH6**, **LSM3**, **YTHDC2**, plus multiple RNA helicase-like factors (DDX10, DHX33, etc.)

**How this maps to HIV:**
- HIV RNA fate is tightly regulated; host RNA surveillance can shape splicing/export/translation.
- But in an siRNA screen where knockdown blocks infection, this module is often interpreted as: **general RNA metabolism factors needed for HIV gene expression** (or for maintaining a permissive cellular RNA environment).
- Some of these can affect HIV by altering **availability of host splicing/export machinery** or by changing stability of viral transcripts.

**B) mRNA export machinery (NPC-adjacent plus export factors)**
- **THOC2** is in your list (TREX/THO complex component; mRNA export-coupled)
- Together with NUPs, suggests an “RNA export competent nucleus” requirement that could influence viral RNA handling and/or the host transcriptional program required for infection.

---

**5) Viral Assembly, Env Maturation, Budding, and Secretion**

This part requires a nuance: in a **target-cell infection** assay, late-stage “assembly/secretion” genes only matter if your readout depends on new virion production (multi-round spread) or if these genes affect surface proteins and membranes that control entry. Still, your network has a robust secretory/glycosylation signature.

**A) Secretory pathway / protein secretion**

**Term:**
- Hallmark: **PROTEIN SECRETION**

**Genes (representative):**
- **TMED2, GOSR2, COG2, ARF1, RAB2A**, etc.

**How this maps to HIV (late stage and/or indirect effects):**
- For producing cells: Env and host factors required for budding/particle composition depend on the secretory system.
- For target cells: secretory pathway genes can influence **maturation and surface display** of CD4/CXCR4 and other membrane proteins.

**B) N-linked glycosylation (very strong)**

**Terms:**
- KEGG: **N-GLYCAN BIOSYNTHESIS**
- Reactome: **ASPARAGINE N-LINKED GLYCOSYLATION**
- GO BP: **PROTEIN N-LINKED GLYCOSYLATION VIA ASPARAGINE**
- HPO: **ABNORMAL GLYCOSYLATION**

**Genes:**
- **STT3A, DDOST, DPM1, MGAT1**, plus trafficking/complex assembly genes (**COG2/3/4**, etc.)

**How this maps to HIV:**
- In producer cells, N-glycosylation is essential for proper **Env folding, trafficking, and infectivity**.
- In target cells, glycosylation affects **host receptor/cofactor maturation**, membrane protein stability, and endocytic routing—so it can still appear as an “entry-support” module even if you’re not measuring virion production.

**C) Ubiquitination / Cullin-RING ligase complexes (budding or restriction-factor balance)**

**Terms:**
- GO BP: **PROTEIN POLYUBIQUITINATION**
- GOCC: **CULLIN RING UBIQUITIN LIGASE COMPLEX**
- GO MF: **ENZYME SUBSTRATE ADAPTOR ACTIVITY**

**Genes:**

- **FBXW11, HUWE1, RNF26, RNF170, KBTBD7, DCAF1**, etc.

**How this maps to HIV:**

- Ubiquitin machinery can influence:
  - trafficking and degradation of receptors/cofactors
  - innate immune components and restriction factors
  - late budding-related pathways (though your network doesn't explicitly show ESCRT genes)

---

**"So where does this network hit the HIV lifecycle hardest?"**

If I had to rank the stages your network most strongly supports (based purely on topology and repeated term convergence):

1. **Entry / early post-entry membrane remodeling**
(AP2/clathrin + PIP5K1C + ARP2/3 + actin coupling)
2. **Nuclear access / nuclear transport**
(dense NUP module + TNPO3 + NPC structural terms)
3. **Post-integration gene expression**
(explicit HIV transcription elongation + mediator/coregulators)
4. **RNA handling (export/decay)**
(nuclear RNA decay/exosome + RNA catalytic activity)
5. **Secretory/glycosylation**
(could be late-stage if your readout includes production/spread; otherwise likely indirect via surface protein maturation)

---

**Quick "assay interpretation" (to connect lifecycle mapping to your screen)**

Your screen can look like this network under two common designs:

**If your readout was**
**single-round reporter expression**
**(e.g., GFP/luciferase after one cycle)**

Then the strongest drivers are:

- entry/cortical actin/endocytosis competence
- nuclear pore/import factors
- transcription/elongation machinery

**If your readout was**
**multi-round replication**
**or**
**p24/viral production**

Then the secretory/glycosylation machinery becomes much more directly interpretable as:

- Env maturation
- assembly/budding support

---

If you tell me one detail—**was the readout single-round (reporter) or multi-round (spread/p24)**—I can tighten this into a "most likely blocked step" shortlist for each major hub (AP2M1, PIP5K1C, NUPs, TNPO3, EXOSC3/5, STT3A/DDOST).

---

**Prompt: is there a key component in HIV infection that is missing from my network?**

Response:
Yes — a few **canonical HIV lifecycle "workhorse" host components** are notably *absent* from the network you pasted. That doesn't mean they're not important; it usually means **your screen/readout/filters/cell line** didn't make them appear as *enriched hubs*.

Here are the biggest "missing modules," mapped to lifecycle steps, and what their absence suggests.

---

**Viral entry (attachment/co-receptor use)**

**Missing:**
**CCR5**

You have **CD4** and **CXCR4**, but not CCR5.

- If your virus was **X4-tropic** (CXCR4-using) or the cell line doesn't express CCR5, CCR5 won't show up.
- If it *was* R5-tropic and CCR5 is missing, that's a clue that your assay wasn't primarily measuring receptor-level entry limitation (or CCR5 KD wasn't effective / was filtered).

**Interpretation:** your network is consistent with **CD4/CXCR4 contexts**, not macrophage-like CCR5 biology.

---

**Cytoplasmic uncoating / reverse transcription support**

**Missing: classic capsid cofactors like**
**PPIA (Cyclophilin A)**
**and**
**CPSF6**

These often show up in HIV dependency factor lists because they interact with capsid and influence uncoating/nuclear entry.

- You *do* have strong **nuclear pore / transport** signal (NUPs, TNPO3), but not these frequently co-mentioned capsid cofactors.

**Interpretation:** your screen may be dominated by **general trafficking + NPC gating**, rather than the specific "capsid cofactor" axis — or those genes didn't pass filters (e.g., essentiality, knockdown efficiency, redundancy).

---

**Nuclear entry + integration targeting**

**Missing:**
**PSIP1 (LEDGF/p75)**

A classic integration tether (especially relevant for integration site targeting and efficient infection).

- You have abundant NPC components, which fit nuclear entry, but not LEDGF.

**Interpretation:** your "infection" readout may not have been sensitive to integration tethering specifically (or LEDGF KD effects were weaker/partial, or compensated).

---

**HIV transcriptional amplification (Tat/P-TEFb axis)**

**Missing:**
**CDK9 / CCNT1 (Cyclin T1)**
**and related transcription elongation control nodes**

Your network includes a Reactome "HIV transcription elongation" label and mediator/coregulators, but **the core P-TEFb engine** isn't obvious among the hubs shown.

**Interpretation:** the hits you're seeing might be skewed toward:

- **general host transcription infrastructure** (Mediator, ERCC3/ELOA/CTDP1),

rather than the **Tat-specific** elongation machinery, OR those factors were too essential/toxic to score cleanly.

---

**Viral RNA nuclear export (Rev pathway)**

**Missing:**
**XPO1/CRM1**
**and other Rev-export machinery components**

Rev-mediated export of unspliced RNA is central for late gene expression and production.

- If your assay is **single-round reporter** driven early, Rev-export isn't strongly selected.
- If it's multi-round/full replication, you'd expect this more.

**Interpretation:** absence of a Rev-export module often points to a readout emphasizing **early infection or early gene expression**, not late structural gene expression.

---

**Assembly/budding (ESCRT pathway)**

**Missing:**
**TSG101, PDCD6IP (ALIX), CHMPs, VPS4**
**(ESCRT machinery)**

This is the **most striking "late-stage module" missing**.

- Your network is full of **ER/Golgi trafficking + glycosylation**, which can matter for Env maturation, but **true budding machinery** isn't present.

**Interpretation:** your screen likely wasn't measuring **virus production/budding** as the dominant bottleneck (i.e., not a producer-cell budding assay), or those genes were filtered due to essentiality/toxicity.

---

**What this overall suggests about your screen/readout**

Given what you *do* have (clathrin/AP2–PIP5K1C–Arp2/3; NUPs/TNPO3; transcription infrastructure; RNA decay; glycosylation/secretory) and what you *don't* have (CCR5, PPIA/CPSF6, LEDGF, P-TEFb, CRM1, ESCRT), the network looks most consistent with a screen that is heavily sensitive to:

1. **Entry/membrane remodeling + trafficking**
2. **Nuclear access through the pore**
3. **Host transcriptional competence**

…and *less* sensitive to:

- **Rev-dependent late RNA export**
- **budding/ESCRT-mediated release**

---

If you tell me two specifics, I can sharpen this into a crisp "why missing" diagnosis:

1. Was the readout **single-round reporter** vs **multi-round spread/p24**?
2. What cell line and virus tropism (X4 vs R5)?

**Supplementary Table 8**

ORA AI response, Gene Ontology network

Full ChatGPT response for the ORA output over the three Gene Ontology collections, supplied as a JSON list of terms and overlapping genes.

**Executive Summary**

Your ORA reads like two tightly connected modules: (i) a dominant **early secretory/Golgi trafficking axis** (intra-Golgi, ER↔Golgi, and retrograde Golgi→ER) and (ii) a **nucleocytoplasmic transport/mRNA-export axis** centered on the NUP107–160 outer-ring complex. A third, supporting layer involves **protein-complex disassembly/autophagy** and **glyco(lipid) biosynthesis**, consistent with organelle remodeling and membrane reconfiguration under secretory stress.

**Key Biological Processes Identified**

**C5: Gene Ontology – Biological Process (GOBP)**

- **#1 GOLGI VESICLE TRANSPORT:** COG2/3/4, RAB1B/2A/6A/6B/6C, STX5, SCFD1, TRAPPC1/8, TMED2, GOSR2, GOLPH3, SPAST, SPTBN1.
- **#2 INTRA-GOLGI VESICLE-MEDIATED TRANSPORT:** COG2/3/4, RAB6A/B/C, GOSR2, GABARAPL2.
- **#4 RETROGRADE TRANSPORT (GOLGI→ER):** RAB6A/B/C, COG3/4, STX5, SCFD1, GOLPH3.
- **#5 GOLGI ORGANIZATION:** COG2/3/4, GOLPH3, RAB1B/6B, STX5, TMED2, TRAPPC8.
- **#6 ER→GOLGI VESICLE-MEDIATED TRANSPORT:** RAB1B/2A, STX5, SCFD1, GOSR2, TMED2, TRAPPC1/8, COG3, SPAST.
- **#7 ESTABLISHMENT OF PROTEIN LOCALIZATION TO ORGANELLE:** includes GOLPH3, NUPs, TNPO3, TOMM70/VPS53 (protein targeting to organelles).
- **#8–12 Nuclear export / RNA & mRNA export / Nuclear transport: NUP107/133/153/155/160/85**, **RANBP2**, **THOC2**, **TNPO3**, **WNK1**, MDN1, FAM76B.
- **#3 & #13 Protein-complex disassembly (and its regulation): ATG12**, **GABARAPL2**, **UBQLN4**, **SPAST**, **SPTAN1/SPTBN1**, **TAOK1**, **WNK1**.
- **#14 Glycolipid biosynthesis: A4GALT**, **ST3GAL5**, **DPM1**, **PIGK/Y**, **PIGH**, **TM9SF2**.

**C5: Cellular Component (GOCC)**

- **#1 NUCLEAR PORE OUTER RING** and **#2 NUCLEAR PORE: NUP107/133/160/85**, ± **NUP153/155**, **RANBP2**, **RGPD5/8**.
- **#3 CORE MEDIATOR COMPLEX: MED14/28/4/6/7** (transcriptional co-activator scaffold).

**C5: Molecular Function (GOMF)**

- **#1 STRUCTURAL CONSTITUENT OF NUCLEAR PORE:** NUP107/133/153/155/160/85 (confirms CC/BP calls).
- **#2 CATALYTIC ACTIVITY ACTING ON RNA:** RNA helicases/processing (**DDX/DHX**, **EXOSC3/5**, **POLR1/3** subunits, **YTHDC2**, **TRMT5**), supporting the mRNA-biogenesis/export theme.

**Gene–Term Relationship Analysis**

**Recurring "hub" genes (likely key drivers):**

- **Golgi/secretory module (highest density; ranks 1–6):**
    - **COG3/4 (±COG2)** — appear across most Golgi terms; essential for intra-Golgi retrograde tethering.
    - **RAB6A/B/C** — hallmark of **retrograde Golgi→ER** and intra-Golgi trafficking.
    - **STX5 & SCFD1** — t-SNARE and SM protein for ER–Golgi/Golgi fusion.
    - **GOSR2, TMED2, TRAPPC1/8, RAB1B/2A** — ER–Golgi cycling/tethering COPII/COPI interfaces.
    - **GOLPH3** — Golgi morphology/stress sensor; recurs across Golgi-centric and localization terms.
- **Nuclear pore/mRNA-export module (ranks 7–12; GOCC/GOMF corroborate):**
    - **NUP107/133/160/85 (outer ring), NUP153/155**, **RANBP2**, **THOC2**, **TNPO3** — define **NPC architecture** and **TREX-mediated mRNA export**.
    - **WNK1** — repeatedly enriched across transport/disassembly/export terms; a pleiotropic kinase with roles in trafficking/cytoskeleton regulation.
- **Proteostasis/remodeling:**
    - **ATG12, GABARAPL2** (autophagy), **UBQLN4** (ubiquilin), **SPAST** (microtubule severing), **SPTAN1/SPTBN1** (spectrins), **TAOK1** — consistent with turnover of stalled complexes and organelle/cytoskeleton remodeling.
- **Membrane composition:**
    - *A4GALT, ST3GAL5, DPM1, PIG genes** — glyco(lipid)/GPI-anchor synthesis → **Golgi-dependent** receptor/lipid remodeling.

**Patterning & hierarchy from ranks:**

- The **secretory/Golgi cluster dominates the top ranks (#1–6)** → primary biological signal.
- The **nuclear export/NPC cluster sits just below (#7–12)** and is strongly reinforced by GOCC/GOMF, suggesting a **coordinated transcription→export→translation→secretion pipeline**.
- **Disassembly/autophagy** and **glycolipid biosynthesis** round out the picture (cleanup/remodeling and membrane supply), consistent with **adaptive responses to secretory stress**.

**Proposed Biological Hypothesis**

**Coherent story:**

The gene set reflects **cells undergoing acute remodeling of the early secretory pathway**, characterized by **heightened intra-Golgi and retrograde Golgi→ER traffic (RAB6–COG–STX5–SCFD1 axis)** to preserve Golgi homeostasis and recycle cargo/enzymes. To sustain this, cells **up-regulate nucleocytoplasmic transport and mRNA export** via the **NUP107–160 outer ring**, **NUP153/155**, **THOC2**, and **TNPO3**, ensuring rapid delivery of mRNAs encoding trafficking machinery. Concurrently, **autophagy/ubiquilin-mediated disassembly** (ATG12, GABARAPL2, UBQLN4) and **microtubule/spectrin remodeling** (SPAST, SPTAN1/SPTBN1) clear stalled complexes and reorganize organelles. **Glyco(lipid) biosynthesis** (A4GALT, ST3GAL5, PIG*/DPM1) adjusts **membrane composition and cargo receptors**, a hallmark of Golgi-dependent adaptation.

**Net inference:** a **Golgi/ER secretory-stress program** that couples **vesicle trafficking**, **mRNA export**, and **proteostasis** to restore secretory capacity.

**Estimated Experimental Context**

Most likely contexts that generate this composite signature (rank-weighted order):

1. **Acute Golgi disruption / secretory pathway stress** (primary hypothesis)
   - **Why:** Very strong enrichment of **intra-Golgi**, **retrograde Golgi→ER**, **ER→Golgi**, **Golgi organization** with the canonical machinery (**COG3/4, RAB6 isoforms, STX5, SCFD1, TRAPPC, TMED2**). Nuclear export/NPC terms fit a **compensatory increase in macromolecular traffic** to rebuild the pathway.
   - **Typical perturbations: Brefeldin A** or **Golgicide A**, temperature block/release of ER exit, or knockdown of key COPI/COPII tethers.
2. **Genetic dependency screen for retrograde-trafficking cargo (toxin/virus)** (credible alternative)
   - **Why:** The combination of **RAB6–COG–STX5–SCFD1–TRAPPC/TMED2** with **glycolipid biosynthesis** (notably **A4GALT**, **ST3GAL5**) is classic for **Shiga/ricin-like toxin** or certain **polyomavirus** host-factor screens that depend on **Gb3/ganglioside receptors** and **Golgi→ER retrograde routing**.
   - **Note:** The strong **nuclear pore/mRNA export** signal would more strongly support **nuclear-replicating pathogens**; it is less expected for pure toxin phenotypes, which tilts primary inference back to secretory stress unless the screen targeted a **nuclear virus that also exploits the early secretory pathway**.
3. **Induction of a high-secretion cell state** (e.g., early plasmablast differentiation)
   - **Why:** Coordinated rise in **ER/Golgi traffic** and **RNA biogenesis/export** can appear during commitment to professional secretion.
   - **Caveat:** The **retrograde Golgi→ER** and **glycolipid enzyme** emphasis is stronger than typically seen in simple differentiation time courses.

If you can share the phenotype (e.g., “survival after BFA,” “viral replication,” or “differential expression after cytokine X”), we can pin this down to one of the scenarios above with high confidence.

---

**In short:** The data best support **a Golgi/early secretory pathway remodeling program tightly coupled to boosted nuclear export of mRNA**, with **autophagy-mediated cleanup** and **membrane/glycolipid reconfiguration** as enabling processes.

## Supplementary Table 10

Hypoxia signature AI response

Prompt, network and full ChatGPT response for the 75-gene hypoxia response signature.

75 Genes that form the response to Hypoxia were analyzed with iGEA
[ChatGPT Analysis Response]

**Prompt to ChatGPT-5**
You are a computational biologist analyzing an iterative gene set enrichment network. Iterative means that for each library source, the top enriched gene set is found, saved and the genes are removed from the initial gene list. The remaining genes are then tested for enrichment again. Thus, each gene appears only once per library source tested but can be linked to multiple terms that originate from different libraries. The results are provided as a DOT network represents the relationships between genes and gene sets(=terms).

The original gene list included 75 genes and was tested across the following libraries: H: Hallmark Gene Sets, C2: KEGG Legacy, C2: KEGG MEDICUS, C2: Pathway Interaction Database, C2: Reactome Pathways, C2: WikiPathways, C5: Gene Ontology: Biological Process, C5: Gene Ontology: Cellular Component, C5: Gene Ontology: Molecular Function, C5: Human Phenotype Ontology.

**NETWORK STRUCTURE:**

```
graph iterative_enrichment_all {
graph [layout=neato];
node [shape=ellipse];
subgraph cluster_legend_libs {
label="Libraries";
node [shape=box, style=filled, fontcolor="white"];
legend_lib_1 [label="H: Hallmark Gene Sets", fillcolor="#FF6B6B"];
legend_lib_2 [label="C2: KEGG Legacy", fillcolor="#F8C471"];
legend_lib_3 [label="C2: KEGG MEDICUS", fillcolor="#98D8C8"];
legend_lib_4 [label="C2: Pathway Interaction Database", fillcolor="#440154"];
legend_lib_5 [label="C2: Reactome Pathways", fillcolor="#BB8FCE"];
legend_lib_6 [label="C2: WikiPathways", fillcolor="#31688e"];
legend_lib_7 [label="C5: Gene Ontology: Biological Process", fillcolor="#F1948A"];
legend_lib_8 [label="C5: Gene Ontology: Cellular Component", fillcolor="#85C1E9"];
legend_lib_9 [label="C5: Gene Ontology: Molecular Function", fillcolor="#D7BDE2"];
legend_lib_10 [label="C5: Human Phenotype Ontology", fillcolor="#F7DC6F"];
}
"gene_ALDOC" [label="ALDOC", type="gene", width=0.7, height=0.7];
"gene_ATF3" [label="ATF3", type="gene", width=0.7, height=0.7];
"gene_CXCR4" [label="CXCR4", type="gene", width=0.7, height=0.7];
"gene_DDIT4" [label="DDIT4", type="gene", width=0.7, height=0.7];
"gene_EFNA1" [label="EFNA1", type="gene", width=0.7, height=0.7];
"gene_ERO1A" [label="ERO1A", type="gene", width=0.7, height=0.7];
"gene_FAM162A" [label="FAM162A", type="gene", width=0.7, height=0.7];
"gene_GAPDH" [label="GAPDH", type="gene", width=0.7, height=0.7];
"gene_GYS1" [label="GYS1", type="gene", width=0.7, height=0.7];
"gene_KDM3A" [label="KDM3A", type="gene", width=0.7, height=0.7];
"gene_LDHA" [label="LDHA", type="gene", width=0.7, height=0.7];
"gene_LOX" [label="LOX", type="gene", width=0.7, height=0.7];
"gene_MIF" [label="MIF", type="gene", width=0.7, height=0.7];
"gene_MXI1" [label="MXI1", type="gene", width=0.7, height=0.7];
"gene_NDRG1" [label="NDRG1", type="gene", width=0.7, height=0.7];
"gene_NR3C1" [label="NR3C1", type="gene", width=0.7, height=0.7];
"gene_P4HA1" [label="P4HA1", type="gene", width=0.7, height=0.7];
"gene_P4HA2" [label="P4HA2", type="gene", width=0.7, height=0.7];
"gene_PGK1" [label="PGK1", type="gene", width=0.7, height=0.7];
"gene_PIM1" [label="PIM1", type="gene", width=0.7, height=0.7];
"gene_RBPJ" [label="RBPJ", type="gene", width=0.7, height=0.7];
"gene_RRAGD" [label="RRAGD", type="gene", width=0.7, height=0.7];
"gene_STC2" [label="STC2", type="gene", width=0.7, height=0.7];
"gene_TMEM45A" [label="TMEM45A", type="gene", width=0.7, height=0.7];
"gene_WSB1" [label="WSB1", type="gene", width=0.7, height=0.7];
"term_1_HALLMARK_HYPOXIA" [label="HALLMARK: HYPOXIA", style=filled, fillcolor="#FF6B6B", fontcolor="white", type="term", width=2, height=2, library="H: Hallmark Gene Sets"];
"gene_ALDOC" -- "term_1_HALLMARK_HYPOXIA" [color="#FF6B6B"];
"gene_ATF3" -- "term_1_HALLMARK_HYPOXIA" [color="#FF6B6B"];
"gene_CXCR4" -- "term_1_HALLMARK_HYPOXIA" [color="#FF6B6B"];
"gene_DDIT4" -- "term_1_HALLMARK_HYPOXIA" [color="#FF6B6B"];
"gene_EFNA1" -- "term_1_HALLMARK_HYPOXIA" [color="#FF6B6B"];
"gene_ERO1A" -- "term_1_HALLMARK_HYPOXIA" [color="#FF6B6B"];
"gene_FAM162A" -- "term_1_HALLMARK_HYPOXIA" [color="#FF6B6B"];
"gene_GAPDH" -- "term_1_HALLMARK_HYPOXIA" [color="#FF6B6B"];
"gene_GYS1" -- "term_1_HALLMARK_HYPOXIA" [color="#FF6B6B"];
"gene_KDM3A" -- "term_1_HALLMARK_HYPOXIA" [color="#FF6B6B"];
"gene_LDHA" -- "term_1_HALLMARK_HYPOXIA" [color="#FF6B6B"];
"gene_LOX" -- "term_1_HALLMARK_HYPOXIA" [color="#FF6B6B"];
"gene_MIF" -- "term_1_HALLMARK_HYPOXIA" [color="#FF6B6B"];
"gene_MXI1" -- "term_1_HALLMARK_HYPOXIA" [color="#FF6B6B"];
"gene_NDRG1" -- "term_1_HALLMARK_HYPOXIA" [color="#FF6B6B"];
"gene_NR3C1" -- "term_1_HALLMARK_HYPOXIA" [color="#FF6B6B"];
"gene_P4HA1" -- "term_1_HALLMARK_HYPOXIA" [color="#FF6B6B"];
"gene_P4HA2" -- "term_1_HALLMARK_HYPOXIA" [color="#FF6B6B"];
"gene_PGK1" -- "term_1_HALLMARK_HYPOXIA" [color="#FF6B6B"];
"gene_PIM1" -- "term_1_HALLMARK_HYPOXIA" [color="#FF6B6B"];
"gene_RBPJ" -- "term_1_HALLMARK_HYPOXIA" [color="#FF6B6B"];
"gene_RRAGD" -- "term_1_HALLMARK_HYPOXIA" [color="#FF6B6B"];
"gene_STC2" -- "term_1_HALLMARK_HYPOXIA" [color="#FF6B6B"];
"gene_TMEM45A" -- "term_1_HALLMARK_HYPOXIA" [color="#FF6B6B"];
"gene_WSB1" -- "term_1_HALLMARK_HYPOXIA" [color="#FF6B6B"];
"gene_CCNB1" [label="CCNB1", type="gene", width=0.7, height=0.7];
"gene_CYCS" [label="CYCS", type="gene", width=0.7, height=0.7];
"gene_GADD45B" [label="GADD45B", type="gene", width=0.7, height=0.7];
"gene_PGAM1" [label="PGAM1", type="gene", width=0.7, height=0.7];
"term_1_KEGG_GLYCOLYSIS_GLUCONEOGENESIS" [label="KEGG: GLYCOLYSIS GLUCONEOGENESIS", style=filled, fillcolor="#F8C471", fontcolor="white", type="term", width=2, height=2, library="C2: KEGG Legacy"];
"term_2_KEGG_P53_SIGNALING_PATHWAY" [label="KEGG: P53 SIGNALING PATHWAY", style=filled, fillcolor="#F8C471", fontcolor="white", type="term", width=2, height=2, library="C2: KEGG Legacy"];
"gene_ALDOC" -- "term_1_KEGG_GLYCOLYSIS_GLUCONEOGENESIS" [color="#F8C471"];
"gene_CCNB1" -- "term_2_KEGG_P53_SIGNALING_PATHWAY" [color="#F8C471"];
```

"gene_CYCS" -- "term_2_KEGG_P53_SIGNALING_PATHWAY" [color="#F8C471"];
"gene_GADD45B" -- "term_2_KEGG_P53_SIGNALING_PATHWAY" [color="#F8C471"];
"gene_GAPDH" -- "term_1_KEGG_GLYCOLYSIS_GLUCONEOGENESIS" [color="#F8C471"];
"gene_LDHA" -- "term_1_KEGG_GLYCOLYSIS_GLUCONEOGENESIS" [color="#F8C471"];
"gene_PGAM1" -- "term_1_KEGG_GLYCOLYSIS_GLUCONEOGENESIS" [color="#F8C471"];
"gene_PGK1" -- "term_1_KEGG_GLYCOLYSIS_GLUCONEOGENESIS" [color="#F8C471"];
"term_1_KEGG_MEDICUS_REFERENCE_GLYCOLYSIS" [label="KEGG: MEDICUS REFERENCE GLYCOLYSIS", style=filled, fillcolor="#98D8C8", fontcolor="white", type="term", width=2, height=2, library="C2: KEGG MEDICUS"];
"gene_ALDOC" -- "term_1_KEGG_MEDICUS_REFERENCE_GLYCOLYSIS" [color="#98D8C8"];
"gene_GAPDH" -- "term_1_KEGG_MEDICUS_REFERENCE_GLYCOLYSIS" [color="#98D8C8"];
"gene_LDHA" -- "term_1_KEGG_MEDICUS_REFERENCE_GLYCOLYSIS" [color="#98D8C8"];
"gene_PGAM1" -- "term_1_KEGG_MEDICUS_REFERENCE_GLYCOLYSIS" [color="#98D8C8"];
"gene_PGK1" -- "term_1_KEGG_MEDICUS_REFERENCE_GLYCOLYSIS" [color="#98D8C8"];
"gene_BNIP3" [label="BNIP3", type="gene", width=0.7, height=0.7];
"gene_CAD" [label="CAD", type="gene", width=0.7, height=0.7];
"term_1_PID_HIF1_TFPATHWAY" [label="PID: HIF1 TFPATHWAY", style=filled, fillcolor="#440154", fontcolor="white", type="term", width=2, height=2, library="Unknown Library"];
"term_2_PID_MYC_ACTIV_PATHWAY" [label="PID: MYC ACTIV PATHWAY", style=filled, fillcolor="#440154", fontcolor="white", type="term", width=2, height=2, library="Unknown Library"];
"gene_BNIP3" -- "term_1_PID_HIF1_TFPATHWAY" [color="#440154"];
"gene_CAD" -- "term_2_PID_MYC_ACTIV_PATHWAY" [color="#440154"];
"gene_CCNB1" -- "term_2_PID_MYC_ACTIV_PATHWAY" [color="#440154"];
"gene_CXCR4" -- "term_1_PID_HIF1_TFPATHWAY" [color="#440154"];
"gene_GAPDH" -- "term_2_PID_MYC_ACTIV_PATHWAY" [color="#440154"];
"gene_LDHA" -- "term_1_PID_HIF1_TFPATHWAY" [color="#440154"];
"gene_NDRG1" -- "term_1_PID_HIF1_TFPATHWAY" [color="#440154"];
"gene_PGK1" -- "term_1_PID_HIF1_TFPATHWAY" [color="#440154"];
"gene_PIM1" -- "term_2_PID_MYC_ACTIV_PATHWAY" [color="#440154"];
"gene_ACTR1A" [label="ACTR1A", type="gene", width=0.7, height=0.7];
"gene_KDM4B" [label="KDM4B", type="gene", width=0.7, height=0.7];
"gene_KDM4C" [label="KDM4C", type="gene", width=0.7, height=0.7];
"gene_PLOD1" [label="PLOD1", type="gene", width=0.7, height=0.7];
"gene_TUBG1" [label="TUBG1", type="gene", width=0.7, height=0.7];
"term_1_REACTOME_GLUCONEOGENESIS" [label="REACTOME: GLUCONEOGENESIS", style=filled, fillcolor="#BB8FCE", fontcolor="white", type="term", width=2, height=2, library="C2: Reactome Pathways"];
"term_2_REACTOME_COLLAGEN_FORMATION" [label="REACTOME: COLLAGEN FORMATION", style=filled, fillcolor="#BB8FCE", fontcolor="white", type="term", width=2, height=2, library="C2: Reactome Pathways"];
"term_3_REACTOME_HDMS_DEMETHYLATE_HISTONES" [label="REACTOME: HDMS DEMETHYLATE HISTONES", style=filled, fillcolor="#BB8FCE", fontcolor="white", type="term", width=2, height=2, library="C2: Reactome Pathways"];
"term_4_REACTOME_TP53_REGULATES_METABOLIC_GENES" [label="REACTOME: TP53 REGULATES METABOLIC GENES", style=filled, fillcolor="#BB8FCE", fontcolor="white", type="term", width=2, height=2, library="C2: Reactome Pathways"];
"term_5_REACTOME_REGULATION_OF_PLK1_ACTIVITY_AT_G2_M_TRANSITION" [label="REACTOME: REGULATION OF PLK1 ACTIVITY AT G2 M TRANSITION", style=filled, fillcolor="#BB8FCE", fontcolor="white", type="term", width=2, height=2, library="C2: Reactome Pathways"];
"gene_ACTR1A" -- "term_5_REACTOME_REGULATION_OF_PLK1_ACTIVITY_AT_G2_M_TRANSITION" [color="#BB8FCE"];
"gene_ALDOC" -- "term_1_REACTOME_GLUCONEOGENESIS" [color="#BB8FCE"];
"gene_CCNB1" -- "term_5_REACTOME_REGULATION_OF_PLK1_ACTIVITY_AT_G2_M_TRANSITION" [color="#BB8FCE"];
"gene_CYCS" -- "term_4_REACTOME_TP53_REGULATES_METABOLIC_GENES" [color="#BB8FCE"];
"gene_DDIT4" -- "term_4_REACTOME_TP53_REGULATES_METABOLIC_GENES" [color="#BB8FCE"];
"gene_GAPDH" -- "term_1_REACTOME_GLUCONEOGENESIS" [color="#BB8FCE"];
"gene_KDM3A" -- "term_3_REACTOME_HDMS_DEMETHYLATE_HISTONES" [color="#BB8FCE"];
"gene_KDM4B" -- "term_3_REACTOME_HDMS_DEMETHYLATE_HISTONES" [color="#BB8FCE"];
"gene_KDM4C" -- "term_3_REACTOME_HDMS_DEMETHYLATE_HISTONES" [color="#BB8FCE"];
"gene_LOX" -- "term_2_REACTOME_COLLAGEN_FORMATION" [color="#BB8FCE"];
"gene_P4HA1" -- "term_2_REACTOME_COLLAGEN_FORMATION" [color="#BB8FCE"];
"gene_P4HA2" -- "term_2_REACTOME_COLLAGEN_FORMATION" [color="#BB8FCE"];
"gene_PGAM1" -- "term_1_REACTOME_GLUCONEOGENESIS" [color="#BB8FCE"];
"gene_PGK1" -- "term_1_REACTOME_GLUCONEOGENESIS" [color="#BB8FCE"];
"gene_PLOD1" -- "term_2_REACTOME_COLLAGEN_FORMATION" [color="#BB8FCE"];
"gene_RRAGD" -- "term_4_REACTOME_TP53_REGULATES_METABOLIC_GENES" [color="#BB8FCE"];
"gene_TUBG1" -- "term_5_REACTOME_REGULATION_OF_PLK1_ACTIVITY_AT_G2_M_TRANSITION" [color="#BB8FCE"];
"gene_NAMPT" [label="NAMPT", type="gene", width=0.7, height=0.7];
"gene_RARA" [label="RARA", type="gene", width=0.7, height=0.7];
"gene_SLC16A1" [label="SLC16A1", type="gene", width=0.7, height=0.7];
"term_1_WP_GLYCOLYSIS_IN_SENESCENCE" [label="WP: GLYCOLYSIS IN SENESCENCE", style=filled, fillcolor="#31688e", fontcolor="white", type="term", width=2, height=2, library="Unknown Library"];
"term_2_WP_METABOLIC_PATHWAYS_OF_FIBROBLASTS" [label="WP: METABOLIC PATHWAYS OF FIBROBLASTS", style=filled, fillcolor="#31688e", fontcolor="white", type="term", width=2, height=2, library="Unknown Library"];
"term_3_WP_ADIPOGENESIS" [label="WP: ADIPOGENESIS", style=filled, fillcolor="#31688e", fontcolor="white", type="term", width=2, height=2, library="Unknown Library"];
"term_4_WP_HEMATOPOIETIC_STEM_CELL_DIFFERENTIATION" [label="WP: HEMATOPOIETIC STEM CELL DIFFERENTIATION", style=filled, fillcolor="#31688e", fontcolor="white", type="term", width=2, height=2, library="Unknown Library"];
"gene_ALDOC" -- "term_1_WP_GLYCOLYSIS_IN_SENESCENCE" [color="#31688e"];
"gene_CXCR4" -- "term_4_WP_HEMATOPOIETIC_STEM_CELL_DIFFERENTIATION" [color="#31688e"];
"gene_GADD45B" -- "term_3_WP_ADIPOGENESIS" [color="#31688e"];
"gene_GAPDH" -- "term_1_WP_GLYCOLYSIS_IN_SENESCENCE" [color="#31688e"];
"gene_LDHA" -- "term_1_WP_GLYCOLYSIS_IN_SENESCENCE" [color="#31688e"];
"gene_MIF" -- "term_3_WP_ADIPOGENESIS" [color="#31688e"];
"gene_MXI1" -- "term_4_WP_HEMATOPOIETIC_STEM_CELL_DIFFERENTIATION" [color="#31688e"];
"gene_NAMPT" -- "term_3_WP_ADIPOGENESIS" [color="#31688e"];
"gene_NR3C1" -- "term_3_WP_ADIPOGENESIS" [color="#31688e"];
"gene_P4HA1" -- "term_2_WP_METABOLIC_PATHWAYS_OF_FIBROBLASTS" [color="#31688e"];
"gene_P4HA2" -- "term_2_WP_METABOLIC_PATHWAYS_OF_FIBROBLASTS" [color="#31688e"];
"gene_PGK1" -- "term_1_WP_GLYCOLYSIS_IN_SENESCENCE" [color="#31688e"];
"gene_PIM1" -- "term_4_WP_HEMATOPOIETIC_STEM_CELL_DIFFERENTIATION" [color="#31688e"];
"gene_PLOD1" -- "term_2_WP_METABOLIC_PATHWAYS_OF_FIBROBLASTS" [color="#31688e"];
"gene_RARA" -- "term_3_WP_ADIPOGENESIS" [color="#31688e"];
"gene_SLC16A1" -- "term_2_WP_METABOLIC_PATHWAYS_OF_FIBROBLASTS" [color="#31688e"];
"gene_CRKL" [label="CRKL", type="gene", width=0.7, height=0.7];
"gene_FH" [label="FH", type="gene", width=0.7, height=0.7];
"gene_HILPDA" [label="HILPDA", type="gene", width=0.7, height=0.7];
"gene_ILF3" [label="ILF3", type="gene", width=0.7, height=0.7];
"gene_KLF10" [label="KLF10", type="gene", width=0.7, height=0.7];
"gene_PJA2" [label="PJA2", type="gene", width=0.7, height=0.7];
"gene_TGFBR1" [label="TGFBR1", type="gene", width=0.7, height=0.7];
"gene_TPCN1" [label="TPCN1", type="gene", width=0.7, height=0.7];
"gene_VDAC1" [label="VDAC1", type="gene", width=0.7, height=0.7];
"term_1_GOBP_PYRUVATE_METABOLIC_PROCESS" [label="GOBP: PYRUVATE METABOLIC PROCESS", style=filled, fillcolor="#F1948A", fontcolor="white", type="term", width=2, height=2, library="C5: Gene Ontology: Biological Process"];
"term_2_GOBP_CELLULAR_RESPONSE_TO_OXYGEN_LEVELS" [label="GOBP: CELLULAR RESPONSE TO OXYGEN LEVELS", style=filled, fillcolor="#F1948A", fontcolor="white", type="term", width=2, height=2, library="C5: Gene Ontology: Biological Process"];
"term_3_GOBP_NUCLEAR_RECEPTOR_MEDIATED_SIGNALING_PATHWAY" [label="GOBP: NUCLEAR RECEPTOR MEDIATED SIGNALING PATHWAY", style=filled, fillcolor="#F1948A", fontcolor="white", type="term", width=2, height=2, library="C5: Gene Ontology: Biological Process"];
"term_4_GOBP_COLLAGEN_FIBRIL_ORGANIZATION" [label="GOBP: COLLAGEN FIBRIL ORGANIZATION", style=filled, fillcolor="#F1948A", fontcolor="white", type="term", width=2, height=2, library="C5: Gene Ontology: Biological Process"];
"term_5_GOBP_RESPONSE_TO_STARVATION" [label="GOBP: RESPONSE TO STARVATION", style=filled, fillcolor="#F1948A", fontcolor="white", type="term", width=2, height=2, library="C5: Gene Ontology: Biological Process"];
"term_6_GOBP_SYMBIONT_ENTRY_INTO_HOST" [label="GOBP: SYMBIONT ENTRY INTO HOST", style=filled, fillcolor="#F1948A", fontcolor="white", type="term", width=2, height=2, library="C5: Gene Ontology: Biological Process"];
"term_7_GOBP_POSITIVE_REGULATION_OF_MAPK_CASCADE" [label="GOBP: POSITIVE REGULATION OF MAPK CASCADE", style=filled, fillcolor="#F1948A", fontcolor="white", type="term", width=2, height=2, library="C5: Gene Ontology: Biological Process"];
"term_8_GOBP_GENERATION_OF_PRECURSOR_METABOLITES_AND_ENERGY" [label="GOBP: GENERATION OF PRECURSOR METABOLITES AND ENERGY", style=filled, fillcolor="#F1948A", fontcolor="white", type="term", width=2, height=2, library="C5: Gene Ontology: Biological Process"];
"gene_ALDOC" -- "term_1_GOBP_PYRUVATE_METABOLIC_PROCESS" [color="#F1948A"];
"gene_ATF3" -- "term_5_GOBP_RESPONSE_TO_STARVATION" [color="#F1948A"];
"gene_BNIP3" -- "term_2_GOBP_CELLULAR_RESPONSE_TO_OXYGEN_LEVELS" [color="#F1948A"];
"gene_CAD" -- "term_5_GOBP_RESPONSE_TO_STARVATION" [color="#F1948A"];
"gene_CCNB1" -- "term_2_GOBP_CELLULAR_RESPONSE_TO_OXYGEN_LEVELS" [color="#F1948A"];
"gene_CRKL" -- "term_3_GOBP_NUCLEAR_RECEPTOR_MEDIATED_SIGNALING_PATHWAY" [color="#F1948A"];
"gene_CXCR4" -- "term_6_GOBP_SYMBIONT_ENTRY_INTO_HOST" [color="#F1948A"];
"gene_CYCS" -- "term_8_GOBP_GENERATION_OF_PRECURSOR_METABOLITES_AND_ENERGY" [color="#F1948A"];
"gene_DDIT4" -- "term_1_GOBP_PYRUVATE_METABOLIC_PROCESS" [color="#F1948A"];
"gene_EFNA1" -- "term_7_GOBP_POSITIVE_REGULATION_OF_MAPK_CASCADE" [color="#F1948A"];
"gene_ERO1A" -- "term_2_GOBP_CELLULAR_RESPONSE_TO_OXYGEN_LEVELS" [color="#F1948A"];
"gene_FAM162A" -- "term_2_GOBP_CELLULAR_RESPONSE_TO_OXYGEN_LEVELS" [color="#F1948A"];
"gene_FH" -- "term_8_GOBP_GENERATION_OF_PRECURSOR_METABOLITES_AND_ENERGY" [color="#F1948A"];
"gene_GADD45B" -- "term_7_GOBP_POSITIVE_REGULATION_OF_MAPK_CASCADE" [color="#F1948A"];
"gene_GAPDH" -- "term_1_GOBP_PYRUVATE_METABOLIC_PROCESS" [color="#F1948A"];
"gene_GYS1" -- "term_8_GOBP_GENERATION_OF_PRECURSOR_METABOLITES_AND_ENERGY" [color="#F1948A"];
"gene_HILPDA" -- "term_2_GOBP_CELLULAR_RESPONSE_TO_OXYGEN_LEVELS" [color="#F1948A"];
"gene_ILF3" -- "term_6_GOBP_SYMBIONT_ENTRY_INTO_HOST" [color="#F1948A"];
"gene_KDM3A" -- "term_3_GOBP_NUCLEAR_RECEPTOR_MEDIATED_SIGNALING_PATHWAY" [color="#F1948A"];
"gene_KDM4C" -- "term_3_GOBP_NUCLEAR_RECEPTOR_MEDIATED_SIGNALING_PATHWAY" [color="#F1948A"];
"gene_KLF10" -- "term_5_GOBP_RESPONSE_TO_STARVATION" [color="#F1948A"];
"gene_LDHA" -- "term_1_GOBP_PYRUVATE_METABOLIC_PROCESS" [color="#F1948A"];
"gene_LOX" -- "term_4_GOBP_COLLAGEN_FIBRIL_ORGANIZATION" [color="#F1948A"];
"gene_MIF" -- "term_7_GOBP_POSITIVE_REGULATION_OF_MAPK_CASCADE" [color="#F1948A"];
"gene_NAMPT" -- "term_7_GOBP_POSITIVE_REGULATION_OF_MAPK_CASCADE" [color="#F1948A"];
"gene_NDRG1" -- "term_2_GOBP_CELLULAR_RESPONSE_TO_OXYGEN_LEVELS" [color="#F1948A"];
"gene_NR3C1" -- "term_3_GOBP_NUCLEAR_RECEPTOR_MEDIATED_SIGNALING_PATHWAY" [color="#F1948A"];
"gene_P4HA1" -- "term_4_GOBP_COLLAGEN_FIBRIL_ORGANIZATION" [color="#F1948A"];
"gene_PGAM1" -- "term_1_GOBP_PYRUVATE_METABOLIC_PROCESS" [color="#F1948A"];
"gene_PGK1" -- "term_1_GOBP_PYRUVATE_METABOLIC_PROCESS" [color="#F1948A"];
"gene_PIM1" -- "term_3_GOBP_NUCLEAR_RECEPTOR_MEDIATED_SIGNALING_PATHWAY" [color="#F1948A"];
"gene_PJA2" -- "term_7_GOBP_POSITIVE_REGULATION_OF_MAPK_CASCADE" [color="#F1948A"];
"gene_PLOD1" -- "term_4_GOBP_COLLAGEN_FIBRIL_ORGANIZATION" [color="#F1948A"];

"gene_RARA" -- "term_3_GOBP_NUCLEAR_RECEPTOR_MEDIATED_SIGNALING_PATHWAY" [color="#F1948A"];
"gene_RBPJ" -- "term_8_GOBP_GENERATION_OF_PRECURSOR_METABOLITES_AND_ENERGY" [color="#F1948A"];
"gene_RRAGD" -- "term_5_GOBP_RESPONSE_TO_STARVATION" [color="#F1948A"];
"gene_SLC16A1" -- "term_1_GOBP_PYRUVATE_METABOLIC_PROCESS" [color="#F1948A"];
"gene_STC2" -- "term_2_GOBP_CELLULAR_RESPONSE_TO_OXYGEN_LEVELS" [color="#F1948A"];
"gene_TGFBR1" -- "term_4_GOBP_COLLAGEN_FIBRIL_ORGANIZATION" [color="#F1948A"];
"gene_TPCN1" -- "term_6_GOBP_SYMBIONT_ENTRY_INTO_HOST" [color="#F1948A"];
"gene_VDAC1" -- "term_1_GOBP_PYRUVATE_METABOLIC_PROCESS" [color="#F1948A"];
"gene_ASCC1" [label="ASCC1", type="gene", width=0.7, height=0.7];
"gene_ASPH" [label="ASPH", type="gene", width=0.7, height=0.7];
"gene_ATF7IP" [label="ATF7IP", type="gene", width=0.7, height=0.7];
"gene_EDEM3" [label="EDEM3", type="gene", width=0.7, height=0.7];
"gene_MRPL4" [label="MRPL4", type="gene", width=0.7, height=0.7];
"gene_MRPS12" [label="MRPS12", type="gene", width=0.7, height=0.7];
"gene_NARF" [label="NARF", type="gene", width=0.7, height=0.7];
"gene_NXF1" [label="NXF1", type="gene", width=0.7, height=0.7];
"gene_RAB8B" [label="RAB8B", type="gene", width=0.7, height=0.7];
"gene_SPAG4" [label="SPAG4", type="gene", width=0.7, height=0.7];
"term_1_GOCC_ENDOPLASMIC_RETICULUM_LUMEN" [label="GOCC: ENDOPLASMIC RETICULUM LUMEN", style=filled, fillcolor="#85C1E9", fontcolor="white", type="term", width=2, height=2, library="C5: Gene Ontology: Cellular Component"];
"term_2_GOCC_MITOCHONDRIAL_MATRIX" [label="GOCC: MITOCHONDRIAL MATRIX", style=filled, fillcolor="#85C1E9", fontcolor="white", type="term", width=2, height=2, library="C5: Gene Ontology: Cellular Component"];
"term_3_GOCC_RECYCLING_ENDOSOME" [label="GOCC: RECYCLING ENDOSOME", style=filled, fillcolor="#85C1E9", fontcolor="white", type="term", width=2, height=2, library="C5: Gene Ontology: Cellular Component"];
"term_4_GOCC_TRANSCRIPTION_REGULATOR_COMPLEX" [label="GOCC: TRANSCRIPTION REGULATOR COMPLEX", style=filled, fillcolor="#85C1E9", fontcolor="white", type="term", width=2, height=2, library="C5: Gene Ontology: Cellular Component"];
"term_5_GOCC_FICOLIN_1_RICH_GRANULE_LUMEN" [label="GOCC: FICOLIN 1 RICH GRANULE LUMEN", style=filled, fillcolor="#85C1E9", fontcolor="white", type="term", width=2, height=2, library="C5: Gene Ontology: Cellular Component"];
"term_6_GOCC_NUCLEAR_ENVELOPE" [label="GOCC: NUCLEAR ENVELOPE", style=filled, fillcolor="#85C1E9", fontcolor="white", type="term", width=2, height=2, library="C5: Gene Ontology: Cellular Component"];
"gene_ALDOC" -- "term_5_GOCC_FICOLIN_1_RICH_GRANULE_LUMEN" [color="#85C1E9"];
"gene_ASCC1" -- "term_4_GOCC_TRANSCRIPTION_REGULATOR_COMPLEX" [color="#85C1E9"];
"gene_ASPH" -- "term_1_GOCC_ENDOPLASMIC_RETICULUM_LUMEN" [color="#85C1E9"];
"gene_ATF3" -- "term_4_GOCC_TRANSCRIPTION_REGULATOR_COMPLEX" [color="#85C1E9"];
"gene_ATF7IP" -- "term_4_GOCC_TRANSCRIPTION_REGULATOR_COMPLEX" [color="#85C1E9"];
"gene_BNIP3" -- "term_6_GOCC_NUCLEAR_ENVELOPE" [color="#85C1E9"];
"gene_CCNB1" -- "term_2_GOCC_MITOCHONDRIAL_MATRIX" [color="#85C1E9"];
"gene_EDEM3" -- "term_1_GOCC_ENDOPLASMIC_RETICULUM_LUMEN" [color="#85C1E9"];
"gene_ERO1A" -- "term_1_GOCC_ENDOPLASMIC_RETICULUM_LUMEN" [color="#85C1E9"];
"gene_FH" -- "term_2_GOCC_MITOCHONDRIAL_MATRIX" [color="#85C1E9"];
"gene_GAPDH" -- "term_6_GOCC_NUCLEAR_ENVELOPE" [color="#85C1E9"];
"gene_MIF" -- "term_5_GOCC_FICOLIN_1_RICH_GRANULE_LUMEN" [color="#85C1E9"];
"gene_MRPL4" -- "term_2_GOCC_MITOCHONDRIAL_MATRIX" [color="#85C1E9"];
"gene_MRPS12" -- "term_2_GOCC_MITOCHONDRIAL_MATRIX" [color="#85C1E9"];
"gene_MXI1" -- "term_4_GOCC_TRANSCRIPTION_REGULATOR_COMPLEX" [color="#85C1E9"];
"gene_NARF" -- "term_6_GOCC_NUCLEAR_ENVELOPE" [color="#85C1E9"];
"gene_NDRG1" -- "term_3_GOCC_RECYCLING_ENDOSOME" [color="#85C1E9"];
"gene_NR3C1" -- "term_2_GOCC_MITOCHONDRIAL_MATRIX" [color="#85C1E9"];
"gene_NXF1" -- "term_6_GOCC_NUCLEAR_ENVELOPE" [color="#85C1E9"];
"gene_P4HA1" -- "term_1_GOCC_ENDOPLASMIC_RETICULUM_LUMEN" [color="#85C1E9"];
"gene_P4HA2" -- "term_1_GOCC_ENDOPLASMIC_RETICULUM_LUMEN" [color="#85C1E9"];
"gene_PGAM1" -- "term_5_GOCC_FICOLIN_1_RICH_GRANULE_LUMEN" [color="#85C1E9"];
"gene_PGK1" -- "term_2_GOCC_MITOCHONDRIAL_MATRIX" [color="#85C1E9"];
"gene_RAB8B" -- "term_3_GOCC_RECYCLING_ENDOSOME" [color="#85C1E9"];
"gene_RARA" -- "term_4_GOCC_TRANSCRIPTION_REGULATOR_COMPLEX" [color="#85C1E9"];
"gene_RBPJ" -- "term_4_GOCC_TRANSCRIPTION_REGULATOR_COMPLEX" [color="#85C1E9"];
"gene_SPAG4" -- "term_6_GOCC_NUCLEAR_ENVELOPE" [color="#85C1E9"];
"gene_STC2" -- "term_1_GOCC_ENDOPLASMIC_RETICULUM_LUMEN" [color="#85C1E9"];
"gene_TPCN1" -- "term_3_GOCC_RECYCLING_ENDOSOME" [color="#85C1E9"];
"gene_TUBG1" -- "term_3_GOCC_RECYCLING_ENDOSOME" [color="#85C1E9"];
"gene_VDAC1" -- "term_2_GOCC_MITOCHONDRIAL_MATRIX" [color="#85C1E9"];
"gene_CLK3" [label="CLK3", type="gene", width=0.7, height=0.7];
"gene_GRK6" [label="GRK6", type="gene", width=0.7, height=0.7];
"gene_INSIG2" [label="INSIG2", type="gene", width=0.7, height=0.7];
"gene_OXSR1" [label="OXSR1", type="gene", width=0.7, height=0.7];
"gene_RIOX1" [label="RIOX1", type="gene", width=0.7, height=0.7];
"gene_RSBN1" [label="RSBN1", type="gene", width=0.7, height=0.7];
"term_1_GOMF_2_OXOGLUTARATE_DEPENDENT_DIOXYGENASE_ACTIVITY" [label="GOMF: 2 OXOGLUTARATE DEPENDENT DIOXYGENASE ACTIVITY", style=filled, fillcolor="#D7BDE2", fontcolor="white", type="term", width=2, height=2, library="C5: Gene Ontology: Molecular Function"];
"term_2_GOMF_PROTEIN_SERINE_THREONINE_KINASE_ACTIVITY" [label="GOMF: PROTEIN SERINE THREONINE KINASE ACTIVITY", style=filled, fillcolor="#D7BDE2", fontcolor="white", type="term", width=2, height=2, library="C5: Gene Ontology: Molecular Function"];
"term_3_GOMF_STEROID_BINDING" [label="GOMF: STEROID BINDING", style=filled, fillcolor="#D7BDE2", fontcolor="white", type="term", width=2, height=2, library="C5: Gene Ontology: Molecular Function"];
"gene_ASPH" -- "term_1_GOMF_2_OXOGLUTARATE_DEPENDENT_DIOXYGENASE_ACTIVITY" [color="#D7BDE2"];
"gene_CCNB1" -- "term_2_GOMF_PROTEIN_SERINE_THREONINE_KINASE_ACTIVITY" [color="#D7BDE2"];
"gene_CLK3" -- "term_2_GOMF_PROTEIN_SERINE_THREONINE_KINASE_ACTIVITY" [color="#D7BDE2"];
"gene_GRK6" -- "term_2_GOMF_PROTEIN_SERINE_THREONINE_KINASE_ACTIVITY" [color="#D7BDE2"];
"gene_INSIG2" -- "term_3_GOMF_STEROID_BINDING" [color="#D7BDE2"];
"gene_KDM3A" -- "term_1_GOMF_2_OXOGLUTARATE_DEPENDENT_DIOXYGENASE_ACTIVITY" [color="#D7BDE2"];
"gene_KDM4B" -- "term_1_GOMF_2_OXOGLUTARATE_DEPENDENT_DIOXYGENASE_ACTIVITY" [color="#D7BDE2"];
"gene_KDM4C" -- "term_1_GOMF_2_OXOGLUTARATE_DEPENDENT_DIOXYGENASE_ACTIVITY" [color="#D7BDE2"];
"gene_NR3C1" -- "term_3_GOMF_STEROID_BINDING" [color="#D7BDE2"];
"gene_OXSR1" -- "term_2_GOMF_PROTEIN_SERINE_THREONINE_KINASE_ACTIVITY" [color="#D7BDE2"];
"gene_P4HA1" -- "term_1_GOMF_2_OXOGLUTARATE_DEPENDENT_DIOXYGENASE_ACTIVITY" [color="#D7BDE2"];
"gene_P4HA2" -- "term_1_GOMF_2_OXOGLUTARATE_DEPENDENT_DIOXYGENASE_ACTIVITY" [color="#D7BDE2"];
"gene_PGK1" -- "term_2_GOMF_PROTEIN_SERINE_THREONINE_KINASE_ACTIVITY" [color="#D7BDE2"];
"gene_PIM1" -- "term_2_GOMF_PROTEIN_SERINE_THREONINE_KINASE_ACTIVITY" [color="#D7BDE2"];
"gene_PLOD1" -- "term_1_GOMF_2_OXOGLUTARATE_DEPENDENT_DIOXYGENASE_ACTIVITY" [color="#D7BDE2"];
"gene_RIOX1" -- "term_1_GOMF_2_OXOGLUTARATE_DEPENDENT_DIOXYGENASE_ACTIVITY" [color="#D7BDE2"];
"gene_RSBN1" -- "term_1_GOMF_2_OXOGLUTARATE_DEPENDENT_DIOXYGENASE_ACTIVITY" [color="#D7BDE2"];
"gene_TGFBR1" -- "term_2_GOMF_PROTEIN_SERINE_THREONINE_KINASE_ACTIVITY" [color="#D7BDE2"];
"gene_VDAC1" -- "term_3_GOMF_STEROID_BINDING" [color="#D7BDE2"];
"gene_NF2" [label="NF2", type="gene", width=0.7, height=0.7];
"term_1_HP_AORTIC_DISSECTION" [label="HP: AORTIC DISSECTION", style=filled, fillcolor="#F7DC6F", fontcolor="white", type="term", width=2, height=2, library="C5: Human Phenotype Ontology"];
"term_2_HP_ABNORMAL_CRANIAL_NERVE_MORPHOLOGY" [label="HP: ABNORMAL CRANIAL NERVE MORPHOLOGY", style=filled, fillcolor="#F7DC6F", fontcolor="white", type="term", width=2, height=2, library="C5: Human Phenotype Ontology"];
"gene_FH" -- "term_2_HP_ABNORMAL_CRANIAL_NERVE_MORPHOLOGY" [color="#F7DC6F"];
"gene_LOX" -- "term_1_HP_AORTIC_DISSECTION" [color="#F7DC6F"];
"gene_NF2" -- "term_2_HP_ABNORMAL_CRANIAL_NERVE_MORPHOLOGY" [color="#F7DC6F"];
"gene_NR3C1" -- "term_2_HP_ABNORMAL_CRANIAL_NERVE_MORPHOLOGY" [color="#F7DC6F"];
"gene_P4HA2" -- "term_1_HP_AORTIC_DISSECTION" [color="#F7DC6F"];
"gene_PLOD1" -- "term_1_HP_AORTIC_DISSECTION" [color="#F7DC6F"];
"gene_TGFBR1" -- "term_1_HP_AORTIC_DISSECTION" [color="#F7DC6F"];
}

**NETWORK INTERPRETATION GUIDE:**

**Node Types:**
- **Gene nodes** (type="gene"): Individual genes from the input gene set
- **Term nodes** (type="term"): Enriched biological pathways/processes/gene sets
- **Library Source**: Each term node includes a "library" attribute indicating the source database

**Library Sources and Their Biological Context:**

- **H: Hallmark gene sets**: Curated gene sets representing well-defined biological states or processes
- **C2: BioCarta**: Canonical pathways from BioCarta database
- **C2: KEGG MEDICUS**: Metabolic and signaling pathways from KEGG
- **C2: Pathway Interaction Database**: Curated human signaling pathways
- **C2: Reactome Pathways**: Expert-curated biological pathways
- **C2: WikiPathways**: Community-curated biological pathways
- **C5: Gene Ontology: Biological Process**: Biological processes from Gene Ontology
- **C5: Gene Ontology: Cellular Component**: Cellular components from Gene Ontology
- **C5: Gene Ontology: Molecular Function**: Molecular functions from Gene Ontology
- **C5: Human Phenotype Ontology**: Human phenotypes and diseases
- **Protein Interaction**: Protein-protein interaction networks

**Edge Connections:**
- Each edge (gene -- term) represents a gene's membership in that biological term
- More connections indicate genes that are part of multiple enriched processes
- The network topology shows which genes are central to the biological response

**Iteration Information:**
- Term nodes are labeled with iteration numbers (term_1_, term_2_, etc.)
- Earlier iterations (lower numbers) represent the most statistically significant enrichments
- Ignore iteration number for biological interpretation, instead focus on network hubs densely connected for interesting biological insights

**ANALYSIS REQUEST:**

Please analyze this network and provide:

1. **Key Biological Insights:**
   - What are the most significant biological processes/pathways identified?
   - Which genes appear to be central to the biological response?

2. **Network Topology Analysis:**
   - Which genes are "hub" genes (connected to multiple terms)?
   - Are there distinct clusters or modules in the network?
   - What does the connectivity pattern suggest about biological organization?

3. **Biological Hypothesis:**

- Based on the network structure, what biological hypothesis can you generate?
- How do the different library sources support or complement each other?
- Try to formulate one hypothesis based on the most central terms and how they possibly related to one another.

4. **Estimated Experimental Context**
   - Can you hypothesize on what was the experiment that generated this result?

**RESPONSE STRUCTURE:**
- **Executive Summary** (2-3 sentences)
- **Key Biological Processes Identified** (by library source)
- **Network Topology Insights**
- **Proposed Biological Hypothesis**
- **Estimated Experimental Context**

**IMPORTANT NOTES:**
- Focus on biological interpretation, not statistical significance or iteration number
- Consider the functional relationships between connected genes and terms
- Look for unexpected connections that might reveal novel or unexpected biology
- Consider the broader biological context and literature
- Note that protein interaction library could shed light on physical interaction between genes while terms indicate functional interaction
- Different library sources may provide complementary biological insights

—

**Executive Summary**

This iterative enrichment network is dominated by a **hypoxia/HIF-driven program** tightly coupled to a **glycolytic metabolic shift**, with a second strong axis of **extracellular matrix (ECM)/collagen biogenesis and remodeling**. A smaller but coherent set of links points to **p53/cell-cycle control** and **chromatin/histone demethylation**, suggesting a coordinated stress-adaptation state rather than a single isolated pathway.

**Key Biological Processes Identified (by library source)**

**H: Hallmark Gene Sets**

- **HALLMARK: HYPOXIA** is the single most central "state" term and captures many of the network's high-degree genes.
  - Core hypoxia/metabolic genes: **ALDOC, GAPDH, LDHA, PGK1, DDIT4, NDRG1, LOX, P4HA1, P4HA2, ERO1A, FAM162A, CXCR4, STC2, WSB1**, etc.

- Interpretation: a classic oxygen-stress transcriptional program (HIF-centric) with downstream metabolic rewiring and remodeling of the microenvironment.

**C2: KEGG (Legacy + MEDICUS)**

- **KEGG: GLYCOLYSIS / GLUCONEOGENESIS** and **KEGG MEDICUS: GLYCOLYSIS** reinforce the metabolic core.
  - Strong overlap: **ALDOC, GAPDH, LDHA, PGAM1, PGK1**.
- **KEGG: p53 signaling** appears as an additional stress-control node:
  - **CCNB1, CYCS, GADD45B** (and conceptually ties to checkpointing/apoptosis and damage response).

**C2: Pathway Interaction Database (PID)**

- **PID: HIF1 TF pathway** matches the Hallmark hypoxia theme directly:
  - **BNIP3, CXCR4, LDHA, NDRG1, PGK1** (canonical HIF outputs).
- **PID: MYC activation** adds a proliferative/metabolic regulator layer:
  - **CAD, CCNB1, GAPDH, PIM1** (a plausible MYC-linked anabolic/proliferative overlay on the hypoxia state).

**C2: Reactome**

Reactome splits the biology into interpretable modules:

- **Gluconeogenesis**: **ALDOC, GAPDH, LDHA, PGAM1, PGK1** (metabolic axis).
- **Collagen formation**: **LOX, P4HA1, P4HA2, PLOD1** (ECM maturation/crosslinking axis).
- **HDMs demethylate histones**: **KDM3A, KDM4B, KDM4C** (epigenetic/chromatin adaptation).
- **TP53 regulates metabolic genes**: **CYCS, DDIT4, RRAGD** (stress → metabolism coupling).
- **PLK1 activity at G2/M transition**: **CCNB1 + cytoskeletal/spindle-associated genes (e.g., TUBG1, ACTR1A)** (cell-cycle machinery).

**C2: WikiPathways**

These terms “contextualize” the same genes into cellular programs:

- **Glycolysis in senescence**: again anchors **ALDOC/GAPDH/LDHA/PGK1** (stress-associated glycolytic phenotype).
- **Metabolic pathways of fibroblasts**: ties metabolism to stromal/ECM biology (**P4HA1/2, PLOD1, SLC16A1**).
- **Hematopoietic stem cell differentiation**: links **CXCR4, MXI1, PIM1** to a differentiation/migration axis.
- **Adipogenesis**: connects **NR3C1, RARA, NAMPT, GADD45B, MIF** (nuclear receptor signaling + stress metabolism).

**C5: Gene Ontology (Biological Process)**

GO-BP terms are consistent with the above but add “meta-annotations”:

- **Cellular response to oxygen levels**: enriched with **BNIP3, ERO1A, FAM162A, HILPDA, NDRG1, STC2**, etc.
- **Pyruvate metabolic process**: captures glycolytic flux genes (**ALDOC, GAPDH, LDHA, PGAM1, PGK1, DDIT4, SLC16A1**).
- **Collagen fibril organization**: ECM module (**LOX, P4HA1, P4HA2, PLOD1, TGFBR1**).
- **Response to starvation** and **positive regulation of MAPK cascade**: stress signaling overlay (**ATF3, CAD, KLF10, RRAGD**, plus **EFNA1/GADD45B/MIF/NAMPT/PJA2**).

**C5: Gene Ontology (Cellular Component)**

Cellular localization hints at organelle-level stress:

- **Endoplasmic reticulum lumen**: **ERO1A, ASPH, EDEM3, P4HA1/2, STC2** (protein folding/processing + collagen modification machinery fits here).
- **Mitochondrial matrix**: **FH, MRPL4, MRPS12, VDAC1, PGK1, CCNB1** (energy/stress and mitochondrial coupling).
- **Nuclear envelope**: includes **BNIP3, GAPDH, NARF, NXF1, SPAG4** (stress-associated nuclear architecture/transport; not the dominant biology but a consistent signal).
- **Recycling endosome**: **NDRG1, RAB8B, TPCN1, TUBG1** (trafficking/membrane remodeling; plausibly tied to migration/adaptation).

**C5: Gene Ontology (Molecular Function)**

- **2-oxoglutarate–dependent dioxygenase activity**: **KDM3A/KDM4B/KDM4C** plus collagen-related enzymes (**P4HA1/2, PLOD1**) and **ASPH/RIOX1/RSBN1**.
    - This is a *nice mechanistic bridge*: oxygen- and 2OG-dependent enzymes are directly impacted by hypoxia and can mediate adaptive transcriptional and ECM changes.
- **Protein serine/threonine kinase activity**: **PIM1, OXSR1, GRK6, CLK3, TGFBR1, PGK1** (signaling control + stress kinases).
- **Steroid binding**: **NR3C1** (glucocorticoid receptor) and **INSIG2/VDAC1**-linked annotations reinforce nuclear receptor/metabolic regulation.

**C5: Human Phenotype Ontology**

- **Aortic dissection** connects to **LOX, P4HA2, PLOD1, TGFBR1**.
    - This is a strong phenotypic “readout” of the **collagen/ECM crosslinking + TGF-β** module (vascular wall integrity).
- **Abnormal cranial nerve morphology**: linked here via **FH, NF2, NR3C1** (less central; likely reflects pleiotropy of broadly important genes).

---

**Network Topology Insights**

**1) Hub genes (high connectivity across multiple terms/libraries)**

**Major hubs (appear repeatedly across hypoxia + glycolysis terms):**

- **ALDOC, GAPDH, LDHA, PGK1, PGAM1**

These form the **metabolic backbone** appearing in Hallmark Hypoxia + KEGG glycolysis + Reactome gluconeogenesis + WP glycolysis-in-senescence + GO pyruvate metabolism.

**ECM/collagen module hubs:**

- **LOX, P4HA1, P4HA2, PLOD1, TGFBR1**
  These show up in Reactome collagen formation + GO collagen fibril organization + WP fibroblast metabolism + HPO aortic dissection, making a very coherent second hub cluster.

**Stress-regulatory and state-control hubs:**

- **DDIT4, NDRG1, BNIP3, CXCR4, STC2, ERO1A, FAM162A**
  These are classic hypoxia-adaptation genes (metabolism, survival, migration, ER stress/protein processing).

**Cell-cycle / checkpoint hubs (smaller module):**

- **CCNB1, CYCS, GADD45B**
  They anchor the p53/cell-cycle terms (KEGG p53, Reactome PLK1/G2M, GO oxygen response in one case).

**Epigenetic hubs:**

- **KDM3A, KDM4B, KDM4C**
  Cluster tightly around the histone demethylation/2OG-dioxygenase activity terms.

**2) Clusters/modules you can read directly off connectivity**

You can basically partition the network into **four modules**:

1. **Hypoxia–glycolysis module (largest, densest)**
   - Central term: Hallmark Hypoxia
   - Reinforced by KEGG glycolysis, Reactome gluconeogenesis, GO pyruvate metabolism, WP glycolysis in senescence.
2. **ECM/collagen remodeling module**
   - Central term: Reactome collagen formation + GO collagen fibril organization
   - Phenotype anchor: HPO aortic dissection
   - Mechanistic tie: ER lumen localization (collagen processing) + 2OG dioxygenases.
3. **Stress checkpoint / p53–cell cycle module**
   - KEGG p53 + Reactome PLK1/G2M + Reactome TP53 metabolic genes
   - Suggests either mixed cell states (some proliferative) or coordinated checkpoint control under stress.
4. **Chromatin adaptation module**
   - Reactome histone demethylation + GO 2OG dioxygenase activity
   - Likely a regulatory layer enabling durable transcriptional remodeling under oxygen/nutrient stress.

**3) What the connectivity pattern suggests biologically**

- The network is **not random multi-pathway noise**: it is **two dominant, functionally compatible programs** (hypoxia metabolism + ECM remodeling), with **regulatory layers** (p53/cell-cycle and chromatin enzymes) that make sense for cells adapting to stress.
- The iterative enrichment design makes this clearer: within each library, you're seeing the **top "explainers"** of the remaining genes. The repeated appearance of the same themes across independent libraries argues the biology is robust.

---

**Proposed Biological Hypothesis**

A single integrative hypothesis that matches the central terms and hubs:

**Hypothesis:**
Your gene list reflects a **hypoxia-driven adaptive remodeling state** in which **HIF signaling** induces a **glycolytic shift** (ALDOC/GAPDH/LDHA/PGK1/PGAM1) and simultaneously activates an **ECM maturation/remodeling program** (P4HA1/2, PLOD1, LOX) potentially mediated or reinforced by **TGF-β signaling (TGFBR1)**. This state is stabilized by **chromatin remodeling via 2OG-dependent histone demethylases (KDM3A/KDM4B/KDM4C)** and constrained/reshaped by **stress checkpoint pathways (p53/cell-cycle components like CCNB1, GADD45B, CYCS)**.

**How the libraries complement each other**

- **Hallmark + PID** give the *state driver*: hypoxia/HIF and MYC.
- **KEGG + Reactome** provide *mechanistic detail*: glycolysis/gluconeogenesis, collagen formation, p53 metabolic regulation, G2/M control.
- **GO (BP/CC/MF)** supplies *cellular interpretation*: pyruvate metabolism, oxygen response, ER lumen (collagen processing), and the key enzyme class (2OG dioxygenases).
- **WikiPathways** adds *cell-type/program framing*: fibroblast metabolism and senescence-associated glycolysis.
- **HPO** offers a *phenotypic readout* that is unusually informative here: the collagen/LOX/TGFBR1 axis mapping onto vascular wall integrity (aortic dissection) strongly supports the ECM remodeling module being real and biologically consequential.

---

**Estimated Experimental Context**

Most plausible experimental scenarios that would generate this specific combination (hypoxia + glycolysis + collagen remodeling + TGF-β + epigenetic dioxygenases):

1. **Cells exposed to hypoxia (or hypoxia mimetic) with microenvironment remodeling**
   - Example contexts: tumor cells, stromal fibroblasts, or endothelial/perivascular cells under low $O_2$.
   - The **LOX/P4HA/PLOD1** ECM maturation signature is especially common in **hypoxic tumors** and **activated stroma**.
2. **Fibroblast activation / pro-fibrotic signaling under metabolic stress**

    - The presence of **TGFBR1** plus multiple collagen-modifying enzymes and WP "metabolic pathways of fibroblasts" fits **TGF-β–driven fibroblast remodeling**, potentially in low oxygen (e.g., wound healing, fibrosis-like programs).
3. **Stress adaptation with partial checkpoint engagement**
    - The p53/G2M elements suggest either:
        - a mixed population (some proliferating, some stressed), or
        - a synchronized stress response where proliferation is being regulated rather than fully shut down.

If I had to pick **one**: this looks most like **transcriptional profiling (bulk RNA-seq or microarray)** of cells in a **hypoxic / ischemic-like condition** (or a solid-tumor microenvironment model), with a strong **ECM remodeling** component consistent with **stroma activation or invasive adaptation**.

---

If you want, paste the *full* original 75-gene list (or confirm that the genes shown are the full set that survived to the DOT output), and I can:

- rank hubs quantitatively from the DOT (degree centrality),
- explicitly label the modules and list "module-defining" genes,
- and suggest 3–5 targeted follow-up experiments (e.g., HIF inhibition vs TGFBR1 inhibition, collagen crosslinking assays, lactate flux).

## Supplementary Table 11

Vaccination signature AI response

Prompt, network and full ChatGPT response for the 426-gene influenza vaccination signature.

Document Content:
Structure iGEA Prompt
ChatGPT Response
Follow up with experiment context
ChatGPT response to experiment context
Follow up: what is inconsistent or novel?
ChatGPT Response to inconsistency and novelty

You are a computational biologist analyzing an iterative gene set enrichment network. Iterative means that for each library source, the top enriched gene set is found, saved and the genes are removed from the initial gene list. The remaining genes are then tested for enrichment again. Thus, each gene appears only once per library source tested but can be linked to multiple terms that originate from different libraries. The results are provided as a DOT network represents the relationships between genes and gene sets(=terms).

The original gene list included 426 genes and was tested across the following libraries: H: Hallmark Gene Sets, C2: KEGG Legacy, C2: KEGG MEDICUS, C2: Pathway Interaction Database, C2: Reactome Pathways, C2: WikiPathways, C5: Gene Ontology: Biological Process, C5: Gene Ontology: Cellular Component, C5: Gene Ontology: Molecular Function, C5: Human Phenotype Ontology.

**NETWORK STRUCTURE:**

```
graph iterative_enrichment_all {
graph [layout=neato];
node [shape=ellipse];
subgraph cluster_legend_libs {
label="Libraries";
node [shape=box, style=filled, fontcolor="white"];
legend_lib_1 [label="H: Hallmark Gene Sets", fillcolor="#FF6B6B"];
legend_lib_2 [label="C2: KEGG Legacy", fillcolor="#F8C471"];
legend_lib_3 [label="C2: KEGG MEDICUS", fillcolor="#98D8C8"];
legend_lib_4 [label="C2: Pathway Interaction Database", fillcolor="#440154"];
legend_lib_5 [label="C2: Reactome Pathways", fillcolor="#BB8FCE"];
legend_lib_6 [label="C2: WikiPathways", fillcolor="#31688e"];
legend_lib_7 [label="C5: Gene Ontology: Biological Process", fillcolor="#F1948A"];
legend_lib_8 [label="C5: Gene Ontology: Cellular Component", fillcolor="#85C1E9"];
legend_lib_9 [label="C5: Gene Ontology: Molecular Function", fillcolor="#D7BDE2"];
legend_lib_10 [label="C5: Human Phenotype Ontology", fillcolor="#F7DC6F"];
}
"gene_ACLY" [label="ACLY", type="gene", width=0.7, height=0.7];
"gene_ACTR3" [label="ACTR3", type="gene", width=0.7, height=0.7];
"gene_ADIPOR2" [label="ADIPOR2", type="gene", width=0.7, height=0.7];
"gene_CASP1" [label="CASP1", type="gene", width=0.7, height=0.7];
"gene_CASP10" [label="CASP10", type="gene", width=0.7, height=0.7];
"gene_CD36" [label="CD36", type="gene", width=0.7, height=0.7];
"gene_CORO1A" [label="CORO1A", type="gene", width=0.7, height=0.7];
"gene_CR1" [label="CR1", type="gene", width=0.7, height=0.7];
"gene_CSF2RB" [label="CSF2RB", type="gene", width=0.7, height=0.7];
"gene_CSF3R" [label="CSF3R", type="gene", width=0.7, height=0.7];
"gene_CTSD" [label="CTSD", type="gene", width=0.7, height=0.7];
"gene_FGL2" [label="FGL2", type="gene", width=0.7, height=0.7];
"gene_GCA" [label="GCA", type="gene", width=0.7, height=0.7];
"gene_GLRX" [label="GLRX", type="gene", width=0.7, height=0.7];
"gene_HMBS" [label="HMBS", type="gene", width=0.7, height=0.7];
"gene_IDH1" [label="IDH1", type="gene", width=0.7, height=0.7];
"gene_IFI30" [label="IFI30", type="gene", width=0.7, height=0.7];
"gene_IFNAR1" [label="IFNAR1", type="gene", width=0.7, height=0.7];
"gene_IFNGR2" [label="IFNGR2", type="gene", width=0.7, height=0.7];
"gene_IL10RB" [label="IL10RB", type="gene", width=0.7, height=0.7];
"gene_IL17RA" [label="IL17RA", type="gene", width=0.7, height=0.7];
"gene_IRF2" [label="IRF2", type="gene", width=0.7, height=0.7];
"gene_LAMP2" [label="LAMP2", type="gene", width=0.7, height=0.7];
"gene_LRP1" [label="LRP1", type="gene", width=0.7, height=0.7];
"gene_M6PR" [label="M6PR", type="gene", width=0.7, height=0.7];
"gene_MAP2K3" [label="MAP2K3", type="gene", width=0.7, height=0.7];
"gene_MSRB1" [label="MSRB1", type="gene", width=0.7, height=0.7];
"gene_PPP2CB" [label="PPP2CB", type="gene", width=0.7, height=0.7];
"gene_PRCP" [label="PRCP", type="gene", width=0.7, height=0.7];
"gene_PRDX1" [label="PRDX1", type="gene", width=0.7, height=0.7];
"gene_PRKCD" [label="PRKCD", type="gene", width=0.7, height=0.7];
"gene_PSEN1" [label="PSEN1", type="gene", width=0.7, height=0.7];
"gene_SERPINA1" [label="SERPINA1", type="gene", width=0.7, height=0.7];
"gene_SKAP2" [label="SKAP2", type="gene", width=0.7, height=0.7];
"gene_SLC6A6" [label="SLC6A6", type="gene", width=0.7, height=0.7];
"gene_STAM2" [label="STAM2", type="gene", width=0.7, height=0.7];
"gene_TIMP2" [label="TIMP2", type="gene", width=0.7, height=0.7];
"gene_TNFRSF1A" [label="TNFRSF1A", type="gene", width=0.7, height=0.7];
"gene_TPI1" [label="TPI1", type="gene", width=0.7, height=0.7];
"term_1_HALLMARK_COMPLEMENT" [label="HALLMARK: COMPLEMENT", style=filled, fillcolor="#FF6B6B", fontcolor="white", type="term", width=2, height=2, library="H: Hallmark Gene Sets"];
"term_2_HALLMARK_MTORC1_SIGNALING" [label="HALLMARK: MTORC1 SIGNALING", style=filled, fillcolor="#FF6B6B", fontcolor="white", type="term", width=2, height=2, library="H: Hallmark Gene Sets"];
"term_3_HALLMARK_IL6_JAK_STAT3_SIGNALING" [label="HALLMARK: IL6 JAK STAT3 SIGNALING", style=filled, fillcolor="#FF6B6B", fontcolor="white", type="term", width=2, height=2, library="H: Hallmark Gene Sets"];
"gene_ACLY" -- "term_2_HALLMARK_MTORC1_SIGNALING" [color="#FF6B6B"];
"gene_ACTR3" -- "term_2_HALLMARK_MTORC1_SIGNALING" [color="#FF6B6B"];
"gene_ADIPOR2" -- "term_2_HALLMARK_MTORC1_SIGNALING" [color="#FF6B6B"];
"gene_CASP1" -- "term_1_HALLMARK_COMPLEMENT" [color="#FF6B6B"];
"gene_CASP10" -- "term_1_HALLMARK_COMPLEMENT" [color="#FF6B6B"];
"gene_CD36" -- "term_1_HALLMARK_COMPLEMENT" [color="#FF6B6B"];
"gene_CORO1A" -- "term_2_HALLMARK_MTORC1_SIGNALING" [color="#FF6B6B"];
"gene_CR1" -- "term_1_HALLMARK_COMPLEMENT" [color="#FF6B6B"];
"gene_CSF2RB" -- "term_3_HALLMARK_IL6_JAK_STAT3_SIGNALING" [color="#FF6B6B"];
"gene_CSF3R" -- "term_3_HALLMARK_IL6_JAK_STAT3_SIGNALING" [color="#FF6B6B"];
"gene_CTSD" -- "term_1_HALLMARK_COMPLEMENT" [color="#FF6B6B"];
"gene_FGL2" -- "term_2_HALLMARK_MTORC1_SIGNALING" [color="#FF6B6B"];
"gene_GCA" -- "term_1_HALLMARK_COMPLEMENT" [color="#FF6B6B"];
"gene_GLRX" -- "term_2_HALLMARK_MTORC1_SIGNALING" [color="#FF6B6B"];
"gene_HMBS" -- "term_2_HALLMARK_MTORC1_SIGNALING" [color="#FF6B6B"];
"gene_IDH1" -- "term_2_HALLMARK_MTORC1_SIGNALING" [color="#FF6B6B"];
"gene_IFI30" -- "term_2_HALLMARK_MTORC1_SIGNALING" [color="#FF6B6B"];
"gene_IFNAR1" -- "term_3_HALLMARK_IL6_JAK_STAT3_SIGNALING" [color="#FF6B6B"];
"gene_IFNGR2" -- "term_3_HALLMARK_IL6_JAK_STAT3_SIGNALING" [color="#FF6B6B"];
"gene_IL10RB" -- "term_3_HALLMARK_IL6_JAK_STAT3_SIGNALING" [color="#FF6B6B"];
"gene_IL17RA" -- "term_3_HALLMARK_IL6_JAK_STAT3_SIGNALING" [color="#FF6B6B"];
"gene_IRF2" -- "term_1_HALLMARK_COMPLEMENT" [color="#FF6B6B"];
"gene_LAMP2" -- "term_1_HALLMARK_COMPLEMENT" [color="#FF6B6B"];
"gene_LRP1" -- "term_1_HALLMARK_COMPLEMENT" [color="#FF6B6B"];
"gene_M6PR" -- "term_2_HALLMARK_MTORC1_SIGNALING" [color="#FF6B6B"];
"gene_MAP2K3" -- "term_2_HALLMARK_MTORC1_SIGNALING" [color="#FF6B6B"];
"gene_MSRB1" -- "term_1_HALLMARK_COMPLEMENT" [color="#FF6B6B"];
"gene_PPP2CB" -- "term_1_HALLMARK_COMPLEMENT" [color="#FF6B6B"];
"gene_PRCP" -- "term_1_HALLMARK_COMPLEMENT" [color="#FF6B6B"];
"gene_PRDX1" -- "term_2_HALLMARK_MTORC1_SIGNALING" [color="#FF6B6B"];
"gene_PRKCD" -- "term_1_HALLMARK_COMPLEMENT" [color="#FF6B6B"];
"gene_PSEN1" -- "term_1_HALLMARK_COMPLEMENT" [color="#FF6B6B"];
"gene_SERPINA1" -- "term_1_HALLMARK_COMPLEMENT" [color="#FF6B6B"];
"gene_SKAP2" -- "term_2_HALLMARK_MTORC1_SIGNALING" [color="#FF6B6B"];
"gene_SLC6A6" -- "term_2_HALLMARK_MTORC1_SIGNALING" [color="#FF6B6B"];
"gene_STAM2" -- "term_3_HALLMARK_IL6_JAK_STAT3_SIGNALING" [color="#FF6B6B"];
"gene_TIMP2" -- "term_1_HALLMARK_COMPLEMENT" [color="#FF6B6B"];
"gene_TNFRSF1A" -- "term_3_HALLMARK_IL6_JAK_STAT3_SIGNALING" [color="#FF6B6B"];
"gene_TPI1" -- "term_2_HALLMARK_MTORC1_SIGNALING" [color="#FF6B6B"];
"gene_AP3S2" [label="AP3S2", type="gene", width=0.7, height=0.7];
"gene_APAF1" [label="APAF1", type="gene", width=0.7, height=0.7];
"gene_ARPC1B" [label="ARPC1B", type="gene", width=0.7, height=0.7];
"gene_ATP6AP1" [label="ATP6AP1", type="gene", width=0.7, height=0.7];
"gene_ATP6V0A1" [label="ATP6V0A1", type="gene", width=0.7, height=0.7];
"gene_ATP6V0D1" [label="ATP6V0D1", type="gene", width=0.7, height=0.7];
"gene_CLTA" [label="CLTA", type="gene", width=0.7, height=0.7];
"gene_DFFA" [label="DFFA", type="gene", width=0.7, height=0.7];
"gene_ENTPD4" [label="ENTPD4", type="gene", width=0.7, height=0.7];
"gene_FPGT" [label="FPGT", type="gene", width=0.7, height=0.7];
"gene_GAA" [label="GAA", type="gene", width=0.7, height=0.7];
"gene_GBA1" [label="GBA1", type="gene", width=0.7, height=0.7];
"gene_GNS" [label="GNS", type="gene", width=0.7, height=0.7];
"gene_GSN" [label="GSN", type="gene", width=0.7, height=0.7];
"gene_HCK" [label="HCK", type="gene", width=0.7, height=0.7];
"gene_HK1" [label="HK1", type="gene", width=0.7, height=0.7];
"gene_HK3" [label="HK3", type="gene", width=0.7, height=0.7];
"gene_MAPK1" [label="MAPK1", type="gene", width=0.7, height=0.7];
"gene_NAGA" [label="NAGA", type="gene", width=0.7, height=0.7];
"gene_NAGK" [label="NAGK", type="gene", width=0.7, height=0.7];
"gene_NCF1" [label="NCF1", type="gene", width=0.7, height=0.7];
"gene_NPL" [label="NPL", type="gene", width=0.7, height=0.7];
"gene_PAK1" [label="PAK1", type="gene", width=0.7, height=0.7];
"gene_PANK2" [label="PANK2", type="gene", width=0.7, height=0.7];
"gene_PPCDC" [label="PPCDC", type="gene", width=0.7, height=0.7];
"gene_PRKACA" [label="PRKACA", type="gene", width=0.7, height=0.7];
"gene_PRKAR1A" [label="PRKAR1A", type="gene", width=0.7, height=0.7];
"gene_SLC11A1" [label="SLC11A1", type="gene", width=0.7, height=0.7];
"gene_SORT1" [label="SORT1", type="gene", width=0.7, height=0.7];
"gene_SULT1A1" [label="SULT1A1", type="gene", width=0.7, height=0.7];
"gene_SULT1A3" [label="SULT1A3", type="gene", width=0.7, height=0.7];
```

```
"gene_SULT1A4" [label="SULT1A4", type="gene", width=0.7, height=0.7];
"gene_SYK" [label="SYK", type="gene", width=0.7, height=0.7];
"gene_TCIRG1" [label="TCIRG1", type="gene", width=0.7, height=0.7];
"gene_VAV1" [label="VAV1", type="gene", width=0.7, height=0.7];
"gene_VNN2" [label="VNN2", type="gene", width=0.7, height=0.7];
"term_1_KEGG_LYSOSOME" [label="KEGG: LYSOSOME", style=filled, fillcolor="#F8C471", fontcolor="white", type="term", width=2, height=2, library="C2: KEGG Legacy"];
"term_2_KEGG_FC_GAMMA_R_MEDIATED_PHAGOCYTOSIS" [label="KEGG: FC GAMMA R MEDIATED PHAGOCYTOSIS", style=filled, fillcolor="#F8C471", fontcolor="white", type="term", width=2, height=2, library="C2: KEGG Legacy"];
"term_3_KEGG_SULFUR_METABOLISM" [label="KEGG: SULFUR METABOLISM", style=filled, fillcolor="#F8C471", fontcolor="white", type="term", width=2, height=2, library="C2: KEGG Legacy"];
"term_4_KEGG_AMINO_SUGAR_AND_NUCLEOTIDE_SUGAR_METABOLISM" [label="KEGG: AMINO SUGAR AND NUCLEOTIDE SUGAR METABOLISM", style=filled, fillcolor="#F8C471", fontcolor="white", type="term", width=2, height=2, library="C2: KEGG Legacy"];
"term_5_KEGG_PANTOTHENATE_AND_COA_BIOSYNTHESIS" [label="KEGG: PANTOTHENATE AND COA BIOSYNTHESIS", style=filled, fillcolor="#F8C471", fontcolor="white", type="term", width=2, height=2, library="C2: KEGG Legacy"];
"term_6_KEGG_APOPTOSIS" [label="KEGG: APOPTOSIS", style=filled, fillcolor="#F8C471", fontcolor="white", type="term", width=2, height=2, library="C2: KEGG Legacy"];
"gene_AP3S2" -- "term_1_KEGG_LYSOSOME" [color="#F8C471"];
"gene_APAF1" -- "term_6_KEGG_APOPTOSIS" [color="#F8C471"];
"gene_ARPC1B" -- "term_2_KEGG_FC_GAMMA_R_MEDIATED_PHAGOCYTOSIS" [color="#F8C471"];
"gene_ATP6AP1" -- "term_1_KEGG_LYSOSOME" [color="#F8C471"];
"gene_ATP6V0A1" -- "term_1_KEGG_LYSOSOME" [color="#F8C471"];
"gene_ATP6V0D1" -- "term_1_KEGG_LYSOSOME" [color="#F8C471"];
"gene_CASP10" -- "term_6_KEGG_APOPTOSIS" [color="#F8C471"];
"gene_CLTA" -- "term_1_KEGG_LYSOSOME" [color="#F8C471"];
"gene_CSF2RB" -- "term_6_KEGG_APOPTOSIS" [color="#F8C471"];
"gene_CTSD" -- "term_1_KEGG_LYSOSOME" [color="#F8C471"];
"gene_DFFA" -- "term_6_KEGG_APOPTOSIS" [color="#F8C471"];
"gene_ENTPD4" -- "term_1_KEGG_LYSOSOME" [color="#F8C471"];
"gene_FPGT" -- "term_4_KEGG_AMINO_SUGAR_AND_NUCLEOTIDE_SUGAR_METABOLISM" [color="#F8C471"];
"gene_GAA" -- "term_1_KEGG_LYSOSOME" [color="#F8C471"];
"gene_GBA1" -- "term_1_KEGG_LYSOSOME" [color="#F8C471"];
"gene_GNS" -- "term_1_KEGG_LYSOSOME" [color="#F8C471"];
"gene_GSN" -- "term_2_KEGG_FC_GAMMA_R_MEDIATED_PHAGOCYTOSIS" [color="#F8C471"];
"gene_HCK" -- "term_2_KEGG_FC_GAMMA_R_MEDIATED_PHAGOCYTOSIS" [color="#F8C471"];
"gene_HK1" -- "term_4_KEGG_AMINO_SUGAR_AND_NUCLEOTIDE_SUGAR_METABOLISM" [color="#F8C471"];
"gene_HK3" -- "term_4_KEGG_AMINO_SUGAR_AND_NUCLEOTIDE_SUGAR_METABOLISM" [color="#F8C471"];
"gene_LAMP2" -- "term_1_KEGG_LYSOSOME" [color="#F8C471"];
"gene_M6PR" -- "term_1_KEGG_LYSOSOME" [color="#F8C471"];
"gene_MAPK1" -- "term_2_KEGG_FC_GAMMA_R_MEDIATED_PHAGOCYTOSIS" [color="#F8C471"];
"gene_NAGA" -- "term_1_KEGG_LYSOSOME" [color="#F8C471"];
"gene_NAGK" -- "term_4_KEGG_AMINO_SUGAR_AND_NUCLEOTIDE_SUGAR_METABOLISM" [color="#F8C471"];
"gene_NCF1" -- "term_2_KEGG_FC_GAMMA_R_MEDIATED_PHAGOCYTOSIS" [color="#F8C471"];
"gene_NPL" -- "term_4_KEGG_AMINO_SUGAR_AND_NUCLEOTIDE_SUGAR_METABOLISM" [color="#F8C471"];
"gene_PAK1" -- "term_2_KEGG_FC_GAMMA_R_MEDIATED_PHAGOCYTOSIS" [color="#F8C471"];
"gene_PANK2" -- "term_5_KEGG_PANTOTHENATE_AND_COA_BIOSYNTHESIS" [color="#F8C471"];
"gene_PPCDC" -- "term_5_KEGG_PANTOTHENATE_AND_COA_BIOSYNTHESIS" [color="#F8C471"];
"gene_PRKACA" -- "term_6_KEGG_APOPTOSIS" [color="#F8C471"];
"gene_PRKAR1A" -- "term_6_KEGG_APOPTOSIS" [color="#F8C471"];
"gene_PRKCD" -- "term_2_KEGG_FC_GAMMA_R_MEDIATED_PHAGOCYTOSIS" [color="#F8C471"];
"gene_SLC11A1" -- "term_1_KEGG_LYSOSOME" [color="#F8C471"];
"gene_SORT1" -- "term_1_KEGG_LYSOSOME" [color="#F8C471"];
"gene_SULT1A1" -- "term_3_KEGG_SULFUR_METABOLISM" [color="#F8C471"];
"gene_SULT1A3" -- "term_3_KEGG_SULFUR_METABOLISM" [color="#F8C471"];
"gene_SULT1A4" -- "term_3_KEGG_SULFUR_METABOLISM" [color="#F8C471"];
"gene_SYK" -- "term_2_KEGG_FC_GAMMA_R_MEDIATED_PHAGOCYTOSIS" [color="#F8C471"];
"gene_TCIRG1" -- "term_1_KEGG_LYSOSOME" [color="#F8C471"];
"gene_TNFRSF1A" -- "term_6_KEGG_APOPTOSIS" [color="#F8C471"];
"gene_VAV1" -- "term_2_KEGG_FC_GAMMA_R_MEDIATED_PHAGOCYTOSIS" [color="#F8C471"];
"gene_VNN2" -- "term_5_KEGG_PANTOTHENATE_AND_COA_BIOSYNTHESIS" [color="#F8C471"];
"gene_CREB5" [label="CREB5", type="gene", width=0.7, height=0.7];
"gene_NCF2" [label="NCF2", type="gene", width=0.7, height=0.7];
"gene_NCF4" [label="NCF4", type="gene", width=0.7, height=0.7];
"term_1_KEGG_MEDICUS_REFERENCE_PRNP_PI3K_NOX2_SIGNALING_PATHWAY" [label="KEGG: MEDICUS REFERENCE PRNP PI3K NOX2 SIGNALING PATHWAY", style=filled, fillcolor="#98D8C8", fontcolor="white", type="term", width=2, height=2, library="C2: KEGG MEDICUS"];
"gene_CREB5" -- "term_1_KEGG_MEDICUS_REFERENCE_PRNP_PI3K_NOX2_SIGNALING_PATHWAY" [color="#98D8C8"];
"gene_MAPK1" -- "term_1_KEGG_MEDICUS_REFERENCE_PRNP_PI3K_NOX2_SIGNALING_PATHWAY" [color="#98D8C8"];
"gene_NCF1" -- "term_1_KEGG_MEDICUS_REFERENCE_PRNP_PI3K_NOX2_SIGNALING_PATHWAY" [color="#98D8C8"];
"gene_NCF2" -- "term_1_KEGG_MEDICUS_REFERENCE_PRNP_PI3K_NOX2_SIGNALING_PATHWAY" [color="#98D8C8"];
"gene_NCF4" -- "term_1_KEGG_MEDICUS_REFERENCE_PRNP_PI3K_NOX2_SIGNALING_PATHWAY" [color="#98D8C8"];
"gene_PRKCD" -- "term_1_KEGG_MEDICUS_REFERENCE_PRNP_PI3K_NOX2_SIGNALING_PATHWAY" [color="#98D8C8"];
"gene_ARAP1" [label="ARAP1", type="gene", width=0.7, height=0.7];
"gene_CSK" [label="CSK", type="gene", width=0.7, height=0.7];
"gene_IQGAP1" [label="IQGAP1", type="gene", width=0.7, height=0.7];
"gene_MAP2K4" [label="MAP2K4", type="gene", width=0.7, height=0.7];
"gene_PTPRJ" [label="PTPRJ", type="gene", width=0.7, height=0.7];
"gene_RBP1" [label="RBP1", type="gene", width=0.7, height=0.7];
"gene_RXRA" [label="RXRA", type="gene", width=0.7, height=0.7];
"gene_VDR" [label="VDR", type="gene", width=0.7, height=0.7];
"gene_YWHAE" [label="YWHAE", type="gene", width=0.7, height=0.7];
"term_1_PID_PDGFRB_PATHWAY" [label="PID: PDGFRB PATHWAY", style=filled, fillcolor="#440154", fontcolor="white", type="term", width=2, height=2, library="Unknown Library"];
"term_2_PID_CASPASE_PATHWAY" [label="PID: CASPASE PATHWAY", style=filled, fillcolor="#440154", fontcolor="white", type="term", width=2, height=2, library="Unknown Library"];
"term_3_PID_RETINOIC_ACID_PATHWAY" [label="PID: RETINOIC ACID PATHWAY", style=filled, fillcolor="#440154", fontcolor="white", type="term", width=2, height=2, library="Unknown Library"];
"gene_ACTR3" -- "term_1_PID_PDGFRB_PATHWAY" [color="#440154"];
"gene_APAF1" -- "term_2_PID_CASPASE_PATHWAY" [color="#440154"];
"gene_ARAP1" -- "term_1_PID_PDGFRB_PATHWAY" [color="#440154"];
"gene_ARPC1B" -- "term_1_PID_PDGFRB_PATHWAY" [color="#440154"];
"gene_CASP1" -- "term_2_PID_CASPASE_PATHWAY" [color="#440154"];
"gene_CASP10" -- "term_2_PID_CASPASE_PATHWAY" [color="#440154"];
"gene_CSK" -- "term_1_PID_PDGFRB_PATHWAY" [color="#440154"];
"gene_DFFA" -- "term_2_PID_CASPASE_PATHWAY" [color="#440154"];
"gene_GSN" -- "term_2_PID_CASPASE_PATHWAY" [color="#440154"];
"gene_HCK" -- "term_1_PID_PDGFRB_PATHWAY" [color="#440154"];
"gene_IQGAP1" -- "term_1_PID_PDGFRB_PATHWAY" [color="#440154"];
"gene_LRP1" -- "term_1_PID_PDGFRB_PATHWAY" [color="#440154"];
"gene_MAP2K4" -- "term_1_PID_PDGFRB_PATHWAY" [color="#440154"];
"gene_MAPK1" -- "term_1_PID_PDGFRB_PATHWAY" [color="#440154"];
"gene_PAK1" -- "term_1_PID_PDGFRB_PATHWAY" [color="#440154"];
"gene_PRKACA" -- "term_3_PID_RETINOIC_ACID_PATHWAY" [color="#440154"];
"gene_PRKCD" -- "term_1_PID_PDGFRB_PATHWAY" [color="#440154"];
"gene_PTPRJ" -- "term_1_PID_PDGFRB_PATHWAY" [color="#440154"];
"gene_RBP1" -- "term_3_PID_RETINOIC_ACID_PATHWAY" [color="#440154"];
"gene_RXRA" -- "term_3_PID_RETINOIC_ACID_PATHWAY" [color="#440154"];
"gene_TNFRSF1A" -- "term_2_PID_CASPASE_PATHWAY" [color="#440154"];
"gene_VDR" -- "term_3_PID_RETINOIC_ACID_PATHWAY" [color="#440154"];
"gene_YWHAE" -- "term_1_PID_PDGFRB_PATHWAY" [color="#440154"];
"gene_ADA2" [label="ADA2", type="gene", width=0.7, height=0.7];
"gene_ADCY7" [label="ADCY7", type="gene", width=0.7, height=0.7];
"gene_AHCYL1" [label="AHCYL1", type="gene", width=0.7, height=0.7];
"gene_ANPEP" [label="ANPEP", type="gene", width=0.7, height=0.7];
"gene_APEH" [label="APEH", type="gene", width=0.7, height=0.7];
"gene_ARF3" [label="ARF3", type="gene", width=0.7, height=0.7];
"gene_ARHGAP9" [label="ARHGAP9", type="gene", width=0.7, height=0.7];
"gene_ARSD" [label="ARSD", type="gene", width=0.7, height=0.7];
"gene_ASGR2" [label="ASGR2", type="gene", width=0.7, height=0.7];
"gene_ATG7" [label="ATG7", type="gene", width=0.7, height=0.7];
"gene_ATP6V1A" [label="ATP6V1A", type="gene", width=0.7, height=0.7];
"gene_ATP6V1B2" [label="ATP6V1B2", type="gene", width=0.7, height=0.7];
"gene_BRI3" [label="BRI3", type="gene", width=0.7, height=0.7];
"gene_BTK" [label="BTK", type="gene", width=0.7, height=0.7];
"gene_CAP1" [label="CAP1", type="gene", width=0.7, height=0.7];
"gene_CAPZB" [label="CAPZB", type="gene", width=0.7, height=0.7];
"gene_CD300A" [label="CD300A", type="gene", width=0.7, height=0.7];
"gene_CD4" [label="CD4", type="gene", width=0.7, height=0.7];
"gene_CD53" [label="CD53", type="gene", width=0.7, height=0.7];
"gene_CD58" [label="CD58", type="gene", width=0.7, height=0.7];
"gene_CD93" [label="CD93", type="gene", width=0.7, height=0.7];
"gene_CFD" [label="CFD", type="gene", width=0.7, height=0.7];
"gene_CPPED1" [label="CPPED1", type="gene", width=0.7, height=0.7];
"gene_CSF1R" [label="CSF1R", type="gene", width=0.7, height=0.7];
"gene_CSTB" [label="CSTB", type="gene", width=0.7, height=0.7];
"gene_CYFIP1" [label="CYFIP1", type="gene", width=0.7, height=0.7];
"gene_DBT" [label="DBT", type="gene", width=0.7, height=0.7];
"gene_ENTPD1" [label="ENTPD1", type="gene", width=0.7, height=0.7];
"gene_FPR2" [label="FPR2", type="gene", width=0.7, height=0.7];
"gene_GRN" [label="GRN", type="gene", width=0.7, height=0.7];
"gene_HSPA6" [label="HSPA6", type="gene", width=0.7, height=0.7];
"gene_ITGAX" [label="ITGAX", type="gene", width=0.7, height=0.7];
"gene_JAML" [label="JAML", type="gene", width=0.7, height=0.7];
"gene_JUP" [label="JUP", type="gene", width=0.7, height=0.7];
"gene_KDELR1" [label="KDELR1", type="gene", width=0.7, height=0.7];
"gene_KDELR2" [label="KDELR2", type="gene", width=0.7, height=0.7];
"gene_KIR2DL2" [label="KIR2DL2", type="gene", width=0.7, height=0.7];
"gene_KIR3DL1" [label="KIR3DL1", type="gene", width=0.7, height=0.7];
"gene_KIR3DL2" [label="KIR3DL2", type="gene", width=0.7, height=0.7];
"gene_LAIR1" [label="LAIR1", type="gene", width=0.7, height=0.7];
"gene_LILRA1" [label="LILRA1", type="gene", width=0.7, height=0.7];
"gene_LILRA2" [label="LILRA2", type="gene", width=0.7, height=0.7];
"gene_LILRA3" [label="LILRA3", type="gene", width=0.7, height=0.7];
"gene_LILRA6" [label="LILRA6", type="gene", width=0.7, height=0.7];
"gene_LILRB1" [label="LILRB1", type="gene", width=0.7, height=0.7];
"gene_LILRB2" [label="LILRB2", type="gene", width=0.7, height=0.7];
"gene_LILRB3" [label="LILRB3", type="gene", width=0.7, height=0.7];
"gene_MAPKAPK3" [label="MAPKAPK3", type="gene", width=0.7, height=0.7];
"gene_MNDA" [label="MNDA", type="gene", width=0.7, height=0.7];
"gene_MOGS" [label="MOGS", type="gene", width=0.7, height=0.7];
"gene_MOSPD2" [label="MOSPD2", type="gene", width=0.7, height=0.7];
"gene_NOD2" [label="NOD2", type="gene", width=0.7, height=0.7];
"gene_ORMDL2" [label="ORMDL2", type="gene", width=0.7, height=0.7];
"gene_OS9" [label="OS9", type="gene", width=0.7, height=0.7];
"gene_OSBPL11" [label="OSBPL11", type="gene", width=0.7, height=0.7];
"gene_PADI2" [label="PADI2", type="gene", width=0.7, height=0.7];
"gene_PECAM1" [label="PECAM1", type="gene", width=0.7, height=0.7];
"gene_PILRA" [label="PILRA", type="gene", width=0.7, height=0.7];
"gene_PPM1L" [label="PPM1L", type="gene", width=0.7, height=0.7];
"gene_PTAFR" [label="PTAFR", type="gene", width=0.7, height=0.7];
"gene_PTPN6" [label="PTPN6", type="gene", width=0.7, height=0.7];
"gene_PYCARD" [label="PYCARD", type="gene", width=0.7, height=0.7];
"gene_QPCT" [label="QPCT", type="gene", width=0.7, height=0.7];
"gene_RAB10" [label="RAB10", type="gene", width=0.7, height=0.7];
"gene_RAB1B" [label="RAB1B", type="gene", width=0.7, height=0.7];
"gene_RAB27A" [label="RAB27A", type="gene", width=0.7, height=0.7];
"gene_RAB4B" [label="RAB4B", type="gene", width=0.7, height=0.7];
"gene_RAB7A" [label="RAB7A", type="gene", width=0.7, height=0.7];
"gene_RIGI" [label="RIGI", type="gene", width=0.7, height=0.7];
"gene_RNF135" [label="RNF135", type="gene", width=0.7, height=0.7];
"gene_RPS6KA1" [label="RPS6KA1", type="gene", width=0.7, height=0.7];
"gene_S100A11" [label="S100A11", type="gene", width=0.7, height=0.7];
"gene_SEC23IP" [label="SEC23IP", type="gene", width=0.7, height=0.7];
"gene_SERPINB1" [label="SERPINB1", type="gene", width=0.7, height=0.7];
"gene_SIGLEC10" [label="SIGLEC10", type="gene", width=0.7, height=0.7];
"gene_SIGLEC12" [label="SIGLEC12", type="gene", width=0.7, height=0.7];
"gene_SIGLEC5" [label="SIGLEC5", type="gene", width=0.7, height=0.7];
"gene_SIGLEC7" [label="SIGLEC7", type="gene", width=0.7, height=0.7];
"gene_SIRPA" [label="SIRPA", type="gene", width=0.7, height=0.7];
"gene_SIRPB1" [label="SIRPB1", type="gene", width=0.7, height=0.7];
"gene_SLAMF7" [label="SLAMF7", type="gene", width=0.7, height=0.7];
"gene_SPTLC2" [label="SPTLC2", type="gene", width=0.7, height=0.7];
"gene_SPTSSA" [label="SPTSSA", type="gene", width=0.7, height=0.7];
"gene_ST8SIA4" [label="ST8SIA4", type="gene", width=0.7, height=0.7];
"gene_STX3" [label="STX3", type="gene", width=0.7, height=0.7];
"gene_TMEM179B" [label="TMEM179B", type="gene", width=0.7, height=0.7];
```

```
"gene_VRK3" [label="VRK3", type="gene", width=0.7, height=0.7];
"term_10_REACTOME_RHOH_GTPASE_CYCLE" [label="REACTOME: RHOH GTPASE CYCLE", style=filled, fillcolor="#BB8FCE", fontcolor="white", type="term", width=2, height=2, library="C2: Reactome Pathways"];
"term_1_REACTOME_NEUTROPHIL_DEGRANULATION" [label="REACTOME: NEUTROPHIL DEGRANULATION", style=filled, fillcolor="#BB8FCE", fontcolor="white", type="term", width=2, height=2, library="C2: Reactome Pathways"];
"term_2_REACTOME_ROS_AND_RNS_PRODUCTION_IN_PHAGOCYTES" [label="REACTOME: ROS AND RNS PRODUCTION IN PHAGOCYTES", style=filled, fillcolor="#BB8FCE", fontcolor="white", type="term", width=2, height=2, library="C2: Reactome Pathways"];
"term_3_REACTOME_OTHER_INTERLEUKIN_SIGNALING" [label="REACTOME: OTHER INTERLEUKIN SIGNALING", style=filled, fillcolor="#BB8FCE", fontcolor="white", type="term", width=2, height=2, library="C2: Reactome Pathways"];
"term_4_REACTOME_INTERLEUKIN_17_SIGNALING" [label="REACTOME: INTERLEUKIN 17 SIGNALING", style=filled, fillcolor="#BB8FCE", fontcolor="white", type="term", width=2, height=2, library="C2: Reactome Pathways"];
"term_5_REACTOME_IMMUNOREGULATORY_INTERACTIONS_BETWEEN_A_LYMPHOID_AND_A_NON_LYMPHOID_CELL" [label="REACTOME: IMMUNOREGULATORY INTERACTIONS BETWEEN A LYMPHOID AND A NON LYMPHOID CELL", style=filled, fillcolor="#BB8FCE", fontcolor="white", type="term", width=2, height=2, library="C2: Reactome Pathways"];
"term_6_REACTOME_LEISHMANIA_INFECTION" [label="REACTOME: LEISHMANIA INFECTION", style=filled, fillcolor="#BB8FCE", fontcolor="white", type="term", width=2, height=2, library="C2: Reactome Pathways"];
"term_7_REACTOME_NF_KB_ACTIVATION_THROUGH_FADD_RIP_1_PATHWAY_MEDIATED_BY_CASPASE_8_AND_10" [label="REACTOME: NF KB ACTIVATION THROUGH FADD RIP 1 PATHWAY MEDIATED BY CASPASE 8 AND 10", style=filled, fillcolor="#BB8FCE", fontcolor="white", type="term", width=2, height=2, library="C2: Reactome Pathways"];
"term_8_REACTOME_ASPARAGINE_N_LINKED_GLYCOSYLATION" [label="REACTOME: ASPARAGINE N LINKED GLYCOSYLATION", style=filled, fillcolor="#BB8FCE", fontcolor="white", type="term", width=2, height=2, library="C2: Reactome Pathways"];
"term_9_REACTOME_SPHINGOLIPID_METABOLISM" [label="REACTOME: SPHINGOLIPID METABOLISM", style=filled, fillcolor="#BB8FCE", fontcolor="white", type="term", width=2, height=2, library="C2: Reactome Pathways"];
"gene_ACLY" -- "term_1_REACTOME_NEUTROPHIL_DEGRANULATION" [color="#BB8FCE"];
"gene_ACTR3" -- "term_6_REACTOME_LEISHMANIA_INFECTION" [color="#BB8FCE"];
"gene_ADA2" -- "term_1_REACTOME_NEUTROPHIL_DEGRANULATION" [color="#BB8FCE"];
"gene_ADCY7" -- "term_6_REACTOME_LEISHMANIA_INFECTION" [color="#BB8FCE"];
"gene_AHCYL1" -- "term_6_REACTOME_LEISHMANIA_INFECTION" [color="#BB8FCE"];
"gene_ANPEP" -- "term_1_REACTOME_NEUTROPHIL_DEGRANULATION" [color="#BB8FCE"];
"gene_APAF1" -- "term_1_REACTOME_NEUTROPHIL_DEGRANULATION" [color="#BB8FCE"];
"gene_APEH" -- "term_1_REACTOME_NEUTROPHIL_DEGRANULATION" [color="#BB8FCE"];
"gene_ARF3" -- "term_8_REACTOME_ASPARAGINE_N_LINKED_GLYCOSYLATION" [color="#BB8FCE"];
"gene_ARHGAP9" -- "term_1_REACTOME_NEUTROPHIL_DEGRANULATION" [color="#BB8FCE"];
"gene_ARPC1B" -- "term_6_REACTOME_LEISHMANIA_INFECTION" [color="#BB8FCE"];
"gene_ARSD" -- "term_9_REACTOME_SPHINGOLIPID_METABOLISM" [color="#BB8FCE"];
"gene_ASGR2" -- "term_8_REACTOME_ASPARAGINE_N_LINKED_GLYCOSYLATION" [color="#BB8FCE"];
"gene_ATG7" -- "term_1_REACTOME_NEUTROPHIL_DEGRANULATION" [color="#BB8FCE"];
"gene_ATP6V0A1" -- "term_1_REACTOME_NEUTROPHIL_DEGRANULATION" [color="#BB8FCE"];
"gene_ATP6V0D1" -- "term_2_REACTOME_ROS_AND_RNS_PRODUCTION_IN_PHAGOCYTES" [color="#BB8FCE"];
"gene_ATP6V1A" -- "term_2_REACTOME_ROS_AND_RNS_PRODUCTION_IN_PHAGOCYTES" [color="#BB8FCE"];
"gene_ATP6V1B2" -- "term_2_REACTOME_ROS_AND_RNS_PRODUCTION_IN_PHAGOCYTES" [color="#BB8FCE"];
"gene_BRI3" -- "term_1_REACTOME_NEUTROPHIL_DEGRANULATION" [color="#BB8FCE"];
"gene_BTK" -- "term_6_REACTOME_LEISHMANIA_INFECTION" [color="#BB8FCE"];
"gene_CAP1" -- "term_1_REACTOME_NEUTROPHIL_DEGRANULATION" [color="#BB8FCE"];
"gene_CAPZB" -- "term_8_REACTOME_ASPARAGINE_N_LINKED_GLYCOSYLATION" [color="#BB8FCE"];
"gene_CASP1" -- "term_6_REACTOME_LEISHMANIA_INFECTION" [color="#BB8FCE"];
"gene_CASP10" -- "term_7_REACTOME_NF_KB_ACTIVATION_THROUGH_FADD_RIP_1_PATHWAY_MEDIATED_BY_CASPASE_8_AND_10" [color="#BB8FCE"];
"gene_CD300A" -- "term_1_REACTOME_NEUTROPHIL_DEGRANULATION" [color="#BB8FCE"];
"gene_CD36" -- "term_1_REACTOME_NEUTROPHIL_DEGRANULATION" [color="#BB8FCE"];
"gene_CD4" -- "term_3_REACTOME_OTHER_INTERLEUKIN_SIGNALING" [color="#BB8FCE"];
"gene_CD53" -- "term_1_REACTOME_NEUTROPHIL_DEGRANULATION" [color="#BB8FCE"];
"gene_CD58" -- "term_1_REACTOME_NEUTROPHIL_DEGRANULATION" [color="#BB8FCE"];
"gene_CD93" -- "term_1_REACTOME_NEUTROPHIL_DEGRANULATION" [color="#BB8FCE"];
"gene_CFD" -- "term_1_REACTOME_NEUTROPHIL_DEGRANULATION" [color="#BB8FCE"];
"gene_CPPED1" -- "term_1_REACTOME_NEUTROPHIL_DEGRANULATION" [color="#BB8FCE"];
"gene_CR1" -- "term_1_REACTOME_NEUTROPHIL_DEGRANULATION" [color="#BB8FCE"];
"gene_CSF1R" -- "term_3_REACTOME_OTHER_INTERLEUKIN_SIGNALING" [color="#BB8FCE"];
"gene_CSF3R" -- "term_3_REACTOME_OTHER_INTERLEUKIN_SIGNALING" [color="#BB8FCE"];
"gene_CSK" -- "term_10_REACTOME_RHOH_GTPASE_CYCLE" [color="#BB8FCE"];
"gene_CSTB" -- "term_1_REACTOME_NEUTROPHIL_DEGRANULATION" [color="#BB8FCE"];
"gene_CTSD" -- "term_1_REACTOME_NEUTROPHIL_DEGRANULATION" [color="#BB8FCE"];
"gene_CYFIP1" -- "term_1_REACTOME_NEUTROPHIL_DEGRANULATION" [color="#BB8FCE"];
"gene_DBT" -- "term_10_REACTOME_RHOH_GTPASE_CYCLE" [color="#BB8FCE"];
"gene_ENTPD1" -- "term_6_REACTOME_LEISHMANIA_INFECTION" [color="#BB8FCE"];
"gene_FGL2" -- "term_1_REACTOME_NEUTROPHIL_DEGRANULATION" [color="#BB8FCE"];
"gene_FPGT" -- "term_8_REACTOME_ASPARAGINE_N_LINKED_GLYCOSYLATION" [color="#BB8FCE"];
"gene_FPR2" -- "term_1_REACTOME_NEUTROPHIL_DEGRANULATION" [color="#BB8FCE"];
"gene_GAA" -- "term_1_REACTOME_NEUTROPHIL_DEGRANULATION" [color="#BB8FCE"];
"gene_GBA1" -- "term_9_REACTOME_SPHINGOLIPID_METABOLISM" [color="#BB8FCE"];
"gene_GCA" -- "term_1_REACTOME_NEUTROPHIL_DEGRANULATION" [color="#BB8FCE"];
"gene_GNS" -- "term_1_REACTOME_NEUTROPHIL_DEGRANULATION" [color="#BB8FCE"];
"gene_GRN" -- "term_1_REACTOME_NEUTROPHIL_DEGRANULATION" [color="#BB8FCE"];
"gene_GSN" -- "term_1_REACTOME_NEUTROPHIL_DEGRANULATION" [color="#BB8FCE"];
"gene_HCK" -- "term_6_REACTOME_LEISHMANIA_INFECTION" [color="#BB8FCE"];
"gene_HK1" -- "term_8_REACTOME_ASPARAGINE_N_LINKED_GLYCOSYLATION" [color="#BB8FCE"];
"gene_HK3" -- "term_1_REACTOME_NEUTROPHIL_DEGRANULATION" [color="#BB8FCE"];
"gene_HSPA6" -- "term_1_REACTOME_NEUTROPHIL_DEGRANULATION" [color="#BB8FCE"];
"gene_IDH1" -- "term_1_REACTOME_NEUTROPHIL_DEGRANULATION" [color="#BB8FCE"];
"gene_IL10RB" -- "term_3_REACTOME_OTHER_INTERLEUKIN_SIGNALING" [color="#BB8FCE"];
"gene_IL17RA" -- "term_4_REACTOME_INTERLEUKIN_17_SIGNALING" [color="#BB8FCE"];
"gene_IQGAP1" -- "term_1_REACTOME_NEUTROPHIL_DEGRANULATION" [color="#BB8FCE"];
"gene_ITGAX" -- "term_1_REACTOME_NEUTROPHIL_DEGRANULATION" [color="#BB8FCE"];
"gene_JAML" -- "term_5_REACTOME_IMMUNOREGULATORY_INTERACTIONS_BETWEEN_A_LYMPHOID_AND_A_NON_LYMPHOID_CELL" [color="#BB8FCE"];
"gene_JUP" -- "term_1_REACTOME_NEUTROPHIL_DEGRANULATION" [color="#BB8FCE"];
"gene_KDELR1" -- "term_8_REACTOME_ASPARAGINE_N_LINKED_GLYCOSYLATION" [color="#BB8FCE"];
"gene_KDELR2" -- "term_8_REACTOME_ASPARAGINE_N_LINKED_GLYCOSYLATION" [color="#BB8FCE"];
"gene_KIR2DL2" -- "term_5_REACTOME_IMMUNOREGULATORY_INTERACTIONS_BETWEEN_A_LYMPHOID_AND_A_NON_LYMPHOID_CELL" [color="#BB8FCE"];
"gene_KIR3DL1" -- "term_5_REACTOME_IMMUNOREGULATORY_INTERACTIONS_BETWEEN_A_LYMPHOID_AND_A_NON_LYMPHOID_CELL" [color="#BB8FCE"];
"gene_KIR3DL2" -- "term_5_REACTOME_IMMUNOREGULATORY_INTERACTIONS_BETWEEN_A_LYMPHOID_AND_A_NON_LYMPHOID_CELL" [color="#BB8FCE"];
"gene_LAIR1" -- "term_1_REACTOME_NEUTROPHIL_DEGRANULATION" [color="#BB8FCE"];
"gene_LAMP2" -- "term_1_REACTOME_NEUTROPHIL_DEGRANULATION" [color="#BB8FCE"];
"gene_LILRA1" -- "term_5_REACTOME_IMMUNOREGULATORY_INTERACTIONS_BETWEEN_A_LYMPHOID_AND_A_NON_LYMPHOID_CELL" [color="#BB8FCE"];
"gene_LILRA2" -- "term_5_REACTOME_IMMUNOREGULATORY_INTERACTIONS_BETWEEN_A_LYMPHOID_AND_A_NON_LYMPHOID_CELL" [color="#BB8FCE"];
"gene_LILRA3" -- "term_1_REACTOME_NEUTROPHIL_DEGRANULATION" [color="#BB8FCE"];
"gene_LILRA6" -- "term_5_REACTOME_IMMUNOREGULATORY_INTERACTIONS_BETWEEN_A_LYMPHOID_AND_A_NON_LYMPHOID_CELL" [color="#BB8FCE"];
"gene_LILRB1" -- "term_5_REACTOME_IMMUNOREGULATORY_INTERACTIONS_BETWEEN_A_LYMPHOID_AND_A_NON_LYMPHOID_CELL" [color="#BB8FCE"];
"gene_LILRB2" -- "term_1_REACTOME_NEUTROPHIL_DEGRANULATION" [color="#BB8FCE"];
"gene_LILRB3" -- "term_1_REACTOME_NEUTROPHIL_DEGRANULATION" [color="#BB8FCE"];
"gene_M6PR" -- "term_9_REACTOME_SPHINGOLIPID_METABOLISM" [color="#BB8FCE"];
"gene_MAP2K3" -- "term_4_REACTOME_INTERLEUKIN_17_SIGNALING" [color="#BB8FCE"];
"gene_MAP2K4" -- "term_4_REACTOME_INTERLEUKIN_17_SIGNALING" [color="#BB8FCE"];
"gene_MAPK1" -- "term_1_REACTOME_NEUTROPHIL_DEGRANULATION" [color="#BB8FCE"];
"gene_MAPKAPK3" -- "term_4_REACTOME_INTERLEUKIN_17_SIGNALING" [color="#BB8FCE"];
"gene_MNDA" -- "term_1_REACTOME_NEUTROPHIL_DEGRANULATION" [color="#BB8FCE"];
"gene_MOGS" -- "term_8_REACTOME_ASPARAGINE_N_LINKED_GLYCOSYLATION" [color="#BB8FCE"];
"gene_MOSPD2" -- "term_1_REACTOME_NEUTROPHIL_DEGRANULATION" [color="#BB8FCE"];
"gene_NAGK" -- "term_8_REACTOME_ASPARAGINE_N_LINKED_GLYCOSYLATION" [color="#BB8FCE"];
"gene_NCF1" -- "term_2_REACTOME_ROS_AND_RNS_PRODUCTION_IN_PHAGOCYTES" [color="#BB8FCE"];
"gene_NCF2" -- "term_2_REACTOME_ROS_AND_RNS_PRODUCTION_IN_PHAGOCYTES" [color="#BB8FCE"];
"gene_NCF4" -- "term_2_REACTOME_ROS_AND_RNS_PRODUCTION_IN_PHAGOCYTES" [color="#BB8FCE"];
"gene_NOD2" -- "term_4_REACTOME_INTERLEUKIN_17_SIGNALING" [color="#BB8FCE"];
"gene_NPL" -- "term_8_REACTOME_ASPARAGINE_N_LINKED_GLYCOSYLATION" [color="#BB8FCE"];
"gene_ORMDL2" -- "term_9_REACTOME_SPHINGOLIPID_METABOLISM" [color="#BB8FCE"];
"gene_OS9" -- "term_8_REACTOME_ASPARAGINE_N_LINKED_GLYCOSYLATION" [color="#BB8FCE"];
"gene_OSBPL11" -- "term_10_REACTOME_RHOH_GTPASE_CYCLE" [color="#BB8FCE"];
"gene_PADI2" -- "term_1_REACTOME_NEUTROPHIL_DEGRANULATION" [color="#BB8FCE"];
"gene_PAK1" -- "term_10_REACTOME_RHOH_GTPASE_CYCLE" [color="#BB8FCE"];
"gene_PECAM1" -- "term_1_REACTOME_NEUTROPHIL_DEGRANULATION" [color="#BB8FCE"];
"gene_PILRA" -- "term_5_REACTOME_IMMUNOREGULATORY_INTERACTIONS_BETWEEN_A_LYMPHOID_AND_A_NON_LYMPHOID_CELL" [color="#BB8FCE"];
"gene_PPM1L" -- "term_9_REACTOME_SPHINGOLIPID_METABOLISM" [color="#BB8FCE"];
"gene_PPP2CB" -- "term_4_REACTOME_INTERLEUKIN_17_SIGNALING" [color="#BB8FCE"];
"gene_PRCP" -- "term_1_REACTOME_NEUTROPHIL_DEGRANULATION" [color="#BB8FCE"];
"gene_PRKACA" -- "term_6_REACTOME_LEISHMANIA_INFECTION" [color="#BB8FCE"];
"gene_PRKAR1A" -- "term_6_REACTOME_LEISHMANIA_INFECTION" [color="#BB8FCE"];
"gene_PRKCD" -- "term_1_REACTOME_NEUTROPHIL_DEGRANULATION" [color="#BB8FCE"];
"gene_PSEN1" -- "term_1_REACTOME_NEUTROPHIL_DEGRANULATION" [color="#BB8FCE"];
"gene_PTAFR" -- "term_1_REACTOME_NEUTROPHIL_DEGRANULATION" [color="#BB8FCE"];
"gene_PTPN6" -- "term_1_REACTOME_NEUTROPHIL_DEGRANULATION" [color="#BB8FCE"];
"gene_PTPRJ" -- "term_1_REACTOME_NEUTROPHIL_DEGRANULATION" [color="#BB8FCE"];
"gene_PYCARD" -- "term_1_REACTOME_NEUTROPHIL_DEGRANULATION" [color="#BB8FCE"];
"gene_QPCT" -- "term_1_REACTOME_NEUTROPHIL_DEGRANULATION" [color="#BB8FCE"];
"gene_RAB10" -- "term_1_REACTOME_NEUTROPHIL_DEGRANULATION" [color="#BB8FCE"];
"gene_RAB1B" -- "term_8_REACTOME_ASPARAGINE_N_LINKED_GLYCOSYLATION" [color="#BB8FCE"];
"gene_RAB27A" -- "term_1_REACTOME_NEUTROPHIL_DEGRANULATION" [color="#BB8FCE"];
"gene_RAB4B" -- "term_1_REACTOME_NEUTROPHIL_DEGRANULATION" [color="#BB8FCE"];
"gene_RAB7A" -- "term_1_REACTOME_NEUTROPHIL_DEGRANULATION" [color="#BB8FCE"];
"gene_RIGI" -- "term_7_REACTOME_NF_KB_ACTIVATION_THROUGH_FADD_RIP_1_PATHWAY_MEDIATED_BY_CASPASE_8_AND_10" [color="#BB8FCE"];
"gene_RNF135" -- "term_7_REACTOME_NF_KB_ACTIVATION_THROUGH_FADD_RIP_1_PATHWAY_MEDIATED_BY_CASPASE_8_AND_10" [color="#BB8FCE"];
"gene_RPS6KA1" -- "term_4_REACTOME_INTERLEUKIN_17_SIGNALING" [color="#BB8FCE"];
"gene_S100A11" -- "term_1_REACTOME_NEUTROPHIL_DEGRANULATION" [color="#BB8FCE"];
"gene_SEC23IP" -- "term_8_REACTOME_ASPARAGINE_N_LINKED_GLYCOSYLATION" [color="#BB8FCE"];
"gene_SERPINA1" -- "term_1_REACTOME_NEUTROPHIL_DEGRANULATION" [color="#BB8FCE"];
"gene_SERPINB1" -- "term_1_REACTOME_NEUTROPHIL_DEGRANULATION" [color="#BB8FCE"];
"gene_SIGLEC10" -- "term_5_REACTOME_IMMUNOREGULATORY_INTERACTIONS_BETWEEN_A_LYMPHOID_AND_A_NON_LYMPHOID_CELL" [color="#BB8FCE"];
"gene_SIGLEC12" -- "term_5_REACTOME_IMMUNOREGULATORY_INTERACTIONS_BETWEEN_A_LYMPHOID_AND_A_NON_LYMPHOID_CELL" [color="#BB8FCE"];
"gene_SIGLEC5" -- "term_1_REACTOME_NEUTROPHIL_DEGRANULATION" [color="#BB8FCE"];
"gene_SIGLEC7" -- "term_5_REACTOME_IMMUNOREGULATORY_INTERACTIONS_BETWEEN_A_LYMPHOID_AND_A_NON_LYMPHOID_CELL" [color="#BB8FCE"];
"gene_SIRPA" -- "term_1_REACTOME_NEUTROPHIL_DEGRANULATION" [color="#BB8FCE"];
"gene_SIRPB1" -- "term_1_REACTOME_NEUTROPHIL_DEGRANULATION" [color="#BB8FCE"];
"gene_SLAMF7" -- "term_5_REACTOME_IMMUNOREGULATORY_INTERACTIONS_BETWEEN_A_LYMPHOID_AND_A_NON_LYMPHOID_CELL" [color="#BB8FCE"];
"gene_SLC11A1" -- "term_1_REACTOME_NEUTROPHIL_DEGRANULATION" [color="#BB8FCE"];
"gene_SPTLC2" -- "term_9_REACTOME_SPHINGOLIPID_METABOLISM" [color="#BB8FCE"];
"gene_SPTSSA" -- "term_9_REACTOME_SPHINGOLIPID_METABOLISM" [color="#BB8FCE"];
"gene_ST8SIA4" -- "term_8_REACTOME_ASPARAGINE_N_LINKED_GLYCOSYLATION" [color="#BB8FCE"];
"gene_STX3" -- "term_3_REACTOME_OTHER_INTERLEUKIN_SIGNALING" [color="#BB8FCE"];
"gene_SYK" -- "term_6_REACTOME_LEISHMANIA_INFECTION" [color="#BB8FCE"];
"gene_TCIRG1" -- "term_1_REACTOME_NEUTROPHIL_DEGRANULATION" [color="#BB8FCE"];
"gene_TIMP2" -- "term_1_REACTOME_NEUTROPHIL_DEGRANULATION" [color="#BB8FCE"];
"gene_TMEM179B" -- "term_1_REACTOME_NEUTROPHIL_DEGRANULATION" [color="#BB8FCE"];
"gene_VAV1" -- "term_6_REACTOME_LEISHMANIA_INFECTION" [color="#BB8FCE"];
"gene_VRK3" -- "term_4_REACTOME_INTERLEUKIN_17_SIGNALING" [color="#BB8FCE"];
"gene_APBB1IP" [label="APBB1IP", type="gene", width=0.7, height=0.7];
"gene_CAPG" [label="CAPG", type="gene", width=0.7, height=0.7];
"gene_CARD8" [label="CARD8", type="gene", width=0.7, height=0.7];
"gene_DAGLB" [label="DAGLB", type="gene", width=0.7, height=0.7];
"gene_DAXX" [label="DAXX", type="gene", width=0.7, height=0.7];
"gene_FKBP15" [label="FKBP15", type="gene", width=0.7, height=0.7];
"gene_GAPT" [label="GAPT", type="gene", width=0.7, height=0.7];
"gene_KCNE3" [label="KCNE3", type="gene", width=0.7, height=0.7];
"gene_LYL1" [label="LYL1", type="gene", width=0.7, height=0.7];
"gene_NAIP" [label="NAIP", type="gene", width=0.7, height=0.7];
"gene_NLRP12" [label="NLRP12", type="gene", width=0.7, height=0.7];
"gene_OGDH" [label="OGDH", type="gene", width=0.7, height=0.7];
"gene_PPARGC1A" [label="PPARGC1A", type="gene", width=0.7, height=0.7];
"gene_PPP1R18" [label="PPP1R18", type="gene", width=0.7, height=0.7];
"gene_RBM47" [label="RBM47", type="gene", width=0.7, height=0.7];
"gene_SMC1A" [label="SMC1A", type="gene", width=0.7, height=0.7];
"gene_TFEB" [label="TFEB", type="gene", width=0.7, height=0.7];
"gene_VMP1" [label="VMP1", type="gene", width=0.7, height=0.7];
"term_1_WP_TYROBP_CAUSAL_NETWORK_IN_MICROGLIA" [label="WP: TYROBP CAUSAL NETWORK IN MICROGLIA", style=filled, fillcolor="#31688e", fontcolor="white", type="term", width=2, height=2, library="Unknown Library"];
"term_2_WP_MICROGLIA_PATHOGEN_PHAGOCYTOSIS_PATHWAY" [label="WP: MICROGLIA PATHOGEN PHAGOCYTOSIS PATHWAY", style=filled, fillcolor="#31688e", fontcolor="white", type="term", width=2, height=2, library="Unknown Library"];
"term_3_WP_FAS_LIGAND_PATHWAY_AND_STRESS_INDUCTION_OF_HEAT_SHOCK_PROTEINS" [label="WP: FAS LIGAND PATHWAY AND STRESS INDUCTION OF HEAT SHOCK PROTEINS", style=filled, fillcolor="#31688e", fontcolor="white", type="term", width=2, height=2, library="Unknown Library"];
"term_4_WP_CANNABINOID_RECEPTOR_SIGNALING" [label="WP: CANNABINOID RECEPTOR SIGNALING", style=filled, fillcolor="#31688e", fontcolor="white", type="term", width=2, height=2, library="Unknown Library"];
"term_5_WP_NUCLEOTIDEBINDING_OLIGOMERIZATION_DOMAIN_NOD_PATHWAY" [label="WP: NUCLEOTIDEBINDING OLIGOMERIZATION DOMAIN NOD PATHWAY", style=filled, fillcolor="#31688e", fontcolor="white", type="term", width=2, height=2, library="Unknown Library"];
"term_6_WP_CONSTITUTIVE_ANDROSTANE_RECEPTOR_PATHWAY" [label="WP: CONSTITUTIVE ANDROSTANE RECEPTOR PATHWAY", style=filled, fillcolor="#31688e", fontcolor="white", type="term", width=2, height=2, library="Unknown Library"];
"term_7_WP_TCA_CYCLE_AND_DEFICIENCY_OF_PYRUVATE_DEHYDROGENASE_COMPLEX_PDHC" [label="WP: TCA CYCLE AND DEFICIENCY OF PYRUVATE DEHYDROGENASE COMPLEX PDHC", style=filled, fillcolor="#31688e", fontcolor="white", type="term", width=2, height=2, library="Unknown Library"];
"term_8_WP_AUTOPHAGY_IN_PANCREATIC_DUCTAL_ADENOCARCINOMA" [label="WP: AUTOPHAGY IN PANCREATIC DUCTAL ADENOCARCINOMA", style=filled, fillcolor="#31688e", fontcolor="white", type="term", width=2, height=2, library="Unknown Library"];
"gene_ACLY" -- "term_7_WP_TCA_CYCLE_AND_DEFICIENCY_OF_PYRUVATE_DEHYDROGENASE_COMPLEX_PDHC" [color="#31688e"];
"gene_ADCY7" -- "term_4_WP_CANNABINOID_RECEPTOR_SIGNALING" [color="#31688e"];
"gene_APAF1" -- "term_3_WP_FAS_LIGAND_PATHWAY_AND_STRESS_INDUCTION_OF_HEAT_SHOCK_PROTEINS" [color="#31688e"];
"gene_APBB1IP" -- "term_1_WP_TYROBP_CAUSAL_NETWORK_IN_MICROGLIA" [color="#31688e"];
"gene_ARPC1B" -- "term_2_WP_MICROGLIA_PATHOGEN_PHAGOCYTOSIS_PATHWAY" [color="#31688e"];
"gene_ATG7" -- "term_8_WP_AUTOPHAGY_IN_PANCREATIC_DUCTAL_ADENOCARCINOMA" [color="#31688e"];
```

```
"gene_CAPG" -- "term_1_WP_TYROBP_CAUSAL_NETWORK_IN_MICROGLIA" [color="#31688e"];
"gene_CARD8" -- "term_5_WP_NUCLEOTIDEBINDING_OLIGOMERIZATION_DOMAIN_NOD_PATHWAY" [color="#31688e"];
"gene_CASP1" -- "term_5_WP_NUCLEOTIDEBINDING_OLIGOMERIZATION_DOMAIN_NOD_PATHWAY" [color="#31688e"];
"gene_CASP10" -- "term_3_WP_FAS_LIGAND_PATHWAY_AND_STRESS_INDUCTION_OF_HEAT_SHOCK_PROTEINS" [color="#31688e"];
"gene_CD4" -- "term_1_WP_TYROBP_CAUSAL_NETWORK_IN_MICROGLIA" [color="#31688e"];
"gene_DAGLB" -- "term_4_WP_CANNABINOID_RECEPTOR_SIGNALING" [color="#31688e"];
"gene_DAXX" -- "term_3_WP_FAS_LIGAND_PATHWAY_AND_STRESS_INDUCTION_OF_HEAT_SHOCK_PROTEINS" [color="#31688e"];
"gene_DFFA" -- "term_3_WP_FAS_LIGAND_PATHWAY_AND_STRESS_INDUCTION_OF_HEAT_SHOCK_PROTEINS" [color="#31688e"];
"gene_FKBP15" -- "term_1_WP_TYROBP_CAUSAL_NETWORK_IN_MICROGLIA" [color="#31688e"];
"gene_GAPT" -- "term_1_WP_TYROBP_CAUSAL_NETWORK_IN_MICROGLIA" [color="#31688e"];
"gene_HCK" -- "term_2_WP_MICROGLIA_PATHOGEN_PHAGOCYTOSIS_PATHWAY" [color="#31688e"];
"gene_IDH1" -- "term_7_WP_TCA_CYCLE_AND_DEFICIENCY_OF_PYRUVATE_DEHYDROGENASE_COMPLEX_PDHC" [color="#31688e"];
"gene_ITGAX" -- "term_1_WP_TYROBP_CAUSAL_NETWORK_IN_MICROGLIA" [color="#31688e"];
"gene_KCNE3" -- "term_1_WP_TYROBP_CAUSAL_NETWORK_IN_MICROGLIA" [color="#31688e"];
"gene_LYL1" -- "term_1_WP_TYROBP_CAUSAL_NETWORK_IN_MICROGLIA" [color="#31688e"];
"gene_MAP2K4" -- "term_3_WP_FAS_LIGAND_PATHWAY_AND_STRESS_INDUCTION_OF_HEAT_SHOCK_PROTEINS" [color="#31688e"];
"gene_MAPK1" -- "term_4_WP_CANNABINOID_RECEPTOR_SIGNALING" [color="#31688e"];
"gene_MAPKAPK3" -- "term_3_WP_FAS_LIGAND_PATHWAY_AND_STRESS_INDUCTION_OF_HEAT_SHOCK_PROTEINS" [color="#31688e"];
"gene_NAIP" -- "term_5_WP_NUCLEOTIDEBINDING_OLIGOMERIZATION_DOMAIN_NOD_PATHWAY" [color="#31688e"];
"gene_NCF1" -- "term_2_WP_MICROGLIA_PATHOGEN_PHAGOCYTOSIS_PATHWAY" [color="#31688e"];
"gene_NCF2" -- "term_1_WP_TYROBP_CAUSAL_NETWORK_IN_MICROGLIA" [color="#31688e"];
"gene_NCF4" -- "term_2_WP_MICROGLIA_PATHOGEN_PHAGOCYTOSIS_PATHWAY" [color="#31688e"];
"gene_NLRP12" -- "term_5_WP_NUCLEOTIDEBINDING_OLIGOMERIZATION_DOMAIN_NOD_PATHWAY" [color="#31688e"];
"gene_NOD2" -- "term_5_WP_NUCLEOTIDEBINDING_OLIGOMERIZATION_DOMAIN_NOD_PATHWAY" [color="#31688e"];
"gene_OGDH" -- "term_7_WP_TCA_CYCLE_AND_DEFICIENCY_OF_PYRUVATE_DEHYDROGENASE_COMPLEX_PDHC" [color="#31688e"];
"gene_PAK1" -- "term_3_WP_FAS_LIGAND_PATHWAY_AND_STRESS_INDUCTION_OF_HEAT_SHOCK_PROTEINS" [color="#31688e"];
"gene_PPARGC1A" -- "term_6_WP_CONSTITUTIVE_ANDROSTANE_RECEPTOR_PATHWAY" [color="#31688e"];
"gene_PPP1R18" -- "term_1_WP_TYROBP_CAUSAL_NETWORK_IN_MICROGLIA" [color="#31688e"];
"gene_PRKACA" -- "term_4_WP_CANNABINOID_RECEPTOR_SIGNALING" [color="#31688e"];
"gene_PRKAR1A" -- "term_4_WP_CANNABINOID_RECEPTOR_SIGNALING" [color="#31688e"];
"gene_PTPN6" -- "term_2_WP_MICROGLIA_PATHOGEN_PHAGOCYTOSIS_PATHWAY" [color="#31688e"];
"gene_PYCARD" -- "term_1_WP_TYROBP_CAUSAL_NETWORK_IN_MICROGLIA" [color="#31688e"];
"gene_RBM47" -- "term_1_WP_TYROBP_CAUSAL_NETWORK_IN_MICROGLIA" [color="#31688e"];
"gene_RPS6KA1" -- "term_1_WP_TYROBP_CAUSAL_NETWORK_IN_MICROGLIA" [color="#31688e"];
"gene_RXRA" -- "term_6_WP_CONSTITUTIVE_ANDROSTANE_RECEPTOR_PATHWAY" [color="#31688e"];
"gene_SIGLEC7" -- "term_2_WP_MICROGLIA_PATHOGEN_PHAGOCYTOSIS_PATHWAY" [color="#31688e"];
"gene_SMC1A" -- "term_6_WP_CONSTITUTIVE_ANDROSTANE_RECEPTOR_PATHWAY" [color="#31688e"];
"gene_SULT1A1" -- "term_6_WP_CONSTITUTIVE_ANDROSTANE_RECEPTOR_PATHWAY" [color="#31688e"];
"gene_SYK" -- "term_2_WP_MICROGLIA_PATHOGEN_PHAGOCYTOSIS_PATHWAY" [color="#31688e"];
"gene_TCIRG1" -- "term_1_WP_TYROBP_CAUSAL_NETWORK_IN_MICROGLIA" [color="#31688e"];
"gene_TFEB" -- "term_8_WP_AUTOPHAGY_IN_PANCREATIC_DUCTAL_ADENOCARCINOMA" [color="#31688e"];
"gene_VAV1" -- "term_2_WP_MICROGLIA_PATHOGEN_PHAGOCYTOSIS_PATHWAY" [color="#31688e"];
"gene_VMP1" -- "term_8_WP_AUTOPHAGY_IN_PANCREATIC_DUCTAL_ADENOCARCINOMA" [color="#31688e"];
"gene_ABCA7" [label="ABCA7", type="gene", width=0.7, height=0.7];
"gene_APH1B" [label="APH1B", type="gene", width=0.7, height=0.7];
"gene_CCR5" [label="CCR5", type="gene", width=0.7, height=0.7];
"gene_CIDEB" [label="CIDEB", type="gene", width=0.7, height=0.7];
"gene_CNIH4" [label="CNIH4", type="gene", width=0.7, height=0.7];
"gene_CORO1B" [label="CORO1B", type="gene", width=0.7, height=0.7];
"gene_CX3CR1" [label="CX3CR1", type="gene", width=0.7, height=0.7];
"gene_DGAT2" [label="DGAT2", type="gene", width=0.7, height=0.7];
"gene_DMXL2" [label="DMXL2", type="gene", width=0.7, height=0.7];
"gene_EIF4E2" [label="EIF4E2", type="gene", width=0.7, height=0.7];
"gene_FAR2" [label="FAR2", type="gene", width=0.7, height=0.7];
"gene_GK" [label="GK", type="gene", width=0.7, height=0.7];
"gene_HAL" [label="HAL", type="gene", width=0.7, height=0.7];
"gene_IFITM2" [label="IFITM2", type="gene", width=0.7, height=0.7];
"gene_IFITM3" [label="IFITM3", type="gene", width=0.7, height=0.7];
"gene_ILK" [label="ILK", type="gene", width=0.7, height=0.7];
"gene_JAGN1" [label="JAGN1", type="gene", width=0.7, height=0.7];
"gene_LRRK2" [label="LRRK2", type="gene", width=0.7, height=0.7];
"gene_LYST" [label="LYST", type="gene", width=0.7, height=0.7];
"gene_MCCC2" [label="MCCC2", type="gene", width=0.7, height=0.7];
"gene_MGST2" [label="MGST2", type="gene", width=0.7, height=0.7];
"gene_MIR21" [label="MIR21", type="gene", width=0.7, height=0.7];
"gene_MPEG1" [label="MPEG1", type="gene", width=0.7, height=0.7];
"gene_NAAA" [label="NAAA", type="gene", width=0.7, height=0.7];
"gene_PMVK" [label="PMVK", type="gene", width=0.7, height=0.7];
"gene_PRELID1" [label="PRELID1", type="gene", width=0.7, height=0.7];
"gene_RAB8A" [label="RAB8A", type="gene", width=0.7, height=0.7];
"gene_RTN1" [label="RTN1", type="gene", width=0.7, height=0.7];
"gene_SEC23B" [label="SEC23B", type="gene", width=0.7, height=0.7];
"gene_SLC12A6" [label="SLC12A6", type="gene", width=0.7, height=0.7];
"gene_SNX1" [label="SNX1", type="gene", width=0.7, height=0.7];
"gene_STX6" [label="STX6", type="gene", width=0.7, height=0.7];
"gene_SYT11" [label="SYT11", type="gene", width=0.7, height=0.7];
"gene_TBXAS1" [label="TBXAS1", type="gene", width=0.7, height=0.7];
"gene_VIPAS39" [label="VIPAS39", type="gene", width=0.7, height=0.7];
"gene_WASHC5" [label="WASHC5", type="gene", width=0.7, height=0.7];
"gene_WBP1L" [label="WBP1L", type="gene", width=0.7, height=0.7];
"term_10_GOBP_MONOATOMIC_ANION_HOMEOSTASIS" [label="GOBP: MONOATOMIC ANION HOMEOSTASIS", style=filled, fillcolor="#F1948A", fontcolor="white", type="term", width=2, height=2, library="C5: Gene Ontology: Biological Process"];
"term_1_GOBP_PHAGOCYTOSIS" [label="GOBP: PHAGOCYTOSIS", style=filled, fillcolor="#F1948A", fontcolor="white", type="term", width=2, height=2, library="C5: Gene Ontology: Biological Process"];
"term_2_GOBP_CYTOKINE_MEDIATED_SIGNALING_PATHWAY" [label="GOBP: CYTOKINE MEDIATED SIGNALING PATHWAY", style=filled, fillcolor="#F1948A", fontcolor="white", type="term", width=2, height=2, library="C5: Gene Ontology: Biological Process"];
"term_3_GOBP_VACUOLAR_ACIDIFICATION" [label="GOBP: VACUOLAR ACIDIFICATION", style=filled, fillcolor="#F1948A", fontcolor="white", type="term", width=2, height=2, library="C5: Gene Ontology: Biological Process"];
"term_4_GOBP_AMIDE_METABOLIC_PROCESS" [label="GOBP: AMIDE METABOLIC PROCESS", style=filled, fillcolor="#F1948A", fontcolor="white", type="term", width=2, height=2, library="C5: Gene Ontology: Biological Process"];
"term_5_GOBP_GOLGI_VESICLE_TRANSPORT" [label="GOBP: GOLGI VESICLE TRANSPORT", style=filled, fillcolor="#F1948A", fontcolor="white", type="term", width=2, height=2, library="C5: Gene Ontology: Biological Process"];
"term_6_GOBP_ALCOHOL_CATABOLIC_PROCESS" [label="GOBP: ALCOHOL CATABOLIC PROCESS", style=filled, fillcolor="#F1948A", fontcolor="white", type="term", width=2, height=2, library="C5: Gene Ontology: Biological Process"];
"term_7_GOBP_ANTIBACTERIAL_INNATE_IMMUNE_RESPONSE" [label="GOBP: ANTIBACTERIAL INNATE IMMUNE RESPONSE", style=filled, fillcolor="#F1948A", fontcolor="white", type="term", width=2, height=2, library="C5: Gene Ontology: Biological Process"];
"term_8_GOBP_ARP2_3_COMPLEX_MEDIATED_ACTIN_NUCLEATION" [label="GOBP: ARP2 3 COMPLEX MEDIATED ACTIN NUCLEATION", style=filled, fillcolor="#F1948A", fontcolor="white", type="term", width=2, height=2, library="C5: Gene Ontology: Biological Process"];
"term_9_GOBP_REGULATION_OF_PEPTIDASE_ACTIVITY" [label="GOBP: REGULATION OF PEPTIDASE ACTIVITY", style=filled, fillcolor="#F1948A", fontcolor="white", type="term", width=2, height=2, library="C5: Gene Ontology: Biological Process"];
"gene_ABCA7" -- "term_1_GOBP_PHAGOCYTOSIS" [color="#F1948A"];
"gene_ACLY" -- "term_4_GOBP_AMIDE_METABOLIC_PROCESS" [color="#F1948A"];
"gene_ACTR3" -- "term_8_GOBP_ARP2_3_COMPLEX_MEDIATED_ACTIN_NUCLEATION" [color="#F1948A"];
"gene_ADIPOR2" -- "term_2_GOBP_CYTOKINE_MEDIATED_SIGNALING_PATHWAY" [color="#F1948A"];
"gene_ANPEP" -- "term_4_GOBP_AMIDE_METABOLIC_PROCESS" [color="#F1948A"];
"gene_AP3S2" -- "term_5_GOBP_GOLGI_VESICLE_TRANSPORT" [color="#F1948A"];
"gene_APEH" -- "term_4_GOBP_AMIDE_METABOLIC_PROCESS" [color="#F1948A"];
"gene_APH1B" -- "term_4_GOBP_AMIDE_METABOLIC_PROCESS" [color="#F1948A"];
"gene_ARAP1" -- "term_1_GOBP_PHAGOCYTOSIS" [color="#F1948A"];
"gene_ARF3" -- "term_5_GOBP_GOLGI_VESICLE_TRANSPORT" [color="#F1948A"];
"gene_ARPC1B" -- "term_8_GOBP_ARP2_3_COMPLEX_MEDIATED_ACTIN_NUCLEATION" [color="#F1948A"];
"gene_ATP6AP1" -- "term_3_GOBP_VACUOLAR_ACIDIFICATION" [color="#F1948A"];
"gene_ATP6V0A1" -- "term_3_GOBP_VACUOLAR_ACIDIFICATION" [color="#F1948A"];
"gene_ATP6V0D1" -- "term_3_GOBP_VACUOLAR_ACIDIFICATION" [color="#F1948A"];
"gene_ATP6V1A" -- "term_3_GOBP_VACUOLAR_ACIDIFICATION" [color="#F1948A"];
"gene_ATP6V1B2" -- "term_3_GOBP_VACUOLAR_ACIDIFICATION" [color="#F1948A"];
"gene_BTK" -- "term_1_GOBP_PHAGOCYTOSIS" [color="#F1948A"];
"gene_CARD8" -- "term_2_GOBP_CYTOKINE_MEDIATED_SIGNALING_PATHWAY" [color="#F1948A"];
"gene_CASP1" -- "term_2_GOBP_CYTOKINE_MEDIATED_SIGNALING_PATHWAY" [color="#F1948A"];
"gene_CCR5" -- "term_2_GOBP_CYTOKINE_MEDIATED_SIGNALING_PATHWAY" [color="#F1948A"];
"gene_CD300A" -- "term_1_GOBP_PHAGOCYTOSIS" [color="#F1948A"];
"gene_CD36" -- "term_1_GOBP_PHAGOCYTOSIS" [color="#F1948A"];
"gene_CD4" -- "term_2_GOBP_CYTOKINE_MEDIATED_SIGNALING_PATHWAY" [color="#F1948A"];
"gene_CD93" -- "term_1_GOBP_PHAGOCYTOSIS" [color="#F1948A"];
"gene_CIDEB" -- "term_5_GOBP_GOLGI_VESICLE_TRANSPORT" [color="#F1948A"];
"gene_CNIH4" -- "term_5_GOBP_GOLGI_VESICLE_TRANSPORT" [color="#F1948A"];
"gene_CORO1A" -- "term_1_GOBP_PHAGOCYTOSIS" [color="#F1948A"];
"gene_CORO1B" -- "term_8_GOBP_ARP2_3_COMPLEX_MEDIATED_ACTIN_NUCLEATION" [color="#F1948A"];
"gene_CR1" -- "term_9_GOBP_REGULATION_OF_PEPTIDASE_ACTIVITY" [color="#F1948A"];
"gene_CSF1R" -- "term_2_GOBP_CYTOKINE_MEDIATED_SIGNALING_PATHWAY" [color="#F1948A"];
"gene_CSF2RB" -- "term_2_GOBP_CYTOKINE_MEDIATED_SIGNALING_PATHWAY" [color="#F1948A"];
"gene_CSF3R" -- "term_2_GOBP_CYTOKINE_MEDIATED_SIGNALING_PATHWAY" [color="#F1948A"];
"gene_CSK" -- "term_1_GOBP_PHAGOCYTOSIS" [color="#F1948A"];
"gene_CX3CR1" -- "term_2_GOBP_CYTOKINE_MEDIATED_SIGNALING_PATHWAY" [color="#F1948A"];
"gene_CYFIP1" -- "term_8_GOBP_ARP2_3_COMPLEX_MEDIATED_ACTIN_NUCLEATION" [color="#F1948A"];
"gene_DGAT2" -- "term_4_GOBP_AMIDE_METABOLIC_PROCESS" [color="#F1948A"];
"gene_DMXL2" -- "term_3_GOBP_VACUOLAR_ACIDIFICATION" [color="#F1948A"];
"gene_EIF4E2" -- "term_2_GOBP_CYTOKINE_MEDIATED_SIGNALING_PATHWAY" [color="#F1948A"];
"gene_FAR2" -- "term_4_GOBP_AMIDE_METABOLIC_PROCESS" [color="#F1948A"];
"gene_FPR2" -- "term_1_GOBP_PHAGOCYTOSIS" [color="#F1948A"];
"gene_GBA1" -- "term_4_GOBP_AMIDE_METABOLIC_PROCESS" [color="#F1948A"];
"gene_GK" -- "term_6_GOBP_ALCOHOL_CATABOLIC_PROCESS" [color="#F1948A"];
"gene_GRN" -- "term_3_GOBP_VACUOLAR_ACIDIFICATION" [color="#F1948A"];
"gene_GSN" -- "term_1_GOBP_PHAGOCYTOSIS" [color="#F1948A"];
"gene_HAL" -- "term_4_GOBP_AMIDE_METABOLIC_PROCESS" [color="#F1948A"];
"gene_HCK" -- "term_1_GOBP_PHAGOCYTOSIS" [color="#F1948A"];
"gene_IDH1" -- "term_4_GOBP_AMIDE_METABOLIC_PROCESS" [color="#F1948A"];
"gene_IFITM2" -- "term_2_GOBP_CYTOKINE_MEDIATED_SIGNALING_PATHWAY" [color="#F1948A"];
"gene_IFITM3" -- "term_2_GOBP_CYTOKINE_MEDIATED_SIGNALING_PATHWAY" [color="#F1948A"];
"gene_IFNAR1" -- "term_2_GOBP_CYTOKINE_MEDIATED_SIGNALING_PATHWAY" [color="#F1948A"];
"gene_IFNGR2" -- "term_2_GOBP_CYTOKINE_MEDIATED_SIGNALING_PATHWAY" [color="#F1948A"];
"gene_IL10RB" -- "term_2_GOBP_CYTOKINE_MEDIATED_SIGNALING_PATHWAY" [color="#F1948A"];
"gene_IL17RA" -- "term_2_GOBP_CYTOKINE_MEDIATED_SIGNALING_PATHWAY" [color="#F1948A"];
"gene_ILK" -- "term_2_GOBP_CYTOKINE_MEDIATED_SIGNALING_PATHWAY" [color="#F1948A"];
"gene_JAGN1" -- "term_2_GOBP_CYTOKINE_MEDIATED_SIGNALING_PATHWAY" [color="#F1948A"];
"gene_KCNE3" -- "term_10_GOBP_MONOATOMIC_ANION_HOMEOSTASIS" [color="#F1948A"];
"gene_KDELR1" -- "term_5_GOBP_GOLGI_VESICLE_TRANSPORT" [color="#F1948A"];
"gene_KDELR2" -- "term_5_GOBP_GOLGI_VESICLE_TRANSPORT" [color="#F1948A"];
"gene_LAMP2" -- "term_3_GOBP_VACUOLAR_ACIDIFICATION" [color="#F1948A"];
"gene_LILRA1" -- "term_2_GOBP_CYTOKINE_MEDIATED_SIGNALING_PATHWAY" [color="#F1948A"];
"gene_LILRA2" -- "term_2_GOBP_CYTOKINE_MEDIATED_SIGNALING_PATHWAY" [color="#F1948A"];
"gene_LILRA3" -- "term_2_GOBP_CYTOKINE_MEDIATED_SIGNALING_PATHWAY" [color="#F1948A"];
"gene_LILRA6" -- "term_2_GOBP_CYTOKINE_MEDIATED_SIGNALING_PATHWAY" [color="#F1948A"];
"gene_LILRB1" -- "term_2_GOBP_CYTOKINE_MEDIATED_SIGNALING_PATHWAY" [color="#F1948A"];
"gene_LILRB2" -- "term_2_GOBP_CYTOKINE_MEDIATED_SIGNALING_PATHWAY" [color="#F1948A"];
"gene_LILRB3" -- "term_2_GOBP_CYTOKINE_MEDIATED_SIGNALING_PATHWAY" [color="#F1948A"];
"gene_LRP1" -- "term_1_GOBP_PHAGOCYTOSIS" [color="#F1948A"];
"gene_LRRK2" -- "term_5_GOBP_GOLGI_VESICLE_TRANSPORT" [color="#F1948A"];
"gene_LYST" -- "term_1_GOBP_PHAGOCYTOSIS" [color="#F1948A"];
"gene_MAPK1" -- "term_2_GOBP_CYTOKINE_MEDIATED_SIGNALING_PATHWAY" [color="#F1948A"];
"gene_MCCC2" -- "term_4_GOBP_AMIDE_METABOLIC_PROCESS" [color="#F1948A"];
"gene_MGST2" -- "term_4_GOBP_AMIDE_METABOLIC_PROCESS" [color="#F1948A"];
"gene_MIR21" -- "term_2_GOBP_CYTOKINE_MEDIATED_SIGNALING_PATHWAY" [color="#F1948A"];
"gene_MPEG1" -- "term_7_GOBP_ANTIBACTERIAL_INNATE_IMMUNE_RESPONSE" [color="#F1948A"];
"gene_NAAA" -- "term_4_GOBP_AMIDE_METABOLIC_PROCESS" [color="#F1948A"];
"gene_NAGK" -- "term_4_GOBP_AMIDE_METABOLIC_PROCESS" [color="#F1948A"];
"gene_NAIP" -- "term_2_GOBP_CYTOKINE_MEDIATED_SIGNALING_PATHWAY" [color="#F1948A"];
"gene_NCF2" -- "term_1_GOBP_PHAGOCYTOSIS" [color="#F1948A"];
"gene_NCF4" -- "term_1_GOBP_PHAGOCYTOSIS" [color="#F1948A"];
"gene_NOD2" -- "term_7_GOBP_ANTIBACTERIAL_INNATE_IMMUNE_RESPONSE" [color="#F1948A"];
"gene_NPL" -- "term_4_GOBP_AMIDE_METABOLIC_PROCESS" [color="#F1948A"];
"gene_OGDH" -- "term_4_GOBP_AMIDE_METABOLIC_PROCESS" [color="#F1948A"];
"gene_ORMDL2" -- "term_4_GOBP_AMIDE_METABOLIC_PROCESS" [color="#F1948A"];
"gene_PADI2" -- "term_2_GOBP_CYTOKINE_MEDIATED_SIGNALING_PATHWAY" [color="#F1948A"];
"gene_PAK1" -- "term_1_GOBP_PHAGOCYTOSIS" [color="#F1948A"];
"gene_PANK2" -- "term_4_GOBP_AMIDE_METABOLIC_PROCESS" [color="#F1948A"];
"gene_PECAM1" -- "term_1_GOBP_PHAGOCYTOSIS" [color="#F1948A"];
"gene_PMVK" -- "term_4_GOBP_AMIDE_METABOLIC_PROCESS" [color="#F1948A"];
"gene_PPCDC" -- "term_4_GOBP_AMIDE_METABOLIC_PROCESS" [color="#F1948A"];
"gene_PPP2CB" -- "term_2_GOBP_CYTOKINE_MEDIATED_SIGNALING_PATHWAY" [color="#F1948A"];
"gene_PRCP" -- "term_4_GOBP_AMIDE_METABOLIC_PROCESS" [color="#F1948A"];
"gene_PRELID1" -- "term_9_GOBP_REGULATION_OF_PEPTIDASE_ACTIVITY" [color="#F1948A"];
"gene_PRKACA" -- "term_1_GOBP_PHAGOCYTOSIS" [color="#F1948A"];
"gene_PRKCD" -- "term_1_GOBP_PHAGOCYTOSIS" [color="#F1948A"];
"gene_PSEN1" -- "term_4_GOBP_AMIDE_METABOLIC_PROCESS" [color="#F1948A"];
```

"gene_PTPN6" -- "term_2_GOBP_CYTOKINE_MEDIATED_SIGNALING_PATHWAY" [color="#F1948A"];
"gene_PTPRJ" -- "term_1_GOBP_PHAGOCYTOSIS" [color="#F1948A"];
"gene_PYCARD" -- "term_1_GOBP_PHAGOCYTOSIS" [color="#F1948A"];
"gene_RAB10" -- "term_5_GOBP_GOLGI_VESICLE_TRANSPORT" [color="#F1948A"];
"gene_RAB1B" -- "term_5_GOBP_GOLGI_VESICLE_TRANSPORT" [color="#F1948A"];
"gene_RAB27A" -- "term_1_GOBP_PHAGOCYTOSIS" [color="#F1948A"];
"gene_RAB4B" -- "term_5_GOBP_GOLGI_VESICLE_TRANSPORT" [color="#F1948A"];
"gene_RAB7A" -- "term_1_GOBP_PHAGOCYTOSIS" [color="#F1948A"];
"gene_RAB8A" -- "term_5_GOBP_GOLGI_VESICLE_TRANSPORT" [color="#F1948A"];
"gene_RBM47" -- "term_2_GOBP_CYTOKINE_MEDIATED_SIGNALING_PATHWAY" [color="#F1948A"];
"gene_RTN1" -- "term_4_GOBP_AMIDE_METABOLIC_PROCESS" [color="#F1948A"];
"gene_SEC23B" -- "term_5_GOBP_GOLGI_VESICLE_TRANSPORT" [color="#F1948A"];
"gene_SEC23IP" -- "term_5_GOBP_GOLGI_VESICLE_TRANSPORT" [color="#F1948A"];
"gene_SERPINB1" -- "term_9_GOBP_REGULATION_OF_PEPTIDASE_ACTIVITY" [color="#F1948A"];
"gene_SIRPA" -- "term_1_GOBP_PHAGOCYTOSIS" [color="#F1948A"];
"gene_SIRPB1" -- "term_1_GOBP_PHAGOCYTOSIS" [color="#F1948A"];
"gene_SLC11A1" -- "term_1_GOBP_PHAGOCYTOSIS" [color="#F1948A"];
"gene_SLC12A6" -- "term_10_GOBP_MONOATOMIC_ANION_HOMEOSTASIS" [color="#F1948A"];
"gene_SNX1" -- "term_5_GOBP_GOLGI_VESICLE_TRANSPORT" [color="#F1948A"];
"gene_SORT1" -- "term_5_GOBP_GOLGI_VESICLE_TRANSPORT" [color="#F1948A"];
"gene_SPTLC2" -- "term_4_GOBP_AMIDE_METABOLIC_PROCESS" [color="#F1948A"];
"gene_SPTSSA" -- "term_4_GOBP_AMIDE_METABOLIC_PROCESS" [color="#F1948A"];
"gene_ST8SIA4" -- "term_4_GOBP_AMIDE_METABOLIC_PROCESS" [color="#F1948A"];
"gene_STX6" -- "term_5_GOBP_GOLGI_VESICLE_TRANSPORT" [color="#F1948A"];
"gene_SULT1A1" -- "term_6_GOBP_ALCOHOL_CATABOLIC_PROCESS" [color="#F1948A"];
"gene_SULT1A3" -- "term_6_GOBP_ALCOHOL_CATABOLIC_PROCESS" [color="#F1948A"];
"gene_SULT1A4" -- "term_6_GOBP_ALCOHOL_CATABOLIC_PROCESS" [color="#F1948A"];
"gene_SYK" -- "term_1_GOBP_PHAGOCYTOSIS" [color="#F1948A"];
"gene_SYT11" -- "term_1_GOBP_PHAGOCYTOSIS" [color="#F1948A"];
"gene_TBXAS1" -- "term_10_GOBP_MONOATOMIC_ANION_HOMEOSTASIS" [color="#F1948A"];
"gene_TCIRG1" -- "term_3_GOBP_VACUOLAR_ACIDIFICATION" [color="#F1948A"];
"gene_TFEB" -- "term_7_GOBP_ANTIBACTERIAL_INNATE_IMMUNE_RESPONSE" [color="#F1948A"];
"gene_TIMP2" -- "term_9_GOBP_REGULATION_OF_PEPTIDASE_ACTIVITY" [color="#F1948A"];
"gene_TNFRSF1A" -- "term_2_GOBP_CYTOKINE_MEDIATED_SIGNALING_PATHWAY" [color="#F1948A"];
"gene_TPI1" -- "term_6_GOBP_ALCOHOL_CATABOLIC_PROCESS" [color="#F1948A"];
"gene_VAV1" -- "term_1_GOBP_PHAGOCYTOSIS" [color="#F1948A"];
"gene_VIPAS39" -- "term_1_GOBP_PHAGOCYTOSIS" [color="#F1948A"];
"gene_VNN2" -- "term_4_GOBP_AMIDE_METABOLIC_PROCESS" [color="#F1948A"];
"gene_WASHC5" -- "term_8_GOBP_ARP2_3_COMPLEX_MEDIATED_ACTIN_NUCLEATION" [color="#F1948A"];
"gene_WBP1L" -- "term_2_GOBP_CYTOKINE_MEDIATED_SIGNALING_PATHWAY" [color="#F1948A"];
"gene_ARAP3" [label="ARAP3", type="gene", width=0.7, height=0.7];
"gene_BEST1" [label="BEST1", type="gene", width=0.7, height=0.7];
"gene_CLEC4E" [label="CLEC4E", type="gene", width=0.7, height=0.7];
"gene_DRAM1" [label="DRAM1", type="gene", width=0.7, height=0.7];
"gene_EFHD2" [label="EFHD2", type="gene", width=0.7, height=0.7];
"gene_FGD2" [label="FGD2", type="gene", width=0.7, height=0.7];
"gene_FGD4" [label="FGD4", type="gene", width=0.7, height=0.7];
"gene_KCNK6" [label="KCNK6", type="gene", width=0.7, height=0.7];
"gene_LAT2" [label="LAT2", type="gene", width=0.7, height=0.7];
"gene_MBOAT7" [label="MBOAT7", type="gene", width=0.7, height=0.7];
"gene_MICU1" [label="MICU1", type="gene", width=0.7, height=0.7];
"gene_OSTM1" [label="OSTM1", type="gene", width=0.7, height=0.7];
"gene_PACSIN2" [label="PACSIN2", type="gene", width=0.7, height=0.7];
"gene_RIPOR2" [label="RIPOR2", type="gene", width=0.7, height=0.7];
"gene_RRAGD" [label="RRAGD", type="gene", width=0.7, height=0.7];
"gene_SELPLG" [label="SELPLG", type="gene", width=0.7, height=0.7];
"gene_TM9SF1" [label="TM9SF1", type="gene", width=0.7, height=0.7];
"gene_TRIM14" [label="TRIM14", type="gene", width=0.7, height=0.7];
"gene_UBA1" [label="UBA1", type="gene", width=0.7, height=0.7];
"gene_VPS35" [label="VPS35", type="gene", width=0.7, height=0.7];
"gene_WDFY3" [label="WDFY3", type="gene", width=0.7, height=0.7];
"gene_ZYX" [label="ZYX", type="gene", width=0.7, height=0.7];
"term_10_GOCC_ORGANELLE_MEMBRANE_CONTACT_SITE" [label="GOCC: ORGANELLE MEMBRANE CONTACT SITE", style=filled, fillcolor="#85C1E9", fontcolor="white", type="term", width=2, height=2, library="C5: Gene Ontology: Cellular Component"];
"term_1_GOCC_FICOLIN_1_RICH_GRANULE" [label="GOCC: FICOLIN 1 RICH GRANULE", style=filled, fillcolor="#85C1E9", fontcolor="white", type="term", width=2, height=2, library="C5: Gene Ontology: Cellular Component"];
"term_2_GOCC_PHAGOCYTIC_VESICLE" [label="GOCC: PHAGOCYTIC VESICLE", style=filled, fillcolor="#85C1E9", fontcolor="white", type="term", width=2, height=2, library="C5: Gene Ontology: Cellular Component"];
"term_3_GOCC_VACUOLAR_MEMBRANE" [label="GOCC: VACUOLAR MEMBRANE", style=filled, fillcolor="#85C1E9", fontcolor="white", type="term", width=2, height=2, library="C5: Gene Ontology: Cellular Component"];
"term_4_GOCC_CELL_LEADING_EDGE" [label="GOCC: CELL LEADING EDGE", style=filled, fillcolor="#85C1E9", fontcolor="white", type="term", width=2, height=2, library="C5: Gene Ontology: Cellular Component"];
"term_5_GOCC_CANONICAL_INFLAMMASOME_COMPLEX" [label="GOCC: CANONICAL INFLAMMASOME COMPLEX", style=filled, fillcolor="#85C1E9", fontcolor="white", type="term", width=2, height=2, library="C5: Gene Ontology: Cellular Component"];
"term_6_GOCC_MEMBRANE_MICRODOMAIN" [label="GOCC: MEMBRANE MICRODOMAIN", style=filled, fillcolor="#85C1E9", fontcolor="white", type="term", width=2, height=2, library="C5: Gene Ontology: Cellular Component"];
"term_7_GOCC_PALMITOYLTRANSFERASE_COMPLEX" [label="GOCC: PALMITOYLTRANSFERASE COMPLEX", style=filled, fillcolor="#85C1E9", fontcolor="white", type="term", width=2, height=2, library="C5: Gene Ontology: Cellular Component"];
"term_8_GOCC_AZUROPHIL_GRANULE_LUMEN" [label="GOCC: AZUROPHIL GRANULE LUMEN", style=filled, fillcolor="#85C1E9", fontcolor="white", type="term", width=2, height=2, library="C5: Gene Ontology: Cellular Component"];
"term_9_GOCC_COPI_COATED_VESICLE" [label="GOCC: COPI COATED VESICLE", style=filled, fillcolor="#85C1E9", fontcolor="white", type="term", width=2, height=2, library="C5: Gene Ontology: Cellular Component"];
"gene_ABCA7" -- "term_4_GOCC_CELL_LEADING_EDGE" [color="#85C1E9"];
"gene_ACLY" -- "term_1_GOCC_FICOLIN_1_RICH_GRANULE" [color="#85C1E9"];
"gene_ACTR3" -- "term_4_GOCC_CELL_LEADING_EDGE" [color="#85C1E9"];
"gene_ADA2" -- "term_8_GOCC_AZUROPHIL_GRANULE_LUMEN" [color="#85C1E9"];
"gene_AHCYL1" -- "term_10_GOCC_ORGANELLE_MEMBRANE_CONTACT_SITE" [color="#85C1E9"];
"gene_ANPEP" -- "term_3_GOCC_VACUOLAR_MEMBRANE" [color="#85C1E9"];
"gene_APAF1" -- "term_1_GOCC_FICOLIN_1_RICH_GRANULE" [color="#85C1E9"];
"gene_APBB1IP" -- "term_4_GOCC_CELL_LEADING_EDGE" [color="#85C1E9"];
"gene_APEH" -- "term_1_GOCC_FICOLIN_1_RICH_GRANULE" [color="#85C1E9"];
"gene_ARAP1" -- "term_4_GOCC_CELL_LEADING_EDGE" [color="#85C1E9"];
"gene_ARAP3" -- "term_4_GOCC_CELL_LEADING_EDGE" [color="#85C1E9"];
"gene_ATG7" -- "term_1_GOCC_FICOLIN_1_RICH_GRANULE" [color="#85C1E9"];
"gene_ATP6AP1" -- "term_3_GOCC_VACUOLAR_MEMBRANE" [color="#85C1E9"];
"gene_ATP6V0A1" -- "term_1_GOCC_FICOLIN_1_RICH_GRANULE" [color="#85C1E9"];
"gene_ATP6V0D1" -- "term_2_GOCC_PHAGOCYTIC_VESICLE" [color="#85C1E9"];
"gene_ATP6V1A" -- "term_3_GOCC_VACUOLAR_MEMBRANE" [color="#85C1E9"];
"gene_ATP6V1B2" -- "term_3_GOCC_VACUOLAR_MEMBRANE" [color="#85C1E9"];
"gene_BEST1" -- "term_6_GOCC_MEMBRANE_MICRODOMAIN" [color="#85C1E9"];
"gene_BRI3" -- "term_3_GOCC_VACUOLAR_MEMBRANE" [color="#85C1E9"];
"gene_BTK" -- "term_6_GOCC_MEMBRANE_MICRODOMAIN" [color="#85C1E9"];
"gene_CAP1" -- "term_8_GOCC_AZUROPHIL_GRANULE_LUMEN" [color="#85C1E9"];
"gene_CAPG" -- "term_4_GOCC_CELL_LEADING_EDGE" [color="#85C1E9"];
"gene_CAPZB" -- "term_4_GOCC_CELL_LEADING_EDGE" [color="#85C1E9"];
"gene_CARD8" -- "term_5_GOCC_CANONICAL_INFLAMMASOME_COMPLEX" [color="#85C1E9"];
"gene_CASP1" -- "term_5_GOCC_CANONICAL_INFLAMMASOME_COMPLEX" [color="#85C1E9"];
"gene_CD300A" -- "term_1_GOCC_FICOLIN_1_RICH_GRANULE" [color="#85C1E9"];
"gene_CD36" -- "term_2_GOCC_PHAGOCYTIC_VESICLE" [color="#85C1E9"];
"gene_CD4" -- "term_6_GOCC_MEMBRANE_MICRODOMAIN" [color="#85C1E9"];
"gene_CD58" -- "term_1_GOCC_FICOLIN_1_RICH_GRANULE" [color="#85C1E9"];
"gene_CD93" -- "term_1_GOCC_FICOLIN_1_RICH_GRANULE" [color="#85C1E9"];
"gene_CFD" -- "term_1_GOCC_FICOLIN_1_RICH_GRANULE" [color="#85C1E9"];
"gene_CIDEB" -- "term_9_GOCC_COPI_COATED_VESICLE" [color="#85C1E9"];
"gene_CLEC4E" -- "term_2_GOCC_PHAGOCYTIC_VESICLE" [color="#85C1E9"];
"gene_CLTA" -- "term_3_GOCC_VACUOLAR_MEMBRANE" [color="#85C1E9"];
"gene_CORO1A" -- "term_2_GOCC_PHAGOCYTIC_VESICLE" [color="#85C1E9"];
"gene_CORO1B" -- "term_4_GOCC_CELL_LEADING_EDGE" [color="#85C1E9"];
"gene_CPPED1" -- "term_8_GOCC_AZUROPHIL_GRANULE_LUMEN" [color="#85C1E9"];
"gene_CR1" -- "term_1_GOCC_FICOLIN_1_RICH_GRANULE" [color="#85C1E9"];
"gene_CSF3R" -- "term_3_GOCC_VACUOLAR_MEMBRANE" [color="#85C1E9"];
"gene_CSTB" -- "term_1_GOCC_FICOLIN_1_RICH_GRANULE" [color="#85C1E9"];
"gene_CTSD" -- "term_1_GOCC_FICOLIN_1_RICH_GRANULE" [color="#85C1E9"];
"gene_CYFIP1" -- "term_4_GOCC_CELL_LEADING_EDGE" [color="#85C1E9"];
"gene_DAGLB" -- "term_3_GOCC_VACUOLAR_MEMBRANE" [color="#85C1E9"];
"gene_DRAM1" -- "term_3_GOCC_VACUOLAR_MEMBRANE" [color="#85C1E9"];
"gene_EFHD2" -- "term_6_GOCC_MEMBRANE_MICRODOMAIN" [color="#85C1E9"];
"gene_ENTPD1" -- "term_6_GOCC_MEMBRANE_MICRODOMAIN" [color="#85C1E9"];
"gene_ENTPD4" -- "term_3_GOCC_VACUOLAR_MEMBRANE" [color="#85C1E9"];
"gene_FGD2" -- "term_4_GOCC_CELL_LEADING_EDGE" [color="#85C1E9"];
"gene_FGD4" -- "term_4_GOCC_CELL_LEADING_EDGE" [color="#85C1E9"];
"gene_FGL2" -- "term_1_GOCC_FICOLIN_1_RICH_GRANULE" [color="#85C1E9"];
"gene_FPR2" -- "term_1_GOCC_FICOLIN_1_RICH_GRANULE" [color="#85C1E9"];
"gene_GAA" -- "term_1_GOCC_FICOLIN_1_RICH_GRANULE" [color="#85C1E9"];
"gene_GBA1" -- "term_3_GOCC_VACUOLAR_MEMBRANE" [color="#85C1E9"];
"gene_GCA" -- "term_8_GOCC_AZUROPHIL_GRANULE_LUMEN" [color="#85C1E9"];
"gene_GNS" -- "term_1_GOCC_FICOLIN_1_RICH_GRANULE" [color="#85C1E9"];
"gene_GRN" -- "term_3_GOCC_VACUOLAR_MEMBRANE" [color="#85C1E9"];
"gene_GSN" -- "term_1_GOCC_FICOLIN_1_RICH_GRANULE" [color="#85C1E9"];
"gene_HCK" -- "term_6_GOCC_MEMBRANE_MICRODOMAIN" [color="#85C1E9"];
"gene_HK1" -- "term_6_GOCC_MEMBRANE_MICRODOMAIN" [color="#85C1E9"];
"gene_HK3" -- "term_1_GOCC_FICOLIN_1_RICH_GRANULE" [color="#85C1E9"];
"gene_HSPA6" -- "term_1_GOCC_FICOLIN_1_RICH_GRANULE" [color="#85C1E9"];
"gene_IDH1" -- "term_1_GOCC_FICOLIN_1_RICH_GRANULE" [color="#85C1E9"];
"gene_IFITM2" -- "term_3_GOCC_VACUOLAR_MEMBRANE" [color="#85C1E9"];
"gene_IFITM3" -- "term_3_GOCC_VACUOLAR_MEMBRANE" [color="#85C1E9"];
"gene_ILK" -- "term_4_GOCC_CELL_LEADING_EDGE" [color="#85C1E9"];
"gene_IQGAP1" -- "term_4_GOCC_CELL_LEADING_EDGE" [color="#85C1E9"];
"gene_ITGAX" -- "term_1_GOCC_FICOLIN_1_RICH_GRANULE" [color="#85C1E9"];
"gene_JUP" -- "term_1_GOCC_FICOLIN_1_RICH_GRANULE" [color="#85C1E9"];
"gene_KCNE3" -- "term_6_GOCC_MEMBRANE_MICRODOMAIN" [color="#85C1E9"];
"gene_KCNK6" -- "term_3_GOCC_VACUOLAR_MEMBRANE" [color="#85C1E9"];
"gene_KDELR1" -- "term_9_GOCC_COPI_COATED_VESICLE" [color="#85C1E9"];
"gene_KDELR2" -- "term_9_GOCC_COPI_COATED_VESICLE" [color="#85C1E9"];
"gene_LAMP2" -- "term_1_GOCC_FICOLIN_1_RICH_GRANULE" [color="#85C1E9"];
"gene_LAT2" -- "term_6_GOCC_MEMBRANE_MICRODOMAIN" [color="#85C1E9"];
"gene_LILRA3" -- "term_1_GOCC_FICOLIN_1_RICH_GRANULE" [color="#85C1E9"];
"gene_LILRB2" -- "term_1_GOCC_FICOLIN_1_RICH_GRANULE" [color="#85C1E9"];
"gene_LRP1" -- "term_3_GOCC_VACUOLAR_MEMBRANE" [color="#85C1E9"];
"gene_LRRK2" -- "term_2_GOCC_PHAGOCYTIC_VESICLE" [color="#85C1E9"];
"gene_M6PR" -- "term_3_GOCC_VACUOLAR_MEMBRANE" [color="#85C1E9"];
"gene_MAPK1" -- "term_1_GOCC_FICOLIN_1_RICH_GRANULE" [color="#85C1E9"];
"gene_MBOAT7" -- "term_10_GOCC_ORGANELLE_MEMBRANE_CONTACT_SITE" [color="#85C1E9"];
"gene_MICU1" -- "term_10_GOCC_ORGANELLE_MEMBRANE_CONTACT_SITE" [color="#85C1E9"];
"gene_MNDA" -- "term_1_GOCC_FICOLIN_1_RICH_GRANULE" [color="#85C1E9"];
"gene_MOSPD2" -- "term_10_GOCC_ORGANELLE_MEMBRANE_CONTACT_SITE" [color="#85C1E9"];
"gene_MPEG1" -- "term_2_GOCC_PHAGOCYTIC_VESICLE" [color="#85C1E9"];
"gene_NAIP" -- "term_5_GOCC_CANONICAL_INFLAMMASOME_COMPLEX" [color="#85C1E9"];
"gene_NCF1" -- "term_2_GOCC_PHAGOCYTIC_VESICLE" [color="#85C1E9"];
"gene_NCF2" -- "term_2_GOCC_PHAGOCYTIC_VESICLE" [color="#85C1E9"];
"gene_NCF4" -- "term_2_GOCC_PHAGOCYTIC_VESICLE" [color="#85C1E9"];
"gene_NOD2" -- "term_2_GOCC_PHAGOCYTIC_VESICLE" [color="#85C1E9"];
"gene_ORMDL2" -- "term_7_GOCC_PALMITOYLTRANSFERASE_COMPLEX" [color="#85C1E9"];
"gene_OSTM1" -- "term_3_GOCC_VACUOLAR_MEMBRANE" [color="#85C1E9"];
"gene_PACSIN2" -- "term_4_GOCC_CELL_LEADING_EDGE" [color="#85C1E9"];
"gene_PADI2" -- "term_8_GOCC_AZUROPHIL_GRANULE_LUMEN" [color="#85C1E9"];
"gene_PAK1" -- "term_4_GOCC_CELL_LEADING_EDGE" [color="#85C1E9"];
"gene_PECAM1" -- "term_6_GOCC_MEMBRANE_MICRODOMAIN" [color="#85C1E9"];
"gene_PRCP" -- "term_1_GOCC_FICOLIN_1_RICH_GRANULE" [color="#85C1E9"];
"gene_PRKACA" -- "term_6_GOCC_MEMBRANE_MICRODOMAIN" [color="#85C1E9"];
"gene_PRKAR1A" -- "term_6_GOCC_MEMBRANE_MICRODOMAIN" [color="#85C1E9"];
"gene_PRKCD" -- "term_8_GOCC_AZUROPHIL_GRANULE_LUMEN" [color="#85C1E9"];
"gene_PSEN1" -- "term_3_GOCC_VACUOLAR_MEMBRANE" [color="#85C1E9"];
"gene_PTPRJ" -- "term_4_GOCC_CELL_LEADING_EDGE" [color="#85C1E9"];
"gene_PYCARD" -- "term_5_GOCC_CANONICAL_INFLAMMASOME_COMPLEX" [color="#85C1E9"];
"gene_QPCT" -- "term_1_GOCC_FICOLIN_1_RICH_GRANULE" [color="#85C1E9"];
"gene_RAB10" -- "term_2_GOCC_PHAGOCYTIC_VESICLE" [color="#85C1E9"];
"gene_RAB7A" -- "term_2_GOCC_PHAGOCYTIC_VESICLE" [color="#85C1E9"];
"gene_RAB8A" -- "term_2_GOCC_PHAGOCYTIC_VESICLE" [color="#85C1E9"];
"gene_RIGI" -- "term_4_GOCC_CELL_LEADING_EDGE" [color="#85C1E9"];
"gene_RIPOR2" -- "term_4_GOCC_CELL_LEADING_EDGE" [color="#85C1E9"];
"gene_RRAGD" -- "term_3_GOCC_VACUOLAR_MEMBRANE" [color="#85C1E9"];
"gene_S100A11" -- "term_4_GOCC_CELL_LEADING_EDGE" [color="#85C1E9"];
"gene_SELPLG" -- "term_6_GOCC_MEMBRANE_MICRODOMAIN" [color="#85C1E9"];
"gene_SERPINA1" -- "term_1_GOCC_FICOLIN_1_RICH_GRANULE" [color="#85C1E9"];

"gene_SIGLEC5" -- "term_1_GOCC_FICOLIN_1_RICH_GRANULE" [color="#85C1E9"];
"gene_SIRPA" -- "term_1_GOCC_FICOLIN_1_RICH_GRANULE" [color="#85C1E9"];
"gene_SLC11A1" -- "term_1_GOCC_FICOLIN_1_RICH_GRANULE" [color="#85C1E9"];
"gene_SNX1" -- "term_4_GOCC_CELL_LEADING_EDGE" [color="#85C1E9"];
"gene_SORT1" -- "term_3_GOCC_VACUOLAR_MEMBRANE" [color="#85C1E9"];
"gene_SPTLC2" -- "term_7_GOCC_PALMITOYLTRANSFERASE_COMPLEX" [color="#85C1E9"];
"gene_SPTSSA" -- "term_7_GOCC_PALMITOYLTRANSFERASE_COMPLEX" [color="#85C1E9"];
"gene_STX3" -- "term_4_GOCC_CELL_LEADING_EDGE" [color="#85C1E9"];
"gene_STX6" -- "term_2_GOCC_PHAGOCYTIC_VESICLE" [color="#85C1E9"];
"gene_SYK" -- "term_2_GOCC_PHAGOCYTIC_VESICLE" [color="#85C1E9"];
"gene_SYT11" -- "term_2_GOCC_PHAGOCYTIC_VESICLE" [color="#85C1E9"];
"gene_TCIRG1" -- "term_1_GOCC_FICOLIN_1_RICH_GRANULE" [color="#85C1E9"];
"gene_TFEB" -- "term_3_GOCC_VACUOLAR_MEMBRANE" [color="#85C1E9"];
"gene_TIMP2" -- "term_1_GOCC_FICOLIN_1_RICH_GRANULE" [color="#85C1E9"];
"gene_TM9SF1" -- "term_3_GOCC_VACUOLAR_MEMBRANE" [color="#85C1E9"];
"gene_TMEM179B" -- "term_1_GOCC_FICOLIN_1_RICH_GRANULE" [color="#85C1E9"];
"gene_TNFRSF1A" -- "term_6_GOCC_MEMBRANE_MICRODOMAIN" [color="#85C1E9"];
"gene_TRIM14" -- "term_2_GOCC_PHAGOCYTIC_VESICLE" [color="#85C1E9"];
"gene_UBA1" -- "term_3_GOCC_VACUOLAR_MEMBRANE" [color="#85C1E9"];
"gene_VMP1" -- "term_3_GOCC_VACUOLAR_MEMBRANE" [color="#85C1E9"];
"gene_VPS35" -- "term_3_GOCC_VACUOLAR_MEMBRANE" [color="#85C1E9"];
"gene_WDFY3" -- "term_3_GOCC_VACUOLAR_MEMBRANE" [color="#85C1E9"];
"gene_ZYX" -- "term_2_GOCC_PHAGOCYTIC_VESICLE" [color="#85C1E9"];
"gene_APTX" [label="APTX", type="gene", width=0.7, height=0.7];
"gene_CALML4" [label="CALML4", type="gene", width=0.7, height=0.7];
"gene_NCF1B" [label="NCF1B", type="gene", width=0.7, height=0.7];
"gene_NCF1C" [label="NCF1C", type="gene", width=0.7, height=0.7];
"gene_SH3BP2" [label="SH3BP2", type="gene", width=0.7, height=0.7];
"gene_TLR5" [label="TLR5", type="gene", width=0.7, height=0.7];
"gene_TRIOBP" [label="TRIOBP", type="gene", width=0.7, height=0.7];
"gene_UBASH3B" [label="UBASH3B", type="gene", width=0.7, height=0.7];
"term_10_GOMF_PHOSPHATIDYLINOSITOL_3_4_5_TRISPHOSPHATE_BINDING" [label="GOMF: PHOSPHATIDYLINOSITOL 3 4 5 TRISPHOSPHATE BINDING", style=filled, fillcolor="#D7BDE2", fontcolor="white", type="term", width=2, height=2, library="C5: Gene Ontology: Molecular Function"];
"term_1_GOMF_INHIBITORY_MHC_CLASS_I_RECEPTOR_ACTIVITY" [label="GOMF: INHIBITORY MHC CLASS I RECEPTOR ACTIVITY", style=filled, fillcolor="#D7BDE2", fontcolor="white", type="term", width=2, height=2, library="C5: Gene Ontology: Molecular Function"];
"term_2_GOMF_PATTERN_RECOGNITION_RECEPTOR_ACTIVITY" [label="GOMF: PATTERN RECOGNITION RECEPTOR ACTIVITY", style=filled, fillcolor="#D7BDE2", fontcolor="white", type="term", width=2, height=2, library="C5: Gene Ontology: Molecular Function"];
"term_3_GOMF_SUPEROXIDE_GENERATING_NADPH_OXIDASE_ACTIVATOR_ACTIVITY" [label="GOMF: SUPEROXIDE GENERATING NADPH OXIDASE ACTIVATOR ACTIVITY", style=filled, fillcolor="#D7BDE2", fontcolor="white", type="term", width=2, height=2, library="C5: Gene Ontology: Molecular Function"];
"term_4_GOMF_SIALIC_ACID_BINDING" [label="GOMF: SIALIC ACID BINDING", style=filled, fillcolor="#D7BDE2", fontcolor="white", type="term", width=2, height=2, library="C5: Gene Ontology: Molecular Function"];
"term_5_GOMF_ATPASE_COUPLED_ION_TRANSMEMBRANE_TRANSPORTER_ACTIVITY" [label="GOMF: ATPASE COUPLED ION TRANSMEMBRANE TRANSPORTER ACTIVITY", style=filled, fillcolor="#D7BDE2", fontcolor="white", type="term", width=2, height=2, library="C5: Gene Ontology: Molecular Function"];
"term_6_GOMF_PHOSPHOPROTEIN_BINDING" [label="GOMF: PHOSPHOPROTEIN BINDING", style=filled, fillcolor="#D7BDE2", fontcolor="white", type="term", width=2, height=2, library="C5: Gene Ontology: Molecular Function"];
"term_7_GOMF_CYTOKINE_RECEPTOR_ACTIVITY" [label="GOMF: CYTOKINE RECEPTOR ACTIVITY", style=filled, fillcolor="#D7BDE2", fontcolor="white", type="term", width=2, height=2, library="C5: Gene Ontology: Molecular Function"];
"term_8_GOMF_MYOSIN_BINDING" [label="GOMF: MYOSIN BINDING", style=filled, fillcolor="#D7BDE2", fontcolor="white", type="term", width=2, height=2, library="C5: Gene Ontology: Molecular Function"];
"term_9_GOMF_RETROMER_COMPLEX_BINDING" [label="GOMF: RETROMER COMPLEX BINDING", style=filled, fillcolor="#D7BDE2", fontcolor="white", type="term", width=2, height=2, library="C5: Gene Ontology: Molecular Function"];
"gene_APTX" -- "term_6_GOMF_PHOSPHOPROTEIN_BINDING" [color="#D7BDE2"];
"gene_ARAP1" -- "term_10_GOMF_PHOSPHATIDYLINOSITOL_3_4_5_TRISPHOSPHATE_BINDING" [color="#D7BDE2"];
"gene_ARAP3" -- "term_10_GOMF_PHOSPHATIDYLINOSITOL_3_4_5_TRISPHOSPHATE_BINDING" [color="#D7BDE2"];
"gene_ARHGAP9" -- "term_10_GOMF_PHOSPHATIDYLINOSITOL_3_4_5_TRISPHOSPHATE_BINDING" [color="#D7BDE2"];
"gene_ASGR2" -- "term_2_GOMF_PATTERN_RECOGNITION_RECEPTOR_ACTIVITY" [color="#D7BDE2"];
"gene_ATP6V0A1" -- "term_5_GOMF_ATPASE_COUPLED_ION_TRANSMEMBRANE_TRANSPORTER_ACTIVITY" [color="#D7BDE2"];
"gene_ATP6V0D1" -- "term_5_GOMF_ATPASE_COUPLED_ION_TRANSMEMBRANE_TRANSPORTER_ACTIVITY" [color="#D7BDE2"];
"gene_ATP6V1A" -- "term_5_GOMF_ATPASE_COUPLED_ION_TRANSMEMBRANE_TRANSPORTER_ACTIVITY" [color="#D7BDE2"];
"gene_ATP6V1B2" -- "term_5_GOMF_ATPASE_COUPLED_ION_TRANSMEMBRANE_TRANSPORTER_ACTIVITY" [color="#D7BDE2"];
"gene_BTK" -- "term_10_GOMF_PHOSPHATIDYLINOSITOL_3_4_5_TRISPHOSPHATE_BINDING" [color="#D7BDE2"];
"gene_CALML4" -- "term_8_GOMF_MYOSIN_BINDING" [color="#D7BDE2"];
"gene_CARD8" -- "term_2_GOMF_PATTERN_RECOGNITION_RECEPTOR_ACTIVITY" [color="#D7BDE2"];
"gene_CCR5" -- "term_7_GOMF_CYTOKINE_RECEPTOR_ACTIVITY" [color="#D7BDE2"];
"gene_CD36" -- "term_2_GOMF_PATTERN_RECOGNITION_RECEPTOR_ACTIVITY" [color="#D7BDE2"];
"gene_CD4" -- "term_7_GOMF_CYTOKINE_RECEPTOR_ACTIVITY" [color="#D7BDE2"];
"gene_CLEC4E" -- "term_2_GOMF_PATTERN_RECOGNITION_RECEPTOR_ACTIVITY" [color="#D7BDE2"];
"gene_CORO1A" -- "term_8_GOMF_MYOSIN_BINDING" [color="#D7BDE2"];
"gene_CSF2RB" -- "term_7_GOMF_CYTOKINE_RECEPTOR_ACTIVITY" [color="#D7BDE2"];
"gene_CSF3R" -- "term_7_GOMF_CYTOKINE_RECEPTOR_ACTIVITY" [color="#D7BDE2"];
"gene_CX3CR1" -- "term_7_GOMF_CYTOKINE_RECEPTOR_ACTIVITY" [color="#D7BDE2"];
"gene_GSN" -- "term_8_GOMF_MYOSIN_BINDING" [color="#D7BDE2"];
"gene_HCK" -- "term_6_GOMF_PHOSPHOPROTEIN_BINDING" [color="#D7BDE2"];
"gene_IFNAR1" -- "term_7_GOMF_CYTOKINE_RECEPTOR_ACTIVITY" [color="#D7BDE2"];
"gene_IFNGR2" -- "term_7_GOMF_CYTOKINE_RECEPTOR_ACTIVITY" [color="#D7BDE2"];
"gene_IL10RB" -- "term_7_GOMF_CYTOKINE_RECEPTOR_ACTIVITY" [color="#D7BDE2"];
"gene_IL17RA" -- "term_7_GOMF_CYTOKINE_RECEPTOR_ACTIVITY" [color="#D7BDE2"];
"gene_IQGAP1" -- "term_10_GOMF_PHOSPHATIDYLINOSITOL_3_4_5_TRISPHOSPHATE_BINDING" [color="#D7BDE2"];
"gene_KIR3DL1" -- "term_1_GOMF_INHIBITORY_MHC_CLASS_I_RECEPTOR_ACTIVITY" [color="#D7BDE2"];
"gene_LILRA1" -- "term_1_GOMF_INHIBITORY_MHC_CLASS_I_RECEPTOR_ACTIVITY" [color="#D7BDE2"];
"gene_LILRA2" -- "term_1_GOMF_INHIBITORY_MHC_CLASS_I_RECEPTOR_ACTIVITY" [color="#D7BDE2"];
"gene_LILRA3" -- "term_1_GOMF_INHIBITORY_MHC_CLASS_I_RECEPTOR_ACTIVITY" [color="#D7BDE2"];
"gene_LILRA6" -- "term_1_GOMF_INHIBITORY_MHC_CLASS_I_RECEPTOR_ACTIVITY" [color="#D7BDE2"];
"gene_LILRB1" -- "term_1_GOMF_INHIBITORY_MHC_CLASS_I_RECEPTOR_ACTIVITY" [color="#D7BDE2"];
"gene_LILRB2" -- "term_1_GOMF_INHIBITORY_MHC_CLASS_I_RECEPTOR_ACTIVITY" [color="#D7BDE2"];
"gene_LILRB3" -- "term_1_GOMF_INHIBITORY_MHC_CLASS_I_RECEPTOR_ACTIVITY" [color="#D7BDE2"];
"gene_M6PR" -- "term_9_GOMF_RETROMER_COMPLEX_BINDING" [color="#D7BDE2"];
"gene_MAPK1" -- "term_6_GOMF_PHOSPHOPROTEIN_BINDING" [color="#D7BDE2"];
"gene_NCF1" -- "term_3_GOMF_SUPEROXIDE_GENERATING_NADPH_OXIDASE_ACTIVATOR_ACTIVITY" [color="#D7BDE2"];
"gene_NCF1B" -- "term_3_GOMF_SUPEROXIDE_GENERATING_NADPH_OXIDASE_ACTIVATOR_ACTIVITY" [color="#D7BDE2"];
"gene_NCF1C" -- "term_3_GOMF_SUPEROXIDE_GENERATING_NADPH_OXIDASE_ACTIVATOR_ACTIVITY" [color="#D7BDE2"];
"gene_NCF2" -- "term_3_GOMF_SUPEROXIDE_GENERATING_NADPH_OXIDASE_ACTIVATOR_ACTIVITY" [color="#D7BDE2"];
"gene_NCF4" -- "term_3_GOMF_SUPEROXIDE_GENERATING_NADPH_OXIDASE_ACTIVATOR_ACTIVITY" [color="#D7BDE2"];
"gene_NOD2" -- "term_2_GOMF_PATTERN_RECOGNITION_RECEPTOR_ACTIVITY" [color="#D7BDE2"];
"gene_PTAFR" -- "term_2_GOMF_PATTERN_RECOGNITION_RECEPTOR_ACTIVITY" [color="#D7BDE2"];
"gene_PTPN6" -- "term_6_GOMF_PHOSPHOPROTEIN_BINDING" [color="#D7BDE2"];
"gene_PYCARD" -- "term_2_GOMF_PATTERN_RECOGNITION_RECEPTOR_ACTIVITY" [color="#D7BDE2"];
"gene_RAB10" -- "term_8_GOMF_MYOSIN_BINDING" [color="#D7BDE2"];
"gene_RAB27A" -- "term_8_GOMF_MYOSIN_BINDING" [color="#D7BDE2"];
"gene_RAB7A" -- "term_9_GOMF_RETROMER_COMPLEX_BINDING" [color="#D7BDE2"];
"gene_RAB8A" -- "term_8_GOMF_MYOSIN_BINDING" [color="#D7BDE2"];
"gene_RIGI" -- "term_2_GOMF_PATTERN_RECOGNITION_RECEPTOR_ACTIVITY" [color="#D7BDE2"];
"gene_SH3BP2" -- "term_6_GOMF_PHOSPHOPROTEIN_BINDING" [color="#D7BDE2"];
"gene_SIGLEC10" -- "term_4_GOMF_SIALIC_ACID_BINDING" [color="#D7BDE2"];
"gene_SIGLEC12" -- "term_4_GOMF_SIALIC_ACID_BINDING" [color="#D7BDE2"];
"gene_SIGLEC5" -- "term_4_GOMF_SIALIC_ACID_BINDING" [color="#D7BDE2"];
"gene_SIGLEC7" -- "term_4_GOMF_SIALIC_ACID_BINDING" [color="#D7BDE2"];
"gene_SORT1" -- "term_9_GOMF_RETROMER_COMPLEX_BINDING" [color="#D7BDE2"];
"gene_ST8SIA4" -- "term_4_GOMF_SIALIC_ACID_BINDING" [color="#D7BDE2"];
"gene_SYK" -- "term_6_GOMF_PHOSPHOPROTEIN_BINDING" [color="#D7BDE2"];
"gene_TCIRG1" -- "term_5_GOMF_ATPASE_COUPLED_ION_TRANSMEMBRANE_TRANSPORTER_ACTIVITY" [color="#D7BDE2"];
"gene_TLR5" -- "term_2_GOMF_PATTERN_RECOGNITION_RECEPTOR_ACTIVITY" [color="#D7BDE2"];
"gene_TRIOBP" -- "term_8_GOMF_MYOSIN_BINDING" [color="#D7BDE2"];
"gene_UBASH3B" -- "term_6_GOMF_PHOSPHOPROTEIN_BINDING" [color="#D7BDE2"];
"gene_VAV1" -- "term_6_GOMF_PHOSPHOPROTEIN_BINDING" [color="#D7BDE2"];
"gene_YWHAE" -- "term_6_GOMF_PHOSPHOPROTEIN_BINDING" [color="#D7BDE2"];
"gene_C9ORF72" [label="C9ORF72", type="gene", width=0.7, height=0.7];
"gene_DDB2" [label="DDB2", type="gene", width=0.7, height=0.7];
"gene_HYCC1" [label="HYCC1", type="gene", width=0.7, height=0.7];
"gene_KCNJ2" [label="KCNJ2", type="gene", width=0.7, height=0.7];
"gene_PEPD" [label="PEPD", type="gene", width=0.7, height=0.7];
"gene_PHKA2" [label="PHKA2", type="gene", width=0.7, height=0.7];
"gene_PLPBP" [label="PLPBP", type="gene", width=0.7, height=0.7];
"gene_RNF31" [label="RNF31", type="gene", width=0.7, height=0.7];
"gene_SASH1" [label="SASH1", type="gene", width=0.7, height=0.7];
"gene_SEMA4A" [label="SEMA4A", type="gene", width=0.7, height=0.7];
"gene_SIL1" [label="SIL1", type="gene", width=0.7, height=0.7];
"gene_TAF4" [label="TAF4", type="gene", width=0.7, height=0.7];
"gene_TET2" [label="TET2", type="gene", width=0.7, height=0.7];
"gene_TMEM107" [label="TMEM107", type="gene", width=0.7, height=0.7];
"gene_TMEM218" [label="TMEM218", type="gene", width=0.7, height=0.7];
"gene_TYMP" [label="TYMP", type="gene", width=0.7, height=0.7];
"gene_USB1" [label="USB1", type="gene", width=0.7, height=0.7];
"term_10_HP_MACULAR_DEGENERATION" [label="HP: MACULAR DEGENERATION", style=filled, fillcolor="#F7DC6F", fontcolor="white", type="term", width=2, height=2, library="C5: Human Phenotype Ontology"];
"term_1_HP_SPLENOMEGALY" [label="HP: SPLENOMEGALY", style=filled, fillcolor="#F7DC6F", fontcolor="white", type="term", width=2, height=2, library="C5: Human Phenotype Ontology"];
"term_2_HP_AGITATION" [label="HP: AGITATION", style=filled, fillcolor="#F7DC6F", fontcolor="white", type="term", width=2, height=2, library="C5: Human Phenotype Ontology"];
"term_3_HP_MUSCLE_FLACCIDITY" [label="HP: MUSCLE FLACCIDITY", style=filled, fillcolor="#F7DC6F", fontcolor="white", type="term", width=2, height=2, library="C5: Human Phenotype Ontology"];
"term_4_HP_ABNORMAL_MOTOR_NERVE_CONDUCTION_VELOCITY" [label="HP: ABNORMAL MOTOR NERVE CONDUCTION VELOCITY", style=filled, fillcolor="#F7DC6F", fontcolor="white", type="term", width=2, height=2, library="C5: Human Phenotype Ontology"];
"term_5_HP_CUTANEOUS_ABSCESS" [label="HP: CUTANEOUS ABSCESS", style=filled, fillcolor="#F7DC6F", fontcolor="white", type="term", width=2, height=2, library="C5: Human Phenotype Ontology"];
"term_6_HP_METABOLIC_ACIDOSIS" [label="HP: METABOLIC ACIDOSIS", style=filled, fillcolor="#F7DC6F", fontcolor="white", type="term", width=2, height=2, library="C5: Human Phenotype Ontology"];
"term_7_HP_APRAXIA" [label="HP: APRAXIA", style=filled, fillcolor="#F7DC6F", fontcolor="white", type="term", width=2, height=2, library="C5: Human Phenotype Ontology"];
"term_8_HP_ABNORMAL_MORPHOLOGY_OF_THE_CONJUNCTIVAL_VASCULATURE" [label="HP: ABNORMAL MORPHOLOGY OF THE CONJUNCTIVAL VASCULATURE", style=filled, fillcolor="#F7DC6F", fontcolor="white", type="term", width=2, height=2, library="C5: Human Phenotype Ontology"];
"term_9_HP_ABNORMAL_PLATELET_VOLUME" [label="HP: ABNORMAL PLATELET VOLUME", style=filled, fillcolor="#F7DC6F", fontcolor="white", type="term", width=2, height=2, library="C5: Human Phenotype Ontology"];
"gene_ABCA7" -- "term_2_HP_AGITATION" [color="#F7DC6F"];
"gene_ADA2" -- "term_1_HP_SPLENOMEGALY" [color="#F7DC6F"];
"gene_APTX" -- "term_7_HP_APRAXIA" [color="#F7DC6F"];
"gene_ARPC1B" -- "term_9_HP_ABNORMAL_PLATELET_VOLUME" [color="#F7DC6F"];
"gene_ATP6AP1" -- "term_1_HP_SPLENOMEGALY" [color="#F7DC6F"];
"gene_ATP6V0A1" -- "term_2_HP_AGITATION" [color="#F7DC6F"];
"gene_ATP6V1B2" -- "term_1_HP_SPLENOMEGALY" [color="#F7DC6F"];
"gene_BEST1" -- "term_10_HP_MACULAR_DEGENERATION" [color="#F7DC6F"];
"gene_BTK" -- "term_5_HP_CUTANEOUS_ABSCESS" [color="#F7DC6F"];
"gene_C9ORF72" -- "term_7_HP_APRAXIA" [color="#F7DC6F"];
"gene_CASP10" -- "term_1_HP_SPLENOMEGALY" [color="#F7DC6F"];
"gene_CD36" -- "term_9_HP_ABNORMAL_PLATELET_VOLUME" [color="#F7DC6F"];
"gene_CSF1R" -- "term_7_HP_APRAXIA" [color="#F7DC6F"];
"gene_CSF3R" -- "term_1_HP_SPLENOMEGALY" [color="#F7DC6F"];
"gene_DDB2" -- "term_8_HP_ABNORMAL_MORPHOLOGY_OF_THE_CONJUNCTIVAL_VASCULATURE" [color="#F7DC6F"];
"gene_FGD4" -- "term_4_HP_ABNORMAL_MOTOR_NERVE_CONDUCTION_VELOCITY" [color="#F7DC6F"];
"gene_GAA" -- "term_1_HP_SPLENOMEGALY" [color="#F7DC6F"];
"gene_GBA1" -- "term_1_HP_SPLENOMEGALY" [color="#F7DC6F"];
"gene_GK" -- "term_6_HP_METABOLIC_ACIDOSIS" [color="#F7DC6F"];
"gene_GNS" -- "term_1_HP_SPLENOMEGALY" [color="#F7DC6F"];
"gene_GRN" -- "term_2_HP_AGITATION" [color="#F7DC6F"];
"gene_HCK" -- "term_1_HP_SPLENOMEGALY" [color="#F7DC6F"];
"gene_HK1" -- "term_1_HP_SPLENOMEGALY" [color="#F7DC6F"];
"gene_HMBS" -- "term_1_HP_SPLENOMEGALY" [color="#F7DC6F"];
"gene_HYCC1" -- "term_4_HP_ABNORMAL_MOTOR_NERVE_CONDUCTION_VELOCITY" [color="#F7DC6F"];
"gene_IL10RB" -- "term_5_HP_CUTANEOUS_ABSCESS" [color="#F7DC6F"];
"gene_IL17RA" -- "term_5_HP_CUTANEOUS_ABSCESS" [color="#F7DC6F"];
"gene_KCNE3" -- "term_3_HP_MUSCLE_FLACCIDITY" [color="#F7DC6F"];
"gene_KCNJ2" -- "term_3_HP_MUSCLE_FLACCIDITY" [color="#F7DC6F"];
"gene_LAMP2" -- "term_3_HP_MUSCLE_FLACCIDITY" [color="#F7DC6F"];
"gene_LRRK2" -- "term_2_HP_AGITATION" [color="#F7DC6F"];
"gene_LYST" -- "term_1_HP_SPLENOMEGALY" [color="#F7DC6F"];
"gene_MAPK1" -- "term_7_HP_APRAXIA" [color="#F7DC6F"];
"gene_MAPKAPK3" -- "term_10_HP_MACULAR_DEGENERATION" [color="#F7DC6F"];
"gene_MCCC2" -- "term_6_HP_METABOLIC_ACIDOSIS" [color="#F7DC6F"];
"gene_MICU1" -- "term_1_HP_SPLENOMEGALY" [color="#F7DC6F"];
"gene_MOGS" -- "term_1_HP_SPLENOMEGALY" [color="#F7DC6F"];
"gene_MPEG1" -- "term_5_HP_CUTANEOUS_ABSCESS" [color="#F7DC6F"];
"gene_NAGA" -- "term_8_HP_ABNORMAL_MORPHOLOGY_OF_THE_CONJUNCTIVAL_VASCULATURE" [color="#F7DC6F"];
"gene_NCF1" -- "term_1_HP_SPLENOMEGALY" [color="#F7DC6F"];
"gene_NCF2" -- "term_1_HP_SPLENOMEGALY" [color="#F7DC6F"];
"gene_NCF4" -- "term_1_HP_SPLENOMEGALY" [color="#F7DC6F"];
"gene_NLRP12" -- "term_1_HP_SPLENOMEGALY" [color="#F7DC6F"];
"gene_NOD2" -- "term_1_HP_SPLENOMEGALY" [color="#F7DC6F"];
"gene_OGDH" -- "term_6_HP_METABOLIC_ACIDOSIS" [color="#F7DC6F"];
"gene_OSTM1" -- "term_1_HP_SPLENOMEGALY" [color="#F7DC6F"];
"gene_PANK2" -- "term_7_HP_APRAXIA" [color="#F7DC6F"];
"gene_PEPD" -- "term_1_HP_SPLENOMEGALY" [color="#F7DC6F"];
"gene_PHKA2" -- "term_1_HP_SPLENOMEGALY" [color="#F7DC6F"];
"gene_PLPBP" -- "term_6_HP_METABOLIC_ACIDOSIS" [color="#F7DC6F"];

"gene_PRDX1" -- "term_6_HP_METABOLIC_ACIDOSIS" [color="#F7DC6F"];
"gene_PRKAR1A" -- "term_2_HP_AGITATION" [color="#F7DC6F"];
"gene_PRKCD" -- "term_1_HP_SPLENOMEGALY" [color="#F7DC6F"];
"gene_PSEN1" -- "term_2_HP_AGITATION" [color="#F7DC6F"];
"gene_PTPRJ" -- "term_9_HP_ABNORMAL_PLATELET_VOLUME" [color="#F7DC6F"];
"gene_RAB27A" -- "term_1_HP_SPLENOMEGALY" [color="#F7DC6F"];
"gene_RAB7A" -- "term_4_HP_ABNORMAL_MOTOR_NERVE_CONDUCTION_VELOCITY" [color="#F7DC6F"];
"gene_RNF135" -- "term_7_HP_APRAXIA" [color="#F7DC6F"];
"gene_RNF31" -- "term_1_HP_SPLENOMEGALY" [color="#F7DC6F"];
"gene_RRAGD" -- "term_6_HP_METABOLIC_ACIDOSIS" [color="#F7DC6F"];
"gene_SASH1" -- "term_8_HP_ABNORMAL_MORPHOLOGY_OF_THE_CONJUNCTIVAL_VASCULATURE" [color="#F7DC6F"];
"gene_SEC23B" -- "term_1_HP_SPLENOMEGALY" [color="#F7DC6F"];
"gene_SEMA4A" -- "term_10_HP_MACULAR_DEGENERATION" [color="#F7DC6F"];
"gene_SERPINA1" -- "term_1_HP_SPLENOMEGALY" [color="#F7DC6F"];
"gene_SH3BP2" -- "term_10_HP_MACULAR_DEGENERATION" [color="#F7DC6F"];
"gene_SIL1" -- "term_3_HP_MUSCLE_FLACCIDITY" [color="#F7DC6F"];
"gene_SLC12A6" -- "term_4_HP_ABNORMAL_MOTOR_NERVE_CONDUCTION_VELOCITY" [color="#F7DC6F"];
"gene_SLC6A6" -- "term_10_HP_MACULAR_DEGENERATION" [color="#F7DC6F"];
"gene_SMC1A" -- "term_2_HP_AGITATION" [color="#F7DC6F"];
"gene_SPTLC2" -- "term_4_HP_ABNORMAL_MOTOR_NERVE_CONDUCTION_VELOCITY" [color="#F7DC6F"];
"gene_STX3" -- "term_6_HP_METABOLIC_ACIDOSIS" [color="#F7DC6F"];
"gene_SYK" -- "term_1_HP_SPLENOMEGALY" [color="#F7DC6F"];
"gene_TAF4" -- "term_7_HP_APRAXIA" [color="#F7DC6F"];
"gene_TBXAS1" -- "term_1_HP_SPLENOMEGALY" [color="#F7DC6F"];
"gene_TCIRG1" -- "term_1_HP_SPLENOMEGALY" [color="#F7DC6F"];
"gene_TET2" -- "term_1_HP_SPLENOMEGALY" [color="#F7DC6F"];
"gene_TMEM107" -- "term_7_HP_APRAXIA" [color="#F7DC6F"];
"gene_TMEM218" -- "term_7_HP_APRAXIA" [color="#F7DC6F"];
"gene_TNFRSF1A" -- "term_1_HP_SPLENOMEGALY" [color="#F7DC6F"];
"gene_TPI1" -- "term_1_HP_SPLENOMEGALY" [color="#F7DC6F"];
"gene_TYMP" -- "term_4_HP_ABNORMAL_MOTOR_NERVE_CONDUCTION_VELOCITY" [color="#F7DC6F"];
"gene_USB1" -- "term_1_HP_SPLENOMEGALY" [color="#F7DC6F"];
"gene_VIPAS39" -- "term_6_HP_METABOLIC_ACIDOSIS" [color="#F7DC6F"];
"gene_VPS35" -- "term_2_HP_AGITATION" [color="#F7DC6F"];
}

**NETWORK INTERPRETATION GUIDE:**

**Node Types:**
- **Gene nodes** (type="gene"): Individual genes from the input gene set
- **Term nodes** (type="term"): Enriched biological pathways/processes/gene sets
- **Library Source**: Each term node includes a "library" attribute indicating the source database

**Library Sources and Their Biological Context:**
- **H: Hallmark gene sets**: Curated gene sets representing well-defined biological states or processes
- **C2: BioCarta**: Canonical pathways from BioCarta database
- **C2: KEGG MEDICUS**: Metabolic and signaling pathways from KEGG
- **C2: Pathway Interaction Database**: Curated human signaling pathways
- **C2: Reactome Pathways**: Expert-curated biological pathways
- **C2: WikiPathways**: Community-curated biological pathways
- **C5: Gene Ontology: Biological Process**: Biological processes from Gene Ontology
- **C5: Gene Ontology: Cellular Component**: Cellular components from Gene Ontology
- **C5: Gene Ontology: Molecular Function**: Molecular functions from Gene Ontology
- **C5: Human Phenotype Ontology**: Human phenotypes and diseases
- **Protein Interaction**: Protein-protein interaction networks

**Edge Connections:**
- Each edge (gene -- term) represents a gene's membership in that biological term
- More connections indicate genes that are part of multiple enriched processes
- The network topology shows which genes are central to the biological response

**Iteration Information:**
- Term nodes are labeled with iteration numbers (term_1_, term_2_, etc.)
- Earlier iterations (lower numbers) represent the most statistically significant enrichments
- Ignore iteration number for biological interpretation, instead focus on network hubs densely connected for interesting biological insights

**ANALYSIS REQUEST:**

Please analyze this network and provide:

1. **Key Biological Insights:**
   - What are the most significant biological processes/pathways identified?
   - Which genes appear to be central to the biological response?

2. **Network Topology Analysis:**
   - Which genes are "hub" genes (connected to multiple terms)?
   - Are there distinct clusters or modules in the network?
   - What does the connectivity pattern suggest about biological organization?

3. **Biological Hypothesis:**
   - Based on the network structure, what biological hypothesis can you generate?
   - How do the different library sources support or complement each other?
   - Try to formulate one hypothesis based on the most central terms and how they possibly related to one another.

4. **Estimated Experimental Context**
   - Can you hypothesize on what was the experiment that generated this result?

**RESPONSE STRUCTURE:**
- **Executive Summary** (2-3 sentences)
- **Key Biological Processes Identified** (by library source)

- **Network Topology Insights**
- **Proposed Biological Hypothesis**
- **Estimated Experimental Context**

**IMPORTANT NOTES:**
- Focus on biological interpretation, not statistical significance or iteration number
- Consider the functional relationships between connected genes and terms
- Look for unexpected connections that might reveal novel or unexpected biology
- Consider the broader biological context and literature
- Note that protein interaction library could shed light on physical interaction between genes while terms indicate functional interaction
- Different library sources may provide complementary biological insights

---

**Executive Summary**

Across essentially every library, the network converges on **innate immune / phagocyte biology**—especially **phagocytosis, FcγR signaling, neutrophil degranulation, complement activation, and ROS/NADPH-oxidase activity**—with a secondary layer of **inflammatory cytokine signaling (IL6–JAK–STAT3, IL-17)**. A smaller but coherent metabolic/trafficking component (lysosome/vacuolar acidification, glycosylation, sphingolipid metabolism, mTORC1/CoA/TCA) suggests an immune-cell activation state with major **endolysosomal remodeling**.

**Key Biological Processes Identified (by library source)**

**H: Hallmark Gene Sets**

**1) Complement (HALLMARK: COMPLEMENT)**
- Genes mapped here include: **CR1, CASP1, CASP10, CD36, CTSD, LAMP2, LRP1, SERPINA1, TIMP2, PRKCD, IRF2, MSRB1, PPP2CB**, etc.
- Interpretation: this reflects **myeloid innate immune activation**, complement engagement/opsonization, and inflammatory effector activity.

**2) mTORC1 signaling (HALLMARK: MTORC1 SIGNALING)**
- Includes metabolic/housekeeping + immune-relevant genes: **ACLY, IDH1, HMBS, TPI1, PRDX1, GLRX**, and others (also **CORO1A, SKAP2, IFI30**).
- Interpretation: suggests **metabolic rewiring** consistent with activated immune cells (increased biosynthesis, redox control, glycolysis/TCA coupling).

**3) IL6–JAK–STAT3 signaling (HALLMARK: IL6 JAK STAT3 SIGNALING)**
- Includes **CSF2RB, CSF3R, IFNAR1, IFNGR2, IL10RB, IL17RA, STAM2, TNFRSF1A**.
- Interpretation: a broad **inflammatory cytokine receptor/signaling state**, spanning interferon, IL-10 family receptor components, TNF receptor, and colony-stimulating factor receptors.

**C2: KEGG Legacy**

**Lysosome / endolysosomal machinery (KEGG: LYSOSOME)**
- Strong set: **ATP6AP1, ATP6V0A1, ATP6V0D1, CLTA, CTSD, GAA, GBA1, GNS, LAMP2, M6PR, NAGA, SLC11A1, SORT1, TCIRG1, ENTPD4**
- Interpretation: **lysosomal biogenesis, acidification, cargo trafficking, antigen processing**—typical of phagocytes (macrophages/microglia/neutrophils).

**FcγR-mediated phagocytosis (KEGG: FC GAMMA R MEDIATED PHAGOCYTOSIS)**
- Includes **SYK, VAV1, PAK1, PRKCD, HCK, ARPC1B, MAPK1, NCF1, GSN**
- Interpretation: canonical **immune receptor → actin remodeling → engulfment**, with downstream oxidative burst capability.

**Apoptosis (KEGG: APOPTOSIS)**
- Includes **APAF1, CASP10, DFFA, TNFRSF1A, PRKACA, PRKAR1A, CSF2RB**
- Interpretation: a "cell fate / inflammatory death" axis that often co-occurs with immune activation (and also with inflammasome/caspase wiring).

**Amino sugar & nucleotide sugar metabolism**
- **HK1, HK3, NAGK, NPL, FPGT**
- Interpretation: ties to **glycosylation substrate supply**, immune activation, and lysosomal/endosomal function.

**Pantothenate and CoA biosynthesis**
- **PANK2, PPCDC, VNN2**

- Interpretation: supports **lipid metabolism and mitochondrial/energetic remodeling**.

---

**C2: KEGG MEDICUS**

**PRNP–PI3K–NOX2 signaling pathway**

- **NCF1/2/4, PRKCD, MAPK1, CREB5**
- Interpretation: this is a very "oxidative burst"–flavored module: **NOX2/NADPH oxidase activation**, downstream signaling, and transcriptional output.

---

**C2: Pathway Interaction Database (PID)**

**PDGFRB pathway**

- **ACTR3, ARAP1, ARPC1B, CSK, HCK, IQGAP1, MAP2K4, MAPK1, PAK1, PRKCD, PTPRJ, YWHAE, LRP1**
- Interpretation: not necessarily "PDGFRB-driven signaling" per se, but a signature rich in **actin dynamics + SRC-family kinase wiring + MAPK**, which also overlaps strongly with phagocytosis/migration machinery.

**Caspase pathway / Retinoic acid pathway**

- **APAF1, CASP1, CASP10, DFFA, TNFRSF1A** (caspase)
- **RXRA, VDR, RBP1, PRKACA** (retinoic acid)
- Interpretation: a second axis: **inflammatory/cell death signaling** plus **nuclear receptor differentiation/immune modulation** (RA/VDR).

---

**C2: Reactome Pathways**

This is where the "story" becomes most integrated.

**Neutrophil degranulation (very dominant)**

- A *huge* number of genes land here: **FPR2, ITGAX, CD36, CR1, CFD, CTSD, GRN, LAMP2, PRKCD, PTPN6, PYCARD, RAB7A, RAB27A, RAB10, SERPINA1, etc.**
- Interpretation: strongly supports **activated innate immune cells with granule/vesicle trafficking and antimicrobial effector programs**.

**ROS and RNS production in phagocytes**

- **NCF1/2/4, ATP6V1A/B2, ATP6V0D1**
- Interpretation: oxidative burst + phagosomal/lysosomal acidification as a coupled effector system.

**Interleukin signaling / IL-17 signaling / immunoregulatory interactions**

- **IL17RA, MAP2K3, MAP2K4, MAPKAPK3, RPS6KA1, PPP2CB, NOD2**
- plus immune inhibitory receptor and cell-cell interaction genes: **KIRs, LILRs, SIGLECs, SLAMF7, JAML**
- Interpretation: a strong **inflammatory immune communication** module that looks like a mixture of myeloid and NK/lymphoid regulatory components.

**NF-κB activation through FADD/RIP1 via caspase-8/10**

- **CASP10, RIGI (DDX58), RNF135**
- Interpretation: points to **innate sensing + NF-κB** activation and programmed inflammatory signaling.

**Sphingolipid metabolism**

- **GBA1, ORMDL2, PPM1L, SPTLC2, SPTSSA, M6PR**
- Interpretation: membrane lipid remodeling that often accompanies immune activation and vesicle trafficking.

**Asparagine N-linked glycosylation**

- **OS9, MOGS, KDELR1/2, SEC23IP, RAB1B, ARF3**
- Interpretation: ER/Golgi processing—often upregulated in secretory/immune activation states.

---

**C2: WikiPathways**

These add "cell-type flavor" and mechanistic narratives:

**TYROBP causal network in microglia + Microglia pathogen phagocytosis pathway**

- Includes classic myeloid/microglia phagocytosis signaling components: **SYK, VAV1, NCF genes, ARPC1B, PTPN6, SIGLEC7, ITGAX, APBB1IP, CAPG, RPS6KA1**
- Interpretation: this pushes the interpretation toward **microglia-like / macrophage-like phagocytes** (and the inclusion of TYROBP-associated wiring is very suggestive).

**NOD pathway / inflammasome-adjacent**

- **NOD2, CASP1, PYCARD, NAIP, NLRP12, CARD8**
- Interpretation: **innate sensing + inflammasome / inflammatory caspase machinery**.

**Cannabinoid receptor signaling**

- **DAGLB, ADCY7, PRKACA, PRKAR1A, MAPK1**
- Interpretation: could be a genuine signal (e.g., immune modulation through GPCR/cAMP), or a "shared signaling scaffold" enrichment because many immune pathways converge on MAPK/PKA/cAMP nodes.

---

**C5: GO Biological Process**

This library is very consistent with the above and helps separate modules:

**Phagocytosis**

- Massive membership: **SYK, VAV1, CD36, CSK, PAK1, LRP1, SLC11A1, NCF2/4, PTPRJ, PECAM1, FPR2, etc.**
- Interpretation: core functional program is **particle uptake / clearance / immune complex handling**.

**Cytokine-mediated signaling**

- **CCR5, CX3CR1, IFNAR1, IFNGR2, IL10RB, IL17RA, CSF1R/2RB/3R, TNFRSF1A**
- Interpretation: suggests **activated immune signaling environment** (chemokine receptor + IFN/TNF/IL17/CSFs).

**Vacuolar acidification**

- *ATP6 genes, TCIRG1, LAMP2, GRN, DMXL2**
- Interpretation: again, **phagolysosome maturation**.

**Golgi vesicle transport**

- **RAB10/1B/4B/8A, SEC23B, STX6, SNX1, SORT1, KDELR1/2**
- Interpretation: **vesicle trafficking / secretory pathway** reprogramming.

**Antibacterial innate immune response**

- Includes **MPEG1, NOD2, TFEB**
- Interpretation: not just uptake—also **microbicidal program + lysosome biogenesis (TFEB)**.

---

**C5: GO Cellular Component**

This provides subcellular structure context:

- **Phagocytic vesicle / vacuolar membrane / azurophil granule lumen**
- **Canonical inflammasome complex** (CASP1/PYCARD/NAIP/CARD8)
- **Membrane microdomain** (immune receptor platforms)
- Interpretation: immune receptors + inflammasome + phagosome + granules all in one picture → **professional phagocyte** signature.

---

**C5: GO Molecular Function**

This library highlights "mechanistic levers":

- **Superoxide-generating NADPH oxidase activator activity** (NCF1/2/4 + paralogs)
- **Pattern recognition receptor activity** (NOD2, CLEC4E, TLR5, CD36, etc.)
- **Cytokine receptor activity** (IL17RA, IFNAR1/IFNGR2, CSF receptors, CCR5/CX3CR1)
- **Phosphoprotein binding / PIP3 binding** (HCK, SYK, PTPN6, ARAPs, IQGAP1)
- Interpretation: coherent **PRR → signaling kinase → PI3K → cytoskeleton + NADPH oxidase** architecture.

---

**C5: Human Phenotype Ontology**

You should treat HPO here as "disease/clinical echo" rather than direct biology.

- **Splenomegaly** connects to many immune/lysosomal/hematologic genes (NCF genes, BTK, lysosomal genes, etc.) → consistent with immune dysregulation / myeloid activation / storage-like phenotypes.
- **Cutaneous abscess** connects to **BTK, IL10RB, IL17RA, MPEG1** → strongly consistent with **impaired bacterial defense / neutrophil function / IL-17 axis**.
- Interpretation: supports a theme of **innate immune dysfunction/activation**, sometimes seen in immunodeficiency/inflammatory syndromes.

---

**Network Topology Insights**

**1) Hub genes (multi-term connectors)**

A few genes clearly act as *bridges across many libraries/terms*:

**Core phagocyte signaling hubs**

- **PRKCD**: appears in FcγR phagocytosis, NOX2 signaling, PDGFRB pathway, complement-associated sets, neutrophil degranulation; classic immune signaling kinase.
- **MAPK1**: shows up in FcγR, PDGFRB, NOX2 signaling, cannabinoid receptor signaling, and broader inflammatory modules.
- **SYK** and **VAV1**: phagocytosis and immune receptor signaling hubs.

**Oxidative burst module hubs**

- **NCF1, NCF2, NCF4**: strongly repeated (ROS production, NOX2 signaling, phagocytosis, HPO immune phenotypes). These are among the strongest "mechanistic anchors" in the network.

**Innate sensing / inflammasome hubs**

- **CASP1, PYCARD, NOD2, NAIP, NLRP12, CARD8**: cluster into a PRR–inflammasome module, also linking to cytokine signaling and phagosome biology.

**Endolysosomal/acidification hubs**

- **ATP6V0/ATP6V1 family, TCIRG1, LAMP2, CTSD, GAA, GBA1, M6PR**: recur across KEGG lysosome, GO vacuolar membrane/acidification, Reactome degranulation and ROS.

**Immune receptor/regulation surface hubs**

- **CD36, CR1, ITGAX, CCR5, CX3CR1**, plus inhibitory receptors (**KIRs, LILRs, SIGLECs**) form a surface-receptor module.

If you wanted a short list of "most central" genes to interpret or test first, it would be:
**PRKCD, SYK, VAV1, MAPK1, NCF1/2/4, CASP1, PYCARD, NOD2, ATP6V0D1/ATP6V1A/B2, LAMP2, CTSD.**

---

**2) Distinct clusters/modules**

Even without doing formal community detection, the graph visually/functionally breaks into:

**Module A — Phagocytosis / FcγR / actin remodeling**

- **SYK–VAV1–PAK1–ARPC1B–ACTR3–IQGAP1–CSK–HCK**
- Links strongly to PID PDGFRB pathway, KEGG FcγR phagocytosis, GO phagocytosis, and microglia phagocytosis WP terms.

**Module B — Phagosome maturation / lysosome / acidification**

- *ATP6 + TCIRG1 + LAMP2 + CTSD + M6PR + GAA/GBA1/GNS**
- Links to KEGG lysosome, GO vacuolar acidification, Reactome neutrophil degranulation.

**Module C — Oxidative burst / NOX2**

- **NCF1/2/4** + upstream signaling (**PRKCD, MAPK1**)
- Links to Reactome ROS production, KEGG MEDICUS NOX2 pathway, GO MF NADPH oxidase activator.

**Module D — Innate sensing / inflammasome / inflammatory caspase wiring**

- **NOD2 + CASP1 + PYCARD + NAIP + NLRP12 + CARD8**
- Links to GO CC inflammasome complex, WP NOD pathway, and inflammatory signaling terms.

**Module E — Cytokine and immune communication**

- **IL17RA, IFNAR1, IFNGR2, IL10RB, CSF receptors, TNFRSF1A, CCR5, CX3CR1**
- Links to Hallmark IL6/JAK/STAT3, GO cytokine signaling, Reactome interleukin signaling, plus immune regulatory interactions.

**Module F — Metabolic / regulatory overlay**

- **ACLY, IDH1, TPI1, PRDX1, GLRX, HMBS** and lipid-related (SPTLC2, ORMDL2)
- Links to mTORC1 signaling and sphingolipid metabolism; likely reflecting the metabolic state of activated innate cells.

---

**3) What the connectivity pattern suggests**

- The network is not dominated by a single pathway; instead, it looks like a **coordinated innate immune program** with:
    1. **recognition** (PRRs, complement receptors, Fc receptors),
    2. **engulfment** (actin remodeling),
    3. **phagosome maturation** (acidification + lysosome fusion),
    4. **killing** (oxidative burst, granule release), and
    5. **communication** (cytokine/chemokine signaling, IL-17 axis).
- The fact that the same signaling skeleton is "named differently" across libraries (FcγR phagocytosis, PDGFRB pathway, microglia TYROBP network) suggests the biology is robust: **multiple ontologies recognize the same core cellular state**.

---

**Proposed Biological Hypothesis**

**High-level hypothesis**

**The gene list reflects an activated professional phagocyte state (likely macrophage/microglia/neutrophil-enriched), characterized by FcγR/complement-mediated phagocytosis with strong phagolysosome maturation and NADPH-oxidase oxidative burst, accompanied by IL-17 / cytokine-driven inflammatory signaling and metabolic rewiring (mTORC1, lipid and glycosylation pathways).**

**Mechanistic hypothesis (testable)**

A particularly testable mechanistic thread is:

### PRKCD–SYK–VAV1 signaling coordinates FcγR/complement receptor-driven cytoskeletal remodeling and NOX2 assembly (NCF1/2/4), coupling phagocytosis to oxidative burst; simultaneously, NOD2/inflammasome components (CASP1/PYCARD/NAIP/NLRP12) amplify cytokine output (IL-17/IFN/TNF axis), pushing the system toward an antimicrobial, pro-inflammatory state.

This is supported consistently across:

- **KEGG FcγR phagocytosis + Reactome neutrophil degranulation/ROS** (function + effector)
- **GO CC phagocytic vesicle + inflammasome complex** (subcellular organization)
- **GO MF NADPH oxidase activator + PRR activity** (molecular mechanism)
- **WikiPathways TYROBP microglia network** (cell-type/program context)
- **Hallmark complement + IL6/JAK/STAT3 + mTORC1** (state-level signatures)

**Why the library sources complement each other**

- **Hallmark** captures *state-level themes* (complement, inflammatory cytokines, mTORC1 metabolic activation).
- **KEGG/PID/Reactome** provide *mechanistic wiring diagrams* (FcγR→SYK→VAV1; caspases; ROS).
- **GO** provides *cellular localization and functional granularity* (phagocytic vesicle, vacuolar membrane, NADPH oxidase activator).
- **WikiPathways** adds *contextual narratives* (microglia TYROBP network) that can hint at the **dominant cell type** driving the signal.
- **HPO** provides a *clinical echo* consistent with innate immune dysfunction/activation (abscesses, splenomegaly).

---

**Estimated Experimental Context (what experiment likely produced this)**

Based on the dominance of **phagocytosis, neutrophil degranulation, ROS production**, and **microglia TYROBP network**, the most plausible contexts are:

**Most likely**

**Bulk RNA-seq (or differential gene expression) from an inflamed tissue or immune-cell-enriched sample**, where the signature is driven by:

- **Activated myeloid cells** (macrophages/monocytes) and/or
- **Microglia-like cells** (if the tissue is CNS) and/or
- **Neutrophil infiltration** (if acute inflammation/infection).

Common real-world scenarios that generate this pattern:

- inflamed mucosa (e.g., IBD-like),
- infected tissue,
- neuroinflammation (microglial activation),
- immune-complex–rich inflammation (FcγR/complement),
- response to a bacterial stimulus (LPS, opsonized particles), or
- cytokine stimulation with IL-17/IFN/TNF combinations.

**Alternative but also plausible**

**Single-cell RNA-seq marker gene list** where the selected genes are enriched in a **myeloid cluster** (microglia/macrophage/neutrophil-like), producing strong phagocytosis/lysosome/ROS pathways when aggregated.

**One specific guess, given the WikiPathways microglia terms**

If this list came from brain tissue, the signature is extremely consistent with:

- **microglial activation / disease-associated microglia–like state**,
- potentially with complement and phagolysosomal remodeling.

---

If you want to push this further analytically, the *next best step* would be to compute (from the DOT) **gene degree** (number of connected term nodes) and list the **top 10 hub genes**, then also compute **term degree** to see which terms recruit the most genes per library—those two lists usually make the biological narrative almost trivial to prioritize for validation.

--

**New Prompt:**
The experiment that generated the list was a vaccination study. Healthy individuals were vaccinated and the experiment is a gene expression analysis of genes correlated with HAI response at day 28 by comparing expression of PBMC at day 28 and day 0. Explain the above analysis in this context and how it affects the analysis

**Executive Summary**

Placing this network in the context of an influenza (or similar) vaccination study with PBMC day28 vs day0 expression correlated with **HAI response** strongly reframes the biology: the network looks like a **coordinated innate immune + antigen handling program** (phagocytosis/lysosome/degranulation/complement) that is coupled to **cytokine/JAK–STAT and IL-17 signaling**, and it co-occurs with **metabolic rewiring (mTORC1, lipid/sphingolipid, amino-sugar metabolism)** needed to sustain activated immune cells. In short, the HAI correlates at day 28 appear to be **immune activation state + myeloid/APC involvement** rather than a purely B-cell transcriptional signature.

**Key Biological Processes Identified (interpreted as correlates of protective antibody response)**

**1) Hallmark (H) – "immune activation state + metabolic support"**

**HALLMARK: COMPLEMENT**

- Strongly points to **innate effector activity** and/or activation of complement-related gene programs in PBMC at day 28 (e.g., **CR1, CFD, SERPINA1**, CTSD, CASP1/10 also overlap with inflammatory programs).
- In vaccine correlates studies, complement signals often reflect **monocyte / neutrophil-like transcriptional activity** in PBMC preparations (even if neutrophils are excluded, monocytes can express complement genes), and can track with antigen clearance and immune complex handling.

**HALLMARK: IL6–JAK–STAT3 SIGNALING**

- A classic module for **post-vaccination inflammatory cytokine signaling** and immune-cell differentiation.
- Here it connects to receptors and signaling nodes (**IFNAR1, IFNGR2, IL10RB, IL17RA, TNFRSF1A, CSF2RB, CSF3R**) suggesting a broad cytokine responsiveness state that may facilitate effective T–B help, antigen presentation, and germinal center support (even if not directly measured in PBMC).

**HALLMARK: MTORC1 SIGNALING**

- This is the metabolic/translation axis of immune activation. Its appearance at day 28 implies that individuals with strong HAI responses maintain (or re-activate) metabolic programs consistent with **trained innate-like activation**, sustained antigen handling, or ongoing immune remodeling.
- Connected genes include metabolic enzymes and housekeeping factors (**ACLY, IDH1, TPI1, HMBS**) as well as immune-associated genes (e.g., IFI30).

**Vaccination-context implication:**
This hallmark triad (Complement + IL6/JAK/STAT3 + mTORC1) reads as:

"Individuals with higher antibody titers show a stronger sustained systemic immune activation/metabolic program at day 28."

That's already a useful hypothesis because many vaccines show early innate signatures (days 1–7) as predictors of later antibody titers; seeing it at day 28 suggests **persistent or secondary immune activation** in responders.

**2) KEGG Legacy / KEGG Medicus – "phagocytic machinery + apoptosis + ROS pathways"**

**KEGG: Lysosome** + **GOCC: Vacuolar membrane / phagocytic vesicle**

- Dense enrichment around lysosomal genes (**ATP6V0A1, ATP6V0D1, ATP6V1A/B2, LAMP2, CTSD, GAA, GBA1, TCIRG1, NAGA, GNS, M6PR, SORT1**).
- In PBMC, this almost always indicates **monocytes / dendritic cells** (APCs) and sometimes activated B cells (endosomal processing), but the dominance of lysosome/vacuolar acidification strongly points to **antigen processing and phagosomal maturation**.

**KEGG: Fc gamma R mediated phagocytosis**

- Adds the antibody-effector side: FcγR signaling + actin remodeling (genes include **SYK, VAV1, HCK, NCF1, MAPK1, PAK1, ARPC1B, GSN, PRKCD**).
- This is particularly meaningful for an HAI correlate at day 28: higher HAI responders likely have more immune complexes and/or more active FcγR signaling, or simply a stronger myeloid activation state.

**KEGG Medicus: PRNP–PI3K–NOX2 signaling pathway**

- Essentially a **NOX2/ROS production module** (NCF1/2/4 with PI3K/MAPK/PRKCD).
- This aligns perfectly with Reactome "ROS and RNS production in phagocytes".

**KEGG: Apoptosis + PID Caspase pathway**

- Includes **CASP10, CASP1, APAF1, DFFA**.
- In the vaccine context, apoptosis modules may reflect:

1. regulated turnover of activated immune cells,
2. inflammasome/caspase-1 activation (more inflammatory than apoptotic per se),
3. changes in PBMC composition (e.g., more activated monocytes in responders).

**Vaccination-context implication:**
These KEGG/PID patterns suggest that strong HAI responders show a day-28 PBMC signature enriched for **myeloid effector functions** (phagocytosis, ROS, lysosome) rather than a pure "plasmablast / antibody-secreting cell" signature (which often peaks around day 7). That may reflect:

- ongoing antigen/immune complex processing,
- "trained immunity" or enhanced innate responsiveness in responders,
- or simply the cellular composition differences in PBMC at day 28.

---

### 3) Reactome / WikiPathways – "neutrophil-like modules + IL-17 + microglial/innate networks"

**Reactome: Neutrophil degranulation**

- This is a common PBMC enrichment artifact/feature: it often appears in monocyte and NK modules because many granule/innate genes overlap.
- But when it's dominant *and* paired with FcγR phagocytosis + ROS production, it strongly indicates a **granulocyte-like / myeloid effector activation program**.

**Reactome: IL-17 signaling**

- Driven by **IL17RA, MAP2K3, MAP2K4, MAPKAPK3, PPP2CB, RPS6KA1, NOD2**.
- IL-17 signals in vaccination can represent:
    - Th17 polarization,
    - innate IL-17 axis activation,
    - mucosal-like inflammatory tone.
- Its co-occurrence with phagocytosis/ROS suggests a **pro-inflammatory innate module** that may enhance antigen presentation and B cell help indirectly.

**WikiPathways: TYROBP causal network / Microglia phagocytosis**

- These are innate immune networks heavily used in neuroinflammation, but the genes involved are broadly **myeloid innate signaling**.
- In PBMC vaccination data, they typically reflect **monocyte/macrophage-like activation**.

**Vaccination-context implication:**
Responders show a signature of **myeloid activation** with IL-17–associated inflammatory signaling. This might be a late "maintenance state" that correlates with durable HAI.

---

### 4) GO Biological Process / Cellular Component / Molecular Function – "mechanistic decomposition"

These GO terms help you interpret *how* the myeloid program is implemented:

- **GOBP: Phagocytosis**
- **GOBP: Vacuolar acidification** → classic antigen processing machinery
- **GOBP: Cytokine-mediated signaling pathway** → cytokine responsiveness
- **GOCC: Phagocytic vesicle / vacuolar membrane / azurophil granule lumen / ficolin-1 rich granule** → vesicle/granule identity typical of myeloid cells
- **GOMF: Superoxide-generating NADPH oxidase activator activity** (**NCF1/2/4**) → ROS production
- **GOMF: Pattern recognition receptor activity** (**NOD2, TLR5, CLEC4E**) → innate sensing
- **GOMF: Cytokine receptor activity** (**IFNAR1, IFNGR2, IL10RB, IL17RA, CCR5, CSF3R**) → responsiveness / recruitment
- **GOMF: Sialic acid binding** (**SIGLEC5/7/10/12, ST8SIA4**) → immunomodulation; potentially relevant for influenza (sialic acid biology), although these are mostly inhibitory receptors and glycosylation enzymes.

**Vaccination-context implication:**
GO analysis confirms the phenotype: **innate immune sensing + phagosome maturation + ROS + cytokine responsiveness**, consistent with responders having a more activated APC/myeloid landscape at day 28.

---

### Network Topology Insights (reinterpreted for PBMC day28 correlates)

**Hub genes (multi-library, cross-module connectors)**

The most "hub-like" genes here are those that repeatedly show up across libraries/terms:

- **NCF1 / NCF2 / NCF4**: connect FcγR phagocytosis ↔ ROS production ↔ innate effector modules. These look like a core "NOX2 respiratory burst" axis.
- **MAPK1 / PRKCD / SYK / VAV1 / PAK1 / HCK**: Fc receptor signaling + downstream activation.
- **ATP6V0A1 / ATP6V0D1 / ATP6V1A / ATP6V1B2 / LAMP2 / CTSD / M6PR / TCIRG1**: acidification/lysosome hubs.

- **NOD2 / CASP1 / PYCARD / CARD8**: innate sensing / inflammasome-ish axis.
- **IL17RA, IFNAR1, IFNGR2, IL10RB, TNFRSF1A, CCR5**: cytokine receptor hub layer that links inflammatory signaling to cellular response programs.
- **ACLY, IDH1, TPI1**: metabolic nodes that connect the immune module to "mTORC1 signaling".

**Vaccination interpretation:**
These hubs suggest that the HAI-linked day 28 signature is not one linear pathway, but a coordinated state combining:

1. **myeloid antigen handling** (lysosome/phagosome),
2. **activation signaling** (FcγR/SYK/MAPK),
3. **effector chemistry** (ROS),
4. **cytokine responsiveness**,
5. **metabolic support** (mTORC1).

**Modules / clusters**

Even without explicit community detection, the network visually decomposes into a few major modules:

1. **Phagocytosis / FcγR / ROS module**
   - SYK/VAV1/NCF1/2/4/HCK/MAPK1/PAK1/ARPC1B
2. **Lysosome / vacuolar acidification / vesicle trafficking module**
   - ATP6V* genes, LAMP2, CTSD, TCIRG1, M6PR, SORT1, RABs
3. **Cytokine / IL-17 / IFN / IL-6 axis module**
   - IL17RA, IFNAR1, IFNGR2, IL10RB, TNFRSF1A, CSF3R
4. **Inflammasome / caspase module**
   - CASP1, PYCARD, CARD8 (plus apoptotic components)
5. **Metabolic rewiring module**
   - ACLY, IDH1, TPI1 and other enzymes linked to MTORC1/Hallmark metabolic programs

**Vaccination interpretation:**
This is what you'd expect if the "signal" is driven by **cell-state + cell-type composition** differences at day 28: responders have more (or more activated) myeloid/APC populations and a stronger inflammatory/metabolic tone.

**What the connectivity pattern suggests about biological organization**

- The fact that many genes connect to **innate/myeloid terms across multiple libraries** suggests a robust, coherent biological signal (not a single database artifact).
- The **iterative enrichment** approach amplifies this interpretation: once "top" genes supporting myeloid programs are removed in each library, subsequent iterations still find related myeloid terms (e.g., phagocytosis ↔ lysosome ↔ ROS ↔ degranulation). That persistence means the gene list contains **multiple partially overlapping layers** of innate activation.

---

**Proposed Biological Hypothesis (now explicitly tied to HAI and day 28)**

**Hypothesis**

**High HAI responders exhibit a persistent day-28 PBMC transcriptional program dominated by activated monocyte/APC effector functions—phagocytosis, lysosomal antigen processing, and NOX2-dependent ROS production—coupled to cytokine responsiveness (IL-6/JAK-STAT3, IFN, IL-17) and metabolic activation (mTORC1).**

This program may either:

1. reflect **ongoing immune complex handling and antigen processing**, potentially linked to sustained germinal center activity and affinity maturation, or
2. represent a **trained immunity-like state**, where responders maintain heightened innate responsiveness that supports stronger humoral immunity.

**How the different libraries support/complement each other**

- **Hallmark** gives the "state labels": complement, IL6/JAK/STAT3, mTORC1.
- **KEGG / Reactome** provide the mechanistic wiring: FcγR → SYK/VAV1/MAPK → phagocytosis; lysosome/vacuolar acidification; ROS machinery (NCF1/2/4).
- **GO** decomposes to cellular localization (phagocytic vesicle, granule lumen, vacuolar membrane) and molecular functions (PRR activity, cytokine receptors, NADPH oxidase activator activity).
- **WikiPathways microglia/TYROBP** acts as a *cross-check* for myeloid innate signaling architecture.

So, across orthogonal annotation systems, the same story recurs: **activated innate phagocyte/APC biology is a core correlate of HAI at day 28**.

---

**Estimated Experimental Context (refined with your actual study design)**

Given you compared **PBMC day28 vs day0** and correlated expression change with **HAI at day 28**, the signal is most consistent with:

**(A) A “late innate activation / APC state” signature associated with antibody magnitude**

- Day 28 is relatively late; a classic plasmablast signature often peaks around day 7.
- So the strongest correlates at day 28 likely capture:
  - **persistent immune activation** in responders (perhaps due to stronger antigen persistence, better innate “set point”, or differential exposure/background immunity), and/or
  - **cellular composition shifts** (more monocytes, fewer lymphocytes, etc.).

**(B) A composition signal in PBMC**

The enrichment for “neutrophil degranulation”, lysosome, phagocytosis, azurophilic granule lumen strongly suggests:

- responders have a higher proportion of **CD14+/CD16+ monocytes** or more activated myeloid cells, or
- responders’ PBMC contain contamination or stronger transcriptional similarity to granulocyte programs (less likely if PBMC prep is clean, but the signature can still appear).

**(C) Possible involvement of IL-17/Th17 axis**

The IL-17 signaling module suggests responders have:

- a Th17-skewed inflammatory environment, or
- innate IL-17 axis activation that enhances antigen presentation and B-cell help indirectly.

---

**What this context changes in the analysis (practical implications)**

**1) Don’t interpret “neutrophil degranulation” literally as neutrophils**

In vaccine PBMC data, this term often means **myeloid activation** rather than actual neutrophils (since neutrophils are excluded). It’s still biologically meaningful as a correlate, but it’s telling you about **monocyte/APC effector programs**.

**2) Expect cell-type composition to be a major driver**

The most robust next step is to ask:

- Are these hubs simply reflecting **more monocytes / fewer lymphocytes** in responders at day 28?
  Because that would fully explain enrichment of lysosome/phagocytosis/ROS terms.

If you have cell proportion estimates (flow, CBC, or deconvolution like CIBERSORT/EPIC/ABIS), adjust for them and see if the signature remains.

**3) The presence of mTORC1 + metabolic enzymes becomes supportive evidence, not “metabolism per se”**

In vaccine responders, metabolic pathways often indicate **immune activation and proliferation**, not necessarily systemic metabolic disease.

**4) The network suggests “innate–humoral coupling” rather than “B cell intrinsic signature”**

This is important: your correlates may be highlighting immune environments that enable better humoral response, rather than antibody-producing cells directly.

---

If you want, I can also:

- extract a ranked list of the top cross-library hub genes (e.g., genes appearing in ≥3 libraries) from the DOT,
- propose a follow-up validation plan (cell-type adjustment, module scoring, and a causal model: innate module at day28 → HAI),
- and suggest how to write this up as a coherent “systems vaccinology” narrative.

--

**New prompt:**

is there anything in the network that is inconsistent with the experimental context? or potentially makes sense and novel?

## Response:

Yes — most of the network is **very consistent** with a vaccination/HAI-correlates analysis, but there are a few things that are either (a) potentially **inconsistent / suspicious**, or (b) **novel and worth leaning into** because they’re not the “usual suspects” in vaccine correlates.

Below I’ll separate those cleanly.

---

**What looks potentially inconsistent / could be an artifact**

**1)**
**“Neutrophil degranulation”, “azurophilic granule lumen”, “ficolin-1 rich granule”**
**in**
**PBMC**

PBMC preps should exclude neutrophils, so neutrophil granule terms can look “wrong” at first glance.

But in practice this is **very common** in vaccine PBMC expression:

- **Monocytes** share many of these genes (granule/lysosome proteins, antimicrobial enzymes).
- A tiny amount of granulocyte contamination can strongly influence RNA-seq/microarray.
- Some NK/monocyte activation states map heavily onto “neutrophil” annotated pathways because annotation is coarse.

So it’s not “inconsistent,” but it *does* mean:

> the strongest signal might be **myeloid abundance/activation** rather than a purely lymphocyte/B-cell program.

**How to check quickly:** look for classic neutrophil markers (e.g., *S100A8/A9, FCGR3B, CSF3R high with neutrophil granules*). You do have **CSF3R** and a lot of granule terms, but you *don’t* have the strongest canonical neutrophil transcriptome anchors (S100A8/A9 are absent here). That argues this is **monocyte-like granule biology**, not overt neutrophil contamination.

□ Likely explanation: **monocyte activation / “neutrophil-like” innate module**

---

**2)**
**WikiPathways “microglia” signatures**

You have:

- **TYROBP causal network in microglia**
- **Microglia pathogen phagocytosis pathway**

This sounds off for blood.

But those pathways are basically **generic myeloid innate programs** (TYROBP, Fc receptor signaling, phagocytosis machinery) written in a neuroimmune context. When you see them in blood, it usually means:

> strong **myeloid innate activation** rather than true neurobiology.

□ Not inconsistent; it’s an *annotation framing issue*.

---

**3)**
**Human Phenotype Ontology terms**
**(splenomegaly, agitation, apraxia, macular degeneration, etc.)**

These are not mechanistic pathways. In enrichment, HPO terms often appear because they are **gene collections linked to diseases/phenotypes**.

So it can look weird to interpret them literally (“why is vaccination inducing apraxia?”).

In your case they mostly reflect:

- lysosomal/immune deficiency genes → **splenomegaly**
- neurodegeneration genes (C9ORF72, GRN, PSEN1, VPS35) → **agitation/apraxia**
- retinal genes (BEST1) → **macular degeneration**

□ These should be interpreted as “this gene module overlaps disease gene sets,” not as actual phenotypes induced by the experiment.

---

**4) Some metabolism terms are “oddly specific”**

You have:

- **Sulfur metabolism (SULT1A1/A3/A4)**
- **Pantothenate and CoA biosynthesis (PANK2, PPCDC, VNN2)**
- **Amino sugar and nucleotide sugar metabolism (HK1, HK3, NAGK, NPL, etc.)**

These can be real (immune activation shifts metabolic enzymes), but they can also reflect:

- housekeeping/metabolic enzymes that sneak into signatures due to correlation structure,
- or **cell-type composition** differences (monocytes have distinct metabolic enzyme expression).

⚠ Not inconsistent, but “metabolism” may be secondary rather than primary biology unless these genes are among the strongest correlates.

---

**What looks**
**very consistent**
**with the vaccination / HAI-correlation context**

**A)**
**FcγR-mediated phagocytosis + SYK/VAV1/HCK/PAK1/MAPK1**

This is exactly what you might expect **at day 28**:

- individuals with higher HAI likely have more antibody/immune complexes,
- their monocytes may show enhanced FcγR signaling and phagocytic programs,
- or a “trained” innate state.

**B)**
**Lysosome/vacuolar acidification**

This is classic APC functionality:

- antigen uptake
- processing
- presentation support

Seeing it correlate with HAI makes sense as “better antigen handling → better humoral response.”

**C)**
**IL-6 / JAK-STAT3 and cytokine receptor activity**

Correlates of vaccine immunogenicity often involve STAT signaling — though usually earlier timepoints show stronger predictive power, seeing it at day 28 suggests:

- persistent immune activation in responders
- differences in immune set-point

**D)**
**Pattern recognition receptor activity (NOD2, TLR5, CLEC4E)**

This is a plausible innate sensing layer associated with responsiveness.

---

**What might be**
**novel / interesting**
**(and worth highlighting)**

These are the parts that feel like they could represent something more than generic “innate activation.”

**1)**
**Strong NOX2 / oxidative burst axis (NCF1/2/4 + “ROS and RNS production in phagocytes”)**

This is one of the more striking modules because it’s not always front-and-center in vaccine correlates papers.

**Possible novel angle:**

High HAI responders at day 28 show a persistent **phagocyte respiratory burst readiness** program, suggesting a link between antibody magnitude and innate effector conditioning (or trained immunity-like priming).

Mechanistically, ROS can modulate:

- antigen processing
- cytokine signaling thresholds
- inflammasome dynamics
- APC activation state

This is a coherent and potentially publishable “systems vaccinology” story.

---

**2)**
**SIGLEC / sialic-acid binding axis (SIGLEC5/7/10/12 + ST8SIA4)**

This is *very interesting* in an influenza vaccination context because influenza hemagglutinin binds **sialic acids**, and Siglecs are immune receptors that sense sialylated ligands and often dampen activation.

This module suggests responders differ in:

- inhibitory tuning of innate cells (Siglec expression),
- glycosylation environment (ST8SIA4),
- regulation of immune complex handling.

**Why this might be novel:**

Most vaccine correlates emphasize interferon modules, plasmablasts, Tfh signatures. A Siglec/sialic-acid regulation layer is less commonly emphasized.

Hypothesis:

Responders show altered **sialic-acid–Siglec immunoregulation**, potentially shaping how innate cells interact with HA-containing particles or immune complexes and modulating activation thresholds.

This is a great "unexpected but mechanistically plausible" thread.

---

**3)**
**IL-17 signaling at day 28**

Th17/IL-17 modules can appear in some vaccine studies, but it's not universally a hallmark correlate.

If real, it could imply:

- responders have a distinct inflammatory polarization (Th17-supportive environment),
- IL-17–driven recruitment/activation of myeloid cells supports antigen presentation and antibody responses indirectly.

Novel angle:

durable humoral responders retain a day-28 IL-17-associated innate activation program.

---

**4)**
**Retromer complex binding (SORT1, RAB7A, M6PR) + vesicle trafficking**

This could be just vesicle biology, but it could also matter for:

- antigen routing
- endosome/lysosome maturation
- receptor recycling (including immune receptors)

If this is strong in your data, it suggests:

responders differ in intracellular trafficking programs that may tune antigen processing/presentation.

That's subtle and interesting.

---

**Bottom line**

**Nothing is**
**glaringly inconsistent**
**with a vaccination PBMC day28 vs day0 HAI-correlate analysis.**

But the network clearly leans toward a **myeloid/APC effector program**, which might feel surprising if you expected a B-cell/Tfh signature at day 28 — and that makes **cell composition** and **innate immune set-point** the key interpretation issues.

**The most "novel / worth exploring" signals are:**

1. **NOX2 oxidative burst module (NCF1/2/4)**
2. **Siglec / sialic-acid binding + glycosylation regulation**
3. **IL-17 signaling persistence**
4. **trafficking/retromer antigen routing**

---

If you want, I can give you:

- a short list of "high-value candidate genes" from those novel modules for follow-up (e.g., NCF1/2/4, SIGLEC7/10, ST8SIA4, NOD2, IL17RA, SYK), and
- a concrete analysis plan to test whether these modules remain significant after adjusting for monocyte proportions.

## Supplementary Table 12

### Random gene list AI response

Prompt, network and full ChatGPT response for the random 300-gene list shown in Supplementary Figure 5A.

# Executive Summary

This iterative enrichment network points to a **dominant immune activation / cytokine-signaling program (IL-2/STAT5 and broader interleukin signaling)** that is intertwined with two major supporting “infrastructure” modules: **mitochondrial protein import / inner mitochondrial membrane organization** and **cell-cycle/mitotic spindle–centriole biology**. A small but notable “secretory/Golgi–granule” axis links immune signaling genes with trafficking and processing genes, suggesting an activated immune (or immune-like) state with high metabolic and secretory demand.

---

# Key Biological Processes Identified (by library source)

## H: Hallmark Gene Sets

**HALLMARK: IL2 STAT5 SIGNALING** is the dominant hallmark term, capturing a canonical **T-cell/immune activation** axis.

**Genes connected in your network:**

- **CD/immune/cytokine–linked:** *LIF, CD4 (in Reactome IL signaling), CSF3R, NFKBIZ*
- **immune–adjacent regulators/processing:** *FURIN, XBP1*
- **cellular activation/metabolism/transport:** *SLC2A3* (glucose transporter; often up in activated immune cells)
- **adhesion/immune interaction:** *ITGAV*
- **trafficking:** *SNX9*
- **calcium-binding / activation-related:** *S100A1*

**Interpretation:** This hallmark term suggests the “front-end” biology is **cytokine-driven immune activation**, consistent with STAT5-linked programs (IL-2 family cytokines, T-cell activation, survival, proliferation).

---

## C2: CP: Reactome Pathways

Two Reactome signals appear:

1. **REACTOME: OTHER INTERLEUKIN SIGNALING**
    - *CD4, CSF3R, STXBP2*

      **Interpretation:** Broad interleukin signaling support for the hallmark IL2/STAT5 theme. CSF3R points to granulocyte/innate signaling involvement, while CD4 suggests T-cell identity/signaling.

2. **REACTOME: MITOCHONDRIAL PROTEIN IMPORT**
    - *COQ2, COX17, MTX1, TIMM44, TIMM50*

      **Interpretation:** This is a strong "metabolic infrastructure" module. Activated immune cells often remodel mitochondrial function and import machinery to meet biosynthetic/energetic needs. It may also reflect **mitochondrial stress responses** coupled to immune activation.

---

## C2: CP: KEGG MEDICUS

### KEGG MEDICUS: DYNEIN RECRUITMENT TO THE KINETOCHORE

- *DYNC1H1, DYNLL2, KNTC1, ACTR10*

  **Interpretation:** A clean **mitotic machinery / chromosome segregation** signal. This aligns very strongly with the GO BP module "mitotic nuclear division." It implies a proliferative component (e.g., cycling immune cells).

---

## C2: CP: WikiPathways

### WP: 2Q112 COPY NUMBER VARIATION SYNDROME

- *COX11, FER1L5, KANSL3*

  **Interpretation:** This is likely acting as a **proxy gene set** that contains a mix of genes in your list (especially *KANSL3* which is part of chromatin/transcriptional regulation complexes, and *COX11* mitochondrial respiratory chain assembly). WikiPathways disease/CNV terms often behave as "gene containers," so biologically I'd interpret this as **chromatin/mitochondrial-linked genes co-occurring**, rather than a literal CNV syndrome readout.

---

## C5: GO Biological Process

You have several BP terms forming two major axes:

1. **Mitotic / centrosome module**

- **GOBP: MITOTIC NUCLEAR DIVISION**:

  *ABRAXAS1, CAV2, CEP85, DLGAP5, EDN3, KIF18A, KNTC1, PHIP, PIBF1, USP16*

- **GOBP: CENTRIOLE ELONGATION**:

  *CCDC15, POC1B, VPS4B*

**Interpretation:** This is a robust **cell-cycle/cell division** program, especially centrosome/spindle-related.

2. **Mitochondrial targeting/import module**

- **GOBP: PROTEIN TARGETING TO MITOCHONDRION**:

  *ARIH2, MTX1, TIMM44, TIMM50, TRMT10B*

**Interpretation:** Consistent with Reactome mitochondrial import.

3. **Protein acetylation / ER-stress-adjacent regulatory module**

- **GOBP: PROTEIN ACETYLATION**:

  *DIP2A, NAA50, XBP1*

**Interpretation:** Protein acetylation is broad, but the presence of **XBP1** is interesting because it is a key transcription factor in the **unfolded protein response (UPR)** and secretory differentiation. That ties naturally to the GO CC terms around secretory granule membrane / Golgi lumen.

4. **Host–pathogen / entry-related term**

- **GOBP: MODULATION BY SYMBIONT OF ENTRY INTO HOST**:

  *CD4, FURIN, ITGAV*

**Interpretation:** This is a very informative “cross-library bridge”:

- **CD4** is a classic receptor used by pathogens (e.g., HIV).
- **FURIN** is a host protease used for cleavage/activation of multiple viral/bacterial proteins and host receptors.
- **ITGAV** is a known entry/adhesion integrin used by multiple viruses.

Even if your experiment isn't infection-related, this term often emerges when a set includes **host entry factors + immune activation**.

---

### C5: GO Cellular Component

These terms highlight subcellular "execution sites":

- **GOCC: SECRETORY GRANULE MEMBRANE**

  *CADPS, CAV2, FAM170B, GP2, ITGAV, RND2, SLC2A3, SLC30A5, STXBP2, TRPM2*

  → Strong signal for **secretory vesicle/granule biology**, typical of immune effector cells, endocrine/secretory cells, or activated cells with high vesicle trafficking.

- **GOCC: GOLGI LUMEN**

  *F8, FURIN, LHB, PROS1, RAB33B*

  → Points to **processing/maturation/secretion**, with **FURIN** being a key Golgi protease.

- **GOCC: INNER MITOCHONDRIAL MEMBRANE PROTEIN COMPLEX**

  *MTX1, TIMM44, TIMM50, TRMT10B*

  → Reinforces mitochondrial import/organization.

- **GOCC: SPINDLE POLE**

  *CEP85, CKAP2L, DLGAP5, KNTC1, MTCL1, POC1B, VPS4B*

  → Reinforces mitosis/spindle/centrosome.

---

# Network Topology Insights

### 1) Hub genes (multi-term connectors)

These are genes that “bridge” multiple libraries/terms and likely represent key biological levers:

- **CD4**

  Links **Reactome interleukin signaling** + **GO “symbiont entry”** + BP terms (host interaction)

  → A *core immune identity/signaling hub*.

- **FURIN**

  Links **Hallmark IL2/STAT5**, **GO “symbiont entry”**, **Golgi lumen**

  → A *processing/activation hub* at the secretory/Golgi interface that can affect cytokines, receptors, and pathogen entry.

- **ITGAV**

  Links **Hallmark IL2/STAT5**, **GO “symbiont entry”**, **secretory granule membrane**

  → *adhesion/entry + immune activation bridge*.

- **KNTC1**

  Links **KEGG kinetochore/dynein recruitment**, **GO mitotic division**, **GO spindle pole**

  → A *pure cell division/mitosis hub*.

- **MTX1, TIMM44, TIMM50**

  Link **Reactome mitochondrial import** + **GO mitochondrial targeting/import** + **inner mitochondrial membrane complex**

  → *mitochondrial biogenesis/import hubs*.

- **STXBP2**

  Links **Reactome interleukin signaling** + **secretory granule membrane**

  → Suggests an immune effector secretory component (this gene is famous for cytotoxic granule exocytosis defects in immune disease contexts).

- **CAV2**

  Links **mitotic nuclear division** + **secretory granule membrane**

→ An unusual connector that may reflect membrane dynamics in proliferating/activated cells.

## 2) Modules / clusters

You have **at least three distinct modules**:

1. **Immune signaling / cytokine module**
    - Hallmark IL2/STAT5 + Reactome other interleukin signaling
    - Centered around *CD4, CSF3R, NFKBIZ, LIF*

    **Meaning:** immune activation state, likely lymphoid + myeloid elements or mixed immune signatures.

2. **Proliferation / spindle–centriole module**
    - KEGG dynein/kinetochore + GO mitosis + GO spindle pole + centriole elongation
    - Centered around *KNTC1, DYNC1H1, DLGAP5, KIF18A, CEP85, POC1B*

    **Meaning:** proliferating cells, cycling immune cells (activated T cells) or proliferative tissue context.

3. **Mitochondrial import / inner membrane module**
    - Reactome mitochondrial import + GO mitochondrial targeting + inner IMM complex
    - Centered around *TIMM44/TIMM50/MTX1/COQ2/COX17*

    **Meaning:** metabolic reprogramming, mitochondrial biogenesis, or stress response.

Plus a **secretory/Golgi module**:

- GO secretory granule membrane + Golgi lumen
- Centered around *FURIN, STXBP2, CADPS, RAB33B*

  **Meaning:** activated secretory trafficking and protein processing.

## 3) Connectivity pattern and biological organization

This network looks like a classic **“activation state” architecture**:

- An upstream **immune signaling trigger** (IL2/STAT5 + interleukins)
- A downstream **execution layer** (Golgi/secretory machinery)
- A supportive **resource layer** (mitochondria, import, IMM organization)
- A **growth/expansion layer** (mitosis/spindle/centriole)

That combination is very consistent with **cells responding to cytokine stimulation by proliferating and ramping up metabolism + secretion**.

---

# Proposed Biological Hypothesis (integrating terms + topology)

## Main hypothesis

Your 300-gene list likely reflects a population (or condition) undergoing **cytokine-driven immune activation** (IL-2/STAT5 and broader interleukin signaling) that triggers:

1. **Clonal expansion / proliferation**

   → evidenced by the strong kinetochore/spindle/centriole/mitotic nuclear division modules (KNTC1, DYNC1H1, KIF18A, DLGAP5).

2. **Metabolic remodeling with increased mitochondrial biogenesis/import**

   → evidenced by mitochondrial protein import + inner mitochondrial membrane complex genes (TIMM44, TIMM50, MTX1, COQ2, COX17).

3. **Increased secretory processing and vesicle trafficking**

   → evidenced by secretory granule membrane + Golgi lumen (STXBP2, CADPS, RAB33B) and **FURIN** as a central proteolytic processing hub.

## How library sources complement each other

- **Hallmark** gives the clean “state label”: *IL2/STAT5 signaling* = immune activation.
- **Reactome** provides mechanistic pathway scaffolding: interleukin signaling and mitochondrial import.
- **KEGG** captures cell-cycle machinery from a different angle (kinetochore/dynein recruitment).
- **GO BP/CC** resolves where things happen and expands the mechanistic picture into **cell division structures (spindle pole)** and **subcellular organelles (Golgi, granule membrane, mitochondria)**.
- **WikiPathways** adds a “disease-ish container” that likely reflects the co-occurrence of chromatin/mitochondrial genes, but should be interpreted cautiously.

---

# Estimated Experimental Context (what generated this result?)

Given the strong IL2/STAT5 + interleukin signaling *and* the concurrent mitotic and mitochondrial remodeling signals, the most plausible contexts are:

1. **Immune cell activation / stimulation experiment**
    - e.g., **T cell activation** (anti-CD3/CD28 ± IL-2), **PBMC stimulation**, cytokine exposure, or immune activation in tissue.
    - This fits: IL2/STAT5, CD4 involvement, proliferation genes, metabolic remodeling.
2. **Inflammation-associated tissue profiling**
    - e.g., inflamed tissue biopsy (intestinal, lung, etc.) with infiltrating immune cells.
    - Would naturally produce mixed immune signatures (CD4 + CSF3R) plus activation/proliferation.
3. **Infection or viral entry–relevant perturbation**
    - The presence of the GO “symbiont entry” term with **CD4, FURIN, ITGAV** could reflect:
        - actual infection model, **or**
        - an inflammatory immune activation state enriched for host entry factors.
    - If this came from epithelial/immune mixed samples, the ITGAV + FURIN + trafficking genes could reflect a host defense/entry axis.

**My best single guess:** a **cytokine-stimulated immune cell dataset** (especially T-cell–biased) where activation drives **proliferation** and requires **mitochondrial upscaling**, with secretory machinery activation (possibly cytotoxic/secretory phenotype features).

---

If you want, I can also:

- rank the **top 10 hub genes** quantitatively (degree in this DOT graph),
- propose a **minimal mechanistic cartoon** (cytokine → STAT5 → proliferation + mitochondrial import + Golgi/secretory processing),
- suggest **follow-up validations** (e.g., phospho-STAT5, Ki-67/cell-cycle markers, mitochondrial mass/respiration, degranulation assays).
