## Supplementary Table 9 for "Iterative Gene Enrichment Analysis: interpretable networks for human and AI-assisted biological insights"

| Cluster | Cluster Enrichment Score | Category | Term | Count | List Total | Pop Hits | Pop Total | % |
|---|---|---|---|---|---|---|---|---|
| 1 | 5.68 | GOTERM_CC_DIRECT | nucleoplasm | 90 | 255 | 4061 | 19687 | 33.09 |
| 1 | 5.68 | GOTERM_CC_DIRECT | nucleus | 123 | 255 | 6821 | 19687 | 45.22 |
| 1 | 5.68 | UP_KW_CELLULAR_COMPONENT | Nucleus | 111 | 240 | 5755 | 17335 | 40.81 |
| 2 | 5.35 | GOTERM_BP_DIRECT | protein transport | 29 | 247 | 676 | 18757 | 10.66 |
| 2 | 5.35 | UP_KW_BIOLOGICAL_PROCESS | Protein transport | 28 | 163 | 633 | 11213 | 10.29 |
| 2 | 5.35 | UP_KW_BIOLOGICAL_PROCESS | Transport | 45 | 163 | 2051 | 11213 | 16.54 |
| 3 | 4.99 | REACTOME_PATHWAY | Membrane Trafficking | 29 | 176 | 642 | 11462 | 10.66 |
| 3 | 4.99 | REACTOME_PATHWAY | Asparagine N-linked glycosylation | 19 | 176 | 315 | 11462 | 6.99 |
| 3 | 4.99 | REACTOME_PATHWAY | COPI-mediated anterograde transport | 11 | 176 | 103 | 11462 | 4.04 |
| 3 | 4.99 | REACTOME_PATHWAY | ER to Golgi Anterograde Transport | 13 | 176 | 158 | 11462 | 4.78 |
| 3 | 4.99 | REACTOME_PATHWAY | Transport to the Golgi and subsequent modification | 14 | 176 | 189 | 11462 | 5.15 |
| 3 | 4.99 | REACTOME_PATHWAY | Vesicle-mediated transport | 29 | 176 | 744 | 11462 | 10.66 |
| 3 | 4.99 | REACTOME_PATHWAY | Intra-Golgi and retrograde Golgi-to-ER traffic | 14 | 176 | 208 | 11462 | 5.15 |
| 3 | 4.99 | GOTERM_BP_DIRECT | vesicle-mediated transport | 15 | 247 | 336 | 18757 | 5.51 |
| 3 | 4.99 | REACTOME_PATHWAY | Intra-Golgi traffic | 6 | 176 | 47 | 11462 | 2.21 |
| 4 | 4.80 | REACTOME_PATHWAY | Viral Infection Pathways | 36 | 176 | 890 | 11462 | 13.24 |
| 4 | 4.80 | REACTOME_PATHWAY | Infectious disease | 38 | 176 | 1119 | 11462 | 13.97 |
| 4 | 4.80 | REACTOME_PATHWAY | Disease | 52 | 176 | 1951 | 11462 | 19.12 |
| 4 | 4.80 | REACTOME_PATHWAY | SARS-CoV Infections | 20 | 176 | 483 | 11462 | 7.35 |
| 4 | 4.80 | REACTOME_PATHWAY | SARS-CoV-2 Infection | 15 | 176 | 303 | 11462 | 5.51 |
| 5 | 4.05 | GOTERM_CC_DIRECT | Golgi membrane | 25 | 255 | 718 | 19687 | 9.19 |
| 5 | 4.05 | GOTERM_CC_DIRECT | Golgi apparatus | 37 | 255 | 1463 | 19687 | 13.60 |
| 5 | 4.05 | UP_KW_CELLULAR_COMPONENT | Golgi apparatus | 29 | 240 | 989 | 17335 | 10.66 |
| 6 | 3.98 | REACTOME_PATHWAY | ER to Golgi Anterograde Transport | 13 | 176 | 158 | 11462 | 4.78 |
| 6 | 3.98 | GOTERM_BP_DIRECT | endoplasmic reticulum to Golgi vesicle-mediated transport | 10 | 247 | 128 | 18757 | 3.68 |

| Cluster | Cluster Enrichment Score | Category | Term | Count | List Total | Pop Hits | Pop Total | % |
|---|---|---|---|---|---|---|---|---|
| 6 | 3.98 | REACTOME_PATHWAY | COPII-mediated vesicle transport | 6 | 176 | 71 | 11462 | 2.21 |
| 7 | 3.34 | REACTOME_PATHWAY | Regulation of Glucokinase by Glucokinase Regulatory Protein | 8 | 176 | 32 | 11462 | 2.94 |
| 7 | 3.34 | REACTOME_PATHWAY | Defective TPR may confer susceptibility towards thyroid papillary carcinoma (TPC) | 8 | 176 | 32 | 11462 | 2.94 |
| 7 | 3.34 | REACTOME_PATHWAY | snRNP Assembly | 9 | 176 | 54 | 11462 | 3.31 |
| 7 | 3.34 | REACTOME_PATHWAY | Metabolism of non-coding RNA | 9 | 176 | 54 | 11462 | 3.31 |
| 7 | 3.34 | REACTOME_PATHWAY | HIV Life Cycle | 13 | 176 | 149 | 11462 | 4.78 |
| 7 | 3.34 | REACTOME_PATHWAY | IPs transport between nucleus and cytosol | 7 | 176 | 30 | 11462 | 2.57 |
| 7 | 3.34 | REACTOME_PATHWAY | IP3 and IP4 transport between cytosol and nucleus | 7 | 176 | 30 | 11462 | 2.57 |
| 7 | 3.34 | REACTOME_PATHWAY | IP6 and IP7 transport between cytosol and nucleus | 7 | 176 | 30 | 11462 | 2.57 |
| 7 | 3.34 | REACTOME_PATHWAY | HIV Infection | 15 | 176 | 220 | 11462 | 5.51 |
| 7 | 3.34 | REACTOME_PATHWAY | Transport of Ribonucleoproteins into the Host Nucleus | 7 | 176 | 32 | 11462 | 2.57 |
| 7 | 3.34 | REACTOME_PATHWAY | NEP/NS2 Interacts with the Cellular Export Machinery | 7 | 176 | 32 | 11462 | 2.57 |
| 7 | 3.34 | REACTOME_PATHWAY | Export of Viral Ribonucleoproteins from Nucleus | 7 | 176 | 34 | 11462 | 2.57 |
| 7 | 3.34 | REACTOME_PATHWAY | Nuclear import of Rev protein | 7 | 176 | 34 | 11462 | 2.57 |
| 7 | 3.34 | REACTOME_PATHWAY | Vpr-mediated nuclear import of PICs | 7 | 176 | 34 | 11462 | 2.57 |
| 7 | 3.34 | REACTOME_PATHWAY | Transport of the SLBP independent Mature mRNA | 7 | 176 | 35 | 11462 | 2.57 |
| 7 | 3.34 | REACTOME_PATHWAY | Rev-mediated nuclear export of HIV RNA | 7 | 176 | 35 | 11462 | 2.57 |
| 7 | 3.34 | REACTOME_PATHWAY | SUMOylation of SUMOylation proteins | 7 | 176 | 35 | 11462 | 2.57 |

| Cluster | Cluster Enrichment Score | Category | Term | Count | List Total | Pop Hits | Pop Total | % |
|---|---|---|---|---|---|---|---|---|
| 7 | 3.34 | REACTOME_PATHWAY | Transport of the SLBP Dependant Mature mRNA | 7 | 176 | 36 | 11462 | 2.57 |
| 7 | 3.34 | REACTOME_PATHWAY | Nuclear Pore Complex (NPC) Disassembly | 7 | 176 | 36 | 11462 | 2.57 |
| 7 | 3.34 | REACTOME_PATHWAY | Interactions of Vpr with host cellular proteins | 7 | 176 | 37 | 11462 | 2.57 |
| 7 | 3.34 | REACTOME_PATHWAY | Interactions of Rev with host cellular proteins | 7 | 176 | 37 | 11462 | 2.57 |
| 7 | 3.34 | GOTERM_MF_DIRECT | structural constituent of nuclear pore | 6 | 252 | 25 | 18532 | 2.21 |
| 7 | 3.34 | REACTOME_PATHWAY | SUMOylation of ubiquitinylation proteins | 7 | 176 | 39 | 11462 | 2.57 |
| 7 | 3.34 | GOTERM_CC_DIRECT | nuclear pore | 9 | 255 | 94 | 19687 | 3.31 |
| 7 | 3.34 | REACTOME_PATHWAY | NS1 Mediated Effects on Host Pathways | 7 | 176 | 41 | 11462 | 2.57 |
| 7 | 3.34 | UP_KW_CELLULAR_COMPONENT | Nuclear pore complex | 7 | 240 | 46 | 17335 | 2.57 |
| 7 | 3.34 | REACTOME_PATHWAY | Transport of Mature mRNA Derived from an Intronless Transcript | 7 | 176 | 42 | 11462 | 2.57 |
| 7 | 3.34 | REACTOME_PATHWAY | Late Phase of HIV Life Cycle | 11 | 176 | 136 | 11462 | 4.04 |
| 7 | 3.34 | REACTOME_PATHWAY | Transport of Mature mRNAs Derived from Intronless Transcripts | 7 | 176 | 43 | 11462 | 2.57 |
| 7 | 3.34 | REACTOME_PATHWAY | Viral Messenger RNA Synthesis | 7 | 176 | 44 | 11462 | 2.57 |
| 7 | 3.34 | GOTERM_BP_DIRECT | nucleocytoplasmic transport | 7 | 247 | 52 | 18757 | 2.57 |
| 7 | 3.34 | REACTOME_PATHWAY | SUMOylation of DNA replication proteins | 7 | 176 | 46 | 11462 | 2.57 |
| 7 | 3.34 | REACTOME_PATHWAY | SUMOylation of RNA binding proteins | 7 | 176 | 48 | 11462 | 2.57 |
| 7 | 3.34 | REACTOME_PATHWAY | Host Interactions of HIV factors | 10 | 176 | 120 | 11462 | 3.68 |
| 7 | 3.34 | REACTOME_PATHWAY | Regulation of HSF1-mediated heat shock response | 8 | 176 | 71 | 11462 | 2.94 |
| 7 | 3.34 | REACTOME_PATHWAY | Nuclear Envelope (NE) Reassembly | 8 | 176 | 72 | 11462 | 2.94 |

| Cluster | Cluster Enrichment Score | Category | Term | Count | List Total | Pop Hits | Pop Total | % |
|---|---|---|---|---|---|---|---|---|
| 7 | 3.34 | REACTOME_PATHWAY | Nuclear Envelope Breakdown | 7 | 176 | 51 | 11462 | 2.57 |
| 7 | 3.34 | KEGG_PATHWAY | Nucleocytoplasmic transport | 9 | 120 | 108 | 8506 | 3.31 |
| 7 | 3.34 | REACTOME_PATHWAY | ISG15 antiviral mechanism | 8 | 176 | 74 | 11462 | 2.94 |
| 7 | 3.34 | REACTOME_PATHWAY | SARS-CoV-2 activates/ modulates innate and adaptive immune responses | 10 | 176 | 126 | 11462 | 3.68 |
| 7 | 3.34 | REACTOME_PATHWAY | Transport of Mature mRNA derived from an Intron-Containing Transcript | 8 | 176 | 75 | 11462 | 2.94 |
| 7 | 3.34 | REACTOME_PATHWAY | Glycolysis | 8 | 176 | 75 | 11462 | 2.94 |
| 7 | 3.34 | REACTOME_PATHWAY | SARS-CoV Infections | 20 | 176 | 483 | 11462 | 7.35 |
| 7 | 3.34 | REACTOME_PATHWAY | Inositol phosphate metabolism | 8 | 176 | 80 | 11462 | 2.94 |
| 7 | 3.34 | UP_KW_BIOLOGICAL_PROCESS | Translocation | 8 | 163 | 85 | 11213 | 2.94 |
| 7 | 3.34 | REACTOME_PATHWAY | SARS-CoV-2 Infection | 15 | 176 | 303 | 11462 | 5.51 |
| 7 | 3.34 | GOTERM_CC_DIRECT | nuclear pore outer ring | 4 | 255 | 10 | 19687 | 1.47 |
| 7 | 3.34 | REACTOME_PATHWAY | tRNA processing in the nucleus | 7 | 176 | 58 | 11462 | 2.57 |
| 7 | 3.34 | REACTOME_PATHWAY | HCMV Early Events | 10 | 176 | 138 | 11462 | 3.68 |
| 7 | 3.34 | REACTOME_PATHWAY | Transport of Mature Transcript to Cytoplasm | 8 | 176 | 84 | 11462 | 2.94 |
| 7 | 3.34 | REACTOME_PATHWAY | tRNA processing | 9 | 176 | 111 | 11462 | 3.31 |
| 7 | 3.34 | REACTOME_PATHWAY | SARS-CoV-2-host interactions | 12 | 176 | 204 | 11462 | 4.41 |
| 7 | 3.34 | REACTOME_PATHWAY | Mitotic Prophase | 10 | 176 | 142 | 11462 | 3.68 |
| 7 | 3.34 | REACTOME_PATHWAY | Metabolism of RNA | 25 | 176 | 734 | 11462 | 9.19 |
| 7 | 3.34 | REACTOME_PATHWAY | Glucose metabolism | 8 | 176 | 86 | 11462 | 2.94 |
| 7 | 3.34 | GOTERM_BP_DIRECT | nephron development | 4 | 247 | 11 | 18757 | 1.47 |
| 7 | 3.34 | REACTOME_PATHWAY | Cellular response to heat stress | 8 | 176 | 91 | 11462 | 2.94 |
| 7 | 3.34 | REACTOME_PATHWAY | Antiviral mechanism by IFN-stimulated genes | 10 | 176 | 150 | 11462 | 3.68 |
| 7 | 3.34 | REACTOME_PATHWAY | Postmitotic nuclear pore | 5 | 176 | 25 | 11462 | 1.84 |

| Cluster | Cluster Enrichment Score | Category | Term | Count | List Total | Pop Hits | Pop Total | % |
|---|---|---|---|---|---|---|---|---|
| | | | complex (NPC) reformation | | | | | |
| 7 | 3.34 | GOTERM_BP_DIRECT | mRNA transport | 8 | 247 | 110 | 18757 | 2.94 |
| 7 | 3.34 | REACTOME_PATHWAY | Cellular responses to stimuli | 28 | 176 | 913 | 11462 | 10.29 |
| 7 | 3.34 | UP_KW_BIOLOGICAL_PROCESS | mRNA transport | 8 | 163 | 103 | 11213 | 2.94 |
| 7 | 3.34 | REACTOME_PATHWAY | SUMOylation of chromatin organization proteins | 7 | 176 | 71 | 11462 | 2.57 |
| 7 | 3.34 | REACTOME_PATHWAY | SLC transporter disorders | 8 | 176 | 98 | 11462 | 2.94 |
| 7 | 3.34 | REACTOME_PATHWAY | HCMV Infection | 10 | 176 | 162 | 11462 | 3.68 |
| 7 | 3.34 | GOTERM_BP_DIRECT | mRNA export from nucleus | 6 | 247 | 59 | 18757 | 2.21 |
| 7 | 3.34 | REACTOME_PATHWAY | SUMOylation of DNA damage response and repair proteins | 7 | 176 | 78 | 11462 | 2.57 |
| 7 | 3.34 | REACTOME_PATHWAY | Transcriptional regulation by small RNAs | 8 | 176 | 107 | 11462 | 2.94 |
| 7 | 3.34 | REACTOME_PATHWAY | Gene Silencing by RNA | 9 | 176 | 139 | 11462 | 3.31 |
| 7 | 3.34 | REACTOME_PATHWAY | SUMO E3 ligases SUMOylate target proteins | 10 | 176 | 181 | 11462 | 3.68 |
| 7 | 3.34 | REACTOME_PATHWAY | SUMOylation | 10 | 176 | 187 | 11462 | 3.68 |
| 7 | 3.34 | GOTERM_CC_DIRECT | nuclear envelope | 10 | 255 | 226 | 19687 | 3.68 |
| 7 | 3.34 | GOTERM_BP_DIRECT | protein import into nucleus | 7 | 247 | 110 | 18757 | 2.57 |
| 7 | 3.34 | WIKIPATHWAYS | Prader Willi and Angelman syndrome | 7 | 138 | 108 | 8952 | 2.57 |
| 7 | 3.34 | REACTOME_PATHWAY | Mitotic Anaphase | 10 | 176 | 218 | 11462 | 3.68 |
| 7 | 3.34 | REACTOME_PATHWAY | Mitotic Metaphase and Anaphase | 10 | 176 | 219 | 11462 | 3.68 |
| 7 | 3.34 | REACTOME_PATHWAY | EML4 and NUDC in mitotic spindle formation | 7 | 176 | 115 | 11462 | 2.57 |
| 7 | 3.34 | REACTOME_PATHWAY | HCMV Late Events | 7 | 176 | 116 | 11462 | 2.57 |
| 7 | 3.34 | REACTOME_PATHWAY | M Phase | 14 | 176 | 403 | 11462 | 5.15 |
| 7 | 3.34 | REACTOME_PATHWAY | Cell Cycle, Mitotic | 17 | 176 | 550 | 11462 | 6.25 |
| 7 | 3.34 | REACTOME_PATHWAY | Processing of Capped Intron-Containing Pre-mRNA | 11 | 176 | 282 | 11462 | 4.04 |
| 7 | 3.34 | REACTOME_PATHWAY | Resolution of Sister Chromatid Cohesion | 7 | 176 | 125 | 11462 | 2.57 |

| Cluster | Cluster Enrichment Score | Category | Term | Count | List Total | Pop Hits | Pop Total | % |
|---|---|---|---|---|---|---|---|---|
| 7 | 3.34 | REACTOME_PATHWAY | Disorders of transmembrane transporters | 8 | 176 | 163 | 11462 | 2.94 |
| 7 | 3.34 | REACTOME_PATHWAY | Amplification of signal from the kinetochores | 6 | 176 | 92 | 11462 | 2.21 |
| 7 | 3.34 | REACTOME_PATHWAY | Amplification of signal from unattached kinetochores via a MAD2 inhibitory signal | 6 | 176 | 92 | 11462 | 2.21 |
| 7 | 3.34 | REACTOME_PATHWAY | Influenza Viral RNA Transcription and Replication | 7 | 176 | 137 | 11462 | 2.57 |
| 7 | 3.34 | REACTOME_PATHWAY | RHO GTPases Activate Formins | 7 | 176 | 138 | 11462 | 2.57 |
| 7 | 3.34 | GOTERM_CC_DIRECT | nuclear membrane | 9 | 255 | 268 | 19687 | 3.31 |
| 7 | 3.34 | REACTOME_PATHWAY | Mitotic Spindle Checkpoint | 6 | 176 | 109 | 11462 | 2.21 |
| 7 | 3.34 | REACTOME_PATHWAY | Interferon Signaling | 10 | 176 | 279 | 11462 | 3.68 |
| 7 | 3.34 | KEGG_PATHWAY | Amyotrophic lateral sclerosis | 11 | 120 | 364 | 8506 | 4.04 |
| 7 | 3.34 | REACTOME_PATHWAY | Cell Cycle | 18 | 176 | 686 | 11462 | 6.62 |
| 7 | 3.34 | REACTOME_PATHWAY | Cytokine Signaling in Immune system | 20 | 176 | 793 | 11462 | 7.35 |
| 7 | 3.34 | REACTOME_PATHWAY | Influenza Infection | 7 | 176 | 158 | 11462 | 2.57 |
| 7 | 3.34 | REACTOME_PATHWAY | Separation of Sister Chromatids | 7 | 176 | 176 | 11462 | 2.57 |
| 7 | 3.34 | REACTOME_PATHWAY | RHO GTPase Effectors | 10 | 176 | 325 | 11462 | 3.68 |
| 7 | 3.34 | REACTOME_PATHWAY | Mitotic Prometaphase | 7 | 176 | 202 | 11462 | 2.57 |
| 7 | 3.34 | GOTERM_CC_DIRECT | kinetochore | 5 | 255 | 168 | 19687 | 1.84 |
| 7 | 3.34 | REACTOME_PATHWAY | Metabolism of carbohydrates and carbohydrate derivatives | 8 | 176 | 303 | 11462 | 2.94 |
| 7 | 3.34 | REACTOME_PATHWAY | Cell Cycle Checkpoints | 7 | 176 | 288 | 11462 | 2.57 |
| 7 | 3.34 | GOTERM_CC_DIRECT | chromosome, centromeric region | 4 | 255 | 166 | 19687 | 1.47 |
| 7 | 3.34 | UP_KW_CELLULAR_COMPONENT | Kinetochore | 3 | 240 | 106 | 17335 | 1.10 |
| 7 | 3.34 | UP_KW_CELLULAR_COMPONENT | Centromere | 3 | 240 | 150 | 17335 | 1.10 |
| 8 | 2.75 | REACTOME_PATHWAY | Retrograde transport at the | 7 | 176 | 51 | 11462 | 2.57 |

| Cluster | Cluster Enrichment Score | Category | Term | Count | List Total | Pop Hits | Pop Total | % |
|---|---|---|---|---|---|---|---|---|
| | | | Trans-Golgi-Network | | | | | |
| 8 | 2.75 | UP_KW_DISEASE | Congenital disorder of glycosylation | 7 | 92 | 54 | 4854 | 2.57 |
| 8 | 2.75 | REACTOME_PATHWAY | Intra-Golgi traffic | 6 | 176 | 47 | 11462 | 2.21 |
| 8 | 2.75 | GOTERM_CC_DIRECT | Golgi transport complex | 3 | 255 | 10 | 19687 | 1.10 |
| 8 | 2.75 | GOTERM_BP_DIRECT | retrograde transport, vesicle recycling within Golgi | 3 | 247 | 10 | 18757 | 1.10 |
| 8 | 2.75 | GOTERM_CC_DIRECT | trans-Golgi network membrane | 6 | 255 | 112 | 19687 | 2.21 |
| 9 | 2.23 | GOTERM_BP_DIRECT | intra-Golgi vesicle-mediated transport | 7 | 247 | 33 | 18757 | 2.57 |
| 9 | 2.23 | GOTERM_BP_DIRECT | retrograde vesicle-mediated transport, Golgi to endoplasmic reticulum | 8 | 247 | 51 | 18757 | 2.94 |
| 9 | 2.23 | UP_SEQ_FEATURE | MOTIF:Switch 2 | 5 | 261 | 47 | 19119 | 1.84 |
| 9 | 2.23 | UP_SEQ_FEATURE | MOTIF:Switch 1 | 5 | 261 | 47 | 19119 | 1.84 |
| 9 | 2.23 | GOTERM_BP_DIRECT | protein localization to Golgi membrane | 3 | 247 | 8 | 18757 | 1.10 |
| 9 | 2.23 | GOTERM_BP_DIRECT | retrograde transport, endosome to Golgi | 6 | 247 | 87 | 18757 | 2.21 |
| 9 | 2.23 | GOTERM_MF_DIRECT | GTP binding | 12 | 252 | 387 | 18532 | 4.41 |
| 9 | 2.23 | GOTERM_MF_DIRECT | GTPase activity | 11 | 252 | 347 | 18532 | 4.04 |
| 9 | 2.23 | UP_KW_PTM | Prenylation | 7 | 208 | 174 | 14032 | 2.57 |
| 9 | 2.23 | UP_SEQ_FEATURE | LIPID:S-geranylgeranyl cysteine | 5 | 261 | 111 | 19119 | 1.84 |
| 9 | 2.23 | GOTERM_MF_DIRECT | G protein activity | 4 | 252 | 73 | 18532 | 1.47 |
| 9 | 2.23 | REACTOME_PATHWAY | RAB geranylgeranylation | 4 | 176 | 66 | 11462 | 1.47 |
| 9 | 2.23 | GOTERM_MF_DIRECT | GDP binding | 3 | 252 | 79 | 18532 | 1.10 |
| 10 | 2.20 | REACTOME_PATHWAY | Transcriptional regulation of white adipocyte differentiation | 9 | 176 | 83 | 11462 | 3.31 |
| 10 | 2.20 | REACTOME_PATHWAY | Adipogenesis | 9 | 176 | 109 | 11462 | 3.31 |
| 10 | 2.20 | GOTERM_CC_DIRECT | core mediator complex | 5 | 255 | 26 | 19687 | 1.84 |
| 10 | 2.20 | WIKIPATHWAYS | RNA polymerase II transcription the | 7 | 138 | 69 | 8952 | 2.57 |

| Cluster | Cluster Enrichment Score | Category | Term | Count | List Total | Pop Hits | Pop Total | % |
|---|---|---|---|---|---|---|---|---|
| | | | preinitiation complex | | | | | |
| 10 | 2.20 | GOTERM_CC_DIRECT | mediator complex | 5 | 255 | 38 | 19687 | 1.84 |
| 10 | 2.20 | REACTOME_PATHWAY | Respiratory Syncytial Virus Infection Pathway | 8 | 176 | 121 | 11462 | 2.94 |
| 10 | 2.20 | GOTERM_BP_DIRECT | positive regulation of transcription elongation by RNA polymerase II | 5 | 247 | 57 | 18757 | 1.84 |
| 10 | 2.20 | GOTERM_BP_DIRECT | RNA polymerase II preinitiation complex assembly | 5 | 247 | 61 | 18757 | 1.84 |
| 10 | 2.20 | GOTERM_BP_DIRECT | positive regulation of transcription initiation by RNA polymerase II | 5 | 247 | 62 | 18757 | 1.84 |
| 10 | 2.20 | REACTOME_PATHWAY | PPARA activates gene expression | 7 | 176 | 117 | 11462 | 2.57 |
| 10 | 2.20 | REACTOME_PATHWAY | Regulation of lipid metabolism by PPARalpha | 7 | 176 | 118 | 11462 | 2.57 |
| 10 | 2.20 | GOTERM_BP_DIRECT | somatic stem cell population maintenance | 5 | 247 | 65 | 18757 | 1.84 |
| 10 | 2.20 | REACTOME_PATHWAY | Epigenetic regulation by WDR5-containing histone modifying complexes | 8 | 176 | 167 | 11462 | 2.94 |
| 10 | 2.20 | REACTOME_PATHWAY | RSV-host interactions | 6 | 176 | 98 | 11462 | 2.21 |
| 10 | 2.20 | REACTOME_PATHWAY | MLL4 and MLL3 complexes regulate expression of PPARG target genes in adipogenesis and hepatic steatosis | 7 | 176 | 135 | 11462 | 2.57 |
| 10 | 2.20 | REACTOME_PATHWAY | Epigenetic regulation of adipogenesis genes by MLL3 and MLL4 complexes | 7 | 176 | 135 | 11462 | 2.57 |
| 10 | 2.20 | REACTOME_PATHWAY | Epigenetic regulation of gene expression by MLL3 and MLL4 complexes | 7 | 176 | 135 | 11462 | 2.57 |
| 10 | 2.20 | REACTOME_PATHWAY | Epigenetic regulation of gene expression | 11 | 176 | 321 | 11462 | 4.04 |
| 10 | 2.20 | GOTERM_MF_DIRECT | transcription coregulator activity | 6 | 252 | 136 | 18532 | 2.21 |
| 10 | 2.20 | GOTERM_MF_DIRECT | transcription coactivator activity | 9 | 252 | 292 | 18532 | 3.31 |

| Cluster | Cluster Enrichment Score | Category | Term | Count | List Total | Pop Hits | Pop Total | % |
|---|---|---|---|---|---|---|---|---|
| 10 | 2.20 | REACTOME_PATHWAY | Metabolism of lipids | 14 | 176 | 756 | 11462 | 5.15 |
| 11 | 2.09 | GOTERM_CC_DIRECT | endoplasmic reticulum membrane | 29 | 255 | 1194 | 19687 | 10.66 |
| 11 | 2.09 | GOTERM_CC_DIRECT | endoplasmic reticulum | 34 | 255 | 1712 | 19687 | 12.50 |
| 11 | 2.09 | UP_KW_CELLULAR_COMPONENT | Endoplasmic reticulum | 29 | 240 | 1381 | 17335 | 10.66 |
| 12 | 2.03 | REACTOME_PATHWAY | Intra-Golgi and retrograde Golgi-to-ER traffic | 14 | 176 | 208 | 11462 | 5.15 |
| 12 | 2.03 | REACTOME_PATHWAY | Golgi-to-ER retrograde transport | 7 | 176 | 137 | 11462 | 2.57 |
| 12 | 2.03 | REACTOME_PATHWAY | COPI-dependent Golgi-to-ER retrograde traffic | 5 | 176 | 103 | 11462 | 1.84 |
| 12 | 2.03 | REACTOME_PATHWAY | MHC class II antigen presentation | 4 | 176 | 128 | 11462 | 1.47 |
| 13 | 1.92 | REACTOME_PATHWAY | Signaling by Rho GTPases | 21 | 176 | 708 | 11462 | 7.72 |
| 13 | 1.92 | REACTOME_PATHWAY | Signaling by Rho GTPases, Miro GTPases and RHOBTB3 | 21 | 176 | 724 | 11462 | 7.72 |
| 13 | 1.92 | REACTOME_PATHWAY | Signal Transduction | 51 | 176 | 2613 | 11462 | 18.75 |
| 14 | 1.79 | UP_SEQ_FEATURE | CROSSLNK:Glycyl lysine isopeptide (Lys-Gly) (interchain with G-Cter in SUMO2) | 30 | 261 | 1272 | 19119 | 11.03 |
| 14 | 1.79 | UP_KW_PTM | Isopeptide bond | 40 | 208 | 1895 | 14032 | 14.71 |
| 14 | 1.79 | UP_KW_PTM | Ubl conjugation | 52 | 208 | 2781 | 14032 | 19.12 |
| 15 | 1.70 | UP_KW_MOLECULAR_FUNCTION | Helicase | 8 | 165 | 132 | 11620 | 2.94 |
| 15 | 1.70 | GOTERM_MF_DIRECT | helicase activity | 8 | 252 | 144 | 18532 | 2.94 |
| 15 | 1.70 | GOTERM_MF_DIRECT | RNA helicase activity | 6 | 252 | 77 | 18532 | 2.21 |
| 15 | 1.70 | UP_SEQ_FEATURE | DOMAIN:Helicase ATP-binding | 7 | 261 | 116 | 19119 | 2.57 |
| 15 | 1.70 | UP_SEQ_FEATURE | DOMAIN:Helicase C-terminal | 6 | 261 | 108 | 19119 | 2.21 |
| 15 | 1.70 | GOTERM_MF_DIRECT | ATP hydrolysis activity | 12 | 252 | 431 | 18532 | 4.41 |
| 15 | 1.70 | UP_SEQ_FEATURE | MOTIF:DEAD box | 3 | 261 | 34 | 19119 | 1.10 |
| 15 | 1.70 | UP_SEQ_FEATURE | MOTIF:Q motif | 3 | 261 | 38 | 19119 | 1.10 |
| 15 | 1.70 | GOTERM_MF_DIRECT | DNA helicase activity | 3 | 252 | 42 | 18532 | 1.10 |

| Cluster | Cluster Enrichment Score | Category | Term | Count | List Total | Pop Hits | Pop Total | % |
|---|---|---|---|---|---|---|---|---|
| 15 | 1.70 | GOTERM_MF_DIRECT | nucleic acid binding | 13 | 252 | 606 | 18532 | 4.78 |
| 16 | 1.68 | REACTOME_PATHWAY | Interleukin-12 signaling | 5 | 176 | 49 | 11462 | 1.84 |
| 16 | 1.68 | REACTOME_PATHWAY | Interleukin-12 family signaling | 5 | 176 | 59 | 11462 | 1.84 |
| 16 | 1.68 | REACTOME_PATHWAY | Gene and protein expression by JAK-STAT signaling after Interleukin-12 stimulation | 4 | 176 | 40 | 11462 | 1.47 |
| 16 | 1.68 | REACTOME_PATHWAY | Signaling by Interleukins | 12 | 176 | 468 | 11462 | 4.41 |
| 17 | 1.63 | GOTERM_CC_DIRECT | endoplasmic reticulum-Golgi intermediate compartment membrane | 6 | 255 | 81 | 19687 | 2.21 |
| 17 | 1.63 | REACTOME_PATHWAY | COPII-mediated vesicle transport | 6 | 176 | 71 | 11462 | 2.21 |
| 17 | 1.63 | REACTOME_PATHWAY | Cargo concentration in the ER | 3 | 176 | 33 | 11462 | 1.10 |
| 17 | 1.63 | GOTERM_CC_DIRECT | ER to Golgi transport vesicle membrane | 3 | 255 | 61 | 19687 | 1.10 |
| 18 | 1.57 | GOTERM_BP_DIRECT | GPI anchor biosynthetic process | 4 | 247 | 33 | 18757 | 1.47 |
| 18 | 1.57 | REACTOME_PATHWAY | Post-translational modification: synthesis of GPI-anchored proteins | 6 | 176 | 95 | 11462 | 2.21 |
| 18 | 1.57 | WIKIPATHWAYS | Glycosylphosphatidyl inositol anchor pathway | 3 | 138 | 26 | 8952 | 1.10 |
| 18 | 1.57 | KEGG_PATHWAY | Glycosylphosphatidylinositol (GPI)-anchor biosynthesis | 3 | 120 | 30 | 8506 | 1.10 |
| 19 | 1.50 | REACTOME_PATHWAY | Gene expression (Transcription) | 39 | 176 | 1598 | 11462 | 14.34 |
| 19 | 1.50 | REACTOME_PATHWAY | Generic Transcription Pathway | 26 | 176 | 1253 | 11462 | 9.56 |
| 19 | 1.50 | REACTOME_PATHWAY | RNA Polymerase II Transcription | 28 | 176 | 1375 | 11462 | 10.29 |
| 20 | 1.49 | REACTOME_PATHWAY | Nef Mediated CD4 Down-regulation | 3 | 176 | 10 | 11462 | 1.10 |
| 20 | 1.49 | REACTOME_PATHWAY | Nef-mediates down modulation of cell surface receptors by recruiting them to clathrin adapters | 3 | 176 | 23 | 11462 | 1.10 |
| 20 | 1.49 | REACTOME_PATHWAY | The role of Nef in HIV-1 | 3 | 176 | 30 | 11462 | 1.10 |

| Cluster | Cluster Enrichment Score | Category | Term | Count | List Total | Pop Hits | Pop Total | % |
|---|---|---|---|---|---|---|---|---|
| | | | replication and disease pathogenesis | | | | | |
| 21 | 1.45 | UP_KW_BIOLOGICAL_PROCESS | Transcription | 49 | 163 | 2460 | 11213 | 18.01 |
| 21 | 1.45 | UP_KW_MOLECULAR_FUNCTION | Activator | 19 | 165 | 723 | 11620 | 6.99 |
| 21 | 1.45 | UP_KW_BIOLOGICAL_PROCESS | Transcription regulation | 46 | 163 | 2393 | 11213 | 16.91 |
| 21 | 1.45 | GOTERM_BP_DIRECT | regulation of transcription by RNA polymerase II | 25 | 247 | 1618 | 18757 | 9.19 |
| 22 | 1.43 | WIKIPATHWAYS | miR targeted genes in lymphocytes | 14 | 138 | 423 | 8952 | 5.15 |
| 22 | 1.43 | WIKIPATHWAYS | miR targeted genes in leukocytes | 6 | 138 | 127 | 8952 | 2.21 |
| 22 | 1.43 | WIKIPATHWAYS | miR targeted genes in muscle cell | 10 | 138 | 352 | 8952 | 3.68 |
| 23 | 1.36 | UP_SEQ_FEATURE | REPEAT:Spectrin 3 | 3 | 261 | 22 | 19119 | 1.10 |
| 23 | 1.36 | UP_SEQ_FEATURE | REPEAT:Spectrin 1 | 3 | 261 | 26 | 19119 | 1.10 |
| 23 | 1.36 | UP_SEQ_FEATURE | REPEAT:Spectrin 2 | 3 | 261 | 26 | 19119 | 1.10 |
| 24 | 1.31 | GOTERM_CC_DIRECT | cortical actin cytoskeleton | 5 | 255 | 58 | 19687 | 1.84 |
| 24 | 1.31 | GOTERM_MF_DIRECT | actin binding | 11 | 252 | 400 | 18532 | 4.04 |
| 24 | 1.31 | UP_KW_MOLECULAR_FUNCTION | Actin-binding | 8 | 165 | 288 | 11620 | 2.94 |
| 24 | 1.31 | GOTERM_MF_DIRECT | actin filament binding | 6 | 252 | 211 | 18532 | 2.21 |
| 25 | 1.30 | UP_SEQ_FEATURE | DOMAIN:RanBD1 1 | 3 | 261 | 8 | 19119 | 1.10 |
| 25 | 1.30 | UP_SEQ_FEATURE | DOMAIN:RanBD1 2 | 3 | 261 | 8 | 19119 | 1.10 |
| 25 | 1.30 | GOTERM_BP_DIRECT | NLS-bearing protein import into nucleus | 4 | 247 | 27 | 18757 | 1.47 |
| 25 | 1.30 | UP_SEQ_FEATURE | DOMAIN:RanBD1 | 3 | 261 | 12 | 19119 | 1.10 |
| 25 | 1.30 | UP_SEQ_FEATURE | REPEAT:TPR | 5 | 261 | 97 | 19119 | 1.84 |
| 25 | 1.30 | UP_SEQ_FEATURE | REPEAT:TPR 1 | 6 | 261 | 159 | 19119 | 2.21 |
| 25 | 1.30 | UP_SEQ_FEATURE | REPEAT:TPR 2 | 6 | 261 | 159 | 19119 | 2.21 |
| 25 | 1.30 | UP_SEQ_FEATURE | REPEAT:TPR 3 | 5 | 261 | 145 | 19119 | 1.84 |
| 25 | 1.30 | UP_SEQ_FEATURE | REPEAT:TPR 7 | 3 | 261 | 65 | 19119 | 1.10 |
| 25 | 1.30 | UP_SEQ_FEATURE | REPEAT:TPR 6 | 3 | 261 | 76 | 19119 | 1.10 |
| 25 | 1.30 | UP_SEQ_FEATURE | REPEAT:TPR 5 | 3 | 261 | 81 | 19119 | 1.10 |

| Cluster | Cluster Enrichment Score | Category | Term | Count | List Total | Pop Hits | Pop Total | % |
|---|---|---|---|---|---|---|---|---|
| 25 | 1.30 | UP_SEQ_FEATURE | REPEAT:TPR 4 | 3 | 261 | 100 | 19119 | 1.10 |
| 26 | 1.22 | REACTOME_PATHWAY | Developmental Biology | 35 | 176 | 1507 | 11462 | 12.87 |
| 26 | 1.22 | KEGG_PATHWAY | Endocytosis | 8 | 120 | 249 | 8506 | 2.94 |
| 26 | 1.22 | REACTOME_PATHWAY | Axon guidance | 13 | 176 | 541 | 11462 | 4.78 |
| 26 | 1.22 | REACTOME_PATHWAY | Nervous system development | 13 | 176 | 566 | 11462 | 4.78 |
| 27 | 1.21 | REACTOME_PATHWAY | Butyrate Response Factor 1 (BRF1) binds and destabilizes mRNA | 3 | 176 | 17 | 11462 | 1.10 |
| 27 | 1.21 | REACTOME_PATHWAY | KSRP (KHSRP) binds and destabilizes mRNA | 3 | 176 | 17 | 11462 | 1.10 |
| 27 | 1.21 | REACTOME_PATHWAY | Regulation of mRNA stability by proteins that bind AU-rich elements | 3 | 176 | 75 | 11462 | 1.10 |
| 28 | 1.21 | GOTERM_MF_DIRECT | ubiquitin-like ligase-substrate adaptor activity | 6 | 252 | 133 | 18532 | 2.21 |
| 28 | 1.21 | UP_SEQ_FEATURE | REPEAT:Kelch 5 | 4 | 261 | 66 | 19119 | 1.47 |
| 28 | 1.21 | UP_SEQ_FEATURE | REPEAT:Kelch 4 | 4 | 261 | 70 | 19119 | 1.47 |
| 28 | 1.21 | UP_SEQ_FEATURE | REPEAT:Kelch 1 | 4 | 261 | 71 | 19119 | 1.47 |
| 28 | 1.21 | UP_SEQ_FEATURE | REPEAT:Kelch 2 | 4 | 261 | 71 | 19119 | 1.47 |
| 28 | 1.21 | UP_SEQ_FEATURE | REPEAT:Kelch 3 | 4 | 261 | 71 | 19119 | 1.47 |
| 29 | 1.20 | REACTOME_PATHWAY | Nuclear RNA decay | 4 | 176 | 35 | 11462 | 1.47 |
| 29 | 1.20 | GOTERM_CC_DIRECT | nuclear exosome (RNase complex) | 3 | 255 | 16 | 19687 | 1.10 |
| 29 | 1.20 | UP_KW_BIOLOGICAL_PROCESS | rRNA processing | 6 | 163 | 105 | 11213 | 2.21 |
| 29 | 1.20 | GOTERM_CC_DIRECT | exosome (RNase complex) | 3 | 255 | 17 | 19687 | 1.10 |
| 29 | 1.20 | GOTERM_BP_DIRECT | rRNA processing | 7 | 247 | 166 | 18757 | 2.57 |
| 29 | 1.20 | KEGG_PATHWAY | RNA degradation | 4 | 120 | 78 | 8506 | 1.47 |
| 29 | 1.20 | REACTOME_PATHWAY | rRNA processing in the nucleus and cytosol | 5 | 176 | 194 | 11462 | 1.84 |
| 29 | 1.20 | REACTOME_PATHWAY | rRNA processing | 5 | 176 | 211 | 11462 | 1.84 |
| 29 | 1.20 | REACTOME_PATHWAY | Major pathway of rRNA processing in the nucleolus and cytosol | 4 | 176 | 184 | 11462 | 1.47 |
| 30 | 1.17 | GOTERM_BP_DIRECT | positive regulation of | 5 | 247 | 71 | 18757 | 1.84 |

| Cluster | Cluster Enrichment Score | Category | Term | Count | List Total | Pop Hits | Pop Total | % |
|---|---|---|---|---|---|---|---|---|
| | | | TORC1 signaling | | | | | |
| 30 | 1.17 | GOTERM_BP_DIRECT | cellular response to amino acid starvation | 4 | 247 | 68 | 18757 | 1.47 |
| 30 | 1.17 | KEGG_PATHWAY | mTOR signaling pathway | 4 | 120 | 155 | 8506 | 1.47 |
| 31 | 1.15 | REACTOME_PATHWAY | Signaling by NOTCH | 11 | 176 | 230 | 11462 | 4.04 |
| 31 | 1.15 | REACTOME_PATHWAY | Signaling by NOTCH3 | 4 | 176 | 48 | 11462 | 1.47 |
| 31 | 1.15 | GOTERM_BP_DIRECT | positive regulation of cell growth | 4 | 247 | 88 | 18757 | 1.47 |
| 31 | 1.15 | REACTOME_PATHWAY | Diseases of signal transduction by growth factor receptors and second messengers | 11 | 176 | 458 | 11462 | 4.04 |
| 31 | 1.15 | REACTOME_PATHWAY | ESR-mediated signaling | 6 | 176 | 222 | 11462 | 2.21 |
| 31 | 1.15 | REACTOME_PATHWAY | Signaling by Nuclear Receptors | 7 | 176 | 296 | 11462 | 2.57 |
| 32 | 1.04 | WIKIPATHWAYS | TGF beta receptor signaling | 5 | 138 | 55 | 8952 | 1.84 |
| 32 | 1.04 | WIKIPATHWAYS | TGF beta receptor signaling in skeletal dysplasias | 5 | 138 | 59 | 8952 | 1.84 |
| 32 | 1.04 | WIKIPATHWAYS | Embryonic stem cell pluripotency pathways | 5 | 138 | 117 | 8952 | 1.84 |
| 32 | 1.04 | WIKIPATHWAYS | Pluripotent stem cell differentiation pathway | 3 | 138 | 48 | 8952 | 1.10 |
| 32 | 1.04 | KEGG_PATHWAY | Signaling pathways regulating pluripotency of stem cells | 4 | 120 | 143 | 8506 | 1.47 |
| 32 | 1.04 | WIKIPATHWAYS | Pleural mesothelioma | 6 | 138 | 438 | 8952 | 2.21 |
| 33 | 1.04 | REACTOME_PATHWAY | Clathrin-mediated endocytosis | 8 | 176 | 148 | 11462 | 2.94 |
| 33 | 1.04 | UP_KW_BIOLOGICAL_PROCESS | Endocytosis | 4 | 163 | 130 | 11213 | 1.47 |
| 33 | 1.04 | GOTERM_BP_DIRECT | endocytosis | 5 | 247 | 225 | 18757 | 1.84 |
| 34 | 1.03 | WIKIPATHWAYS | MET in type 1 papillary renal cell carcinoma | 5 | 138 | 59 | 8952 | 1.84 |
| 34 | 1.03 | KEGG_PATHWAY | Focal adhesion | 7 | 120 | 202 | 8506 | 2.57 |
| 34 | 1.03 | WIKIPATHWAYS | Integrin mediated cell adhesion | 5 | 138 | 102 | 8952 | 1.84 |
| 34 | 1.03 | KEGG_PATHWAY | Renal cell carcinoma | 4 | 120 | 69 | 8506 | 1.47 |

| Cluster | Cluster Enrichment Score | Category | Term | Count | List Total | Pop Hits | Pop Total | % |
|---|---|---|---|---|---|---|---|---|
| 34 | 1.03 | WIKIPATHWAYS | Focal adhesion | 7 | 138 | 199 | 8952 | 2.57 |
| 34 | 1.03 | WIKIPATHWAYS | BDNF signaling in neurodevelopment | 5 | 138 | 119 | 8952 | 1.84 |
| 34 | 1.03 | GOTERM_BP_DIRECT | phosphatidylinositol 3-kinase/protein kinase B signal transduction | 4 | 247 | 93 | 18757 | 1.47 |
| 34 | 1.03 | KEGG_PATHWAY | Rap1 signaling pathway | 6 | 120 | 210 | 8506 | 2.21 |
| 34 | 1.03 | KEGG_PATHWAY | Neurotrophin signaling pathway | 4 | 120 | 119 | 8506 | 1.47 |
| 34 | 1.03 | WIKIPATHWAYS | Androgen receptor network in prostate cancer | 4 | 138 | 113 | 8952 | 1.47 |
| 35 | 1.00 | REACTOME_PATHWAY | Regulation of CDH1 posttranslational processing and trafficking to plasma membrane | 5 | 176 | 32 | 11462 | 1.84 |
| 35 | 1.00 | REACTOME_PATHWAY | PD-L1(CD274) glycosylation and translocation to plasma membrane | 3 | 176 | 21 | 11462 | 1.10 |
| 35 | 1.00 | REACTOME_PATHWAY | Regulation of CDH1 Expression and Function | 7 | 176 | 202 | 11462 | 2.57 |
| 35 | 1.00 | REACTOME_PATHWAY | Regulation of Expression and Function of Type I Classical Cadherins | 7 | 176 | 202 | 11462 | 2.57 |
| 35 | 1.00 | REACTOME_PATHWAY | Regulation of Homotypic Cell-Cell Adhesion | 7 | 176 | 217 | 11462 | 2.57 |
| 35 | 1.00 | REACTOME_PATHWAY | Regulation of T cell activation by CD28 family | 8 | 176 | 273 | 11462 | 2.94 |
| 35 | 1.00 | REACTOME_PATHWAY | Co-inhibition by PD-1 | 7 | 176 | 223 | 11462 | 2.57 |
| 35 | 1.00 | REACTOME_PATHWAY | Cell-Cell communication | 9 | 176 | 336 | 11462 | 3.31 |
| 35 | 1.00 | REACTOME_PATHWAY | Adherens junctions interactions | 7 | 176 | 240 | 11462 | 2.57 |
| 35 | 1.00 | REACTOME_PATHWAY | Regulation of PD-L1(CD274) expression | 6 | 176 | 195 | 11462 | 2.21 |
| 35 | 1.00 | REACTOME_PATHWAY | Cell-cell junction organization | 7 | 176 | 272 | 11462 | 2.57 |
| 35 | 1.00 | REACTOME_PATHWAY | Cell junction organization | 7 | 176 | 299 | 11462 | 2.57 |
| 35 | 1.00 | REACTOME_PATHWAY | Regulation of PD-L1(CD274) Post-translational modification | 3 | 176 | 86 | 11462 | 1.10 |

| Cluster | Cluster Enrichment Score | Category | Term | Count | List Total | Pop Hits | Pop Total | % |
|---|---|---|---|---|---|---|---|---|
| 36 | 1.00 | REACTOME_PATHWAY | SARS-CoV-1 activates/ modulates innate immune responses | 4 | 176 | 43 | 11462 | 1.47 |
| 36 | 1.00 | REACTOME_PATHWAY | SARS-CoV-1 Infection | 5 | 176 | 145 | 11462 | 1.84 |
| 36 | 1.00 | REACTOME_PATHWAY | SARS-CoV-1-host interactions | 4 | 176 | 100 | 11462 | 1.47 |
| 37 | 1.00 | WIKIPATHWAYS | Cytosolic DNA sensing pathway | 5 | 138 | 73 | 8952 | 1.84 |
| 37 | 1.00 | KEGG_PATHWAY | NOD-like receptor signaling pathway | 7 | 120 | 186 | 8506 | 2.57 |
| 37 | 1.00 | REACTOME_PATHWAY | DDX58/IFIH1-mediated induction of interferon-alpha/beta | 4 | 176 | 83 | 11462 | 1.47 |
| 37 | 1.00 | WIKIPATHWAYS | Novel intracellular components of RIG I like receptor pathway | 3 | 138 | 60 | 8952 | 1.10 |
| 37 | 1.00 | KEGG_PATHWAY | RIG-I-like receptor signaling pathway | 3 | 120 | 72 | 8506 | 1.10 |
| 38 | 0.99 | KEGG_PATHWAY | FoxO signaling pathway | 6 | 120 | 131 | 8506 | 2.21 |
| 38 | 0.99 | WIKIPATHWAYS | Perturbations to host cell autophagy induced by SARS CoV 2 proteins | 3 | 138 | 44 | 8952 | 1.10 |
| 38 | 0.99 | KEGG_PATHWAY | Autophagy - animal | 5 | 120 | 167 | 8506 | 1.84 |
| 39 | 0.98 | GOTERM_BP_DIRECT | protein N-linked glycosylation via asparagine | 4 | 247 | 24 | 18757 | 1.47 |
| 39 | 0.98 | KEGG_PATHWAY | N-Glycan biosynthesis | 4 | 120 | 55 | 8506 | 1.47 |
| 39 | 0.98 | WIKIPATHWAYS | N glycan biosynthesis | 4 | 138 | 57 | 8952 | 1.47 |
| 39 | 0.98 | GOTERM_MF_DIRECT | glycosyltransferase activity | 7 | 252 | 236 | 18532 | 2.57 |
| 39 | 0.98 | UP_KW_MOLECULAR_FUNCTION | Glycosyltransferase | 7 | 165 | 232 | 11620 | 2.57 |
| 39 | 0.98 | KEGG_PATHWAY | Various types of N-glycan biosynthesis | 3 | 120 | 43 | 8506 | 1.10 |
| 39 | 0.98 | REACTOME_PATHWAY | Maturation of spike protein | 3 | 176 | 41 | 11462 | 1.10 |
| 39 | 0.98 | GOTERM_BP_DIRECT | protein N-linked glycosylation | 3 | 247 | 62 | 18757 | 1.10 |
| 39 | 0.98 | REACTOME_PATHWAY | Translation of Structural Proteins | 3 | 176 | 61 | 11462 | 1.10 |
| 39 | 0.98 | REACTOME_PATHWAY | Late SARS-CoV-2 Infection | 3 | 176 | 74 | 11462 | 1.10 |

| Cluster | Cluster Enrichment Score | Category | Term | Count | List Total | Pop Hits | Pop Total | % |
|---|---|---|---|---|---|---|---|---|
| | | | Events | | | | | |
| 39 | 0.98 | KEGG_PATHWAY | Metabolic pathways | 20 | 120 | 1565 | 8506 | 7.35 |
| 40 | 0.97 | GOTERM_MF_DIRECT | DNA-directed RNA polymerase activity | 3 | 252 | 31 | 18532 | 1.10 |
| 40 | 0.97 | KEGG_PATHWAY | RNA polymerase | 3 | 120 | 34 | 8506 | 1.10 |
| 40 | 0.97 | GOTERM_CC_DIRECT | DNA-directed RNA polymerase complex | 3 | 255 | 38 | 19687 | 1.10 |
| 40 | 0.97 | UP_KW_CELLULAR_COMPONENT | DNA-directed RNA polymerase | 3 | 240 | 38 | 17335 | 1.10 |
| 40 | 0.97 | UP_KW_CELLULAR_COMPONENT | DNA-directed RNA polymerase | 3 | 240 | 38 | 17335 | 1.10 |
| 40 | 0.97 | WIKIPATHWAYS | Pyrimidine metabolism | 3 | 138 | 84 | 8952 | 1.10 |
| 41 | 0.92 | WIKIPATHWAYS | Cytosolic DNA sensing pathway | 5 | 138 | 73 | 8952 | 1.84 |
| 41 | 0.92 | REACTOME_PATHWAY | Cytosolic sensors of pathogen-associated DNA | 4 | 176 | 64 | 11462 | 1.47 |
| 41 | 0.92 | KEGG_PATHWAY | Cytosolic DNA-sensing pathway | 4 | 120 | 82 | 8506 | 1.47 |
| 41 | 0.92 | GOTERM_BP_DIRECT | defense response to virus | 6 | 247 | 250 | 18757 | 2.21 |
| 41 | 0.92 | GOTERM_BP_DIRECT | innate immune response | 9 | 247 | 610 | 18757 | 3.31 |
| 42 | 0.86 | WIKIPATHWAYS | Hallmark of cancer sustaining proliferative signaling | 6 | 138 | 98 | 8952 | 2.21 |
| 42 | 0.86 | WIKIPATHWAYS | Wnt beta catenin signaling in leukemia | 3 | 138 | 26 | 8952 | 1.10 |
| 42 | 0.86 | REACTOME_PATHWAY | Signaling by WNT | 6 | 176 | 318 | 11462 | 2.21 |
| 42 | 0.86 | REACTOME_PATHWAY | TCF dependent signaling in response to WNT | 4 | 176 | 220 | 11462 | 1.47 |
| 43 | 0.84 | GOTERM_BP_DIRECT | methylation | 6 | 247 | 204 | 18757 | 2.21 |
| 43 | 0.84 | GOTERM_MF_DIRECT | methyltransferase activity | 6 | 252 | 199 | 18532 | 2.21 |
| 43 | 0.84 | UP_KW_MOLECULAR_FUNCTION | Methyltransferase | 6 | 165 | 199 | 11620 | 2.21 |
| 43 | 0.84 | UP_SEQ_FEATURE | DOMAIN:SET | 3 | 261 | 52 | 19119 | 1.10 |
| 44 | 0.83 | REACTOME_PATHWAY | Macroautophagy | 6 | 176 | 143 | 11462 | 2.21 |
| 44 | 0.83 | REACTOME_PATHWAY | Autophagy | 6 | 176 | 158 | 11462 | 2.21 |
| 44 | 0.83 | GOTERM_BP_DIRECT | autophagosome assembly | 4 | 247 | 98 | 18757 | 1.47 |

| Cluster | Cluster Enrichment Score | Category | Term | Count | List Total | Pop Hits | Pop Total | % |
|---|---|---|---|---|---|---|---|---|
| 44 | 0.83 | KEGG_PATHWAY | Autophagy - animal | 5 | 120 | 167 | 8506 | 1.84 |
| 44 | 0.83 | REACTOME_PATHWAY | Selective autophagy | 3 | 176 | 88 | 11462 | 1.10 |
| 45 | 0.83 | UP_SEQ_FEATURE | DOMAIN:PH | 9 | 261 | 280 | 19119 | 3.31 |
| 45 | 0.83 | UP_SEQ_FEATURE | DOMAIN:DH | 4 | 261 | 71 | 19119 | 1.47 |
| 45 | 0.83 | UP_KW_MOLECULAR_FUNCTION | Guanine-nucleotide releasing factor | 6 | 165 | 168 | 11620 | 2.21 |
| 45 | 0.83 | GOTERM_MF_DIRECT | guanyl-nucleotide exchange factor activity | 7 | 252 | 233 | 18532 | 2.57 |
| 45 | 0.83 | REACTOME_PATHWAY | p75 NTR receptor-mediated signalling | 4 | 176 | 99 | 11462 | 1.47 |
| 45 | 0.83 | GOTERM_BP_DIRECT | regulation of small GTPase mediated signal transduction | 4 | 247 | 120 | 18757 | 1.47 |
| 45 | 0.83 | REACTOME_PATHWAY | Death Receptor Signaling | 5 | 176 | 156 | 11462 | 1.84 |
| 45 | 0.83 | REACTOME_PATHWAY | NRAGE signals death through JNK | 3 | 176 | 60 | 11462 | 1.10 |
| 45 | 0.83 | REACTOME_PATHWAY | Cell death signalling via NRAGE, NRIF and NADE | 3 | 176 | 77 | 11462 | 1.10 |
| 45 | 0.83 | REACTOME_PATHWAY | G alpha (12/13) signalling events | 3 | 176 | 81 | 11462 | 1.10 |
| 46 | 0.82 | KEGG_PATHWAY | PD-L1 expression and PD-1 checkpoint pathway in cancer | 7 | 120 | 89 | 8506 | 2.57 |
| 46 | 0.82 | KEGG_PATHWAY | Shigellosis | 11 | 120 | 249 | 8506 | 4.04 |
| 46 | 0.82 | KEGG_PATHWAY | Yersinia infection | 8 | 120 | 136 | 8506 | 2.94 |
| 46 | 0.82 | KEGG_PATHWAY | Pancreatic cancer | 6 | 120 | 76 | 8506 | 2.21 |
| 46 | 0.82 | WIKIPATHWAYS | Pancreatic adenocarcinoma pathway | 6 | 138 | 89 | 8952 | 2.21 |
| 46 | 0.82 | WIKIPATHWAYS | IL17 signaling | 4 | 138 | 32 | 8952 | 1.47 |
| 46 | 0.82 | KEGG_PATHWAY | Prostate cancer | 6 | 120 | 105 | 8506 | 2.21 |
| 46 | 0.82 | KEGG_PATHWAY | Epithelial cell signaling in Helicobacter pylori infection | 5 | 120 | 70 | 8506 | 1.84 |
| 46 | 0.82 | WIKIPATHWAYS | Hallmark of cancer sustaining proliferative signaling | 6 | 138 | 98 | 8952 | 2.21 |
| 46 | 0.82 | WIKIPATHWAYS | Catabolism of skeletal | 4 | 138 | 40 | 8952 | 1.47 |

| Cluster | Cluster Enrichment Score | Category | Term | Count | List Total | Pop Hits | Pop Total | % |
|---|---|---|---|---|---|---|---|---|
| | | | muscle in cachexia | | | | | |
| 46 | 0.82 | KEGG_PATHWAY | Hepatitis C | 7 | 120 | 158 | 8506 | 2.57 |
| 46 | 0.82 | WIKIPATHWAYS | Prolactin signaling | 5 | 138 | 74 | 8952 | 1.84 |
| 46 | 0.82 | KEGG_PATHWAY | T cell receptor signaling pathway | 6 | 120 | 121 | 8506 | 2.21 |
| 46 | 0.82 | WIKIPATHWAYS | T cell receptor signaling | 5 | 138 | 75 | 8952 | 1.84 |
| 46 | 0.82 | KEGG_PATHWAY | Human T-cell leukemia virus 1 infection | 8 | 120 | 223 | 8506 | 2.94 |
| 46 | 0.82 | WIKIPATHWAYS | Chemokine signaling | 7 | 138 | 165 | 8952 | 2.57 |
| 46 | 0.82 | WIKIPATHWAYS | Androgen receptor signaling | 5 | 138 | 87 | 8952 | 1.84 |
| 46 | 0.82 | WIKIPATHWAYS | T cell activation SARS CoV 2 | 5 | 138 | 88 | 8952 | 1.84 |
| 46 | 0.82 | KEGG_PATHWAY | NOD-like receptor signaling pathway | 7 | 120 | 186 | 8506 | 2.57 |
| 46 | 0.82 | WIKIPATHWAYS | NRP1 triggered signaling in pancreatic cancer | 4 | 138 | 53 | 8952 | 1.47 |
| 46 | 0.82 | WIKIPATHWAYS | B cell receptor signaling | 5 | 138 | 92 | 8952 | 1.84 |
| 46 | 0.82 | WIKIPATHWAYS | Modulators of TCR signaling and T cell activation | 4 | 138 | 61 | 8952 | 1.47 |
| 46 | 0.82 | WIKIPATHWAYS | IL19 signaling | 3 | 138 | 28 | 8952 | 1.10 |
| 46 | 0.82 | WIKIPATHWAYS | T cell antigen receptor TCR pathway during Staphylococcus aureus infection | 4 | 138 | 62 | 8952 | 1.47 |
| 46 | 0.82 | REACTOME_PATHWAY | Turbulent (oscillatory, disturbed) flow shear stress activates signaling by PIEZO1 and integrins in endothelial cells | 3 | 176 | 29 | 11462 | 1.10 |
| 46 | 0.82 | KEGG_PATHWAY | Human immunodeficiency virus 1 infection | 7 | 120 | 212 | 8506 | 2.57 |
| 46 | 0.82 | WIKIPATHWAYS | AGE RAGE pathway | 4 | 138 | 66 | 8952 | 1.47 |
| 46 | 0.82 | WIKIPATHWAYS | MAPK signaling | 8 | 138 | 246 | 8952 | 2.94 |
| 46 | 0.82 | WIKIPATHWAYS | RAC1 PAK1 p38 MMP2 pathway | 4 | 138 | 68 | 8952 | 1.47 |
| 46 | 0.82 | WIKIPATHWAYS | Resistin as a regulator of inflammation | 3 | 138 | 33 | 8952 | 1.10 |

| Cluster | Cluster Enrichment Score | Category | Term | Count | List Total | Pop Hits | Pop Total | % |
|---|---|---|---|---|---|---|---|---|
| 46 | 0.82 | KEGG_PATHWAY | Human papillomavirus infection | 9 | 120 | 331 | 8506 | 3.31 |
| 46 | 0.82 | KEGG_PATHWAY | Human cytomegalovirus infection | 7 | 120 | 224 | 8506 | 2.57 |
| 46 | 0.82 | WIKIPATHWAYS | Leptin signaling | 4 | 138 | 71 | 8952 | 1.47 |
| 46 | 0.82 | REACTOME_PATHWAY | TCR signaling | 5 | 176 | 115 | 11462 | 1.84 |
| 46 | 0.82 | WIKIPATHWAYS | BDNF signaling in neurodevelopment | 5 | 138 | 119 | 8952 | 1.84 |
| 46 | 0.82 | WIKIPATHWAYS | Neovascularization processes | 3 | 138 | 37 | 8952 | 1.10 |
| 46 | 0.82 | WIKIPATHWAYS | Thymic stromal lymphopoietin TSLP signaling | 3 | 138 | 37 | 8952 | 1.10 |
| 46 | 0.82 | REACTOME_PATHWAY | Extra-nuclear estrogen signaling | 4 | 176 | 76 | 11462 | 1.47 |
| 46 | 0.82 | KEGG_PATHWAY | Osteoclast differentiation | 5 | 120 | 134 | 8506 | 1.84 |
| 46 | 0.82 | KEGG_PATHWAY | Apoptosis | 5 | 120 | 136 | 8506 | 1.84 |
| 46 | 0.82 | KEGG_PATHWAY | MAPK signaling pathway | 8 | 120 | 299 | 8506 | 2.94 |
| 46 | 0.82 | KEGG_PATHWAY | Measles | 5 | 120 | 138 | 8506 | 1.84 |
| 46 | 0.82 | WIKIPATHWAYS | Acute viral myocarditis | 4 | 138 | 85 | 8952 | 1.47 |
| 46 | 0.82 | KEGG_PATHWAY | Th1 and Th2 cell differentiation | 4 | 120 | 93 | 8506 | 1.47 |
| 46 | 0.82 | WIKIPATHWAYS | IL11 signaling | 3 | 138 | 44 | 8952 | 1.10 |
| 46 | 0.82 | WIKIPATHWAYS | Fibrin complement receptor 3 signaling | 3 | 138 | 44 | 8952 | 1.10 |
| 46 | 0.82 | KEGG_PATHWAY | Pathogenic Escherichia coli infection | 6 | 120 | 199 | 8506 | 2.21 |
| 46 | 0.82 | WIKIPATHWAYS | IL26 signaling | 3 | 138 | 47 | 8952 | 1.10 |
| 46 | 0.82 | KEGG_PATHWAY | Chagas disease | 4 | 120 | 101 | 8506 | 1.47 |
| 46 | 0.82 | REACTOME_PATHWAY | Response of endothelial cells to shear stress | 4 | 176 | 93 | 11462 | 1.47 |
| 46 | 0.82 | KEGG_PATHWAY | Rap1 signaling pathway | 6 | 120 | 210 | 8506 | 2.21 |
| 46 | 0.82 | WIKIPATHWAYS | IL4 signaling | 3 | 138 | 50 | 8952 | 1.10 |
| 46 | 0.82 | WIKIPATHWAYS | RANKL RANK signaling | 3 | 138 | 50 | 8952 | 1.10 |
| 46 | 0.82 | KEGG_PATHWAY | C-type lectin receptor | 4 | 120 | 104 | 8506 | 1.47 |

| Cluster | Cluster Enrichment Score | Category | Term | Count | List Total | Pop Hits | Pop Total | % |
|---|---|---|---|---|---|---|---|---|
| | | | signaling pathway | | | | | |
| 46 | 0.82 | KEGG_PATHWAY | Toll-like receptor signaling pathway | 4 | 120 | 106 | 8506 | 1.47 |
| 46 | 0.82 | WIKIPATHWAYS | Network map of SARS CoV 2 signaling | 7 | 138 | 254 | 8952 | 2.57 |
| 46 | 0.82 | KEGG_PATHWAY | HIF-1 signaling pathway | 4 | 120 | 109 | 8506 | 1.47 |
| 46 | 0.82 | KEGG_PATHWAY | Th17 cell differentiation | 4 | 120 | 109 | 8506 | 1.47 |
| 46 | 0.82 | WIKIPATHWAYS | Cardiac hypertrophic response | 3 | 138 | 54 | 8952 | 1.10 |
| 46 | 0.82 | WIKIPATHWAYS | IL1 signaling | 3 | 138 | 54 | 8952 | 1.10 |
| 46 | 0.82 | KEGG_PATHWAY | Chemical carcinogenesis - reactive oxygen species | 6 | 120 | 223 | 8506 | 2.21 |
| 46 | 0.82 | KEGG_PATHWAY | Toxoplasmosis | 4 | 120 | 112 | 8506 | 1.47 |
| 46 | 0.82 | REACTOME_PATHWAY | Cellular responses to mechanical stimuli | 4 | 176 | 103 | 11462 | 1.47 |
| 46 | 0.82 | WIKIPATHWAYS | Oncostatin M signaling | 3 | 138 | 56 | 8952 | 1.10 |
| 46 | 0.82 | KEGG_PATHWAY | Influenza A | 5 | 120 | 172 | 8506 | 1.84 |
| 46 | 0.82 | GOTERM_MF_DIRECT | ubiquitin protein ligase binding | 7 | 252 | 303 | 18532 | 2.57 |
| 46 | 0.82 | KEGG_PATHWAY | Neurotrophin signaling pathway | 4 | 120 | 119 | 8506 | 1.47 |
| 46 | 0.82 | KEGG_PATHWAY | Ras signaling pathway | 6 | 120 | 236 | 8506 | 2.21 |
| 46 | 0.82 | KEGG_PATHWAY | PI3K-Akt signaling pathway | 8 | 120 | 359 | 8506 | 2.94 |
| 46 | 0.82 | KEGG_PATHWAY | Acute myeloid leukemia | 3 | 120 | 67 | 8506 | 1.10 |
| 46 | 0.82 | REACTOME_PATHWAY | KEAP1-NFE2L2 pathway | 4 | 176 | 112 | 11462 | 1.47 |
| 46 | 0.82 | KEGG_PATHWAY | Tuberculosis | 5 | 120 | 180 | 8506 | 1.84 |
| 46 | 0.82 | KEGG_PATHWAY | Adipocytokine signaling pathway | 3 | 120 | 70 | 8506 | 1.10 |
| 46 | 0.82 | KEGG_PATHWAY | Salmonella infection | 6 | 120 | 248 | 8506 | 2.21 |
| 46 | 0.82 | KEGG_PATHWAY | Chemokine signaling pathway | 5 | 120 | 190 | 8506 | 1.84 |
| 46 | 0.82 | WIKIPATHWAYS | Acute myeloid leukemia | 3 | 138 | 68 | 8952 | 1.10 |
| 46 | 0.82 | WIKIPATHWAYS | Urotensin II mediated signaling | 3 | 138 | 69 | 8952 | 1.10 |

| Cluster | Cluster Enrichment Score | Category | Term | Count | List Total | Pop Hits | Pop Total | % |
|---|---|---|---|---|---|---|---|---|
| 46 | 0.82 | KEGG_PATHWAY | Chronic myeloid leukemia | 3 | 120 | 76 | 8506 | 1.10 |
| 46 | 0.82 | KEGG_PATHWAY | Alzheimer disease | 8 | 120 | 383 | 8506 | 2.94 |
| 46 | 0.82 | UP_SEQ_FEATURE | CROSSLNK:Glycyl lysine isopeptide (Lys-Gly) (interchain with G-Cter in ubiquitin) | 7 | 261 | 329 | 19119 | 2.57 |
| 46 | 0.82 | WIKIPATHWAYS | Ras signaling | 5 | 138 | 184 | 8952 | 1.84 |
| 46 | 0.82 | KEGG_PATHWAY | Epstein-Barr virus infection | 5 | 120 | 202 | 8506 | 1.84 |
| 46 | 0.82 | WIKIPATHWAYS | Head and neck squamous cell carcinoma | 3 | 138 | 74 | 8952 | 1.10 |
| 46 | 0.82 | GOTERM_BP_DIRECT | cytokine-mediated signaling pathway | 4 | 247 | 150 | 18757 | 1.47 |
| 46 | 0.82 | KEGG_PATHWAY | Fluid shear stress and atherosclerosis | 4 | 120 | 141 | 8506 | 1.47 |
| 46 | 0.82 | WIKIPATHWAYS | Ebola virus infection in host | 4 | 138 | 129 | 8952 | 1.47 |
| 46 | 0.82 | KEGG_PATHWAY | B cell receptor signaling pathway | 3 | 120 | 82 | 8506 | 1.10 |
| 46 | 0.82 | WIKIPATHWAYS | Focal adhesion PI3K Akt mTOR signaling | 7 | 138 | 304 | 8952 | 2.57 |
| 46 | 0.82 | GOTERM_BP_DIRECT | intracellular signal transduction | 9 | 247 | 494 | 18757 | 3.31 |
| 46 | 0.82 | KEGG_PATHWAY | Pathways in cancer | 10 | 120 | 532 | 8506 | 3.68 |
| 46 | 0.82 | GOTERM_MF_DIRECT | enzyme binding | 8 | 252 | 416 | 18532 | 2.94 |
| 46 | 0.82 | WIKIPATHWAYS | Measles virus infection | 4 | 138 | 136 | 8952 | 1.47 |
| 46 | 0.82 | GOTERM_BP_DIRECT | positive regulation of gene expression | 9 | 247 | 510 | 18757 | 3.31 |
| 46 | 0.82 | KEGG_PATHWAY | Chemical carcinogenesis - receptor activation | 5 | 120 | 216 | 8506 | 1.84 |
| 46 | 0.82 | REACTOME_PATHWAY | Cellular response to chemical stress | 5 | 176 | 198 | 11462 | 1.84 |
| 46 | 0.82 | KEGG_PATHWAY | Small cell lung cancer | 3 | 120 | 92 | 8506 | 1.10 |
| 46 | 0.82 | WIKIPATHWAYS | Cancer pathways | 10 | 138 | 507 | 8952 | 3.68 |
| 46 | 0.82 | REACTOME_PATHWAY | Nuclear events mediated by NFE2L2 | 3 | 176 | 86 | 11462 | 1.10 |
| 46 | 0.82 | WIKIPATHWAYS | Apoptosis | 3 | 138 | 88 | 8952 | 1.10 |

| Cluster | Cluster Enrichment Score | Category | Term | Count | List Total | Pop Hits | Pop Total | % |
|---|---|---|---|---|---|---|---|---|
| 46 | 0.82 | KEGG_PATHWAY | Hepatitis B | 4 | 120 | 162 | 8506 | 1.47 |
| 46 | 0.82 | WIKIPATHWAYS | Hepatitis B infection | 4 | 138 | 152 | 8952 | 1.47 |
| 46 | 0.82 | WIKIPATHWAYS | Corticotropin releasing hormone signaling | 3 | 138 | 93 | 8952 | 1.10 |
| 46 | 0.82 | WIKIPATHWAYS | PI3K Akt signaling | 7 | 138 | 340 | 8952 | 2.57 |
| 46 | 0.82 | WIKIPATHWAYS | Small cell lung cancer | 3 | 138 | 95 | 8952 | 1.10 |
| 46 | 0.82 | WIKIPATHWAYS | Toll like receptor signaling | 3 | 138 | 95 | 8952 | 1.10 |
| 46 | 0.82 | KEGG_PATHWAY | Insulin resistance | 3 | 120 | 108 | 8506 | 1.10 |
| 46 | 0.82 | KEGG_PATHWAY | Herpes simplex virus 1 infection | 4 | 120 | 182 | 8506 | 1.47 |
| 46 | 0.82 | WIKIPATHWAYS | MicroRNAs in cardiomyocyte hypertrophy | 3 | 138 | 104 | 8952 | 1.10 |
| 46 | 0.82 | KEGG_PATHWAY | TNF signaling pathway | 3 | 120 | 118 | 8506 | 1.10 |
| 46 | 0.82 | KEGG_PATHWAY | Sphingolipid signaling pathway | 3 | 120 | 121 | 8506 | 1.10 |
| 46 | 0.82 | KEGG_PATHWAY | Kaposi sarcoma-associated herpesvirus infection | 4 | 120 | 194 | 8506 | 1.47 |
| 46 | 0.82 | WIKIPATHWAYS | Gastrin signaling | 3 | 138 | 115 | 8952 | 1.10 |
| 46 | 0.82 | KEGG_PATHWAY | Relaxin signaling pathway | 3 | 120 | 129 | 8506 | 1.10 |
| 46 | 0.82 | KEGG_PATHWAY | Viral carcinogenesis | 4 | 120 | 203 | 8506 | 1.47 |
| 46 | 0.82 | GOTERM_BP_DIRECT | negative regulation of apoptotic process | 8 | 247 | 540 | 18757 | 2.94 |
| 46 | 0.82 | KEGG_PATHWAY | Lipid and atherosclerosis | 4 | 120 | 214 | 8506 | 1.47 |
| 46 | 0.82 | GOTERM_BP_DIRECT | inflammatory response | 7 | 247 | 469 | 18757 | 2.57 |
| 46 | 0.82 | KEGG_PATHWAY | Alcoholic liver disease | 3 | 120 | 143 | 8506 | 1.10 |
| 46 | 0.82 | KEGG_PATHWAY | cAMP signaling pathway | 4 | 120 | 224 | 8506 | 1.47 |
| 46 | 0.82 | WIKIPATHWAYS | Brain derived neurotrophic factor BDNF signaling | 3 | 138 | 137 | 8952 | 1.10 |
| 46 | 0.82 | KEGG_PATHWAY | Coronavirus disease - COVID-19 | 4 | 120 | 230 | 8506 | 1.47 |
| 46 | 0.82 | WIKIPATHWAYS | Regulation of toll like receptor signaling | 3 | 138 | 145 | 8952 | 1.10 |
| 46 | 0.82 | GOTERM_BP_DIRECT | immune response | 5 | 247 | 556 | 18757 | 1.84 |
| 47 | 0.81 | REACTOME_PATHWAY | MAPK family signaling | 10 | 176 | 313 | 11462 | 3.68 |

| Cluster | Cluster Enrichment Score | Category | Term | Count | List Total | Pop Hits | Pop Total | % |
|---|---|---|---|---|---|---|---|---|
| | | | cascades | | | | | |
| 47 | 0.81 | REACTOME_PATHWAY | RAF/MAP kinase cascade | 8 | 176 | 268 | 11462 | 2.94 |
| 47 | 0.81 | REACTOME_PATHWAY | MAPK1/MAPK3 signaling | 8 | 176 | 273 | 11462 | 2.94 |
| 47 | 0.81 | GOTERM_CC_DIRECT | cell junction | 3 | 255 | 210 | 19687 | 1.10 |
| 48 | 0.80 | GOTERM_BP_DIRECT | cell surface receptor protein tyrosine kinase signaling pathway | 5 | 247 | 124 | 18757 | 1.84 |
| 48 | 0.80 | KEGG_PATHWAY | Rap1 signaling pathway | 6 | 120 | 210 | 8506 | 2.21 |
| 48 | 0.80 | GOTERM_BP_DIRECT | positive regulation of ERK1 and ERK2 cascade | 5 | 247 | 207 | 18757 | 1.84 |
| 49 | 0.80 | GOTERM_MF_DIRECT | ubiquitin protein ligase activity | 9 | 252 | 379 | 18532 | 3.31 |
| 49 | 0.80 | GOTERM_BP_DIRECT | protein polyubiquitination | 5 | 247 | 161 | 18757 | 1.84 |
| 49 | 0.80 | UP_SEQ_FEATURE | ZN_FING:RING-type | 6 | 261 | 218 | 19119 | 2.21 |
| 50 | 0.77 | KEGG_PATHWAY | NOD-like receptor signaling pathway | 7 | 120 | 186 | 8506 | 2.57 |
| 50 | 0.77 | GOTERM_BP_DIRECT | positive regulation of canonical NF-kappaB signal transduction | 7 | 247 | 233 | 18757 | 2.57 |
| 50 | 0.77 | GOTERM_BP_DIRECT | positive regulation of NF-kappaB transcription factor activity | 4 | 247 | 94 | 18757 | 1.47 |
| 50 | 0.77 | REACTOME_PATHWAY | Nucleotide-binding domain, leucine rich repeat containing receptor (NLR) signaling pathways | 3 | 176 | 57 | 11462 | 1.10 |
| 50 | 0.77 | REACTOME_PATHWAY | Interleukin-1 family signaling | 4 | 176 | 142 | 11462 | 1.47 |
| 50 | 0.77 | GOTERM_BP_DIRECT | inflammatory response | 7 | 247 | 469 | 18757 | 2.57 |
| 51 | 0.75 | GOTERM_CC_DIRECT | autophagosome | 4 | 255 | 83 | 19687 | 1.47 |
| 51 | 0.75 | GOTERM_BP_DIRECT | autophagosome assembly | 4 | 247 | 98 | 18757 | 1.47 |
| 51 | 0.75 | UP_KW_BIOLOGICAL_PROCESS | Autophagy | 5 | 163 | 172 | 11213 | 1.84 |
| 51 | 0.75 | GOTERM_BP_DIRECT | autophagy | 5 | 247 | 221 | 18757 | 1.84 |
| 52 | 0.75 | REACTOME_PATHWAY | Signaling by NOTCH3 | 4 | 176 | 48 | 11462 | 1.47 |
| 52 | 0.75 | REACTOME_PATHWAY | NOTCH3 Activation and Transmission of Signal to | 3 | 176 | 25 | 11462 | 1.10 |

| Cluster | Cluster Enrichment Score | Category | Term | Count | List Total | Pop Hits | Pop Total | % |
|---|---|---|---|---|---|---|---|---|
| | | | the Nucleus | | | | | |
| 52 | 0.75 | REACTOME_PATHWAY | Signaling by EGFR | 3 | 176 | 53 | 11462 | 1.10 |
| 52 | 0.75 | GOTERM_BP_DIRECT | positive regulation of cell migration | 5 | 247 | 296 | 18757 | 1.84 |
| 52 | 0.75 | GOTERM_BP_DIRECT | positive regulation of cell population proliferation | 6 | 247 | 542 | 18757 | 2.21 |
| 53 | 0.74 | KEGG_PATHWAY | PD-L1 expression and PD-1 checkpoint pathway in cancer | 7 | 120 | 89 | 8506 | 2.57 |
| 53 | 0.74 | KEGG_PATHWAY | Pancreatic cancer | 6 | 120 | 76 | 8506 | 2.21 |
| 53 | 0.74 | WIKIPATHWAYS | Pancreatic adenocarcinoma pathway | 6 | 138 | 89 | 8952 | 2.21 |
| 53 | 0.74 | KEGG_PATHWAY | Prostate cancer | 6 | 120 | 105 | 8506 | 2.21 |
| 53 | 0.74 | WIKIPATHWAYS | Hallmark of cancer sustaining proliferative signaling | 6 | 138 | 98 | 8952 | 2.21 |
| 53 | 0.74 | KEGG_PATHWAY | Hepatitis C | 7 | 120 | 158 | 8506 | 2.57 |
| 53 | 0.74 | GOTERM_BP_DIRECT | cell population proliferation | 9 | 247 | 266 | 18757 | 3.31 |
| 53 | 0.74 | WIKIPATHWAYS | EGF EGFR signaling | 7 | 138 | 159 | 8952 | 2.57 |
| 53 | 0.74 | KEGG_PATHWAY | FoxO signaling pathway | 6 | 120 | 131 | 8506 | 2.21 |
| 53 | 0.74 | KEGG_PATHWAY | Regulation of actin cytoskeleton | 8 | 120 | 231 | 8506 | 2.94 |
| 53 | 0.74 | WIKIPATHWAYS | NRP1 triggered signaling in pancreatic cancer | 4 | 138 | 53 | 8952 | 1.47 |
| 53 | 0.74 | REACTOME_PATHWAY | Estrogen-dependent nuclear events downstream of ESR-membrane signaling | 3 | 176 | 24 | 11462 | 1.10 |
| 53 | 0.74 | WIKIPATHWAYS | Tyrosine kinase inhibitors in glioblastoma | 3 | 138 | 26 | 8952 | 1.10 |
| 53 | 0.74 | KEGG_PATHWAY | Focal adhesion | 7 | 120 | 202 | 8506 | 2.57 |
| 53 | 0.74 | GOTERM_CC_DIRECT | clathrin-coated endocytic vesicle membrane | 4 | 255 | 71 | 19687 | 1.47 |
| 53 | 0.74 | REACTOME_PATHWAY | Downregulation of ERBB2 signaling | 3 | 176 | 29 | 11462 | 1.10 |
| 53 | 0.74 | WIKIPATHWAYS | MAPK signaling | 8 | 138 | 246 | 8952 | 2.94 |
| 53 | 0.74 | WIKIPATHWAYS | Focal adhesion | 7 | 138 | 199 | 8952 | 2.57 |

| Cluster | Cluster Enrichment Score | Category | Term | Count | List Total | Pop Hits | Pop Total | % |
|---|---|---|---|---|---|---|---|---|
| 53 | 0.74 | KEGG_PATHWAY | Human papillomavirus infection | 9 | 120 | 331 | 8506 | 3.31 |
| 53 | 0.74 | KEGG_PATHWAY | Human cytomegalovirus infection | 7 | 120 | 224 | 8506 | 2.57 |
| 53 | 0.74 | KEGG_PATHWAY | EGFR tyrosine kinase inhibitor resistance | 4 | 120 | 79 | 8506 | 1.47 |
| 53 | 0.74 | WIKIPATHWAYS | Embryonic stem cell pluripotency pathways | 5 | 138 | 117 | 8952 | 1.84 |
| 53 | 0.74 | REACTOME_PATHWAY | Extra-nuclear estrogen signaling | 4 | 176 | 76 | 11462 | 1.47 |
| 53 | 0.74 | GOTERM_BP_DIRECT | phosphatidylinositol 3-kinase/protein kinase B signal transduction | 4 | 247 | 93 | 18757 | 1.47 |
| 53 | 0.74 | KEGG_PATHWAY | MAPK signaling pathway | 8 | 120 | 299 | 8506 | 2.94 |
| 53 | 0.74 | WIKIPATHWAYS | EGFR tyrosine kinase inhibitor resistance | 4 | 138 | 84 | 8952 | 1.47 |
| 53 | 0.74 | UP_KW_DISEASE | Proto-oncogene | 8 | 92 | 233 | 4854 | 2.94 |
| 53 | 0.74 | KEGG_PATHWAY | Phospholipase D signaling pathway | 5 | 120 | 147 | 8506 | 1.84 |
| 53 | 0.74 | KEGG_PATHWAY | Choline metabolism in cancer | 4 | 120 | 98 | 8506 | 1.47 |
| 53 | 0.74 | KEGG_PATHWAY | Rap1 signaling pathway | 6 | 120 | 210 | 8506 | 2.21 |
| 53 | 0.74 | REACTOME_PATHWAY | Signaling by ERBB2 | 3 | 176 | 50 | 11462 | 1.10 |
| 53 | 0.74 | REACTOME_PATHWAY | Intracellular signaling by second messengers | 8 | 176 | 306 | 11462 | 2.94 |
| 53 | 0.74 | KEGG_PATHWAY | Endometrial cancer | 3 | 120 | 58 | 8506 | 1.10 |
| 53 | 0.74 | KEGG_PATHWAY | HIF-1 signaling pathway | 4 | 120 | 109 | 8506 | 1.47 |
| 53 | 0.74 | REACTOME_PATHWAY | Signaling by PTK6 | 3 | 176 | 54 | 11462 | 1.10 |
| 53 | 0.74 | REACTOME_PATHWAY | Signaling by Non-Receptor Tyrosine Kinases | 3 | 176 | 54 | 11462 | 1.10 |
| 53 | 0.74 | KEGG_PATHWAY | Chemical carcinogenesis - reactive oxygen species | 6 | 120 | 223 | 8506 | 2.21 |
| 53 | 0.74 | REACTOME_PATHWAY | PIP3 activates AKT signaling | 7 | 176 | 264 | 11462 | 2.57 |
| 53 | 0.74 | REACTOME_PATHWAY | Cargo recognition for clathrin-mediated endocytosis | 4 | 176 | 107 | 11462 | 1.47 |

| Cluster | Cluster Enrichment Score | Category | Term | Count | List Total | Pop Hits | Pop Total | % |
|---|---|---|---|---|---|---|---|---|
| 53 | 0.74 | KEGG_PATHWAY | Ras signaling pathway | 6 | 120 | 236 | 8506 | 2.21 |
| 53 | 0.74 | KEGG_PATHWAY | PI3K-Akt signaling pathway | 8 | 120 | 359 | 8506 | 2.94 |
| 53 | 0.74 | GOTERM_BP_DIRECT | positive regulation of endothelial cell proliferation | 3 | 247 | 71 | 18757 | 1.10 |
| 53 | 0.74 | GOTERM_BP_DIRECT | epidermal growth factor receptor signaling pathway | 3 | 247 | 71 | 18757 | 1.10 |
| 53 | 0.74 | REACTOME_PATHWAY | ESR-mediated signaling | 6 | 176 | 222 | 11462 | 2.21 |
| 53 | 0.74 | REACTOME_PATHWAY | PI5P, PP2A and IER3 Regulate PI3K/AKT Signaling | 4 | 176 | 113 | 11462 | 1.47 |
| 53 | 0.74 | WIKIPATHWAYS | Endometrial cancer | 3 | 138 | 63 | 8952 | 1.10 |
| 53 | 0.74 | KEGG_PATHWAY | Melanoma | 3 | 120 | 72 | 8506 | 1.10 |
| 53 | 0.74 | KEGG_PATHWAY | Non-small cell lung cancer | 3 | 120 | 72 | 8506 | 1.10 |
| 53 | 0.74 | REACTOME_PATHWAY | Negative regulation of the PI3K/AKT network | 4 | 176 | 120 | 11462 | 1.47 |
| 53 | 0.74 | KEGG_PATHWAY | Glioma | 3 | 120 | 75 | 8506 | 1.10 |
| 53 | 0.74 | REACTOME_PATHWAY | Signaling by Nuclear Receptors | 7 | 176 | 296 | 11462 | 2.57 |
| 53 | 0.74 | WIKIPATHWAYS | Non small cell lung cancer | 3 | 138 | 72 | 8952 | 1.10 |
| 53 | 0.74 | REACTOME_PATHWAY | PTEN Regulation | 4 | 176 | 128 | 11462 | 1.47 |
| 53 | 0.74 | KEGG_PATHWAY | Proteoglycans in cancer | 5 | 120 | 203 | 8506 | 1.84 |
| 53 | 0.74 | WIKIPATHWAYS | Burn wound healing | 3 | 138 | 75 | 8952 | 1.10 |
| 53 | 0.74 | WIKIPATHWAYS | Focal adhesion PI3K Akt mTOR signaling | 7 | 138 | 304 | 8952 | 2.57 |
| 53 | 0.74 | KEGG_PATHWAY | Pathways in cancer | 10 | 120 | 532 | 8506 | 3.68 |
| 53 | 0.74 | KEGG_PATHWAY | ErbB signaling pathway | 3 | 120 | 85 | 8506 | 1.10 |
| 53 | 0.74 | KEGG_PATHWAY | Breast cancer | 4 | 120 | 147 | 8506 | 1.47 |
| 53 | 0.74 | KEGG_PATHWAY | Colorectal cancer | 3 | 120 | 86 | 8506 | 1.10 |
| 53 | 0.74 | KEGG_PATHWAY | Gastric cancer | 4 | 120 | 149 | 8506 | 1.47 |
| 53 | 0.74 | REACTOME_PATHWAY | TP53 Regulates Metabolic Genes | 3 | 176 | 82 | 11462 | 1.10 |
| 53 | 0.74 | KEGG_PATHWAY | Chemical carcinogenesis - receptor activation | 5 | 120 | 216 | 8506 | 1.84 |
| 53 | 0.74 | WIKIPATHWAYS | Cancer pathways | 10 | 138 | 507 | 8952 | 3.68 |

| Cluster | Cluster Enrichment Score | Category | Term | Count | List Total | Pop Hits | Pop Total | % |
|---|---|---|---|---|---|---|---|---|
| 53 | 0.74 | WIKIPATHWAYS | ErbB signaling | 3 | 138 | 90 | 8952 | 1.10 |
| 53 | 0.74 | WIKIPATHWAYS | Regulation of actin cytoskeleton | 4 | 138 | 150 | 8952 | 1.47 |
| 53 | 0.74 | WIKIPATHWAYS | Integrated breast cancer pathway | 4 | 138 | 152 | 8952 | 1.47 |
| 53 | 0.74 | KEGG_PATHWAY | JAK-STAT signaling pathway | 4 | 120 | 167 | 8506 | 1.47 |
| 53 | 0.74 | WIKIPATHWAYS | Breast cancer pathway | 4 | 138 | 154 | 8952 | 1.47 |
| 53 | 0.74 | WIKIPATHWAYS | PI3K Akt signaling | 7 | 138 | 340 | 8952 | 2.57 |
| 53 | 0.74 | KEGG_PATHWAY | Calcium signaling pathway | 5 | 120 | 251 | 8506 | 1.84 |
| 53 | 0.74 | REACTOME_PATHWAY | PI3K/AKT Signaling in Cancer | 3 | 176 | 112 | 11462 | 1.10 |
| 53 | 0.74 | GOTERM_BP_DIRECT | positive regulation of cell migration | 5 | 247 | 296 | 18757 | 1.84 |
| 53 | 0.74 | WIKIPATHWAYS | Effect of omega 3 PUFA on Huntington 39 s disease pathways | 3 | 138 | 120 | 8952 | 1.10 |
| 53 | 0.74 | REACTOME_PATHWAY | Signaling by Receptor Tyrosine Kinases | 9 | 176 | 535 | 11462 | 3.31 |
| 53 | 0.74 | KEGG_PATHWAY | Hepatocellular carcinoma | 3 | 120 | 169 | 8506 | 1.10 |
| 53 | 0.74 | REACTOME_PATHWAY | Signaling by GPCR | 10 | 176 | 715 | 11462 | 3.68 |
| 53 | 0.74 | GOTERM_BP_DIRECT | cell-cell adhesion | 3 | 247 | 225 | 18757 | 1.10 |
| 53 | 0.74 | WIKIPATHWAYS | Pleural mesothelioma | 6 | 138 | 438 | 8952 | 2.21 |
| 53 | 0.74 | GOTERM_BP_DIRECT | positive regulation of cell population proliferation | 6 | 247 | 542 | 18757 | 2.21 |
| 53 | 0.74 | REACTOME_PATHWAY | GPCR downstream signalling | 8 | 176 | 639 | 11462 | 2.94 |
| 53 | 0.74 | UP_KW_MOLECULAR_FUNCTION | Developmental protein | 7 | 165 | 1012 | 11620 | 2.57 |
| 53 | 0.74 | UP_KW_MOLECULAR_FUNCTION | Developmental protein | 7 | 165 | 1012 | 11620 | 2.57 |
| 53 | 0.74 | GOTERM_BP_DIRECT | G protein-coupled receptor signaling pathway | 5 | 247 | 1030 | 18757 | 1.84 |
| 54 | 0.74 | GOTERM_MF_DIRECT | ubiquitin-like ligase-substrate adaptor activity | 6 | 252 | 133 | 18532 | 2.21 |
| 54 | 0.74 | GOTERM_BP_DIRECT | proteasome-mediated ubiquitin-dependent protein catabolic process | 9 | 247 | 354 | 18757 | 3.31 |

| Cluster | Cluster Enrichment Score | Category | Term | Count | List Total | Pop Hits | Pop Total | % |
|---|---|---|---|---|---|---|---|---|
| 54 | 0.74 | REACTOME_PATHWAY | Neddylation | 5 | 176 | 233 | 11462 | 1.84 |
| 54 | 0.74 | REACTOME_PATHWAY | Antigen processing: Ubiquitination & Proteasome degradation | 5 | 176 | 296 | 11462 | 1.84 |
| 55 | 0.71 | GOTERM_CC_DIRECT | lysosomal membrane | 11 | 255 | 407 | 19687 | 4.04 |
| 55 | 0.71 | GOTERM_CC_DIRECT | lysosome | 9 | 255 | 515 | 19687 | 3.31 |
| 55 | 0.71 | UP_KW_CELLULAR_COMPONENT | Lysosome | 7 | 240 | 432 | 17335 | 2.57 |
| 56 | 0.70 | GOTERM_BP_DIRECT | transcription elongation by RNA polymerase II | 3 | 247 | 47 | 18757 | 1.10 |
| 56 | 0.70 | REACTOME_PATHWAY | HIV Transcription Elongation | 3 | 176 | 43 | 11462 | 1.10 |
| 56 | 0.70 | REACTOME_PATHWAY | Formation of HIV-1 elongation complex containing HIV-1 Tat | 3 | 176 | 43 | 11462 | 1.10 |
| 56 | 0.70 | REACTOME_PATHWAY | Tat-mediated elongation of the HIV-1 transcript | 3 | 176 | 43 | 11462 | 1.10 |
| 56 | 0.70 | REACTOME_PATHWAY | Formation of HIV elongation complex in the absence of HIV Tat | 3 | 176 | 45 | 11462 | 1.10 |
| 56 | 0.70 | REACTOME_PATHWAY | Formation of RNA Pol II elongation complex | 3 | 176 | 58 | 11462 | 1.10 |
| 56 | 0.70 | REACTOME_PATHWAY | RNA Polymerase II Transcription Elongation | 3 | 176 | 58 | 11462 | 1.10 |
| 56 | 0.70 | REACTOME_PATHWAY | TP53 Regulates Transcription of DNA Repair Genes | 3 | 176 | 62 | 11462 | 1.10 |
| 56 | 0.70 | REACTOME_PATHWAY | Transcription of the HIV genome | 3 | 176 | 70 | 11462 | 1.10 |
| 56 | 0.70 | REACTOME_PATHWAY | Transcriptional Regulation by TP53 | 8 | 176 | 359 | 11462 | 2.94 |
| 56 | 0.70 | REACTOME_PATHWAY | RNA Polymerase II Pre-transcription Events | 3 | 176 | 81 | 11462 | 1.10 |
| 57 | 0.69 | UP_SEQ_FEATURE | REPEAT:WD 5 | 7 | 261 | 263 | 19119 | 2.57 |
| 57 | 0.69 | UP_SEQ_FEATURE | REPEAT:WD 4 | 7 | 261 | 266 | 19119 | 2.57 |
| 57 | 0.69 | UP_SEQ_FEATURE | REPEAT:WD 3 | 7 | 261 | 269 | 19119 | 2.57 |
| 57 | 0.69 | UP_SEQ_FEATURE | REPEAT:WD 1 | 7 | 261 | 277 | 19119 | 2.57 |
| 57 | 0.69 | UP_SEQ_FEATURE | REPEAT:WD 2 | 7 | 261 | 277 | 19119 | 2.57 |

| Cluster | Cluster Enrichment Score | Category | Term | Count | List Total | Pop Hits | Pop Total | % |
|---|---|---|---|---|---|---|---|---|
| 57 | 0.69 | UP_SEQ_FEATURE | REPEAT:WD 6 | 6 | 261 | 257 | 19119 | 2.21 |
| 57 | 0.69 | UP_SEQ_FEATURE | REPEAT:WD 7 | 5 | 261 | 205 | 19119 | 1.84 |
| 57 | 0.69 | UP_SEQ_FEATURE | REPEAT:WD | 5 | 261 | 205 | 19119 | 1.84 |
| 58 | 0.68 | REACTOME_PATHWAY | Pre-NOTCH Expression and Processing | 6 | 176 | 114 | 11462 | 2.21 |
| 58 | 0.68 | REACTOME_PATHWAY | Pre-NOTCH Transcription and Translation | 4 | 176 | 98 | 11462 | 1.47 |
| 58 | 0.68 | REACTOME_PATHWAY | Oxidative Stress Induced Senescence | 3 | 176 | 125 | 11462 | 1.10 |
| 58 | 0.68 | REACTOME_PATHWAY | Cellular Senescence | 4 | 176 | 197 | 11462 | 1.47 |
| 59 | 0.68 | REACTOME_PATHWAY | Programmed Cell Death | 6 | 176 | 198 | 11462 | 2.21 |
| 59 | 0.68 | REACTOME_PATHWAY | Intrinsic Pathway for Apoptosis | 3 | 176 | 55 | 11462 | 1.10 |
| 59 | 0.68 | REACTOME_PATHWAY | Apoptosis | 5 | 176 | 165 | 11462 | 1.84 |
| 60 | 0.67 | UP_KW_MOLECULAR_FUNCTION | GTPase activation | 7 | 165 | 209 | 11620 | 2.57 |
| 60 | 0.67 | REACTOME_PATHWAY | RHOA GTPase cycle | 5 | 176 | 149 | 11462 | 1.84 |
| 60 | 0.67 | GOTERM_BP_DIRECT | regulation of small GTPase mediated signal transduction | 4 | 247 | 120 | 18757 | 1.47 |
| 60 | 0.67 | REACTOME_PATHWAY | CDC42 GTPase cycle | 3 | 176 | 157 | 11462 | 1.10 |
| 61 | 0.63 | REACTOME_PATHWAY | Constitutive Signaling by NOTCH1 HD+PEST Domain Mutants | 3 | 176 | 58 | 11462 | 1.10 |
| 61 | 0.63 | REACTOME_PATHWAY | Signaling by NOTCH1 PEST Domain Mutants in Cancer | 3 | 176 | 58 | 11462 | 1.10 |
| 61 | 0.63 | REACTOME_PATHWAY | Signaling by NOTCH1 in Cancer | 3 | 176 | 58 | 11462 | 1.10 |
| 61 | 0.63 | REACTOME_PATHWAY | Constitutive Signaling by NOTCH1 PEST Domain Mutants | 3 | 176 | 58 | 11462 | 1.10 |
| 61 | 0.63 | REACTOME_PATHWAY | Signaling by NOTCH1 HD+PEST Domain Mutants in Cancer | 3 | 176 | 58 | 11462 | 1.10 |
| 61 | 0.63 | REACTOME_PATHWAY | Signaling by NOTCH1 | 3 | 176 | 73 | 11462 | 1.10 |
| 62 | 0.61 | GOTERM_BP_DIRECT | symbiont entry into host cell | 4 | 247 | 111 | 18757 | 1.47 |
| 62 | 0.61 | UP_KW_MOLECULAR_FUNCTION | Host cell receptor for virus | 3 | 165 | 73 | 11620 | 1.10 |

| Cluster | Cluster Enrichment Score | Category | Term | Count | List Total | Pop Hits | Pop Total | % |
|---|---|---|---|---|---|---|---|---|
| | | | entry | | | | | |
| 62 | 0.61 | GOTERM_MF_DIRECT | virus receptor activity | 3 | 252 | 81 | 18532 | 1.10 |
| 63 | 0.61 | REACTOME_PATHWAY | Developmental Lineages of the Mammary Gland | 3 | 176 | 39 | 11462 | 1.10 |
| 63 | 0.61 | REACTOME_PATHWAY | Developmental Cell Lineages of the Integumentary System | 3 | 176 | 73 | 11462 | 1.10 |
| 63 | 0.61 | REACTOME_PATHWAY | Developmental Cell Lineages | 4 | 176 | 153 | 11462 | 1.47 |
| 64 | 0.59 | WIKIPATHWAYS | Chemokine signaling | 7 | 138 | 165 | 8952 | 2.57 |
| 64 | 0.59 | GOTERM_BP_DIRECT | regulation of cell migration | 3 | 247 | 123 | 18757 | 1.10 |
| 64 | 0.59 | REACTOME_PATHWAY | GPCR downstream signalling | 8 | 176 | 639 | 11462 | 2.94 |
| 65 | 0.54 | UP_KW_MOLECULAR_FUNCTION | Actin-binding | 8 | 165 | 288 | 11620 | 2.94 |
| 65 | 0.54 | REACTOME_PATHWAY | Sensory processing of sound by outer hair cells of the cochlea | 3 | 176 | 55 | 11462 | 1.10 |
| 65 | 0.54 | REACTOME_PATHWAY | Sensory processing of sound by inner hair cells of the cochlea | 3 | 176 | 68 | 11462 | 1.10 |
| 65 | 0.54 | REACTOME_PATHWAY | Sensory processing of sound | 3 | 176 | 76 | 11462 | 1.10 |
| 65 | 0.54 | REACTOME_PATHWAY | Sensory Perception | 6 | 176 | 647 | 11462 | 2.21 |
| 66 | 0.51 | UP_SEQ_FEATURE | DOMAIN:UPAR/Ly6 | 3 | 261 | 36 | 19119 | 1.10 |
| 66 | 0.51 | GOTERM_CC_DIRECT | side of membrane | 3 | 255 | 145 | 19687 | 1.10 |
| 66 | 0.51 | UP_KW_PTM | GPI-anchor | 3 | 208 | 145 | 14032 | 1.10 |
| 67 | 0.48 | GOTERM_MF_DIRECT | dioxygenase activity | 4 | 252 | 86 | 18532 | 1.47 |
| 67 | 0.48 | UP_KW_MOLECULAR_FUNCTION | Dioxygenase | 4 | 165 | 86 | 11620 | 1.47 |
| 67 | 0.48 | GOTERM_MF_DIRECT | iron ion binding | 4 | 252 | 141 | 18532 | 1.47 |
| 67 | 0.48 | GOTERM_MF_DIRECT | oxidoreductase activity | 5 | 252 | 617 | 18532 | 1.84 |
| 67 | 0.48 | UP_KW_MOLECULAR_FUNCTION | Oxidoreductase | 5 | 165 | 603 | 11620 | 1.84 |
| 68 | 0.47 | UP_SEQ_FEATURE | REPEAT:ANK | 5 | 261 | 185 | 19119 | 1.84 |
| 68 | 0.47 | UP_SEQ_FEATURE | REPEAT:ANK 1 | 6 | 261 | 251 | 19119 | 2.21 |
| 68 | 0.47 | UP_SEQ_FEATURE | REPEAT:ANK 2 | 6 | 261 | 252 | 19119 | 2.21 |

| Cluster | Cluster Enrichment Score | Category | Term | Count | List Total | Pop Hits | Pop Total | % |
|---|---|---|---|---|---|---|---|---|
| 68 | 0.47 | UP_SEQ_FEATURE | REPEAT:ANK 3 | 3 | 261 | 215 | 19119 | 1.10 |
| 69 | 0.46 | REACTOME_PATHWAY | RNA Polymerase I Transcription Initiation | 3 | 176 | 48 | 11462 | 1.10 |
| 69 | 0.46 | REACTOME_PATHWAY | RNA Polymerase I Promoter Clearance | 4 | 176 | 112 | 11462 | 1.47 |
| 69 | 0.46 | REACTOME_PATHWAY | RNA Polymerase I Transcription | 4 | 176 | 113 | 11462 | 1.47 |
| 69 | 0.46 | REACTOME_PATHWAY | B-WICH complex positively regulates rRNA expression | 3 | 176 | 91 | 11462 | 1.10 |
| 69 | 0.46 | REACTOME_PATHWAY | RNA Polymerase I Promoter Escape | 3 | 176 | 91 | 11462 | 1.10 |
| 69 | 0.46 | REACTOME_PATHWAY | Positive epigenetic regulation of rRNA expression | 3 | 176 | 107 | 11462 | 1.10 |
| 69 | 0.46 | REACTOME_PATHWAY | NoRC negatively regulates rRNA expression | 3 | 176 | 107 | 11462 | 1.10 |
| 69 | 0.46 | REACTOME_PATHWAY | Negative epigenetic regulation of rRNA expression | 3 | 176 | 110 | 11462 | 1.10 |
| 70 | 0.45 | GOTERM_BP_DIRECT | chromatin organization | 9 | 247 | 453 | 18757 | 3.31 |
| 70 | 0.45 | GOTERM_BP_DIRECT | chromatin remodeling | 7 | 247 | 324 | 18757 | 2.57 |
| 70 | 0.45 | GOTERM_MF_DIRECT | histone deacetylase binding | 4 | 252 | 134 | 18532 | 1.47 |
| 70 | 0.45 | REACTOME_PATHWAY | Chromatin modifying enzymes | 6 | 176 | 274 | 11462 | 2.21 |
| 70 | 0.45 | REACTOME_PATHWAY | Chromatin organization | 6 | 176 | 289 | 11462 | 2.21 |
| 70 | 0.45 | UP_KW_MOLECULAR_FUNCTION | Chromatin regulator | 6 | 165 | 314 | 11620 | 2.21 |
| 70 | 0.45 | GOTERM_MF_DIRECT | chromatin binding | 8 | 252 | 499 | 18532 | 2.94 |
| 71 | 0.41 | KEGG_PATHWAY | Human immunodeficiency virus 1 infection | 7 | 120 | 212 | 8506 | 2.57 |
| 71 | 0.41 | GOTERM_BP_DIRECT | positive regulation of canonical NF-kappaB signal transduction | 7 | 247 | 233 | 18757 | 2.57 |
| 71 | 0.41 | REACTOME_PATHWAY | TCR signaling | 5 | 176 | 115 | 11462 | 1.84 |
| 71 | 0.41 | REACTOME_PATHWAY | Downstream TCR signaling | 4 | 176 | 93 | 11462 | 1.47 |
| 71 | 0.41 | REACTOME_PATHWAY | Activation of NF-kappaB in B cells | 3 | 176 | 54 | 11462 | 1.10 |

| Cluster | Cluster Enrichment Score | Category | Term | Count | List Total | Pop Hits | Pop Total | % |
|---|---|---|---|---|---|---|---|---|
| 71 | 0.41 | REACTOME_PATHWAY | Downstream signaling events of B Cell Receptor (BCR) | 3 | 176 | 71 | 11462 | 1.10 |
| 71 | 0.41 | REACTOME_PATHWAY | Interleukin-1 family signaling | 4 | 176 | 142 | 11462 | 1.47 |
| 71 | 0.41 | REACTOME_PATHWAY | CLEC7A (Dectin-1) signaling | 3 | 176 | 87 | 11462 | 1.10 |
| 71 | 0.41 | REACTOME_PATHWAY | MyD88 cascade initiated on plasma membrane | 3 | 176 | 97 | 11462 | 1.10 |
| 71 | 0.41 | REACTOME_PATHWAY | Toll Like Receptor 10 (TLR10) Cascade | 3 | 176 | 97 | 11462 | 1.10 |
| 71 | 0.41 | REACTOME_PATHWAY | Toll Like Receptor 5 (TLR5) Cascade | 3 | 176 | 97 | 11462 | 1.10 |
| 71 | 0.41 | REACTOME_PATHWAY | TRAF6 mediated induction of NFkB and MAP kinases upon TLR7/8 or 9 activation | 3 | 176 | 102 | 11462 | 1.10 |
| 71 | 0.41 | REACTOME_PATHWAY | Interleukin-1 signaling | 3 | 176 | 102 | 11462 | 1.10 |
| 71 | 0.41 | REACTOME_PATHWAY | MyD88 dependent cascade initiated on endosome | 3 | 176 | 103 | 11462 | 1.10 |
| 71 | 0.41 | REACTOME_PATHWAY | Toll Like Receptor 3 (TLR3) Cascade | 3 | 176 | 106 | 11462 | 1.10 |
| 71 | 0.41 | REACTOME_PATHWAY | Toll Like Receptor 7/8 (TLR7/8) Cascade | 3 | 176 | 107 | 11462 | 1.10 |
| 71 | 0.41 | REACTOME_PATHWAY | MyD88-independent TLR4 cascade | 3 | 176 | 110 | 11462 | 1.10 |
| 71 | 0.41 | REACTOME_PATHWAY | Toll Like Receptor 9 (TLR9) Cascade | 3 | 176 | 110 | 11462 | 1.10 |
| 71 | 0.41 | REACTOME_PATHWAY | TRIF (TICAM1)-mediated TLR4 signaling | 3 | 176 | 110 | 11462 | 1.10 |
| 71 | 0.41 | REACTOME_PATHWAY | MyD88:MAL(TIRAP) cascade initiated on plasma membrane | 3 | 176 | 113 | 11462 | 1.10 |
| 71 | 0.41 | REACTOME_PATHWAY | Toll Like Receptor TLR6:TLR2 Cascade | 3 | 176 | 113 | 11462 | 1.10 |
| 71 | 0.41 | REACTOME_PATHWAY | Fc epsilon receptor (FCERI) signaling | 4 | 176 | 182 | 11462 | 1.47 |
| 71 | 0.41 | REACTOME_PATHWAY | Toll Like Receptor TLR1:TLR2 Cascade | 3 | 176 | 116 | 11462 | 1.10 |
| 71 | 0.41 | REACTOME_PATHWAY | Toll Like Receptor 2 (TLR2) Cascade | 3 | 176 | 116 | 11462 | 1.10 |

| Cluster | Cluster Enrichment Score | Category | Term | Count | List Total | Pop Hits | Pop Total | % |
|---|---|---|---|---|---|---|---|---|
| 71 | 0.41 | REACTOME_PATHWAY | C-type lectin receptors (CLRs) | 3 | 176 | 129 | 11462 | 1.10 |
| 71 | 0.41 | REACTOME_PATHWAY | FCERI mediated NF-kB activation | 3 | 176 | 130 | 11462 | 1.10 |
| 71 | 0.41 | REACTOME_PATHWAY | Toll Like Receptor 4 (TLR4) Cascade | 3 | 176 | 146 | 11462 | 1.10 |
| 71 | 0.41 | REACTOME_PATHWAY | Signaling by the B Cell Receptor (BCR) | 3 | 176 | 162 | 11462 | 1.10 |
| 71 | 0.41 | REACTOME_PATHWAY | Toll-like Receptor Cascades | 3 | 176 | 171 | 11462 | 1.10 |
| 72 | 0.40 | REACTOME_PATHWAY | Synthesis of PIPs at the plasma membrane | 3 | 176 | 54 | 11462 | 1.10 |
| 72 | 0.40 | REACTOME_PATHWAY | PI Metabolism | 3 | 176 | 84 | 11462 | 1.10 |
| 72 | 0.40 | REACTOME_PATHWAY | Phospholipid metabolism | 3 | 176 | 212 | 11462 | 1.10 |
| 73 | 0.39 | GOTERM_MF_DIRECT | single-stranded RNA binding | 3 | 252 | 49 | 18532 | 1.10 |
| 73 | 0.39 | UP_KW_MOLECULAR_FUNCTION | Ribonucleoprotein | 5 | 165 | 307 | 11620 | 1.84 |
| 73 | 0.39 | GOTERM_CC_DIRECT | ribonucleoprotein complex | 5 | 255 | 403 | 19687 | 1.84 |
| 74 | 0.38 | GOTERM_BP_DIRECT | DNA damage response | 11 | 247 | 578 | 18757 | 4.04 |
| 74 | 0.38 | UP_KW_BIOLOGICAL_PROCESS | DNA damage | 8 | 163 | 434 | 11213 | 2.94 |
| 74 | 0.38 | GOTERM_BP_DIRECT | DNA repair | 7 | 247 | 426 | 18757 | 2.57 |
| 74 | 0.38 | UP_KW_BIOLOGICAL_PROCESS | DNA repair | 6 | 163 | 360 | 11213 | 2.21 |
| 75 | 0.37 | KEGG_PATHWAY | Rap1 signaling pathway | 6 | 120 | 210 | 8506 | 2.21 |
| 75 | 0.37 | KEGG_PATHWAY | IgSF CAM signaling | 7 | 120 | 298 | 8506 | 2.57 |
| 75 | 0.37 | KEGG_PATHWAY | Platelet activation | 4 | 120 | 125 | 8506 | 1.47 |
| 75 | 0.37 | GOTERM_CC_DIRECT | cell-cell junction | 3 | 255 | 208 | 19687 | 1.10 |
| 75 | 0.37 | REACTOME_PATHWAY | Platelet activation, signaling and aggregation | 4 | 176 | 266 | 11462 | 1.47 |
| 75 | 0.37 | REACTOME_PATHWAY | Hemostasis | 7 | 176 | 694 | 11462 | 2.57 |
| 76 | 0.36 | UP_SEQ_FEATURE | REPEAT:17 | 3 | 261 | 58 | 19119 | 1.10 |
| 76 | 0.36 | UP_SEQ_FEATURE | REPEAT:16 | 3 | 261 | 61 | 19119 | 1.10 |
| 76 | 0.36 | UP_SEQ_FEATURE | REPEAT:15 | 3 | 261 | 71 | 19119 | 1.10 |
| 76 | 0.36 | UP_SEQ_FEATURE | REPEAT:14 | 3 | 261 | 79 | 19119 | 1.10 |
| 76 | 0.36 | UP_SEQ_FEATURE | REPEAT:13 | 3 | 261 | 85 | 19119 | 1.10 |

| Cluster | Cluster Enrichment Score | Category | Term | Count | List Total | Pop Hits | Pop Total | % |
|---|---|---|---|---|---|---|---|---|
| 76 | 0.36 | UP_SEQ_FEATURE | REPEAT:12 | 3 | 261 | 91 | 19119 | 1.10 |
| 76 | 0.36 | UP_SEQ_FEATURE | REPEAT:11 | 3 | 261 | 93 | 19119 | 1.10 |
| 76 | 0.36 | UP_SEQ_FEATURE | REPEAT:10 | 3 | 261 | 99 | 19119 | 1.10 |
| 76 | 0.36 | UP_SEQ_FEATURE | REPEAT:9 | 3 | 261 | 109 | 19119 | 1.10 |
| 76 | 0.36 | UP_SEQ_FEATURE | REPEAT:7 | 3 | 261 | 135 | 19119 | 1.10 |
| 76 | 0.36 | UP_SEQ_FEATURE | REPEAT:6 | 3 | 261 | 148 | 19119 | 1.10 |
| 76 | 0.36 | UP_SEQ_FEATURE | REPEAT:5 | 3 | 261 | 165 | 19119 | 1.10 |
| 76 | 0.36 | UP_SEQ_FEATURE | REPEAT:4 | 3 | 261 | 189 | 19119 | 1.10 |
| 76 | 0.36 | UP_SEQ_FEATURE | REPEAT:3 | 3 | 261 | 222 | 19119 | 1.10 |
| 76 | 0.36 | UP_SEQ_FEATURE | REPEAT:1 | 3 | 261 | 247 | 19119 | 1.10 |
| 76 | 0.36 | UP_SEQ_FEATURE | REPEAT:2 | 3 | 261 | 250 | 19119 | 1.10 |
| 77 | 0.34 | GOTERM_BP_DIRECT | proteolysis | 11 | 247 | 625 | 18757 | 4.04 |
| 77 | 0.34 | UP_KW_MOLECULAR_FUNCTION | Thiol protease | 4 | 165 | 155 | 11620 | 1.47 |
| 77 | 0.34 | UP_SEQ_FEATURE | ACT_SITE:Nucleophile | 7 | 261 | 382 | 19119 | 2.57 |
| 77 | 0.34 | GOTERM_BP_DIRECT | protein deubiquitination | 3 | 247 | 113 | 18757 | 1.10 |
| 77 | 0.34 | GOTERM_MF_DIRECT | peptidase activity | 9 | 252 | 534 | 18532 | 3.31 |
| 77 | 0.34 | GOTERM_MF_DIRECT | cysteine-type deubiquitinase activity | 3 | 252 | 117 | 18532 | 1.10 |
| 77 | 0.34 | GOTERM_MF_DIRECT | cysteine-type peptidase activity | 3 | 252 | 154 | 18532 | 1.10 |
| 77 | 0.34 | UP_KW_MOLECULAR_FUNCTION | Protease | 8 | 165 | 540 | 11620 | 2.94 |
| 78 | 0.34 | KEGG_PATHWAY | Chemical carcinogenesis - reactive oxygen species | 6 | 120 | 223 | 8506 | 2.21 |
| 78 | 0.34 | KEGG_PATHWAY | Alzheimer disease | 8 | 120 | 383 | 8506 | 2.94 |
| 78 | 0.34 | KEGG_PATHWAY | Non-alcoholic fatty liver disease | 3 | 120 | 153 | 8506 | 1.10 |
| 78 | 0.34 | WIKIPATHWAYS | Nonalcoholic fatty liver disease | 3 | 138 | 152 | 8952 | 1.10 |
| 78 | 0.34 | KEGG_PATHWAY | Diabetic cardiomyopathy | 3 | 120 | 201 | 8506 | 1.10 |
| 79 | 0.33 | GOTERM_CC_DIRECT | spliceosomal complex | 5 | 255 | 186 | 19687 | 1.84 |
| 79 | 0.33 | GOTERM_BP_DIRECT | RNA splicing | 7 | 247 | 322 | 18757 | 2.57 |
| 79 | 0.33 | UP_KW_BIOLOGICAL_PROCESS | mRNA splicing | 7 | 163 | 306 | 11213 | 2.57 |

| Cluster | Cluster Enrichment Score | Category | Term | Count | List Total | Pop Hits | Pop Total | % |
|---|---|---|---|---|---|---|---|---|
| 79 | 0.33 | GOTERM_BP_DIRECT | mRNA processing | 7 | 247 | 418 | 18757 | 2.57 |
| 79 | 0.33 | UP_KW_BIOLOGICAL_PROCESS | mRNA processing | 7 | 163 | 394 | 11213 | 2.57 |
| 79 | 0.33 | UP_KW_CELLULAR_COMPONENT | Spliceosome | 3 | 240 | 151 | 17335 | 1.10 |
| 79 | 0.33 | GOTERM_BP_DIRECT | mRNA splicing, via spliceosome | 3 | 247 | 193 | 18757 | 1.10 |
| 79 | 0.33 | REACTOME_PATHWAY | mRNA Splicing - Major Pathway | 3 | 176 | 204 | 11462 | 1.10 |
| 79 | 0.33 | REACTOME_PATHWAY | mRNA Splicing | 3 | 176 | 212 | 11462 | 1.10 |
| 80 | 0.32 | KEGG_PATHWAY | Fluid shear stress and atherosclerosis | 4 | 120 | 141 | 8506 | 1.47 |
| 80 | 0.32 | GOTERM_MF_DIRECT | protein homodimerization activity | 11 | 252 | 731 | 18532 | 4.04 |
| 80 | 0.32 | GOTERM_MF_DIRECT | protein kinase binding | 7 | 252 | 486 | 18532 | 2.57 |
| 81 | 0.29 | KEGG_PATHWAY | Alzheimer disease | 8 | 120 | 383 | 8506 | 2.94 |
| 81 | 0.29 | WIKIPATHWAYS | Alzheimer 39 s disease | 6 | 138 | 262 | 8952 | 2.21 |
| 81 | 0.29 | WIKIPATHWAYS | Alzheimer 39 s disease and miRNA effects | 6 | 138 | 267 | 8952 | 2.21 |
| 81 | 0.29 | KEGG_PATHWAY | Pathways of neurodegeneration - multiple diseases | 5 | 120 | 475 | 8506 | 1.84 |
| 81 | 0.29 | GOTERM_MF_DIRECT | signaling receptor binding | 4 | 252 | 424 | 18532 | 1.47 |
| 82 | 0.26 | GOTERM_MF_DIRECT | kinase activity | 13 | 252 | 726 | 18532 | 4.78 |
| 82 | 0.26 | UP_KW_MOLECULAR_FUNCTION | Kinase | 13 | 165 | 741 | 11620 | 4.78 |
| 82 | 0.26 | GOTERM_MF_DIRECT | protein serine kinase activity | 6 | 252 | 364 | 18532 | 2.21 |
| 82 | 0.26 | UP_KW_MOLECULAR_FUNCTION | Serine/threonine-protein kinase | 6 | 165 | 391 | 11620 | 2.21 |
| 82 | 0.26 | UP_SEQ_FEATURE | ACT_SITE:Proton acceptor | 12 | 261 | 883 | 19119 | 4.41 |
| 82 | 0.26 | GOTERM_MF_DIRECT | protein serine/threonine kinase activity | 6 | 252 | 413 | 18532 | 2.21 |
| 82 | 0.26 | UP_SEQ_FEATURE | DOMAIN:Protein kinase | 7 | 261 | 492 | 19119 | 2.57 |
| 82 | 0.26 | GOTERM_BP_DIRECT | protein phosphorylation | 4 | 247 | 261 | 18757 | 1.47 |
| 82 | 0.26 | GOTERM_MF_DIRECT | protein kinase activity | 7 | 252 | 523 | 18532 | 2.57 |
| 83 | 0.23 | GOTERM_MF_DIRECT | RNA polymerase II transcription regulatory region sequence-specific | 8 | 252 | 388 | 18532 | 2.94 |

| Cluster | Cluster Enrichment Score | Category | Term | Count | List Total | Pop Hits | Pop Total | % |
|---|---|---|---|---|---|---|---|---|
| | | | DNA binding | | | | | |
| 83 | 0.23 | UP_SEQ_FEATURE | ZN_FING:C2H2-type 2 | 11 | 261 | 627 | 19119 | 4.04 |
| 83 | 0.23 | UP_SEQ_FEATURE | DOMAIN:KRAB | 7 | 261 | 381 | 19119 | 2.57 |
| 83 | 0.23 | UP_SEQ_FEATURE | ZN_FING:C2H2-type 7 | 8 | 261 | 481 | 19119 | 2.94 |
| 83 | 0.23 | UP_SEQ_FEATURE | ZN_FING:C2H2-type 5 | 9 | 261 | 565 | 19119 | 3.31 |
| 83 | 0.23 | UP_SEQ_FEATURE | ZN_FING:C2H2-type 12 | 5 | 261 | 276 | 19119 | 1.84 |
| 83 | 0.23 | UP_SEQ_FEATURE | ZN_FING:C2H2-type 6 | 8 | 261 | 513 | 19119 | 2.94 |
| 83 | 0.23 | UP_SEQ_FEATURE | ZN_FING:C2H2-type 13 | 4 | 261 | 219 | 19119 | 1.47 |
| 83 | 0.23 | UP_SEQ_FEATURE | ZN_FING:C2H2-type 15 | 3 | 261 | 148 | 19119 | 1.10 |
| 83 | 0.23 | UP_SEQ_FEATURE | ZN_FING:C2H2-type 11 | 5 | 261 | 311 | 19119 | 1.84 |
| 83 | 0.23 | GOTERM_MF_DIRECT | DNA binding | 31 | 252 | 2308 | 18532 | 11.40 |
| 83 | 0.23 | UP_SEQ_FEATURE | ZN_FING:C2H2-type 1 | 8 | 261 | 557 | 19119 | 2.94 |
| 83 | 0.23 | UP_SEQ_FEATURE | ZN_FING:C2H2-type 9 | 6 | 261 | 406 | 19119 | 2.21 |
| 83 | 0.23 | UP_SEQ_FEATURE | ZN_FING:C2H2-type 3 | 9 | 261 | 648 | 19119 | 3.31 |
| 83 | 0.23 | UP_SEQ_FEATURE | ZN_FING:C2H2-type 14 | 3 | 261 | 171 | 19119 | 1.10 |
| 83 | 0.23 | UP_SEQ_FEATURE | DOMAIN:C2H2-type | 7 | 261 | 508 | 19119 | 2.57 |
| 83 | 0.23 | UP_SEQ_FEATURE | ZN_FING:C2H2-type 4 | 8 | 261 | 600 | 19119 | 2.94 |
| 83 | 0.23 | UP_SEQ_FEATURE | ZN_FING:C2H2-type 8 | 6 | 261 | 448 | 19119 | 2.21 |
| 83 | 0.23 | UP_SEQ_FEATURE | ZN_FING:C2H2-type 10 | 5 | 261 | 365 | 19119 | 1.84 |
| 83 | 0.23 | UP_KW_MOLECULAR_FUNCTION | DNA-binding | 27 | 165 | 2074 | 11620 | 9.93 |
| 83 | 0.23 | GOTERM_BP_DIRECT | regulation of DNA-templated transcription | 17 | 247 | 1471 | 18757 | 6.25 |
| 83 | 0.23 | GOTERM_MF_DIRECT | DNA-binding transcription factor activity, RNA polymerase II-specific | 13 | 252 | 1227 | 18532 | 4.78 |
| 84 | 0.18 | UP_KW_BIOLOGICAL_PROCESS | Protein biosynthesis | 4 | 163 | 151 | 11213 | 1.47 |
| 84 | 0.18 | REACTOME_PATHWAY | Translation | 5 | 176 | 371 | 11462 | 1.84 |
| 84 | 0.18 | GOTERM_BP_DIRECT | translation | 3 | 247 | 310 | 18757 | 1.10 |
| 85 | 0.11 | GOTERM_CC_DIRECT | monoatomic ion channel complex | 3 | 255 | 149 | 19687 | 1.10 |
| 85 | 0.11 | UP_KW_MOLECULAR_FUNCTION | Ion channel | 5 | 165 | 354 | 11620 | 1.84 |

| Cluster | Cluster Enrichment Score | Category | Term | Count | List Total | Pop Hits | Pop Total | % |
|---|---|---|---|---|---|---|---|---|
| 85 | 0.11 | GOTERM_BP_DIRECT | monoatomic ion transmembrane transport | 5 | 247 | 413 | 18757 | 1.84 |
| 85 | 0.11 | GOTERM_BP_DIRECT | monoatomic ion transport | 7 | 247 | 667 | 18757 | 2.57 |
| 85 | 0.11 | UP_KW_BIOLOGICAL_PROCESS | Ion transport | 6 | 163 | 636 | 11213 | 2.21 |
| 86 | 0.10 | GOTERM_CC_DIRECT | sperm flagellum | 3 | 255 | 158 | 19687 | 1.10 |
| 86 | 0.10 | UP_KW_CELLULAR_COMPONENT | Flagellum | 3 | 240 | 216 | 17335 | 1.10 |
| 86 | 0.10 | GOTERM_CC_DIRECT | motile cilium | 3 | 255 | 259 | 19687 | 1.10 |
| 86 | 0.10 | UP_KW_CELLULAR_COMPONENT | Cilium | 3 | 240 | 377 | 17335 | 1.10 |
| 87 | 0.07 | GOTERM_BP_DIRECT | innate immune response | 9 | 247 | 610 | 18757 | 3.31 |
| 87 | 0.07 | UP_KW_BIOLOGICAL_PROCESS | Innate immunity | 4 | 163 | 421 | 11213 | 1.47 |
| 87 | 0.07 | GOTERM_BP_DIRECT | immune system process | 7 | 247 | 929 | 18757 | 2.57 |
| 87 | 0.07 | UP_KW_BIOLOGICAL_PROCESS | Immunity | 6 | 163 | 910 | 11213 | 2.21 |
| 88 | 0.05 | UP_SEQ_FEATURE | REPEAT:LRR 4 | 3 | 261 | 259 | 19119 | 1.10 |
| 88 | 0.05 | UP_SEQ_FEATURE | REPEAT:LRR 3 | 3 | 261 | 272 | 19119 | 1.10 |
| 88 | 0.05 | UP_SEQ_FEATURE | REPEAT:LRR 2 | 3 | 261 | 281 | 19119 | 1.10 |
| 88 | 0.05 | UP_SEQ_FEATURE | REPEAT:LRR 1 | 3 | 261 | 284 | 19119 | 1.10 |
| 89 | 0.04 | REACTOME_PATHWAY | Neurotransmitter receptors and postsynaptic signal transmission | 3 | 176 | 209 | 11462 | 1.10 |
| 89 | 0.04 | REACTOME_PATHWAY | Transmission across Chemical Synapses | 3 | 176 | 274 | 11462 | 1.10 |
| 89 | 0.04 | REACTOME_PATHWAY | Neuronal System | 4 | 176 | 416 | 11462 | 1.47 |
| 90 | 0.04 | GOTERM_CC_DIRECT | mitochondrial matrix | 5 | 255 | 450 | 19687 | 1.84 |
| 90 | 0.04 | REACTOME_PATHWAY | Metabolism of amino acids and derivatives | 4 | 176 | 357 | 11462 | 1.47 |
| 90 | 0.04 | UP_KW_CELLULAR_COMPONENT | Mitochondrion | 14 | 240 | 1362 | 17335 | 5.15 |
| 90 | 0.04 | UP_SEQ_FEATURE | TRANSIT:Mitochondrion | 5 | 261 | 560 | 19119 | 1.84 |
| 91 | 0.04 | REACTOME_PATHWAY | Peptide ligand-binding receptors | 3 | 176 | 202 | 11462 | 1.10 |
| 91 | 0.04 | REACTOME_PATHWAY | Class A/1 (Rhodopsin-like receptors) | 3 | 176 | 340 | 11462 | 1.10 |
| 91 | 0.04 | REACTOME_PATHWAY | GPCR ligand binding | 4 | 176 | 474 | 11462 | 1.47 |
| 92 | 0.00 | UP_SEQ_FEATURE | TOPO_DOM:Cytoplasmic | 39 | 261 | 3852 | 19119 | 14.34 |

| Cluster | Cluster Enrichment Score | Category | Term | Count | List Total | Pop Hits | Pop Total | % |
|---|---|---|---|---|---|---|---|---|
| 92 | 0.00 | UP_SEQ_FEATURE | TRANSMEM:Helical | 49 | 261 | 5102 | 19119 | 18.01 |
| 92 | 0.00 | UP_SEQ_FEATURE | TOPO_DOM:Extracellular | 20 | 261 | 2945 | 19119 | 7.35 |
| 92 | 0.00 | UP_SEQ_FEATURE | CARBOHYD:N-linked (GlcNAc...) asparagine | 34 | 261 | 4329 | 19119 | 12.50 |
| 93 | 0.00 | GOTERM_CC_DIRECT | extracellular region | 20 | 255 | 2564 | 19687 | 7.35 |
| 93 | 0.00 | GOTERM_CC_DIRECT | extracellular space | 12 | 255 | 1845 | 19687 | 4.41 |
| 93 | 0.00 | UP_KW_CELLULAR_COMPONENT | Secreted | 13 | 240 | 2104 | 17335 | 4.78 |
| 94 | 0.00 | UP_SEQ_FEATURE | TRANSMEM:Helical; Name=3 | 4 | 261 | 1071 | 19119 | 1.47 |
| 94 | 0.00 | UP_SEQ_FEATURE | TRANSMEM:Helical; Name=4 | 4 | 261 | 1073 | 19119 | 1.47 |
| 94 | 0.00 | UP_SEQ_FEATURE | TRANSMEM:Helical; Name=1 | 4 | 261 | 1088 | 19119 | 1.47 |
| 94 | 0.00 | UP_SEQ_FEATURE | TRANSMEM:Helical; Name=2 | 4 | 261 | 1095 | 19119 | 1.47 |
