## Supplementary File - Peer Review for "Iterative Gene Enrichment Analysis: interpretable networks for human and AI-assisted biological insights"

### Peer review history

Maxim V. Kuleshov, Yair Benita — AION Labs, Rehovot, Israel

#### Purpose of this document

This preprint was submitted to peer-reviewed journals before being posted to bioRxiv. It was ultimately rejected, and the authors do not intend to submit it elsewhere. Rather than leave the work unpublished, we are posting it here together with the complete peer review correspondence so that readers can evaluate the method, the criticisms raised against it, and our responses, and reach their own conclusions.

#### Submission history

**Bioinformatics (Oxford University Press).** Manuscript BIOINF-2025-2109, submitted August 2025. Declined without external review on 8 September 2025 on editorial capacity grounds; the decision letter cites submission volume and a target acceptance rate of 20-25%, and states no assessment of the work's merit.

**BMC Bioinformatics, first submission.** Submission ID 6159728f-72a8-41ce-9350-8146f6a82c2d. Received 20 September 2025; an amended manuscript was requested and received on 22 September; the submission passed technical check and was assigned to an editor on 17 October; reviewers were invited on 24 October; two reviewer reports were received on 11 November; the manuscript was rejected on 19 November 2025. The authors filed a formal appeal, which was granted, and were invited to submit a revised version.

**BMC Bioinformatics, second submission.** Submission ID 6e5f1a44-cee9-4011-8305-0f068bcd1604. Received 2 April 2026; reviewers were invited on 20 April; four reviewer reports were received on 13 May; a revision was requested on 28 May 2026. The revised manuscript was received on 13 June 2026. Reviewers were invited on 5 August, reports were returned on 9 August, and the manuscript was rejected on 25 August 2026. Handling Editor: Alireza Khanteymoori.

From the first submission to BMC Bioinformatics to the final decision, the manuscript was under consideration for eleven months. Dates are taken from the journal's submission status history.

#### Note on the final decision

The final decision did not provide a clear reason for the rejection.

Reviewer 2 reported that the file delivered to them as the "Revised Manuscript" was an email invitation to re-review containing no manuscript content, and recommended rejection on that basis alone, stating that "until the actual manuscript is provided, the review cannot proceed." Our response: the revised manuscript had in fact been prepared and submitted; we validated it within the submission system of the journal. We wrote to the editorial office on 26 August 2026 asking which files had been made available to each reviewer, and followed up on 30 August. We received no reply.

Reviewer 3 wrote that their questions had been addressed and they had no further comment.

Reviewer 4 maintained that a shorter list of terms does not by itself demonstrate better biological inference, but offered no suggestion for how such a demonstration should be made. We accept the premise, and the manuscript does not argue otherwise. Our claim is representational: iGEA produces a more compact and structured representation of enrichment results than standard ORA, and that representation can be inspected directly by a human expert or supplied as a structured input for LLM-assisted exploration. We do not claim that it yields more accurate biological inference, nor that LLM interpretation of it is superior to alternative approaches. The stronger claim did appear in the version these reviewers read – the abstract of the previous revision stated that the network "improves the consistency and usefulness of LLM-based interpretation compared with unstructured enrichment outputs" – and we removed it in response to Reviewer 1, replacing it with an explicit statement that our examples show how iGEA networks can be used as structured inputs, "not that LLM interpretation is superior to alternative approaches." What remains is the representational claim, supported by direct comparison with ORA on the same gene list, comparison with DAVID functional clustering, and empirical benchmarking of network structure against 1,000 random gene lists of matched size.

The choice of validation dataset was deliberate. We selected the 2008 HIV dependency-factor screen precisely because the intervening years established which signals were real: the individual screens of that era produced poorly overlapping gene lists, while subsequent follow-up studies and cross-study meta-analyses converged at the pathway level. This provides an external reference against which recovery can be judged, and the modules iGEA identified corresponded closely to themes identified in those meta-analyses. At the reviewers' request we also benchmarked against additional tools; the corresponding DAVID report spanned 38 pages and remained difficult to synthesise into a coherent hypothesis, whereas applying a current LLM to the iGEA network produced a compact summary of the dominant biological signal.

We consider the evidence presented sufficient to support the claims made in the manuscript, which we have deliberately kept modest, and we leave the judgement to the readers. We would welcome concrete proposals for testing iGEA's ability to recover biological signal, and would be glad to pressure-test the method against them.

#### Contents


1. Editorial decision — *Bioinformatics*, 8 September 2025
2. Point-by-point response to reviewers — *BMC Bioinformatics*, first submission (reports of 11 November 2025)
3. Point-by-point response to reviewers — *BMC Bioinformatics*, second submission (reports of 13 May 2026)
4. Editorial decision and reviewer comments — *BMC Bioinformatics*, 25 August 2026


Reviewer comments are reproduced verbatim within the response documents, each followed by the authors' reply.

### Editorial decision — Bioinformatics, 8 September 2025

From:

To:

CC:

Subject:

Bioinformatics BIOINF-2025-2109 Decision

Body:

08-Sep-2025 BIOINF-2025-2109 Iterative Gene Enrichment Analysis: interpretable networks for human and AI-assisted biological insights

Dear Dr. Benita,

Thank you for giving us the opportunity to consider your manuscript for publication in Bioinformatics.

An associate editor has assessed your manuscript and has decided that the journal is unable to consider the manuscript for further review.

I am sorry to disappoint you, but the journal is currently receiving many more submissions than can be published - over 2000 submissions were received in 2023. Hence only those of the very highest quality and significance are accepted (the editors currently aim for an acceptance rate of ~20-25%).

The journal must therefore formally decline the manuscript and we are returning it to you without further review so that you can resubmit it elsewhere without delay. Please note that we will not consider any further versions of this manuscript.

Best regards, S Ceci Bioinformatics

Date Sent:

08-Sep-2025

Ref: Submission ID 6159728f-72a8-41ce-9350-8146f6a82c2d

We thank the reviewers and the editor for their careful evaluation of our manuscript and for the constructive and insightful comments. We have substantially revised the manuscript to address all major concerns raised during review. In particular, we have strengthened the statistical framework through empirical network-level benchmarking, added quantitative comparisons with standard ORA and existing tools, expanded validation to additional datasets, and clarified the methodological scope and the role of AI-assisted interpretation. We believe these revisions have significantly improved the rigor, clarity, and positioning of the work.

Below, we provide a detailed point-by-point response to all reviewer comments.

Reviewer Comments:

**Reviewer 1**
The manuscript introduces iterative Gene Enrichment Analysis (iGEA), an extension of over-representation-based gene enrichment methods that sequentially removes genes accounted for by top-ranked terms. This produces non-overlapping, more interpretable enrichment outputs and enables network-based visualization across multiple libraries. The authors further demonstrate how iGEA outputs can be summarized by large language models (LLMs) to accelerate hypothesis generation.

The concept is timely, the implementation appears sound, and the method directly addresses redundance, a long-standing limitation of enrichment analysis. The open-source availability and web interface enhance reproducibility and accessibility. Overall, this work is appropriate for publication after targeted improvements to method description, statistical justification, and comparative validation.

Major Comments (Essential for Revision)

Clarify statistical handling of iterations and p-values.

The text currently states that “adjustment of p-values does not apply to iGEA.” Reviewers will expect justification that the iterative procedure does not inflate Type I error rates.

Please explain whether independence between iterations can be assumed and whether alternative correction schemes (e.g., sequential Bonferroni or permutation testing) were explored.

Response: Thank you for raising this important point. In response, we implemented a network-level empirical framework that provides a statistically sound alternative to applying term-level multiple testing corrections, which are not formally valid under the adaptive iterative structure of iGEA.

In iGEA, enrichment is performed iteratively on adaptively reduced gene lists. At each iteration, the most significant term is selected and its overlapping genes are removed before the next round of testing. Because the input gene list changes at every iteration, the null hypothesis underlying standard multiple testing procedures (e.g., FDR or Bonferroni correction) is not preserved across iterations. The tests are not independent, and the total number of hypotheses tested is not fixed in advance. For this reason, conventional term-level multiple testing correction does not formally apply to the full iterative procedure.

In practice, raw p-values in iGEA are used primarily for ranking terms within each iteration and as a stopping criterion. Only one term is retained per iteration per library, so within-iteration correction would not alter the selection outcome. We considered sequential correction schemes (e.g., sequential Bonferroni); however, these methods assume a fixed hypothesis space and do not accommodate the adaptive reduction of the gene list after each selection step.

To address potential inflation of Type I error rates, we adopted an empirical benchmarking strategy at the network level. Rather than assigning significance to individual terms across iterations, we evaluate whether the resulting cross-library gene–term network exhibits structure beyond chance expectation. We generated 20,000 random gene lists (sizes 50–1000, step size 50; 1,000 per size bin) and applied the complete iterative procedure using the same 11 libraries. For each random run, we computed network-level metrics, including cluster size, number of terms, number of edges, average edges per gene, and number of libraries represented. Observed networks are then compared to the corresponding null distributions using z-scores. This framework directly tests whether the integrated biological organization produced by iGEA exceeds random expectation and provides statistical control at the level of the final network representation.

We have clarified this rationale in the revised manuscript.

Provide explicit algorithmic description.
A short pseudocode box or flow diagram would make the iterative logic transparent (input → rank → remove → repeat → stop when $p > \text{threshold}$).
Define the termination criterion precisely (p-value > 0.01? Overlap < 3 genes?).

Response: Pseudocode added to supplements.

Benchmark against existing redundancy-reduction tools.
To demonstrate added value beyond term clustering or semantic similarity (e.g.,

REVIGO, clusterProfiler simplify()), include at least one quantitative comparison: number of terms retained, mean Jaccard similarity, or semantic similarity score before/after reduction.
Even a small-scale benchmark (GO BP subset) would strengthen the claim that iGEA achieves both compactness and biological fidelity.

Response: We thank the reviewer for this constructive suggestion.

Although iGEA is not primarily designed as a redundancy-reduction method, we agree that quantitative comparison strengthens the manuscript. In the revised version, we added a direct comparison using DAVID's functional clustering framework, which represents a widely used redundancy-reduction approach.

In the HIV dataset, DAVID clustering across six libraries (KEGG, Reactome, WikiPathways, and three GO collections) produced 94 clusters comprising 834 terms spanning 38 pages of results. Applying FDR $< 0.05$ reduced the output to 108 terms (Supplementary Table 9). In contrast, iGEA across 11 libraries produced 71 terms integrated into a single cross-library network with statistically validated structure.

This comparison highlights an important conceptual distinction. Redundancy-reduction tools such as DAVID operate downstream of ORA and cluster overlapping terms within or across libraries. While clustering reduces repetition, the resulting outputs remain extensive and require manual synthesis of term labels. By contrast, iGEA restructures enrichment prior to redundancy accumulation and integrates signals across libraries into a compact gene–term network. The primary advantage therefore lies not merely in reducing overlap, but in producing a statistically benchmarked representation that supports module-level biological interpretation.

We have clarified this distinction in the revised manuscript and explicitly reference the DAVID comparison as a quantitative illustration of the difference in output structure.

Quantify redundancy reduction and coverage.
Report simple metrics such as:
average overlap between enriched terms pre- vs post-iteration,
number of unique genes represented, and
percentage of total input genes retained.
These numbers will make the improvement reproducible and objectively measurable.

Response: We thank the reviewer for this helpful suggestion.

In response, we added quantitative redundancy and coverage metrics to the revised manuscript and to the iGEA application. Specifically, for the HIV dataset (GO Biological Process example), we now report:

- Total number of enriched terms (ORA raw: 57; iGEA: 12).
- Average pairwise gene overlap between retained terms.
- Number of unique genes represented.
- Percentage of input genes covered.

In this example, iGEA reduced the number of enriched terms from 57 (ORA raw) to 12 while maintaining substantial gene coverage (70 of 105 genes, 67%), compared to 103 of 105 genes (98%) in ORA. Importantly, overlap between retained terms was markedly reduced due to the one-gene–one-term-per-library constraint. These metrics are now part of every iGEA run.

Expand discussion of limitations and scope.
Acknowledge that iGEA is currently limited to ORA methods and cannot incorporate gene-level weighting as in GSEA.

Response: We have expanded the Discussion section to explicitly clarify that iGEA is currently restricted to ORA-based enrichment and does not incorporate gene-level weighting or ranking as implemented in methods such as GSEA. Because iGEA relies on iterative removal of overlapping genes, it operates on discrete gene sets rather than ranked lists. For studies in which gene-level effect sizes or ranking information are critical, methods such as GSEA may be more appropriate.

Discuss sensitivity to gene-set size: removing large, broad terms early might suppress smaller, specific ones.

We thank the reviewer for raising this important point.

Because iGEA is iterative, selecting very large, broad gene sets early could remove many genes from the input list while providing limited biological specificity, potentially reducing interpretive resolution. To mitigate this, iGEA applies gene-set size filtering (default 10–600 genes per term) and requires a minimum overlap threshold.

Importantly, examination of gene-set size distributions across the libraries used in this study shows that the majority of curated pathway collections fall within this range (typically 60–100% of terms per pathway library). Very large terms (>600 genes) represent only a small fraction of most libraries (generally <4%), primarily within Gene Ontology and Human Phenotype Ontology collections. Conversely, certain resources such as protein interaction libraries contain many very small sets, which justifies enforcing a lower size bound to avoid unstable enrichment driven by minimal overlap.

Thus, the default size range preserves the majority of biologically curated terms while excluding statistical extremes that are either too broad to be informative or too small to be stable. Because enrichment is performed independently across libraries, related biological processes are often captured through complementary terms in other collections, further reducing sensitivity to any single gene-set size threshold.

We have clarified these design considerations in the Discussion.

Add a short statement on computational cost relative to standard ORA.
Response: We thank the reviewer for this suggestion.

We have added a short statement in the Discussion clarifying the computational cost of iGEA relative to standard ORA. Because iGEA performs iterative ORA within each library, computational time increases approximately linearly with the number of iterations. In practice, if a library yields k iterations, the cost is roughly k-fold compared to a single ORA run for that library. In our implementation, iterations are capped at 30 by default, and typical analyses converge well below this limit.

We note that for typical gene list sizes and the 11-library benchmark configuration used in this study, runtime remains practical on standard hardware. The empirical network benchmarking (1,000 random lists per size bin) is computationally intensive, but this is performed offline to generate null distributions and does not need to be recomputed for routine use.

This clarification has been added to the Discussion section.

Clarify AI interpretation component.
Response: We thank the reviewer for this comment and have clarified the role and scope of the AI interpretation component in the revised manuscript.

Large language models (LLMs) are not part of the statistical procedure of iGEA. All enrichment testing, iteration, and network construction are performed independently. The AI component is applied only after the cross-library gene–term network has been generated and statistically benchmarked.

To evaluate the robustness of AI-assisted interpretation, we analyzed randomly generated gene lists whose iGEA networks exhibited null-like statistical characteristics. As reported in the revised manuscript, ChatGPT was able to generate plausible biological narratives even for these random networks. This highlights an important limitation of generative AI systems: they will attempt interpretation regardless of statistical validity. For this reason, we emphasize that network-level statistical benchmarking is a prerequisite for AI-assisted analysis. Only networks that significantly deviate from null expectations should be interpreted biologically.

At the same time, we observed that when applied to statistically significant iGEA networks, the AI remains largely constrained by the topology and content of the gene–term graph. The structured representation restricts interpretation to explicit gene–term relationships and cross-library clusters. For example, in the vaccination dataset, ChatGPT identified neutrophil-associated terms and, when challenged with the PBMC context, reasoned about potential inconsistencies and proposed alternative cellular explanations (e.g., monocyte involvement). This illustrates grounded, iterative discussion constrained by the network structure.

We further clarify that the iGEA network is particularly suitable for AI-assisted interpretation because it encodes cross-library relational structure rather than unstructured term lists. As network size and connectivity increase, manual synthesis becomes more challenging, whereas structured relational input enables systematic identification of modules and higher-level biological themes. In this framework, iGEA provides statistical grounding and structural constraints, while the LLM serves as an interpretive assistant.

These clarifications have been added to the Results and Discussion sections.

Abstract
Streamline final sentences to emphasize results:
“iGEA reduces redundancy while preserving key biological signals, producing compact, interpretable enrichment networks amenable to AI-assisted hypothesis generation.”

Introduction
Separate the description of library expansion from the discussion of redundancy for improved readability.
When citing LLM-based tools, briefly summarize their current limitations to justify iGEA’s necessity.

Methods
Correct minor typographical issues (“individualy” → “individually”).
Standardize parameter notation: define M, n, N only once.
Ensure that the equation in the hypergeometric description is typeset correctly in final formatting.

Results
Include quantitative summary tables comparing regular GEA vs iGEA (number of terms, unique genes, median overlap).
Clarify figure legends—state which libraries (10 total) were used and what colors represent.

Discussion

Reduce redundancy of the word “redundancy” itself; vary language with “overlapping terms,” “semantic duplication,” etc.
Strengthen link to prior HIV meta-analyses by citing which iGEA-derived clusters correspond to published modules.
Slightly shorten the paragraph beginning “Because different libraries employ…” to maintain focus.

Language and Style
Maintain consistent capitalization for Gene Ontology categories (GOBP, GOMF, GOCC).
Replace informal phrasing (“AI system think”) with formal equivalents (“AI systems may infer”).
Review comma use and eliminate run-on sentences in the Implementation and Discussion sections.

The manuscript meets the general criteria for bioinformatics methods papers: clear motivation, implementable code, and demonstrable utility. Once the statistical justification and comparative analyses are added, the work should satisfy publication standards for journals such as Bioinformatics Advances or BMC Bioinformatics.

Response:

We thank the reviewer for the detailed and constructive section-specific suggestions. All points have been addressed in the revised manuscript as outlined below:

Abstract

The final sentences were streamlined to emphasize the key results and interpretive contribution of iGEA, highlighting compact network generation and suitability for AI-assisted hypothesis development.

Introduction

We separated the discussion of library expansion from the description of overlapping term inflation to improve readability and logical flow.

When citing LLM-based tools, we now briefly summarize their current limitations (lack of statistical grounding and sensitivity to unstructured inputs) to clarify the motivation for iGEA.

##### Methods

Minor typographical errors (e.g., “individualy”) were corrected.

Parameter notation was standardized, and variables (M, n, N) are now defined once and used consistently.

The hypergeometric/Fisher’s exact formulation has been checked for correct typesetting in the final formatting.

##### Results

We added quantitative comparison metrics between ORA and iGEA (number of terms, unique genes represented, and overlap measures) to provide objective evaluation.

Figure legends were clarified to explicitly state which libraries were used and to define color annotations.

##### Discussion

Language was revised to reduce repetitive use of “redundancy,” replacing it with terms such as “overlapping terms” and “semantic duplication.”

The connection to prior HIV meta-analyses was strengthened by explicitly mapping iGEA-derived clusters to published modules.

The paragraph beginning “Because different libraries employ…” was shortened for focus and clarity.

##### Language and Style

Capitalization of Gene Ontology categories (GOBP, GOMF, GOCC) was standardized throughout.

Informal phrasing was revised for formal tone (e.g., “AI systems may infer”).

Comma usage and long sentences in the Implementation and Discussion sections were edited for clarity and concision.

**Reviewer 2**

The authors designed and implemented an application that performs over-representation analysis of genes against dictionaries of ontologies, with the particularity that the result is refined iteratively by removing the best hit, recalculating again until certain convergence level is reached.

While the proposal is interesting, there are many issues that need to be solved before this work can be considered for publication:

MAJOR issues:
* The key aspect of this implementation is the removal of the term that is found more significant and the genes of the input that belong to this term under the hypothesis that by doing that, other overlapping terms (in terms of composition of genes) will be not called after that. To assess the value of this idea, I think that the authors should measure the degree of gene overlap between the terms of the collections that are used in this study. Once it is established how strong is this overlap, we could consider to move forward with this new approach.

* David (https://eur05.safelinks.protection.outlook.com/?url=https%3A%2F%2Fdavidbioinformatics.nih.gov%2F&data=05%7C02%7Cyair%40aionlabs.com%7C997ce598d07e49e8eca708de277c8ba7%7C2a5164ef2a974833b270f537bea92263%7C0%7C0%7C638991613063966422%7CUnknown%7CTWFpbGZsb3d8eyJFbXB0eU1hcGkiOnRydWUsIlYiOiIwLjAuMDAwMCIsIlAiOiJXaW4zMiIsIkFOIjoiTWFpbCIsIldUIjoyfQ%3D%3D%7C0%7C%7C%7C&sdata=QowSYRT2DAaWkCrAnLdvIjl%2F%2FvCUgFXnFTKCgDEGVRw%3D&reserved=0) is another classical ORA tool that should be mentioned in the Introduction. David provides a semantical clusterization of the resulting enriched terms. Please, can you see whether this could be useful for your approach (Discussion)?

Response: We thank the reviewer for this constructive suggestion.

Quantifying overlap among enriched terms. In response, we added quantitative overlap reporting to the iGEA web application. For any analysis run, the tool now reports the overlap between ORA and iGEA outputs, including counts of (i) terms retained uniquely by ORA, uniquely by iGEA, and shared between the two, and (ii) unique genes represented by each method and their overlap. This provides an objective, reproducible summary of how the iterative procedure changes term-level redundancy and gene coverage relative to standard ORA for the same libraries and parameters. We reference these metrics in the revised Results/Supplementary materials.

DAVID comparison. We have added DAVID to the Introduction and Discussion as a representative post-ORA clustering approach. To evaluate its utility in our setting, we performed DAVID functional clustering on the HIV gene list using 6 libraries that overlap with the iGEA analysis (three pathway databases and the three Gene Ontology collections). The DAVID output is provided in the Supplementary Material. While DAVID clusters enriched terms, the resulting output remains a large set of clusters and term lists that still require manual synthesis into a coherent hypothesis.

In contrast, iGEA is designed to generate a compact cross-library gene–term network with empirical network-level statistics, providing a structured substrate for interpretation and AI-assisted module-level summarization.

We have incorporated these additions and clarifications in the revised manuscript.

* Enrichr offers a vast number of libraries (by large higher than MSigDB). In particular, the ontologies of the Transcription section are very interesting as they contain published lists of target genes of ChIP-seq experiments and in silico predictions of transcription factors. The authors should make an effort to introduce these libraries inside their system and perform a comprehensive comparison between their results and the results using Enrichr on the same datasets extracted from the literature

Response: We thank the reviewer for this suggestion.

We agree that transcription factor target collections, including those derived from ChIP-seq experiments and in silico predictions, are highly informative for enrichment analysis. We note that MSigDB already includes regulatory target collections (e.g., C3: transcription factor target gene sets), and users can easily activate those in iGEA.

In the current public web implementation, some of the larger regulatory collections are not enabled by default due to memory and deployment constraints associated with the Streamlit hosting environment. However, the underlying iGEA framework is library-agnostic and supports any gene set collection in GMT format, including MSigDB regulatory sets and Enrichr transcription factor libraries. Users can activate and run these collections locally without modification.

We have clarified in the revised manuscript that iGEA supports regulatory gene-set libraries and that limitations in the web interface reflect deployment constraints rather than methodological restrictions.

* The benchmarking of the tool is very limited. Please, examine the literature in search of works in cancer and development/cell differentiation to build a collection of at least 10 published datasets of genes that can scrutinized with this tool, with Enrichr or David and with GSEA, to demonstrate this new tool improves current state of the art in the GEA field.

Response: We thank the reviewer for this suggestion. We agree that broader benchmarking across multiple cancer and developmental datasets would be valuable. However, the primary goal of this manuscript is to introduce and validate the methodological framework of iGEA, particularly its network-level statistical benchmarking and interpretability advantages, rather than to perform a large-scale compendium study.

Objective benchmarking of “interpretation quality” across 10+ datasets is inherently challenging, as commonly reported metrics (e.g., number of terms, clusters, or genes retained) do not directly measure biological insight, and no gold standard exists for hypothesis-generation performance. In addition, comparisons across ORA, DAVID/Enrichr clustering, and GSEA require harmonizing ranked and unranked inputs, which can substantially affect outcomes.

To strengthen generalizability beyond a single example, we included additional datasets in the Supplementary Materials representing different gene list sizes: a small hypoxia signature and a large vaccination-associated transcriptional signature. This demonstrates that iGEA performs consistently across varying input scales. We also implemented an empirical null framework based on 20,000 random gene lists to provide dataset-independent statistical benchmarking. We believe this combination demonstrates methodological robustness while keeping the manuscript focused.

We agree that assembling a larger benchmarking compendium would be a valuable future direction and is well supported by the publicly available implementation.

* To evaluate the improvement of the results using this new tool, instead of ChatGPT, I would expect a more traditional view in terms of a table with number of genes in common, numer of terms in common, P value of the results, number of new terms, network topology attributes, etc.

Response: As noted above, we have implemented explicit quantitative comparison between ORA and iGEA outputs.

The updated iGEA interface now provides a summary table reporting the number of enriched terms (ORA vs iGEA), shared and unique terms, unique genes represented, and overlap statistics, along with raw and FDR-adjusted ORA p-values alongside iterative p-values. In addition, network-level topology metrics (cluster size, number of genes, number of terms, number of edges, average edges per gene, and number of libraries represented) are reported and benchmarked against empirical null distributions derived from random gene lists.

These additions provide a quantitative framework for evaluating differences between ORA and iGEA. AI-assisted interpretation is presented as a complementary layer built upon these statistical summaries rather than as a substitute for them.

* The number of input genes in the 3 files of the data folder of the web server is really low. Please, can you provide examples with at least 1000-2000 genes to be more realistic?
Response: We thank the reviewer for this comment.

iGEA can technically be applied to gene lists of 1000–2000 genes. However, as gene list size increases, the probability of observing enrichment in random gene lists also increases, leading to larger but less dense networks. This behavior is illustrated in Supplementary Figure 6, where permutation analyses show that network size increases with input size while connectivity and interpretive specificity decrease.

For this reason, we recommend applying iGEA primarily to focused gene lists (e.g., <800 genes) derived from biologically meaningful selection criteria. For very large gene sets, ranked enrichment methods such as GSEA may be more appropriate, or users may reduce list size using more stringent statistical thresholds.

We have clarified this guidance in the revised manuscript.

* I got several execution errors in the web server, when loading the demo provided the first time (after that, the rest of examples worked well):
Error processing library H: Hallmark Gene Sets: [Errno 2] No such file or directory: '/mount/src/iterative-gene-enrichment-app/results/run_20251112_083943_427'
Error processing library C2: Reactome Pathways: [Errno 2] No such file or directory: '/mount/src/iterative-gene-enrichment-app/results/run_20251112_083943_427'
Error processing library C5: Gene Ontology: Biological Process: [Errno 2] No such file or directory:
'/mount/src/iterative-gene-enrichment-app/results/run_20251112_083943_427'
Response: We thank the reviewer for reporting this issue.

The error was traced to a transient file-system and storage limitation in the Streamlit Community Cloud hosting environment, which affected the creation of temporary result directories during the first execution. We have corrected the deployment configuration and implemented more robust directory handling and error checking.

In addition, we generated an optimized version of iGEA specifically for web hosting to ensure stable execution under memory and storage constraints. This web version includes a curated subset of libraries and streamlined functionality for reliability, while the full-featured implementation remains available in the GitHub repository.

The updated web application has been tested extensively and the reported error no longer occurs.

MINOR issues:

* ClusterProfiler (https://eur05.safelinks.protection.outlook.com/?url=https%3A%2F%2Fbioconductor.org%2Fpackages%2Frelease%2Fbioc%2Fhtml%2FclusterProfiler.html&data=05%7C02%7Cyair%40aionlabs.com%7C997ce598d07e49e8eca708de277c8ba7%7C2a5164ef2a974833b270f537bea92263%7C0%7C0%7C638991613063990617%7CUnknown%7CTWFpbGZsb3d8eyJFbXB0eU1hcGkiOnRydWUsIlYiOiIwLjAuMDAwMCIsIlAiOiJXaW4zMiIsIkFOIjoiTWFpbCIsIldUIjoyfQ%3D%3D%7C0%7C%7C%7C&sdata=wRuaPAz%2Beme2BSzgISM38M%2FtJoG9MORL1yHg2KVWLuU

%3D&reserved=0) reference should appear in the initial list of references in the Introduction
Reponse: The clusterProfiler reference has now been added to the Introduction and included in the reference list.

* Iterative  analysis: when the genes of the most significant term are removed from the input list, are they remove as well from the background reference (typically, the full set of genes in the genome) for the statistical analysis?

Response: We thank the reviewer for this important methodological question.

In the current implementation, the background (i.e., the universe of genes represented in the library) remains unchanged across iterations. At each iteration, only the input gene list is reduced by removing genes associated with the previously selected term. The hypergeometric/Fisher's exact test is then recalculated using the updated input list against the same fixed background.

We chose to keep the background constant because it represents the reference universe for enrichment testing (e.g., all genes represented in the library or experiment). Modifying the background at each iteration would change the statistical universe and alter the meaning of the enrichment test. Instead, the iterative procedure conditions on the reduced input list while preserving a consistent reference set.

Given that the background is typically much larger than the input list, removing a subset of input genes from the background would have minimal numerical impact on enrichment probabilities, but would complicate interpretation of the null model. For clarity and consistency, we therefore maintain a fixed background across iterations. This has been clarified in the Methods section.

* Enrichment network: it is not clear whether the connection between the iGEA approach and the network graphics is exclusive or the networking approach can be applied to other ORA tools?

Response: We thank the reviewer for this important clarification.

Network-based visualization of enrichment results is not exclusive to iGEA. Tools such as EnrichmentMap and related approaches construct gene–term or term–term networks from standard ORA outputs to facilitate visualization and redundancy reduction. We have clarified this point in the Introduction and added appropriate references.

The distinction is not the use of networks per se, but the structure of the network produced by iGEA. Because iGEA enforces a non-overlapping gene-to-term assignment within each library, the resulting cross-library gene–term network is substantially less dense than typical ORA-derived networks. In this representation, highly connected regions arise from complementary signals across independent libraries rather than from repeated within-library redundancy. This structural difference enables more meaningful network topology analysis.

Furthermore, iGEA does not rely solely on visual inspection of the network. The network representation serves as a structured, statistically benchmarked substrate for downstream analysis, including AI-assisted interpretation. These distinctions have been clarified in the revised manuscript.

* The website needs a section about Documentation and some Tutorials on how to run a real example and how to interpret the results, with screenshots and text explanations
Response: We thank the reviewer for this important suggestion.

In response, we have expanded the documentation substantially. The web application now includes a direct link to user documentation which is hosted on a private website. The new website has been updated with detailed usage instructions, including step-by-step guidance on running analyses and a case study. The web page is available at https://bivas-benita.org/igea.

We believe these additions significantly improve accessibility and usability for new users.

### Point-by-point response to reviewers

We thank the editor and reviewers for their careful evaluation of our revised manuscript and for the constructive comments. We have revised the manuscript substantially to clarify the scope of iGEA, moderate claims regarding LLM-assisted interpretation, improve statistical framing, clarify the meaning of "non-overlapping," correct inconsistencies, add software versioning information, and fix typographical errors. Below we provide a detailed point-by-point response.

#### Reviewer 1

Comment

Both authors are affiliated with AION Labs, which is listed as the sole funding source for the work, and the senior author is the CTO of this company. The current competing interests statement reads, "The authors declare that they have no competing interests." However, employment by the funding entity should certainly be disclosed as a competing interest, even if it did not create bias. The authors need to make a disclosure in this section.

Response

We thank the reviewer for pointing this out. We agree that the original competing interests statement was incomplete. We have revised the declaration to disclose the relationship with AION Labs and the funding source. The revised statement now reads: "Y.B. is an employee of AION Labs. M.K. is a paid software developer. AION Labs provided funding for this work. The authors declare no additional competing interests."

---

Comment

The authors' response to Reviewer 1's concerns about the statement "adjustment of p-values does not apply to iGEA" dismissed sequential rejection schemes as inapplicable because the hypothesis space is not fixed. However, the authors did not engage with the broader family of procedures designed for adaptive testing, including the Yekutieli FDR, which can handle some dependency. Briefly contextualizing iGEA against this literature would strengthen the manuscript.

Response

We thank the reviewer for this important clarification. We agree that our previous response did not sufficiently contextualize iGEA relative to dependency-aware and adaptive testing procedures. We have revised the Statistical Analysis section to clarify the distinction between correlated tests in a fixed hypothesis setting and the adaptive gene-removal procedure used by iGEA. Dependency-aware approaches such as Benjamini–Yekutieli FDR control can address dependence among tests when the input gene list and hypothesis space are fixed. However, in iGEA, the input gene list is modified after each selected term, so later tests depend on earlier selections. We therefore use raw p-values for iterative term selection and evaluate statistical support at the level of the final cross-collection network using empirical random-gene-list benchmarks.

---

Comment

The authors discuss the LLM results honestly, noting that a random gene list can produce convincing output from the LLM and that the network statistical interpretation is important to limit false convincing output. However, the demonstration on the HIV dataset is arguably an open-book test for the model, since it's a heavily cited paper, where the gene lists and biology are almost certainly represented in GPT-5. The hypoxia and vaccination datasets are also well represented in the training data. A more informative test would involve datasets that the model does not have memorized or a quantitative analysis of LLM interpretation that does not depend on memorization. I would recommend considering the methods performed by Hu et al. (Nat Methods 2025, reference 9 in this work), which compared GPT-4 outputs to curator-assigned MSigDB/GO labels and random gene-set negative controls. The authors make a gesture towards this type of work with three random 300-gene lists but do not quantify the comparison and do not benchmark the usefulness of LLM interpretation. If this is more than weeks of work, I would recommend explicitly stating that the work does not seek to validate LLM interpretation correctness.

Response

We thank the reviewer for this thoughtful comment. We agree that the examples presented in the manuscript should not be interpreted as a formal benchmark of LLM correctness, consistency, or utility. We also agree that memorization of the biological context cannot be excluded, particularly for well-known datasets such as HIV dependency factors, hypoxia response signatures, and influenza vaccination datasets. We have therefore revised the manuscript to state explicitly that we do not attempt to validate LLM interpretation correctness in this work. Instead, we present LLMs as an optional exploratory interface for summarizing and navigating the iGEA network. We also added a limitation noting that future studies using blinded datasets and formal benchmarking frameworks, such as the type of approach described by Hu et al., will be required to assess the correctness, consistency, and utility of LLM-assisted interpretation.

——

Comment

In the results and discussion, the claim that the LLM “correctly inferred a viral infection context” is reported as evidence of interpretive success. However, this has not been quantified, and I have concerns about LLM memorization of these contexts. The authors should consider acknowledging the potential of LLM memorization as a limitation of the demonstrations in this manuscript in the Discussion.

Response

We agree. We have revised the Results and Discussion to avoid presenting LLM output as evidence of validated interpretive success. We replaced language such as “correctly inferred” with more cautious wording, such as “proposed a context consistent with the underlying experiment.” We also added an explicit limitation stating that memorization of biological context by modern LLMs cannot be excluded as a contributing factor in these demonstrations.

——

Comment

Enrichment-analysis tools routinely accept quantitative input alongside gene identifiers. iGEA accepts only unranked binary gene lists. The methods note that “iGEA is not designed for ranked gene lists.” The authors should consider expanding this scope limitation statement to include quantitative inputs.

Response

We agree and have expanded the limitation statement. The revised manuscript now clarifies that iGEA is designed for unranked gene lists and is not intended for ranked gene lists or other quantitative gene-level inputs such as fold changes, expression values, effect sizes, or perturbation scores. We also clarify that methods such as GSEA are designed for ranked inputs and are therefore not directly comparable to iGEA.

——

Comment

Abstract: the authors claim that the “consistency and usefulness of LLM-based interpretation compared with unstructured enrichment outputs.” However, this is not evaluated in this work, as noted above, and I find myself concerned about the possibility of false-but-convincing output, especially in cases the model has memorized. I would recommend supporting this with a quantitative analysis of LLM output correctness, as noted above. Otherwise, I recommend removing this claim of improvement from the abstract if the authors accept this limitation of the work, and instead state that they use LLM-based interpretation for summary and rely on network-level statistics to ensure consistency and usefulness, as they have done in this work.

Response

We agree and have modified the abstract accordingly. We removed the claim that iGEA improves the consistency or usefulness of LLM-based interpretation. The abstract now states that the resulting network structure supports standardized prompting and provides a structured representation for LLM-assisted summarization and exploration of enrichment results. This more accurately reflects the scope of the work.

——

Comment

Background, Results & Discussion: There are typos throughout these sections. The authors should consider using spell-checking tools or double-check their spell-check settings.

Response

We thank the reviewer for identifying these errors. We have corrected the listed typographical errors and reviewed the manuscript for additional spelling and grammar issues.

——

Comment

Results: grammar is off in “a network of gene-term is generated,” consider changing to “a gene-term network is generated.”

Response

We agree and have corrected this sentence to “a gene–term network is generated.”

---

Comment

Results (paragraphs 4-5): There is a large cluster with 179 genes / 62 terms, and then another large cluster with 156 genes / 29 terms is mentioned. The authors should adjust their wording to note the relationship between these networks to avoid the interpretation of these as a discrepancy or fix this discrepancy if it is one.

Response

We thank the reviewer for noting this ambiguity. The manuscript wording was unclear. The largest benchmarked cross-collection iGEA cluster contains 179 genes and 62 terms, as reported in Supplementary Table 5. The separate number of 156 genes and 29 terms refers to the reduced GO-only network shown for visualization clarity, not to the full benchmarked network. We have revised the Results section to make this distinction explicit.

---

Comment

Discussion: The authors mention a cutoff in the discussion for the first time in the Discussion, “gene/edge z-scores ≥ 2-2.5).” If this is a recommendation, this should be specified and justified in a different section.

Response

We agree. To avoid introducing an unsupported threshold in the Discussion, we removed the specific z-score recommendation. The revised manuscript now states more generally that network-level statistics should be considered before downstream interpretation, without prescribing a fixed cutoff.

——

Comment

Discussion: The authors should revise the statement “iGEA addresses this challenge by enforcing a one gene-one term constraint within each library, producing a compact non-overlapping enriched terms” to more closely align with the methods. The authors should at least capture the shrinking gene pool element of the method is missing in this summary for readers who skip to the Discussion.

Response

We agree and have revised the Discussion to more closely match the algorithm. The revised text now explains that iGEA iteratively removes genes associated with the selected enriched term within each gene-set collection and that this creates a one-gene–one-term assignment within each collection. We also clarify that the collection-specific outputs are then integrated into a cross-collection gene–term network, where genes may connect to terms from different collections.

——

Comment

Figure 1: The legend should specify which p-value / FDR thresholds were used to select results for each panel.

Response

We agree and have revised the Figure 1 legend. The legend now states that ORA terms are shown after FDR correction at $q < 0.05$ and iGEA terms are shown using the iterative raw p-value threshold $p < 0.01$.

——

Comment

Software: The authors should reference the version of their tool used in creating this manuscript. At a minimum, the authors should create a release in GitHub and cite that version in the manuscript. The authors could also consider archiving this versioned release in Zenodo and referencing the Zenodo DOI in the manuscript.

Response

We thank the reviewer for this suggestion. We have created a versioned GitHub release for the software used in this manuscript and added the release information to the Availability and requirements section. The manuscript now lists the version used in this study as v1.0.0 and provides the corresponding GitHub release link.

---

Comment

Software: The authors should consider adding a link to the streamlit app in the GitHub. It is easy to find the GitHub from https://bivas-benita.org/igea, but not the reverse. Linking the source code for the website would also help navigate between these repositories.

Response

We agree and have updated the GitHub repository to include links to the Streamlit application and related documentation resources.

---

#### Reviewer 2

Comment

This manuscript presents iterative Gene Enrichment Analysis (iGEA), a software framework intended to address the interpretation bottleneck of standard Over-Representation Analysis. The core idea of iteratively removing overlapping genes to produce a more compact set of enriched terms is conceptually interesting. However, the manuscript suffers from several critical weaknesses that undermine its central claims. The experimental validation is insufficient, relying heavily on a single demonstration case without rigorous comparison to standard methods. The claim regarding improved LLM-based interpretation is entirely unsupported by the provided evidence, as no controlled comparative experiment is described. Furthermore, the statistical benchmarking, while present, does not directly validate the algorithm's ability to uncover masked biological processes. The manuscript requires substantial revision to substantiate its claims.

Response

We thank the reviewer for the detailed critique. In the revised manuscript, we have substantially moderated the central claims and clarified the scope of the study. We no longer claim that iGEA improves the correctness, consistency, or usefulness of LLM-based interpretation. Instead, we present iGEA as a software framework that produces compact, empirically benchmarked cross-collection networks that can be inspected by users and optionally explored using conversational AI. We also clarified the comparison with standard ORA, the meaning of "non-overlapping," the termination criteria, and the interpretation of the statistical benchmark. The revised manuscript now distinguishes between three separate points: (i) iGEA prioritizes enriched terms under an iterative procedure, (ii) empirical random-gene-list benchmarking evaluates whether the resulting network exceeds random expectation, and (iii) LLMs are used only as an exploratory summarization interface rather than as evidence of biological correctness.

---

Comment

The claim that iGEA produces "a compact set of non-overlapping enriched terms" is fundamentally misleading. The methodology describes an iterative, within-library procedure that removes overlapping genes to achieve non-redundancy. However, the final output is a cross-library gene-term network. The supplementary material explicitly states that "each gene appears only once per library source tested but can be linked to multiple terms that originate from different libraries." This means a single gene can be assigned to a term in Library A and a different term in Library B, creating cross-library overlap. The claim of "non-overlapping" is therefore only true within a single library, not for the final integrated network. The manuscript states, "This procedure yields a compact set of non-overlapping enriched terms," which is contradicted by the algorithm's own integration step.

Response

We agree that the original wording was imprecise. The non-overlap constraint in iGEA applies only within each gene-set collection, because genes assigned to a selected enriched term are removed before the next iteration within that collection. The final integrated network is not globally non-overlapping, since genes may connect to terms originating from different collections. We have revised the Abstract, Methods, Results, and Discussion to clarify this distinction. The manuscript now states that iGEA produces non-overlapping enriched terms within each collection, and that these collection-specific outputs are subsequently integrated into a cross-collection gene–term network.

——

Comment

The central claim that the iterative procedure "enables detection of additional biological processes that would otherwise remain masked" is not supported by the experimental design. The manuscript demonstrates iGEA on a single list of 275 HIV dependency factors and asserts that it identifies compact modules. However, no direct comparison to a standard ORA output on the same gene list is provided to show which processes were "masked" and then uncovered by iGEA. The only comparison is with DAVID clustering, not with raw ORA. The text states, "For comparison, we performed functional clustering using DAVID across six libraries... This analysis identified 94 clusters comprising 834 terms." This comparison does not validate the claim of detecting masked processes. Without showing the standard ORA output for the HIV list, the reader cannot verify that the processes found by iGEA were indeed masked in the standard output.

Response

We thank the reviewer for raising this point. We agree that the term "masked" required clarification. We did not intend to imply that iGEA proves the discovery of previously hidden biological processes. Rather, we used this term operationally to describe cases in which dominant overlapping terms in standard ORA, together with multiple-testing correction across many related hypotheses, make additional enrichment signals less visible or not retained after FDR filtering. We have revised the manuscript to clarify this point.

We also emphasize that the manuscript does include a direct ORA–iGEA comparison for the HIV gene list. In the GOBP example shown in Figure 1, standard ORA identified 14 FDR-significant terms, whereas iGEA identified 10 terms using the iterative procedure. Four terms were shared between the methods and six terms were unique to iGEA. These iGEA-specific terms were selected after genes associated with dominant enrichment signals had been removed from the remaining input list. Thus, the data show that iGEA can prioritize enriched terms that are not retained by FDR-filtered ORA. We have revised the text to describe these as additional enriched terms prioritized by the iterative procedure, rather than as definitive proof of previously hidden biology.

---

Comment

The claim that the cross-library network "improves the consistency and usefulness of LLM-based interpretation compared with unstructured enrichment outputs" is entirely unsubstantiated by the provided text. The manuscript describes a demonstration where the iGEA network is given to an LLM with a standardized prompt, but it does not describe any controlled experiment comparing LLM outputs from iGEA networks versus unstructured enrichment outputs. The text states, "To demonstrate how iGEA supports LLM-assisted interpretation, we provided the cross-library iGEA network to a large language model..." This is a demonstration, not a comparative experiment. Furthermore, the manuscript undermines its own claim by showing that "When applied to random gene lists, ChatGPT generated plausible biological narratives despite weaker network structure." This demonstrates that the LLM can produce coherent narratives from random data, directly challenging the claim that iGEA's structured output improves interpretation consistency or usefulness.

Response

We agree with the reviewer. We have removed the claim that iGEA improves the consistency or usefulness of LLM-based interpretation. The revised manuscript instead states that iGEA provides a structured representation that supports LLM-assisted summarization and exploration of enrichment results. We also clarified that the LLM component is exploratory and that we do not benchmark LLM correctness, consistency, or utility in this study. The random gene-list analysis is now explicitly framed as evidence that LLM-generated narratives can be plausible even when the underlying network is weak, reinforcing the need to separate statistical evidence from LLM-generated interpretation.

---

Comment

The manuscript lacks a clear definition of the termination condition for the iterative procedure. The text states the iteration continues "until no additional terms meet the user-defined significance criteria." The specific p-value threshold is not provided in the text, making the claim of “compactness” dependent on an unspecified parameter. A very lenient threshold could still produce a long list of terms.

Response

We agree that the termination condition should be clearer. We have revised the Methods section to specify the threshold used in the analyses reported in the manuscript. Unless otherwise stated, iGEA analyses were performed using an iterative raw p-value threshold of $p < 0.01$. We also note that the maximum number of iterations was capped at 30 during benchmarking. These parameters are now explicitly described in the Methods and figure legends where relevant.

——

Comment

The statistical benchmarking with random gene lists validates that the network structure is non-random, but it does not validate the specific claim that the iterative procedure unmasks true biological processes. The text states, “To establish empirical null distributions, iGEA was applied to random gene lists...” This analysis is irrelevant to the claim of detecting masked processes, as it does not test the algorithm on lists where a dominant signal is known to be present.

Response

We agree that the empirical random-gene-list benchmark does not validate the correctness of any specific biological interpretation or prove that iGEA unmasks true biological processes. We have revised the manuscript to avoid that implication. The purpose of the benchmark is to evaluate whether the resulting cross-collection network structure exceeds random expectation. The claim regarding additional terms has been reframed more narrowly: iGEA can prioritize enriched terms that are not retained by FDR-filtered ORA under the iterative procedure. Biological interpretation of those terms remains hypothesis-generating and should be evaluated in the context of the underlying experiment and domain knowledge.

——

Comment

The manuscript acknowledges a critical limitation that directly contradicts the claim of producing a biologically accurate non-overlapping representation. The text states, “The one-term-per-gene-per-library constraint simplifies interpretation but may partition genes that legitimately participate in multiple processes.” This admission weakens the claim that the resulting set is a valid, non-redundant representation of the underlying biology.

Response

We agree that this limitation should be interpreted carefully. iGEA is not intended to produce a complete or exclusive model of all biological roles for each gene. Rather, the one-gene–one-term constraint within each collection is an intentional prioritization strategy designed to reduce within-collection redundancy and produce a more compact representation for interpretation. We have revised the manuscript to avoid implying that the resulting network is a complete non-redundant representation of the underlying biology. The limitation is now stated explicitly: genes may legitimately participate in multiple processes, and iGEA prioritizes one assignment per collection to simplify interpretation.

---

Comment

The LLM interpretation comparison is confounded by different input formats. For the iGEA network, ChatGPT was given the network structure. For the ORA results, the authors state the network was “difficult to provide directly to ChatGPT due to their size” and instead provided “a JSON list of enriched terms and their overlapping genes.” This is a different input format, not just a difference in structure versus unstructured.

Response

We thank the reviewer for this comment and have revised the manuscript to clarify the comparison. We agree that this analysis should not be interpreted as a controlled benchmark of LLM performance. However, the distinction between the ORA and iGEA inputs was not primarily the file format. A gene–term network can be represented equivalently in DOT, JSON, or other structured formats. The practical difference was that the ORA-derived networks were substantially larger and denser, making them difficult to provide directly to the LLM and difficult to interpret interactively.

More importantly, the topology of ORA-derived networks reflects both within-collection redundancy and cross-collection overlap: genes cluster with many related terms because conventional ORA retains multiple overlapping terms from the same collection as well as overlapping terms across collections. In contrast, iGEA removes assigned genes within each collection before subsequent iterations. Therefore, clustering in the integrated iGEA network arises primarily from genes connecting terms across collections, rather than from repeated within-collection redundancy.

We have revised the manuscript to make this distinction clearer. The ORA example is now presented as a practical illustration of the size and redundancy of conventional ORA-derived networks, rather than as a controlled comparison of LLM performance across input formats. We also removed claims that iGEA improves LLM interpretation accuracy. The revised manuscript instead presents LLM-assisted exploration as a demonstration of how compact cross-collection iGEA networks can be summarized and queried conversationally.

---

Comment

The phrase “non-overlapping enriched terms” is used ambiguously. It should be clarified that this property applies only within a single library, not to the final cross-library network.

Response

We agree and have clarified this throughout the manuscript. The revised text now states that iGEA produces non-overlapping enriched terms within each gene-set collection. The final integrated cross-collection network is not globally non-overlapping, because genes may connect to terms from different collections.

---

Comment

The sentence “meaningful interpretation should be supported by significant cross-library network metrics” is vague. The specific z-score thresholds recommended (2-2.5) are not linked to any demonstrated improvement in interpretation quality.

Response

We agree. We removed the specific z-score recommendation and revised the manuscript to avoid implying that a particular threshold has been validated as improving interpretation quality. The manuscript now states more generally that network-level statistics help determine whether the observed network structure exceeds random expectation and should guide downstream interpretation.

——

#### Reviewer 3

Comment

The submission reports a method for the characterization of non-overlapping gene ontology terms. The approach is reasonable, if a bit aggressive, as it iteratively assigns gene sets to distinct ontology sets, so the "rich-get-richer" problem is ameliorated with respect to ontology categories but exacerbated with respect to individual ontologies (e.g., if a single gene has multiple roles/ is involved in multiple ontologies, only one will be retained at the expense of others).

Response

We thank the reviewer for this thoughtful assessment. We agree that the one-gene–one-term assignment is a bit aggressive when viewed within a single gene-set collection, because genes may legitimately participate in multiple biological processes. However, this constraint is applied only within each collection. A central design feature of iGEA is that the collection-specific outputs are subsequently integrated into a cross-collection gene–term network, where the same gene can connect terms from different collections.

Thus, information that may be simplified within one collection can be represented at the network level through complementary annotations from other collections. This is one of the main differences between iGEA and conventional ORA. In standard ORA, interpretation often becomes more difficult as the number of collections increases, because the user must inspect increasingly large and redundant term lists. In iGEA, adding collections can instead enrich the cross-collection network by providing additional biological perspectives while limiting within-collection redundancy.

We have revised the manuscript to clarify that iGEA is not intended to provide a complete model of all gene functions within a single ontology. Rather, it uses within-collection prioritization followed by cross-collection integration to generate a compact but biologically contextualized representation for interpretation. We also retain the potential loss of within-collection multifunctionality as an explicit limitation in the Discussion.

——

Comment

The determination of nonredundant GO projections is not a new problem, and other papers have attempted to treat it, qualitatively or quantitatively (https://journals.plos.org/plosone/article?id=10.1371/journal.pone.0024289, https://academic.oup.com/bioinformatics/article/28/16/2209/325325, https://journals.plos.org/plosone/article?id=10.1371/journal.pone.0021800). The paper would bear briefly discussing how this approach is different from those—as well as, potentially, conceptual similarities to other computational methods that simplify networks by only prioritizing the top link and dropping the rest (occasionally used in GRNs and elsewhere).

Response

We thank the reviewer for this helpful suggestion. We agree that reducing redundancy among Gene Ontology terms is a well-recognized problem and that several prior methods have addressed this using ontology structure, semantic similarity, or related approaches. We have revised the Background to acknowledge this prior work and to cite representative examples.

At the same time, we clarify that iGEA is not designed specifically as a GO-term redundancy-reduction method. The motivation for iGEA is broader: modern enrichment analyses often involve many gene-set collections spanning pathways, ontologies, regulatory signatures, disease signatures, and protein interaction resources. Gene Ontology represents only a subset of this space, and GO-specific redundancy approaches do not by themselves solve the challenge of integrating enrichment results across many collections.

We therefore discuss prior GO redundancy-reduction methods briefly, but we do not expand this into a detailed comparison because the main contribution of iGEA is different. iGEA applies an iterative within-collection enrichment procedure and then integrates the resulting collection-specific outputs into a cross-collection gene–term network. This design is intended to support interpretation across multiple gene-set collections, rather than to provide another GO-only projection or semantic-similarity method.

——

Comment

I strongly recommend against introducing the word “library” here. This is not standard nomenclature, and “library” already means two different things in bioinformatics (a computational package and a set of molecules). If I understand correctly, the authors use “library” to mean “database.”

Response

We thank the reviewer for this suggestion. We have revised the terminology throughout the manuscript to use “gene-set collection” or “collection” in the main methodological descriptions and claims. In a few places, we retain “library” only where it reflects common usage in existing resources or software contexts, but we now define the intended meaning clearly and use “collection” as the primary term.

——

Comment

I do not have much to say about the ChatGPT component. The authors recognize that the results could be somewhat helpful or meaningless (based on a random gene list). Caveat emptor.

Response

We agree with the reviewer’s caution. We have revised the manuscript to emphasize that LLMs are not statistical engines and that plausible LLM-generated narratives can arise even from random gene lists. The revised text presents conversational AI as an optional exploratory interface for summarizing and navigating iGEA networks, not as independent evidence of biological correctness.

—

Comment

I think there is an interpretive context here that is alluded to but not discussed much. The challenge is defined with reference to HIV etiology screening, and the historical issues in coming up with a reproducible gene list, moving on to the discovery of reproducible GO motifs. Even this So the ostensible problem is integrating data across studies or datasets rather than databases. But the study does not actually address this problem—it only looks at merging together different databases for qualitative downstream analysis. Therefore, I think the paper would be significantly strengthened by taking this motivation to its logical conclusion and explaining how the method addresses the issue of poor reproducibility; otherwise, it reads as a solution to a problem that has not been motivated.

Response

We thank the reviewer for this important clarification. We agree that iGEA is not designed to integrate multiple studies or resolve cross-study reproducibility directly. The reason for using the HIV dependency-factor dataset is different. The HIV screening literature provides a useful benchmark because several independent genome-wide screens produced poorly overlapping gene lists, yet subsequent meta-analyses showed that reproducible biological signals could be recovered at the pathway and process level.

Thus, the HIV example allows us to ask whether iGEA, when applied to a single well-characterized screen, can recover pathway-level biology that was later supported by multiple studies and meta-analyses. In this sense, prior HIV meta-analyses provide an approximate pathway-level reference for evaluating interpretability. We have revised the manuscript to explain the motivation of comparing iGEA to cross-study meta-analysis.

—

Comment

Minor issues: Various typos. Non-overalpping, amd, gens and edges, netowrks, MSigDB ([4] version, “Supplementrary Figure 8” (also should be table 8?; possibly some other issues with mislabeling), “that iGEA cluster increase”. Not sure why MS Word does not pick up most of them.

Response

We thank the reviewer for noting these issues. We corrected the listed typographical and formatting errors and reviewed the manuscript for additional spelling, grammar, and reference-formatting issues.

---

#### Reviewer 4

Comment

The permutation test was used to demonstrate the statistical significance of the identified network. However, as the authors noted in their response to Reviewer 2 (“The benchmarking of the tool ...”), the statistics used in the permutation test do not directly measure biological insight or interpretability. Therefore, the manuscript should avoid making overly strong claims regarding statistically grounded biological interpretation. Nevertheless, the comparisons between iGEA and conventional ORA remain informative and valuable for readers and potential users of the method.

Response

We agree with the reviewer. We have revised the manuscript to avoid implying that network-level statistical significance directly measures biological insight or interpretability. The revised text states that empirical network benchmarking evaluates whether the observed cross-collection network structure exceeds random expectation. We also moderated claims regarding “statistically grounded biological interpretation” and now describe the output as an empirically benchmarked network representation that can guide downstream interpretation. The comparisons between iGEA and conventional ORA have been retained and clarified.

---

Comment

If the statistical claims are moderated or removed, the manuscript could potentially be shortened and more tightly focused on the iGEA methodology and its practical utility.

Response

We agree. In response, we moderated the statistical and LLM-related claims and focused the manuscript more tightly on the iGEA methodology, the ORA–iGEA comparison, the empirical network benchmark, and the practical utility of the software. We also clarified that the LLM component is an exploratory interface for summarization and hypothesis generation rather than a validated measure of biological correctness.

### Editorial decision and reviewer comments — BMC Bioinformatics, 25 August 2026

Ref: Submission ID 6e5f1a44-cee9-4011-8305-0f068bcd1604

Dear Dr Benita,

Your manuscript "Iterative Gene Enrichment Analysis: interpretable networks for human and AI-assisted biological insights" has now been assessed. If there are any reviewer comments on your manuscript, you can find them at the end of this email.

Regrettably, your manuscript has been rejected for publication in BMC Bioinformatics.

---

Handling Editor: Alireza Khanteymoori Recommendation: Reject

---

Thank you for the opportunity to review your work. I'm sorry that we cannot be more positive on this occasion and hope you will not be deterred from submitting future work to BMC Bioinformatics.

Kind regards,

Payel Karmakar Assistant Editor BMC Bioinformatics

Reviewer Comments:

Reviewer 2 The revision process is fundamentally compromised because the submitted "Revised Manuscript" is not the actual revised paper; it is an email invitation to re-review, containing no manuscript content. Consequently, none of the claimed revisions, such as corrected typos, revised abstract, updated figures, or terminology changes, can be verified. This absence of the primary document prevents any meaningful assessment of the revision's quality or completeness, and it is a critical failure that undermines the entire review process. The response letter itself also contains typographical errors, which further raises concerns about the rigor of the revision.

Given the critical failure to provide the actual revised manuscript, the manuscript cannot be accepted or considered for further revision at this stage. The authors must resubmit the complete revised manuscript along with a detailed response letter, ensuring that all claimed changes are clearly visible and verifiable. Until the actual manuscript is provided, the review cannot proceed, and the current submission is rejected.

Reviewer 3 My questions have addressed and I have no further comment.

Reviewer 4 Overall, iGEA iteratively identifies the top-enriched term from over-representation analysis by removing the genes associated with terms identified in previous iterations. However, the fundamental rationale for this procedure and, more importantly, evidence that it improves biological inference remain unclear.

The authors argue that a shorter list of terms facilitates interpretation, but a shorter list does not necessarily capture the underlying biology more accurately. This is particularly important given the authors' own observation that LLMs can generate plausible responses even from arbitrary sets of genes and terms. The key question is therefore not whether iGEA produces a concise and plausible interpretation, but whether it produces a biologically meaningful one.

My main concern is that iGEA addresses the problem of reducing the number of terms without sufficiently demonstrating that it addresses the more important problem of identifying biologically relevant signals. Without convincing statistical and biological validation, I remain unconvinced that the method provides a substantive advance beyond generating a shorter list of terms for interpretation.
