## Supplementary File - Previous manuscript versions for "Iterative Gene Enrichment Analysis: interpretable networks for human and AI-assisted biological insights"

### Previously submitted manuscript versions

Maxim V. Kuleshov, Yair Benita — AION Labs, Rehovot, Israel

#### What this document contains

The three manuscript versions submitted to peer-reviewed journals before this preprint was posted, reproduced in full and in chronological order. Intermediate working drafts are not included; every version reproduced here was sent to a journal. Dates are taken from the journals' submission status histories.

The reviewer reports and our point-by-point responses are in the companion file, *Supplementary File - Peer review history.*

| | Submitted | Journal | Length | Outcome |
|---|---|---|---|---|
| **Submission 1** | Aug 2025 | *Bioinformatics* | 3,312 words | Declined without review |
| **Submission 2** | 20 Sep 2025 | *BMC Bioinformatics* | 3,563 words | Rejected; appeal granted |
| **Submission 3** | 2 Apr 2026 | *BMC Bioinformatics* | 5,122 words | Revision requested |

**Submission 1** — *Bioinformatics* (Oxford University Press), manuscript BIOINF-2025-2109. Declined without external review on 8 September 2025.

**Submission 2** — *BMC Bioinformatics*, submission ID 6159728f. Received 20 September 2025. An amendment was requested at technical check and received on 22 September 2025; the manuscript reproduced here is that amended version, which is the one that went to review. Two reviewer reports were received on 11 November 2025 and the manuscript was rejected on 19 November 2025. A formal appeal was granted.

**Submission 3** — *BMC Bioinformatics*, submission ID 6e5f1a44. Received 2 April 2026. Four reviewer reports were received on 13 May 2026 and a revision was requested on 28 May 2026.

The revision that followed was received by the journal on 13 June 2026 and rejected on 25 August 2026. That revision, with subsequent corrections, is the preprint posted here (6,003 words). From the first submission to *BMC Bioinformatics* to the final decision, the manuscript was under consideration for eleven months.

#### How the manuscript changed

**Submission 1 to Submission 2.** Reformatted from the *Bioinformatics* application-note format to the *BMC Bioinformatics* research-article format. No external review had been received, so the scientific content was substantially unchanged.

**Submission 2 to Submission 3.** The largest change in the manuscript's history, and the reason the text grew by roughly half. Prepared in response to the two reviewer reports of 11 November 2025 and to the formal appeal that followed. Additions included an empirical benchmarking framework based on random gene lists, quantitative comparison between over-representation analysis and iGEA, comparison against DAVID functional clustering, validation on two further datasets, and clarification of the statistical methodology and of the role of AI-assisted interpretation.

**Submission 3 to this preprint.** Prepared in response to the four reviewer reports of 13 May 2026. Claims regarding large language model interpretation were moderated, the statistical framing was clarified, the meaning of "non-overlapping" was made precise, the competing interests declaration was corrected to disclose the authors' relationship with AION Labs, software versioning was added, and typographical errors were corrected. The point-by-point response documenting these changes is in the companion peer review history file.

*Bioinformatics*, YYYY, 0–0
doi: 10.1093/bioinformatics/xxxxx
Advance Access Publication Date: DD Month YYYY
Manuscript Category

*Subject Section*

### Iterative Gene Enrichment Analysis: interpretable networks for human and AI-assisted biological insights

Maxim Kuleshov[1], Yair Benita[1*]

[1]AION Labs, 4 Oppenheimer st. Rehovot, Israel.

*

Associate Editor: XXXXXXX

Received on XXXXX; revised on XXXXX; accepted on XXXXX

**Abstract**

**Motivation:** Gene set enrichment analysis (GEA) is a standard approach for interpreting gene lists from omics studies. While widely used, generating a biological hypothesis from lists of enrichment terms is increasingly difficult due to expansion of gene set collections. The same subset of genes often drives significance across many terms in each database, producing long, redundant lists of terms. Large language models (LLMs), have been employed to address this challenge but were found to lack the precision, stability and statistical grounding of enrichment methods, making them useful as summarizers rather than analytical engines. This underscores the need for methods that adress gene set redundancy while retaining statistical rigor.

**Results:** We present iterative Gene Enrichment Analysis (iGEA), a method that sequentially removes genes explained by the top-ranked term in each collection and re-runs enrichment until no significant terms remain. This iterative process produces a compact, non-overlapping set of enriched terms that is easier to interpret. Cross-collection results are represented as a gene–term network, highlighting hubs and clusters that capture key biological processes. Applying iGEA to a hypoxia gene signature and a screen for HIV required host factors demonstrated substantial reduction in redundant terms while preserving biologically meaningful findings. Moreover, structured networks generated by iGEA can be readily summarized by LLMs, enabling rapid, coherent hypothesis generation within minutes. Together, iGEA enhances the interpretability of enrichment analysis and provides a reproducible framework for integrating human expertise with AI-assisted interpretation.

**Availability:** Source code is available at https://github.com/aion-labs/Gene-Enrichment-Analysis.

**Contact:**

**Supplementary information:** Supplementary data are available at *Bioinformatics* online.

#### 1 Introduction

High-throughput molecular profiling studies frequently generate gene lists of interest, most commonly from differential expression analysis, genome-wide association studies, or large-scale perturbation screens. A standard approach to interpret gene lists is gene set enrichment, where the input is compared to curated gene sets to identify biologically informative overlaps. Two broad classes of methods are commonly used: over-representation analysis (ORA), such as the hypergeometric test (Falcon and Gentleman, 2007; Kuleshov *et al.*, 2016), and Gene Set Enrichment Analysis (GSEA), which analyzes ranked gene lists (Subramanian *et al.*, 2005). In this study, we focus on ORA-based methods. The goal of all enrichment approaches is to formulate testable biological hypotheses. To reduce redundancy and improve interpretability of ORA results, we introduce iterative Gene Enrichment Analysis (iGEA).

For the past two decades, the statistical foundations of Gene Enrichment Analysis (GEA) have changed little. Progress has instead come from expansion of gene set databases, such as MSigDB (Liberzon *et al.*, 2011) and Enrichr (Kuleshov *et al.*, 2016), which now contain 30 and

218 collections, respectively (as of August 2025), covering transcriptional regulation, pathways, disease signatures, and functional ontologies. However, this proliferation introduces major interpretability challenges: the same subset of genes often drives significance across dozens of terms, leading to long, redundant lists of enriched terms. In addition, due to false discovery rate (FDR) correction for multiple testing, smaller subsets of genes within the input do not reach the significance threshold. As a result, researchers must manually prioritize redundant terms while missing others, which complicates hypothesis generation.

Large language models (LLMs) have recently been explored as a complementary strategy. For example, GPT-based systems such as ChatGPT can generate plausible narratives about the functions of enriched gene sets (Hu *et al.*, 2025; Joachimiak *et al.*, 2024). While useful as summarizers, these models were found to lack the precision, stability, and statistical grounding of GEA approaches.

Here, we present iterative GEA (iGEA), which is designed to address gene set redundancy and leverage the diversity of gene set collections. Iterative GEA starts with a regular GEA within a specific collection. After identifying the most significant term, overlapping genes are removed from the input list and the analysis is repeated until no significant terms remain. This approach frames enrichment as a different question: beyond the genes already explained, what else remains? Iterative GEA yields a compact, non-overlapping set of enriched terms.

By design, Iterative GEA produces fewer, more interpretable results than regualr GEA and allows for cross-collection visualization as a gene--term network, where each gene can connect to only one enriched term per collection. The cross-collection interaction through genes is an intuitive way to explore the biology driven by groups of genes. We further demonstrate that such networks not only simplify interpretation but also provide a structured framework for conversational exploration with LLMs, enabling rapid generation of coherent biological hypotheses.

#### 2 Methods

**Gene Enrichment Analysis**

To calculate gene set enrichment a hypergeometric test was conducted using SciPy's hypergeom.sf() function, which returns the probability of observing at least the given overlap by chance. For each gene set collection, we first filtered gene sets to include only those within the specified size range selected by the user (10-600 genes per gene set by default). The range was chosen to exclude genes sets that are likely to be less informative. Depending on the parameters chosen, a library specific input and background was calculated as the intersection between the global input or background and the gene sets to be tested within the library. The hypergeometric function was used with the following parameters: (n_overlap - 1, M, n, N) where n_overlap is the observed number of overlapping genes minus one, M is the collection-specific background size, n is the number of genes in the specific gene set being tested, and N is the library specific input gene list size. As an example, if there are 400 genes in the input gene list and 350 of them in the gene sets to be tested, the library specific input size was 350. Similarly, if the background included 10,000 genes but only 4500 were represented in the filtered library, the library specific background was set at 4500. If the a specific pathway term being tested had 50 genes and 15 genes overlap with the library specific input, the p-value will be calculated as hypergeom.sf(15-1, 4500, 50, 350). For the purpose of this study, all regular enrichment analyses were performed using a raw p-value cutoff of 0.01, a minimum of 3 genes overlapping with a gene set, and gene set size in the range of 10 to 600.

**Iterative Gene Enrichment Analysis**

Iterative GEA analysis was performed on each gene set library collection individualy. The process begins with a regular enrichment analysis on the input gene list. The term with the most significant p-value is selected and recorded, and the genes that overlap with this term are removed from the input gene list. This process repeats iteratively: each time, the most significant remaining term is identified and saved, while its overlapping genes are eliminated from the input list. The iteration continues until no significant terms remain that meet the defined p-value threshold.

**Gene Set Libraries**

Gene set libraries were obtained from the MSigDB ((Liberzon *et al.*, 2011) version 2025.1. The database contains many different types of collections and only canonical pathways and Gene Ontology collections were used.

A gene set library was created for protein interaction data using STRING-DB (Szklarczyk *et al.*, 2023), a protein interaction database. Human protein-protein interaction data was obtained from v12.0 filtering for high-confidence interactions with score ≥ 0.9. Each protein in the database was defined as "term" and its interacting partners were defined as the gene set. The resulting library included 12,323 gene sets covering 12,320 genes with an average of 16.4 genes per set. The library was added in GMT format for compatibility with MSigDB.

**Enrichment Network**

The enrichment network allows cross-library interpretation. Iterative GEA results are integrated into by building a DOT language (Graphviz.org) network. The network has 2 types of nodes, a gene name from the input list and a term from the library collection. Edges are only created between genes and terms in which they were identified as significantly enriched and indicate membership. No weights are applied to the edges and nodes are colored by library collection. The network is visualized using Plotly (Plot.ly).

**Interpretation by Artificial Intelligence**

To demonstrate the interpretability of iGEA networks, we provided the results to a large language model (GPT-5, OpenAI) with a standardized prompt that is generated by the enrichment application. A conversational AI can platform can assist biologists in exploring network results and specific clusters or topics within. The standardized prompt is available in Supplementary Table 1.

#### 3 Results

Two previous studies were selected that generated a gene list for biological hypothesis generation and analyzed them in depth. The first, a gene expression signature associated with hypoxia where 5 different cell lines were used to derive core molecular response to hypoxia (Benita *et al.*, 2009). The second, a high throughput screen for host factors required for HIV infection (Brass *et al.*, 2008).

##### 3.1 A core molecular response to hypoxia

The core molecular response to hypoxia included 81 genes of which 80 were mapped to a currently valid gene symbol by the application. Enrichment was calculated for Hallmark and each of the C2 and C5 collections of MSigDB. Overall, 174 terms across 11 collections passed the filtering results (Supplementary Table 2A-B). Applying a false discovery rate (FDR) cutoff of 0.05, 57 terms remained across 8 different collections. Results for Gene Ontology Biological Process are shown in Figure 1. Applying a FDR cutoff of 0.05, 27 terms were significantly enriched using the regular enrichment method while 8 were identified using iGEA. Only 4 terms were identified in both enrichment methods. Importantly, the 4 shared terms were ranked differently and far from one another in the regular enrichment result.

For simplicity, four collections were selected for generating an enrichment network based on the iGEA method: Reactome pathways, Wiki Pathways, Gene Ontology Biological Process and Protein Interactions (Supplementary Table 2C). The network included 3 clusters, the largest is shown in Figure 2. The network included terms associated with response to oxygen, glycolysis, collagen formation and metabolic stress. An AI prompt was passed to ChatGPT 5 with the full network with a request for interpretation (Supplementary Table 2D). The following is the first sentence of the executive summary of ChatGPT 5: “This network resolves into three dominant biological programs: (i) glycolytic rewiring/pyruvate metabolism, (ii) extracellular matrix (ECM) collagen biosynthesis and maturation, and (iii) hypoxia–stress signaling with RNA-processing changes”. As part of the template, ChatGPT is required to estimated the experimental context that led to this result: “Primary or immortalized fibroblasts cultured under hypoxia (±TGF-β), leading to enhanced glycolysis, collagen production/maturation, and hypoxia/ER stress responses with supportive changes in RNA splicing and chromatin state”. Following the analysis, ChatGPT was asked to explain how thermogenesis is associated with hypoxia. The summary of the response was: “While hypoxia typically downregulates canonical thermogenesis, your network suggests a subtler, regulatory adaptation: hypoxia may engage adaptive thermogenic-like pathways—through circadian, metabolic, and signaling nodes—to modulate energy balance and survival in stressed cells. This form of metabolic reprogramming may reflect an emergent, non-canonical thermogenic response that maintains homeostasis under oxygen limitation”.

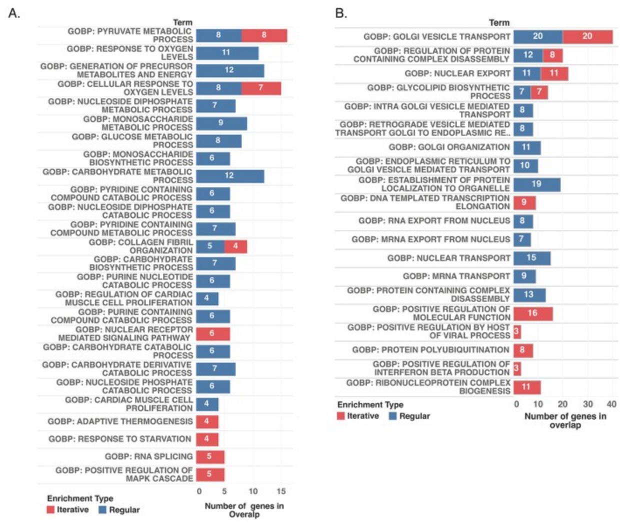


**Fig. 1. Regular and iterative GEA.** A. Gene Ontology Biological Process results of regular (blue) and iterative (red) enrichment analysis. For regular enrichment, an FDR cutoff of 0.05 was applied. The size of the bar and numbers within the bar indicate the number of genes overlapping with the identified gene set. Terms are sorted by p-value for regular enrichment analysis and if not available by iterative p-value. Panel A depicts the Hypoxia signature enrichment while panel B HIV enrichment.

#### 3.2 Host proteins required for HIV infection

In a genome-wide screen to knockdown human proteins using siRNA, 275 genes where identified that when knocked down reduce significantly HIV infection. The 275 genes were subject to a regular and iterative GEA (Supplementary table 3A-B). Regular enrichment identified a total of 417 terms across the 12 libraries, of which 148 terms across 7 libraries had FDR<0.05. iGEA had 78 significant terms across 12 libraries. Golgi and Endoplasmic Reticulum terms were dominant in for Gene Ontology Biological Process in both regular and iterative GEA. However, while the top 4 terms were identified by both methods the remaining terms were unique (Figure 1B). This observation was typical for other libraries.

The iGEA network was built from Reactome pathways, Wiki Pathways, Gene Ontology Biological Process and Protein Interaction for consistency with the hypoxia network and also to increase the interpretation challenge for ChatGPT. The network had one large cluster that included Golgi and transcription terms as well as some immune related terms (Figure 2B and Supplementary Table 3C).

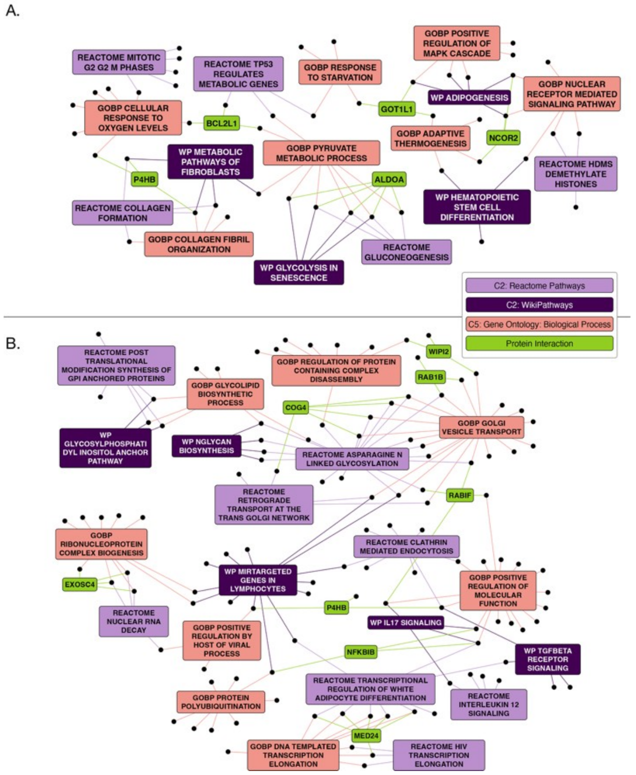


**Fig. 2. Iterative GEA network.** Iterative GEA results shown as a network limited to four libraries (Reactome, Wiki Pathways, Gene Ontology Biological Process and Protein Interactions). Genes from the input list overlapping with a term are shown as black dots. The networks shown are only the largest cluster for Hypoxia (A) and HIV (B). The nodes and edges are colored by library name.

The iGEA network was loaded to ChatGPT for interpretation (Supplementary Table 3D). The first sentence of the executive summary was: “This network resolves into three tightly connected biological modules: (i) a secretory/trafficking axis (ER/Golgi glycosylation, GPI-anchor synthesis, Golgi–TGN–endosome transport), (ii) a nucleus-centered RNA/transcription axis (transcription elongation, nuclear export, RNA exosome), and (iii) an innate/immune signaling axis (NF-κB, IL-12/IL-17, type-I interferon)”. The proposed biological hypothesis was: “An immune cell population (likely CD4⁺ T cells or myeloid cells) underwent innate inflammatory activation that simultaneously engaged NF-κB and type-I interferon programs, necessitating extensive remodeling of nucleocytoplasmic transport and the secretory pathway to produce and traffic immune mediators and receptors". Next, we explained to ChatGPT the context of the HIV experiment and asked to modify the analysis. ChatGPT mapped the results to 10 different processes in the HIV life cycle and provided the following as a summary: The screen recapitulates the entire HIV life cycle in host dependency terms: (i) Entry/trafficking → nuclear import → transcription/export → protein processing → assembly/release.(ii) The highest-degree hubs (e.g., NUPs, MED subunits, COG/STX5/RAB GTPases, NF-κB regulators, EXOSC components) represent multi-stage bottlenecks — disrupting them affects HIV at more than one step".

#### Discussion

For more than two decades, scientists have relied on gene set enrichment analysis (GEA) to interpret results from diverse molecular studies. The goal has always been twofold: to recover known biology as a positive control, and to generate new hypotheses for future investigation. The hypoxia and HIV studies we revisit here are relatively old but were chosen because the original analyses, performed by YB, took months to complete, were experimentally validated, and were published in peer-reviewed journals.

With the proliferation of libraries and gene sets, it has become increasingly difficult, even for experienced scientists, to derive a clear biological hypothesis from the output of conventional GEA. As shown in Figure 1, results often span long lists of overlapping terms. Most users will intuitively gravitate to those that "fit the story," such as response to oxygen in hypoxia or interferon/viral terms in HIV. A common workaround is to restrict the analysis by using fewer libraries and applying more stringent p-value thresholds. While this reduces the number of terms, it also narrows the results to only the most dominant enrichments, often not the most informative. For instance, in the HIV study, the Golgi apparatus consistently emerges as the top hit. Yet this is potentially misleading: damaging Golgi functionality would reduce all receptor levels, including CD4, and thereby block HIV infection. In other words, the most statistically significant term may not be the most biologically relevant.

The iGEA method addresses this challenge by producing a clean list of non-overlapping gene sets per library. Each term can then be viewed as a distinct piece of the biological puzzle. Top results still overlap with conventional enrichment (Figure 1), but iGEA also recovers additional, functionally meaningful terms that are often lost due to FDR penalties. In the hypoxia dataset, for example, iGEA identified five such terms, including response to starvation, which are clearly relevant when inspected in the network (Figure 2A).

A major advantage of iGEA is its natural representation as a gene–term network. Because each gene is assigned to only one term per library, results from multiple collections can be integrated into a coherent whole. In the hypoxia example, some libraries highlight HIF-1, the master regulator of hypoxia; others emphasize glycolysis, the alternative energy source when oxygen is scarce; while others point to the mitochondria, the cell's main oxygen consumer. Viewed together in a network, these complementary insights allow scientists to identify hubs and assemble a mechanistic hypothesis with far greater clarity.

Another strength of this representation is its compatibility with large language models (LLMs). In our proof-of-concept, ChatGPT was able to explain the core biology of iGEA networks within seconds, capturing both expected and unexpected patterns. The most powerful feature, however, was the interactive exploration of results. For example, by prompting ChatGPT to explore the role of thermogenesis in the hypoxia network, we uncovered a potentially relevant angle that had been overlooked in the original study.

Of course, limitations remain. There is no gold standard for benchmarking methods that interpret gene lists. Results depend on choices of libraries, cutoffs, and filters, and no analysis can claim universal truth. Enrichment methods are best viewed as assistants that help biologists formulate hypotheses. From this perspective, we find iGEA's network-based outputs more intuitive and interpretable than long lists of overlapping terms, and importantly, we show that LLMs excel in analyzing and contextualizing this type of data.

#### Acknowledgements

We thank Code Ocean for their support. The original streamlit code was adopted from Code Ocean and written by MK while at Code Ocean. The iGEA method was added with support of AION Labs. Features and UI improvements and tests were added by YB using Cursor (www.cursor.com). Code Ocean also supported this study by providing computing infrastructure and by hosting the application on codeocean.com.

#### Data Availability

Data and code used for this study are available in https://github.com/aion-labs/Gene-Enrichment-Analysis. The application is available on Code Ocean at xxx and is also hosted on the streamlit community cloud for everyone to use.

#### Funding

This work has been supported by AION Labs.
*Conflict of Interest:* none declared.

### Iterative Gene Enrichment Analysis: interpretable networks for human and AI-assisted biological insights

Maxim V. Kuleshov[1], Yair Benita[1*]

[1]AION Labs, 4 Oppenheimer street, Rehovot, Israel.

*

##### Abstract

**Background:** Gene enrichment analysis (GEA) is a standard approach for interpreting gene lists from omics studies. While widely used, generating a biological hypothesis from lists of enrichment terms is increasingly difficult due to the expansion of gene set libraries. The same subset of genes often drives significance across many terms in each library, producing long, redundant lists of terms. Large language models (LLMs) have been employed to address this challenge but were found to lack the precision, stability, and statistical grounding of enrichment methods, making them useful as summarizers rather than analytical engines. This underscores the need for methods that address gene set redundancy while retaining statistical rigor.

**Results:** We present iterative Gene Enrichment Analysis (iGEA), a method that sequentially removes genes explained by the top-ranked term in each library and reruns enrichment until no significant terms remain. This iterative process produces a compact, non-overlapping set of enriched terms that is easier to interpret. Cross-library results are represented as a gene-term network, suitable for network topology analysis that results in hubs and clusters that capture key biological processes effectively. Applying iGEA to a gene list of HIV-required host factors demonstrated a substantial reduction in redundant terms while preserving and expanding biologically meaningful findings. Moreover, structured networks generated by iGEA were effectively summarized by LLMs, enabling rapid, coherent hypothesis generation within minutes. We concluded that iGEA enhances the interpretability of enrichment analysis and provides a reproducible framework for integrating human expertise with AI-assisted interpretation.

**Availability:** Source code is available at https://github.com/aion-labs/Gene-Enrichment-Analysis and application available online on streamlit community cloud at: https://iterative-gene-enrichment.streamlit.app.

**Contact:**

#### Background

High-throughput molecular profiling studies frequently generate gene lists of interest, most commonly from differential expression analysis, genome-wide association studies, or large-scale perturbation screens. A standard approach to interpret gene lists is gene set enrichment, where the input is compared to curated gene sets to identify biologically informative overlaps. Two broad classes of methods are commonly used: over-representation analysis (ORA), such as the hypergeometric test [1, 2], and Gene Set Enrichment Analysis (GSEA), which analyzes ranked gene lists [3]. In this study, we focus on ORA-based methods. The goal of all enrichment approaches is to formulate testable biological hypotheses. To reduce redundancy and improve interpretability of ORA results, we introduce iterative Gene Enrichment Analysis (iGEA).

For the past two decades, the statistical foundations of Gene Enrichment Analysis (GEA) have changed little. Progress has instead come from expansion of gene set databases, such as MSigDB [4] and Enrichr [2], which now contain 30 and 218 collections, respectively (as of August 2025), covering transcriptional regulation, pathways, disease signatures, and functional ontologies. However, this proliferation introduces major interpretability challenges: the same subset of genes often drives significance across dozens of terms, leading to long, redundant lists of enriched terms. In addition, due to false discovery rate (FDR) correction for multiple testing, smaller subsets of genes within the input do not reach the significance threshold. As a result, researchers must manually prioritize redundant terms while missing others, which complicates hypothesis generation.

Large language models (LLMs) have recently been explored as a complementary strategy. For example, GPT-based systems such as ChatGPT can generate plausible narratives about the functions of enriched gene sets [5, 6]. While useful as summarizers, these models were found to lack the precision, stability, and statistical grounding of GEA approaches.

Here, we present iterative GEA (iGEA), which is designed to address gene set redundancy and leverage the diversity of gene set collections. Iterative GEA starts with a regular GEA within a specific collection. After identifying the most significant term, overlapping genes are removed from the input list and the analysis is repeated until no significant terms remain. This approach frames enrichment as a different question: beyond the genes already explained, what else remains? Iterative GEA yields a compact, non-overlapping set of enriched terms.

By design, Iterative GEA produces fewer, more interpretable results than regular GEA and allows for cross-collection visualization as a gene--term network, where each gene can connect to only one enriched term per collection. The cross-collection interaction through genes is an intuitive way to explore the biology driven by groups of genes. We demonstrate iGEA by analyzing a list of 275 genes required for HIV infection that were identified using a genome-wide siRNA knockdown screen [7]. This list of genes, together with additional similar studies [8, 9] were criticized for the lack of cross studies reproducibility [10] and analyzed extensively using cross studies meta-analysis methods [11, 12] which provide a benchmark for the analysis. Our approach simplifies interpretation and also provide a structured framework for conversational exploration with LLMs, enabling rapid generation of coherent biological hypotheses.

### Implementation

##### Gene Enrichment Analysis

To calculate gene set enrichment a hypergeometric test was conducted using SciPy's hypergeom.sf() function, which returns the probability of observing at least the given overlap by chance. For each gene set collection, we first filtered gene sets to include only those within the specified size range selected by the user (10-600 genes per gene set by default). The range was chosen to exclude genes sets that are likely to be less informative. Depending on the parameters chosen, a library specific input and background was calculated as the intersection between the global input or background and the gene sets to be tested within the library. The hypergeometric function was used with the following parameters: (n_overlap - 1, M, n, N) where n_overlap is the observed number of overlapping genes minus one, M is the collection-specific background size, n is the number of genes in the specific gene set being tested, and N is the library specific input gene list size. As an example, if there are 400 genes in the input gene list and 350 of them in the gene sets to be tested, the library specific input size was 350. Similarly, if the background included 10,000 genes but only 4500 were represented in the filtered library, the library specific background was set at 4500. If a specific pathway term being tested had 50 genes and 15 genes overlap with the library specific input, the p-value will be calculated as hypergeom.sf(15-1, 4500, 50, 350). For the purpose of this study, all regular enrichment analyses were performed using a raw p-value cutoff of 0.01, a minimum of 3 genes overlapping with a gene set, and gene set size in the range of 10 to 600.

##### Iterative Gene Enrichment Analysis

Iterative GEA analysis was performed on each gene set library collection individualy. The process begins with a regular enrichment analysis on the input gene list. The term with the most significant p-value is selected and recorded, and the genes that overlap with this term are removed from the input gene list. This process repeats iteratively: each time, the most significant remaining term is identified and saved, while its overlapping genes are eliminated from the input list. The iteration continues until no significant terms remain that meet the defined p-value threshold.

##### Gene Set Libraries

Gene set libraries were obtained from the MSigDB ([4] version 2025.1. The database contains many different types of collections and only canonical pathways and Gene Ontology collections were used.

A gene set library was created for protein interaction data using STRING-DB [13], a protein interaction database. Human protein-protein interaction data was obtained from v12.0 filtering for high-confidence interactions with score ≥ 0.9. Each protein in the database was defined as "term" and its interacting partners were defined as the gene set. The resulting library included 12,323 gene sets covering 12,320 genes with an average of 16.4 genes per set. The library was added in GMT format for compatibility with MSigDB.

##### Enrichment Network

The enrichment network allows cross-library interpretation. Iterative GEA results are integrated into by building a DOT language (Graphviz.org) network. The network has 2 types of nodes, a gene name from the input list and a term from the library collection. Edges are only created between genes and terms in which they were identified as significantly enriched and indicate membership. No weights are applied to the edges and nodes are colored by library collection. The network is visualized using Plotly (Plot.ly).

##### Interpretation by Artificial Intelligence

To demonstrate the interpretability of iGEA networks, we provided the results to a large language model (GPT-5, OpenAI) with a standardized prompt that is generated by the enrichment application. A conversational AI platform can assist biologists in exploring network results and specific clusters or topics within. The standardized prompt is available in Supplementary Table 1.

#### Results

To demonstrate the features of iGEA and compare it with regular GEA, we employed the method to a gene list of 275 HIV required host factors which were identified using a high throughput siRNA screen [7].

Regular GEA identified a total of 252 terms with p-value < 0.01 across the 10 selected libraries, of which 87 terms had adjusted p-value < 0.05 corrected by false discovery rate (FDR) (Supplementary Table 2). iGEA identified 57 significant terms with p-value < 0.01 across the same 10 selected libraries (Supplementary Table 3). Adjustment of p-values does not apply to iGEA. To illustrate the difference, we focus on Gene Ontology Biological Process (GOBP) as an example (Figure 1).

Golgi apparatus and nuclear export were the dominant terms of the regular GEA. The results were plotted as a network of terms-genes to observe term-gene associations. In regular enrichment 14 terms were significant with an adjusted p-value < 0.05 and associated to 57 unique genes from the input list. We observed 4 clusters of genes, the largest included 5 terms

mapping to Golgi related functionality and another large cluster with 6 terms associated with nuclear export (Figure 1A). The iGEA results identified 10 significantly enriched terms (p<0.01) associated with 96 unique genes from the input list. As shown in figure 1A, by definition, each term was linked to a unique subset of genes from the input list. There were 4 terms in common to both GEA, namely: Golgi transport, nuclear export, glycolipid biosynthetic process and protein containing complex disassembly. The iGEA results included 6 additional terms not associated with the 4 shared terms.

To provide cross-library context, the iGEA network was expanded to include all significant terms across 10 libraries. However, we observed that Reactome was unusually well curated for HIV related genes, thus for the purpose of evaluating the method, the analysis was restricted to the 3 Gene Ontology collections: Biological Process, Molecular Function and Cellular Component. None of which captured HIV related terms making interpretation more challenging. The iGEA network included 156 genes linked 29 terms across the 3 collections (Figure 2 and Supplementary Figure 1). To assist in the interpretation of the network, a ChatGPT prompt was created and optimized to generate a report as a starting point for interacting with the AI system (Supplementary Table 4). The executive summary of ChatGPT was: “This network resolves into five tightly connected modules: ER–Golgi trafficking & glycosylation, nuclear pore/export & transcription/RNA processing, ubiquitin–proteasome/Cullin-RING ligases, small-GTPase/vesicle adaptors & endocytosis, and an innate-immune axis centered on POLR3A/F→TOMM70→IFN-β. Hub genes bridging these modules (e.g., NUPs, MED subunits, RAB1B/RAB2A, EXOSC3/5, FBXW11, SPAST, DPM1/STT3A/PIGY) suggest a coordinated state of transcriptional reprogramming, nucleocytoplasmic transport, vesicle remodeling, proteostasis, and antiviral signaling.

As part of the report, ChatGPT was asked to provide a biological hypothesis for the network. The hypothesis suggested was: “The gene set reflects cells mounting an early antiviral/innate-immune response (likely Pol III–RIG-I/MAVS–IFN-β activation) coupled to extensive remodeling of the secretory pathway and proteostasis, enabling cytokine production and surface receptor turnover”.

We next attempted to conduct a similar analysis using regular enrichment. Providing a network structure to ChatGPT failed multiple times, either because the prompt was too large or the system timed out on the response. In cases where a network was analyzed, the topology analysis was biased due to term redundancy within each library (not shown). The optimal approach was found to be a JSON format with enriched terms and their overlapping genes (Supplementary Table 5). The executive summary for the regular enrichment was: “Your ORA reads like two tightly connected modules: (i) a dominant early secretory/Golgi trafficking axis (intra-Golgi, ER↔Golgi, and retrograde Golgi→ER) and (ii) a nucleocytoplasmic transport/mRNA-export axis centered on the NUP107–160 outer-ring complex. A third, supporting layer involves protein-complex disassembly/autophagy and glyco(lipid) biosynthesis, consistent with organelle remodeling and membrane reconfiguration under secretory stress”. The proposed hypothesis and estimated context were focused on remodeling of the early secretory pathway and summarized as: “The data best support a Golgi/early secretory pathway remodeling program tightly coupled to boosted nuclear export of mRNA, with autophagy-mediated cleanup and membrane/glycolipid reconfiguration as enabling processes”.

### Discussion

For more than two decades, scientists have relied on gene set enrichment analysis (GEA) to interpret results from diverse molecular studies. The goal has always been twofold: to recover known biology as a positive control, and to generate new hypotheses for future investigation. The HIV study we revisited here, originally published in 2008, was chosen because the analysis conducted by YB took months to complete, was experimentally validated, and inspired significant follow-up work through similar screens and meta-analyses, all published in peer-reviewed journals.

With the proliferation of libraries and gene sets, it has become increasingly difficult, even for experienced scientists, to derive a clear biological hypothesis from enrichment outputs. Mostly due to the same small subset of genes appearing across many similar terms, while other meaningful signals are missed due to multiple-hypothesis correction. This is illustrated in Figure 1, where regular GEA highlighted two main functions (Golgi apparatus and nuclear export) associated with 57 genes across 14 terms. The redundancy of terms is misleading, making humans and AI system think that the dense clusters of terms are more important than the other. For instance, glycolipid biosynthetic process may be considered as a false positive since it appears to be isolated. By contrast, iGEA revealed 10 distinct terms linked to 96 genes, retaining a representative term from each of the four clusters shown in Figure 1A and adding six additional terms missed by regular GEA due to p-value adjustment. Removing redundancy allows scientists to recognize distinct processes more easily, whether in tables or network visualizations, whereas regular GEA collapses most terms into only a few broad categories.

Because different libraries employ different strategies when constructing gene sets, it is informative to compare results across multiple libraries. The common practice is to inspect the top enriched terms in each library and attempt to generate a hypothesis. However, this approach may be misleading if the underlying driver genes are not considered. Cross-library results can be represented as a gene–term network, but term redundancy makes such networks difficult to interpret, as seen in Figure 1A for Gene Ontology Biological Process. The network can become dense very quickly and network analysis tools would not apply to network topology analysis. In contrast, iGEA networks are more interpretable because each gene contributes to only one term per library, allowing hubs of genes and terms to emerge through analysis of network topology. We also found that conversational AI systems, such as ChatGPT, excel at interpreting these networks when provided with a standardized prompt, enabling users to engage in productive exploration of their results.

The HIV example highlights this advantage: ChatGPT, when applied to an iGEA network built from only three Gene Ontology libraries, correctly identified that the experiment reflected a viral infection, whereas regular GEA emphasized only disruption of the secretory pathway. In addition, it was able to correctly associate key genes within each process as highly connected hubs. For instance, ChatGPT correctly identified the immune antiviral component which is driven by NF-κB and is central to HIV. The immune component is subtle and missed completely by regular GEA.

A common alternative to enrichment-based interpretation is to build a protein interaction map of the input genes and annotate densely connected clusters using regular GEA. This strategy was employed in two HIV meta-analyses [11,

12], which concluded that gene-level overlap across screens was poor, but pathway-level overlap was robust. The five clusters defined by ChatGPT from the iGEA were all recovered in the meta-analysis, suggesting that iGEA was able to recover true biological signals from a single screen.

Limitations remain. iGEA is not compatible with ranked gene enrichment methods such as GSEA [3], which take advantage of gene-level ranking and remain widely used. More broadly, benchmarking methods for biological interpretation is inherently challenging because no gold standard exists; results depend heavily on the choice of libraries, cutoffs, and filters.

Enrichment methods are best regarded as assistants that help biologists formulate hypotheses. From this perspective, we find that iGEA's network-based outputs are more intuitive and interpretable than long, overlapping lists of terms. Importantly, we also demonstrated that large language models are particularly effective in analyzing and contextualizing this type of structured output.

### Conclusions

Iterative GEA provides a robust and interpretable framework for gene enrichment analysis. By addressing gene redundancy across terms, it produces compact, non-overlapping gene-term sets within each library and a cross-library network. The network structure is well suited for applying network analysis tools to identify hubs and simplify interpretation. Importantly, iGEA provides the missing statistical foundation for integration with conversational AI systems, which can efficiently summarize key findings and assist with hypothesis generation. We anticipate that iGEA will serve as a valuable assistant for researchers working with high-throughput molecular data across diverse biological domains.

### Acknowledgements

We thank Code Ocean for their support. The original streamlit code was adopted from Code Ocean and written by MK while at Code Ocean. The iGEA method was added with support of AION Labs. Features and UI improvements and tests were added by YB using Cursor (www.cursor.com). Code Ocean also supported this study by providing computing infrastructure and by hosting the application on codeocean.com.

### Availability and requirements

- Project name: iterative Gene Enrichment Analysis
- Project source code: https://github.com/aion-labs/Gene-Enrichment-Analysis
- Project web server: https://iterative-gene-enrichment.streamlit.app
- Operating system(s): Platform independent
- Programming language: Python
- Other requirements: Python 3.12+
- License: GPL-3.0 license
- Restrictions for non-academics: None

### Funding

This work has been supported by AION Labs.

### Declarations

- Consent to Publish: Not applicable.
- Consent to Participate: Not applicable.
- Data Availability: All data generated or analyzed during this study are included in this published article and its supplementary information files. Source code is openly available at https://github.com/aion-labs/Gene-Enrichment-Analysis. A web server implementation is available at https://iterative-gene-enrichment.streamlit.app.

- Author Contributions: MK implemented the initial code and wrote the software. YB developed the iterative GEA method, added features and tests, and conducted the analyses. Both authors contributed to the manuscript and approved the final version.
- Ethics Approval: Not applicable.
- Competing Interests: The authors declare that they have no competing interests.


#### Figures

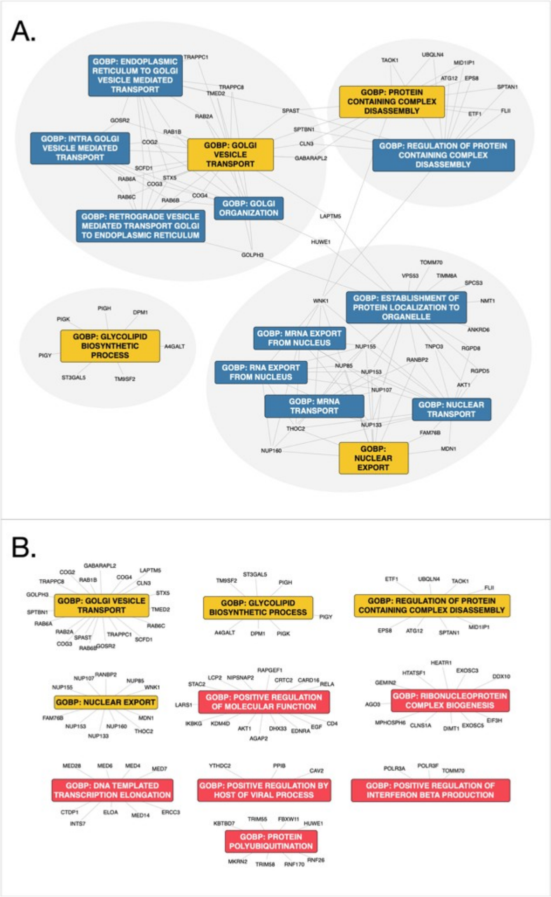


**Fig. 1. Regular and iterative GEA**. Gene Ontology Biological Process results of regular GEA (A) and iGEA (B). Results are shown as a network of term-gene associations. Terms in blue were unique to regular GEA, terms in red unique to iGEA and terms in yellow were captured in both.

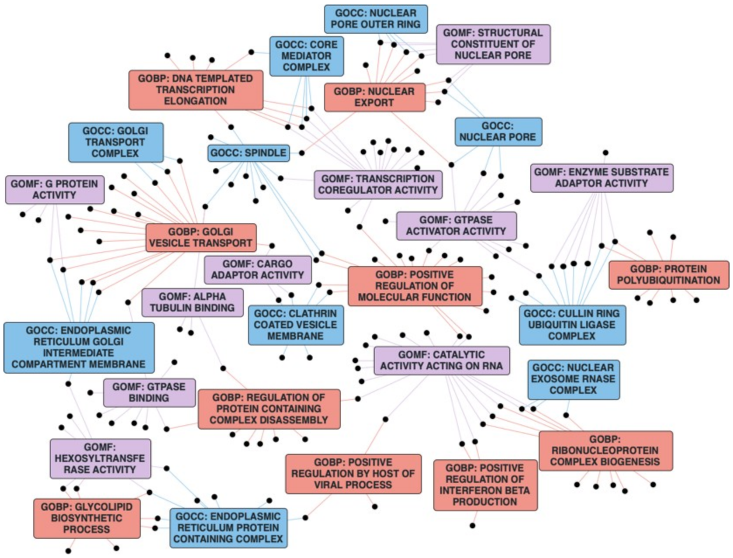


**Fig. 2. Iterative GEA network.** Iterative GEA results shown as a network limited to Gene Ontology libraries (Biological Process (GOBP, red); Molecular Function (GOMF, purple) and Cellular Component (GOCC, blue)). Genes from the input list overlapping with a term are shown as black dots. The network with full gene names is shown in Supplementary Figure 1.

### Iterative Gene Enrichment Analysis: interpretable networks for human and AI-assisted biological insights

Maxim V. Kuleshov[1], Yair Benita[1*]

[1]AION Labs, 4 Oppenheimer street, Rehovot, Israel.

*

##### Abstract

**Background:**
Over-representation analysis (ORA) is widely used to interpret gene lists from high-throughput biological experiments. However, ORA often produces long and fragmented lists of enriched terms that are difficult to translate into coherent, testable biological hypotheses. This challenge is amplified by the rapid growth of gene set libraries and the need to integrate results across multiple collections. Large language models (LLMs) can assist in summarizing enrichment outputs, but their effectiveness is limited by the lack of structured and statistically grounded representations.

**Results:**
We introduce iterative Gene Enrichment Analysis (iGEA), a software framework that transforms ORA-based enrichment outputs into structured, interpretable networks, enabling both human and AI-assisted biological interpretation. iGEA iteratively selects the most significant enriched term, removes its overlapping genes from the input list, and repeats enrichment until no significant terms remain. Applied independently across libraries, this procedure yields a compact set of non-overlapping enriched terms. These results are integrated into a cross-library network in which hubs and modules emerge from shared genes across libraries rather than from within-library redundancy. Using a published set of HIV dependency factors, iGEA identified compact modules spanning secretory trafficking, nucleocytoplasmic transport, proteostasis, and innate immune signaling, enabling rapid hypothesis generation and interactive exploration. The resulting network structure supports standardized prompting and improves the consistency and usefulness of LLM-based interpretation compared with unstructured enrichment outputs.

**Conclusions:**
iGEA provides a software framework for gene enrichment analysis that addresses the interpretation challenge of ORA by generating compact, statistically grounded network representations. This approach facilitates both human reasoning and AI-assisted biological interpretation.

Large language models (LLMs) have recently been explored as a complementary strategy. For example, GPT-based systems such as ChatGPT can generate plausible narratives about the functions of enriched gene sets [9, 10]. In practice, LLMs are most effective when they are given structured, constrained inputs that reflect statistical evidence and expose interpretable relationships. Ustructured term lists provide little topology and encourage narrative drift.

Here, we present iterative Gene Enrichment Analysis (iGEA), a framework designed to produce interpretable, cross-library representations of enrichment results that support hypothesis generation by both humans and LLM-based assistants. iGEA begins with a standard ORA within a given library, selects the most significant term, assigns the overlapping genes to that term, and removes them from the input list before repeating the analysis on the remaining genes. This iterative procedure mitigates the “rich-get-richer” effect inherent to conventional ORA, in which a dominant biological signal (e.g., cell cycle) drives enrichment across many highly overlapping terms. By removing the genes responsible for the primary signal, iGEA enables detection of additional biological processes that would otherwise remain masked. The result is a compact, non-overlapping representation of distinct biological modules rather than a ranked list of partially redundant terms.

The central design goal of iGEA is not simply to reduce redundancy of terms, but to create a structured output that supports biological interpretation. In many enrichment analyses, a dominant signal can drive dozens of overlapping terms, making it straightforward to identify the primary process but difficult to uncover subtler, potentially novel biology beneath it. By enforcing a non-overlapping mapping of genes to terms within each collection, iGEA limits this masking effect and enables additional biological mechanisms to emerge. The resulting cross-library network reflects complementary biological context across collections and is well suited to both network topology analysis and standardized prompting for conversational AI tools.

We demonstrate iGEA by analyzing a list of 275 genes required for HIV infection that were identified using a genome-wide siRNA knockdown screen [11]. This list of genes, together with additional similar studies [12, 13] was criticized for limited cross-study reproducibility [14] and analyzed extensively using cross studies meta-analysis methods [15, 16] which provide a benchmark for the analysis. We show that iGEA enables rapid and coherent biological interpretation by producing a compact cross-library network that can be interrogated directly by human experts or used as a structured substrate for standardized LLM prompting.

### Implementation

##### Gene Enrichment Analysis

To compute gene set enrichment, we performed over-representation analysis (ORA) using a hypergeometric test implemented with SciPy’s hypergeom.sf() function, which returns the probability of observing at least the given overlap by chance. For each gene set collection, we first filtered gene sets by size to include only those within a user-defined range (10–600 genes per gene set by default). This filtering improves interpretability and stability by excluding extremely small gene sets, where minimal overlap can yield disproportionately low p-values, and very large gene sets that are often biologically broad. In an iterative framework, such broad categories could remove many genes from the input list while providing limited biological resolution. The selected default range captures the majority of curated pathway terms across the libraries used in this study while excluding statistical extremes.

For each library, the input gene list and the background gene list were intersected with the genes represented in that library after filtering. The hypergeometric test was then computed using the parameters (n_overlap – 1, M, n_term, N), where n_overlap is the number of overlapping genes between the input list and the term, M is the library-specific background size, n_term is the number of genes in the tested term, and N is the library-specific input size. For example, if the input list contains 400 genes but only 350 are represented in the selected library, then N = 350. If the background contains 10,000 genes but only 4,500 are represented in the selected library, then M = 4,500. If a tested term contains 50 genes and overlaps with 15 genes from the library-specific input list, the p-value is computed as hypergeom.sf(15 – 1, 4500, 50, 350).

**Iterative Gene Enrichment Analysis**

iGEA is performed separately for each gene set library collection. The process begins with an ORA on the full input gene list. The most significant enriched term is selected and recorded. The genes overlapping with this term are then assigned to that term and removed from the input gene list. ORA is repeated on the remaining genes, and the next most significant term is selected. This iterative selection and assignment procedure continues until no additional terms meet the user-defined significance criteria. This process is also described by a pseudo algorithm in supplementary Table 1.

**Gene Set Libraries**

Gene set libraries were obtained from the MSigDB ([4] version 2025.1. For this study, we focused on canonical pathway collections and Gene Ontology collections. In addition, we constructed a protein interaction–derived gene set library using STRING-DB [17]. Human protein–protein interaction data (STRING v12.0) were filtered to retain high-confidence interactions with score ≥ 0.9. Each protein was defined as a “term,” and its interacting partners were used to define the gene set. The resulting library included 12,323 gene sets covering 12,320 genes with an average of 16.4 genes per set. The library was exported in GMT format for compatibility with the MSigDB-based pipeline. Including this interaction-derived library provides an orthogonal perspective that can complement pathway and ontology annotations in the cross-library iGEA network.

**Cross-library iGEA Network**

The primary output of iGEA is a cross-library gene–term network designed for biological interpretation by both human experts and conversational AI tools. Results from individual libraries are integrated into a gene-term network. An edge is created between a gene and a term if the gene was assigned to that term by iGEA in that library. No edge weights are applied. Nodes are colored by library collection to facilitate cross-library interpretation by visualization. Networks were exported using the DOT language (Graphviz) and visualized interactively using Plotly.

**Statistical Analysis of iGEA Networks**

Statistical significance was assessed at the cross-library network level rather than at the individual term level. To establish empirical null distributions, iGEA was applied to random gene lists spanning sizes from 50 to 1000 genes (step size 50). For each gene list size, 1000 random lists were generated, resulting in 20,000 random iGEA runs. Statistical benchmarking was performed using 11 libraries: eight pathway collections (Hallmark, Reactome, KEGG Legacy, KEGG Medicus, BioCarta, Pathway Interaction Database, Canonical Pathways, and WikiPathways) and three Gene Ontology collections (Biological Process, Molecular Function, and Cellular Component).

For each iGEA run, we constructed the cross-library gene–term network and extracted the largest connected cluster. The following cluster-level metrics were computed: total cluster size (genes + terms), number of genes, number of terms, number of edges, average edges per gene, and the number of distinct libraries represented in the cluster. Each observed iGEA result was benchmarked against the null distribution from the closest matching gene list size bin (e.g., an input list of 280 genes was benchmarked against the 300-gene null distribution). For each metric, significance was summarized as a z-score relative to the corresponding null distribution, computed on the largest connected cluster only. This network-level benchmarking provides an empirical measure of whether the observed network structure reflects non-random biological organization and supports more reliable downstream interpretation.

For benchmarking in each iGEA run, the user-selected raw p-value threshold was matched provided it was ≤ 0.01, and the maximum number of iGEA iterations was capped at 30. If the user selected a subset of libraries, benchmarking was performed using the corresponding subset (up to 11 libraries). Because generating 1000 random iGEA networks for every possible parameter combination is computationally intensive, some parameters were not matched during benchmarking, including minimum overlap and maximum gene set size. Random iGEA runs were generated using default settings for these parameters. Regardless of benchmarking constraints, the full iGEA results for the user-selected parameters are always reported

**Interpretation by Artificial Intelligence**

To demonstrate how iGEA supports LLM-assisted interpretation, we provided the cross-library iGEA network to a large language model (GPT-5, OpenAI) using a standardized prompt generated by the iGEA application (Supplementary Table 2). In this workflow, iGEA provides the statistically grounded structure (genes, terms, and network topology), and the LLM operates as an interpretation layer to summarize modules, identify hubs, and propose testable hypotheses. Standardized prompting provides a consistent starting point for user-driven conversational exploration of specific clusters, biological themes, or gene-level relationships in the network.

### Results

To evaluate whether iGEA improves biological interpretation of gene lists, we applied iGEA and standard ORA to a published set of 275 HIV dependency factors identified by a genome-wide siRNA screen [11] (272 mapped to valid human gene symbols). The analysis included 8 pathway libraries and 3 Gene Ontology collections (11 libraries total).

Across the 11 libraries, ORA identified 304 terms with $p < 0.01$, of which 87 (less than one third) remained significant after FDR correction ($q < 0.05$) (Supplementary Table 3). iGEA identified 71 significant terms ($p < 0.01$) across the same libraries (Supplementary Table 3). For five libraries, iGEA and FDR-adjusted ORA produced identical results. In four additional libraries, all FDR-significant ORA terms were captured by iGEA, which also identified additional terms. In the remaining two libraries (Reactome and GO Biological Process), overlap was limited and ORA produced a large number of redundant terms (Supplementary Figure 1).

To illustrate the conceptual differences between ORA and iGEA, and for simplicity, we focus on a single library of Gene Ontology Biological Process (GOBP) as an example. For the HIV input gene list ORA identified 14 significant terms (FDR < 0.05) associated with 57 unique genes, forming four clusters dominated by Golgi-related functions (5 terms) and nuclear export (6 terms) (Figure 1A). In contrast, iGEA identified 10 significant terms ($p < 0.01$) associated with 96 unique genes. By design, each iGEA term was linked to a distinct subset of genes. Four terms were shared between methods (Golgi transport, nuclear export, glycolipid biosynthetic process, and protein-containing complex disassembly), each representing a cluster within ORA, while six additional terms were uniquely identified by iGEA (Figure 1B).

Because iGEA performs iterative enrichment on adaptively reduced gene lists, conventional term-level multiple testing p-value correction is not directly applicable as the null hypothesis is violated. Instead, we assessed significance at the cross-library network level (see Methods). Following iGEA, a network of gene-term is generated for all tested libraries. Clusters can only emerge when genes are linked to terms from different libraries and statiscs were computed on the largest connected cluster and compared to 1000 random gene lists of similar size, in this case 300. The largest connected cluster of the HIV iGEA network contained 179 genes and 62 terms, with an average of 2.3 edges per gene. Compared to 1,000 random gene lists of size 300, these values corresponded to z-scores of 7.38 (cluster size), 8.58 (number of terms), and 3.36 (average edges per gene). The statistical report generated by iGEA is shown in Supplementary Table 5.

The 11-library iGEA network included a single large cluster of 156 genes connected with 29 terms (Supplementary Figure 1). Due to its size, we present a simplified view in Figure 2 by restricting the network to three Gene Ontology collections (Biological Process, Molecular Function, and Cellular Component). The same network with gene names is available in Supplementary Figure 3. Even in this simplified representation, interpretation based solely on visual inspection of the network is challenging. To assist interpretation, we applied a standardized ChatGPT prompt (Supplementary Table 6). ChatGPT summarized the GO-based iGEA network as follows: "This network resolves into five tightly connected modules: ER–Golgi trafficking & glycosylation; nuclear pore/export & transcription/RNA processing; ubiquitin–proteasome/Cullin-RING ligases; small-GTPase/vesicle adaptors & endocytosis; and an innate-immune axis centered on POLR3A/F→TOMM70→IFN-β. Hub genes bridging these modules (e.g., NUPs, MED subunits, RAB1B/RAB2A, EXOSC3/5, FBXW11, SPAST, DPM1/STT3A/PIGY) suggest a coordinated state of transcriptional reprogramming, nucleocytoplasmic transport, vesicle remodeling, proteostasis, and antiviral signaling. The ChatGPT response for the 11 library network is shown in Supplementary Table 7 and can be summarized as dominated by an endomembrane/secretory–trafficking axis (Golgi/ER transport, clathrin endocytosis, Rab/ARF machinery) tightly linked to membrane protein processing (N-linked glycosylation + GPI-anchor synthesis) and a second axis of information flow control (nuclear pore/export + RNA decay). Layered on top is a growth-factor / inflammatory signaling core (EGF–EGFR–PI3K/AKT–NFκB; IL-12/IL-17–JAK/NFκB), suggesting a coordinated program coupling receptor trafficking/processing to signaling outputs.

To assess the reliability of ChatGPT we explicitly asked it to infer the experimental context underlying the gene list. It responded: “A perturbation/screen tied to cell-surface receptor trafficking and inflammatory signaling (EGFR/NFκB/JAK) with a strong contribution from secretory pathway capacity and glycoprotein maturation.” (Supplementary Table 7). It further proposed several possible contexts: “(i) A receptor-dependent signaling readout (e.g., EGFR pathway activation, NFκB reporter activity); (ii) A host–pathogen entry/replication phenotype (viral/bacterial toxin entry); (iii) A secretion-based assay (secreted reporter/cytokine output).” We next asked to map the iGEA network into modules corresponding to the HIV life cycle, ChatGPT generated eight modules mapped to five life cycle stages (Supplementary Figure 4).

We next attempted to interpret ORA results using the same workflow using the 3 Gene Ontology collections. ORA-derived gene-term networks were difficult to provide directly to ChatGPT due to their size. The most practical format was a JSON list of enriched terms and their overlapping genes. ChatGPT summarized the ORA results as follows: “Your ORA reads like two tightly connected modules: (i) a dominant early secretory/Golgi trafficking axis (intra-Golgi, ER<->Golgi, and retrograde Golgi->ER) and (ii) a nucleocytoplasmic transport/mRNA-export axis centered on the NUP107–160 outer-ring complex. A third, supporting layer involves protein-complex disassembly/autophagy and glyco(lipid) biosynthesis, consistent with organelle remodeling and membrane reconfiguration under secretory stress.” When asked to propose the experimental context, ChatGPT concluded: “The data best support a Golgi/early secretory pathway remodeling program tightly coupled to boosted nuclear export of mRNA, with autophagy-mediated cleanup and membrane/glycolipid reconfiguration as enabling processes.” The full response is shown in Supplementrary Figure 8.

**Random gene lists and LLM interpretation**. To assess ChatGPT interpretation further, we generated three random gene lists of size 300 and analyzed them with iGEA using the same 11 libraries. The largest clusters were extracted from these networks and are visulaized in Supplementary Figure 5A–C. All random netowrks were significantly smaller and less connected than the HIV largest cluster (Supplementary Figure 5D). Despite the relatively small size and low density of these random networks, ChatGPT generated plausible hypotheses and experimental context. For example, for Supplementary Figure 5A it proposed: “Your 300-gene list likely reflects a population (or condition) undergoing cytokine-driven immune activation (IL-2/STAT5 and broader interleukin signaling).” We further tested random enrichment with respect to gene list size and found that iGEA cluster increase in size as the gene list size grows but reduce in connectivity (Supplementary Figure 6).

For comparison, we performed functional clustering using DAVID across six libraries (KEGG, Reactome, WikiPathways, and the three Gene Ontology collections) (Supplementary Table 9). This analysis identified 94 clusters comprising 834 terms. Applying FDR < 0.05 reduced the output to 108 terms. The resulting report spanned 38 pages and remained difficult to synthesize into a coherent hypothesis.

We further evaluated iGEA combined with ChatGPT interpretation on additional datasets: (1) a 75-gene hypoxia response signature [18], and (2) a 426-gene transcriptional signature associated with antibody titers following influenza vaccination[19]. In both cases, iGEA networks showed significant enrichment (z-scores > 3 across metrics of gens and edges). ChatGPT correctly inferred the experimental context. For hypoxia, it stated: “Your gene list reflects a hypoxia-driven adaptive remodeling state in which HIF signaling induces a glycolytic shift and simultaneously activates an extracellular matrix maturation/remodeling program.” (Supplementary Table 10). For the vaccination dataset, ChatGPT identified the signature as most likely reflecting bulk RNA-seq from an inflamed, immune-cell–rich tissue driven by activated myeloid/microglial cells and/or neutrophil infiltration, as seen in infections, IBD-like inflammation, neuroinflammation, immune-complex inflammation, or strong bacterial/cytokine stimulation (Supplementary Table 11). Follow-up queries further demonstrated the utility of structured conversational exploration. When asked what was inconsistent between the vaccination dataset and the iGEA network, ChatGPT noted that neutrophil degranulation terms may be unexpected in PBMC samples, but suggested that similar mechanisms in monocytes could explain the observation.

#### Discussion

For more than two decades, gene set over-representation analysis (ORA) has been a central tool for interpreting molecular datasets. Its purpose is twofold: to recover known biology as a positive control and to generate new, testable hypotheses. We revisited a well-characterized HIV siRNA screen [11] because it required extensive manual interpretation and was later validated experimentally and through meta-analyses[16], providing a robust benchmark for evaluating interpretability.

In common practice, ORA results are interpreted by assembling biological hypotheses from enriched term labels. This approach is feasible when the number of terms is limited and when deep domain expertise is available. However, as gene set libraries have expanded, interpretation has increasingly become the bottleneck. Redundancy reduction methods can help organize enriched terms, but browsing clustered labels remains challenging. In the HIV dataset presented here, ER–Golgi trafficking emerged as the dominant signal across libraries and methods. This is expected: HIV induces broad reconfiguration of host protein maturation, trafficking, and membrane composition [19]. The challenge is not identifying this primary signal, but uncovering the subtler processes beneath it that may reveal new biology.

iGEA addresses this challenge by enforcing a one gene-one term constraint within each library, producing a compact non-overalpping enriched terms. This constraint enables construction of a cross-library gene–term network in which hubs arise from complementary signals across libraries rather than repeated within-library overlap. Because gene set libraries are curated from different biological perspectives, cross-library connectivity captures multiple aspects of the same biological system. The resulting network provides a structured overview of the processes represented in the gene list. However, interpreting network topology and gene-term relationships remains cognitively demanding.

This is where conversational AI tools become useful. When provided with the structured iGEA network, ChatGPT was able to generate coherent module-level summaries and propose plausible experimental contexts within minutes. Importantly, the network provides statistical grounding and constrains the model's interpretation, allowing follow-up questions to remain anchored in the topology of the data.

Using only the 3 Gene Ontology collections for visualization clarity, ChatGPT correctly inferred a viral infection context despite the dominance of ER–Golgi-related terms and the absence of explicit HIV annotations. It also highlighted immune antiviral components driven by NF-κB, which were not emphasized in FDR-filtered ORA outputs. When expanded to all 11 libraries, the model generated higher-resolution modules amd was able to map them properly to stages of the HIV life cycle, an interpretation that was difficult to assemble in the original study, took months and lacked the statistical grounding shown here.

An alternative strategy is to construct protein interaction networks and annotate dense clusters using ORA. This approach was used in prior HIV meta-analyses [15, 16], which concluded that pathway-level overlap across screens was robust despite limited gene-level reproducibility. Notably, the five clusters identified by ChatGPT from the iGEA network, were all identified in those meta-analyses, suggesting that iGEA can recover pathway-level biology from a single screen.

Large language models, however, are not statistical engines. When applied to random gene lists, ChatGPT generated plausible biological narratives despite weaker network structure. As expected, larger random gene lists are more likely to produce apparent enrichment by chance. These observations underscore the importance of network-level statistical benchmarking prior to LLM-assisted interpretation. In practice, meaningful interpretation should be supported by significant cross-library network metrics (e.g., gene/edge z-scores $\geq$ 2–2.5) and in addition by correctly interfering the experimental context.

Rather than focusing on term-level significance within individual libraries, iGEA assesses significance at the cross-library network level. By comparing network structure to empirical null distributions generated from random gene lists, we evaluate whether the integrated biological organization exceeds chance expectations. This network-level benchmarking is statistically robust; however, generating 1,000 random runs per parameter set is computationally intensive. We therefore recommend first running iGEA with the 11 libraries used for benchmarking and default parameters to assess significance. If network-level significance is achieved, alternative parameter settings can be explored under the assumption that the observed structure exceeds random expectation. For users requiring parameter-specific benchmarking, the provided code allows re-computation of random gene list statistics under customized settings.

Beyond the HIV example, iGEA combined with LLM-assisted interpretation performed consistently across gene lists of different sizes, including a compact hypoxia signature and a larger vaccination-associated transcriptional signature. In both cases, network-level statistics supported non-random organization and the inferred experimental context was biologically coherent. However, because the likelihood of random enrichment increases with gene list size, we recommend limiting input lists to fewer than ~800 genes to maintain robust statistical discrimination.

Several limitations remain. iGEA is not designed for ranked gene lists and is therefore not directly comparable to GSEA [3]. The one-term-per-gene-per-library constraint simplifies interpretation but may partition genes that legitimately participate in multiple processes. As with all enrichment analyses, results depend on library selection, parameter thresholds, and background definition. Computational cost is also higher than standard ORA, as each iteration requires a separate enrichment calculation.

### Conclusions

For years, enrichment analysis has left researchers with the most difficult step: assembling overlapping term labels into a coherent biological hypothesis. Redundancy-reduction tools improved organization but did not resolve the interpretive burden, and network visualizations alone are often too dense for direct synthesis. LLMs are powerful summarizers, yet without structured and statistically grounded inputs they risk producing plausible but unanchored narratives. iGEA bridges this gap by coupling a cross-library network representation with empirical network-level statistics, providing a structured foundation for LLM interpretation. In this framework, statistical rigor constrains the hypothesis space, and conversational AI accelerates biological synthesis.

### Figures

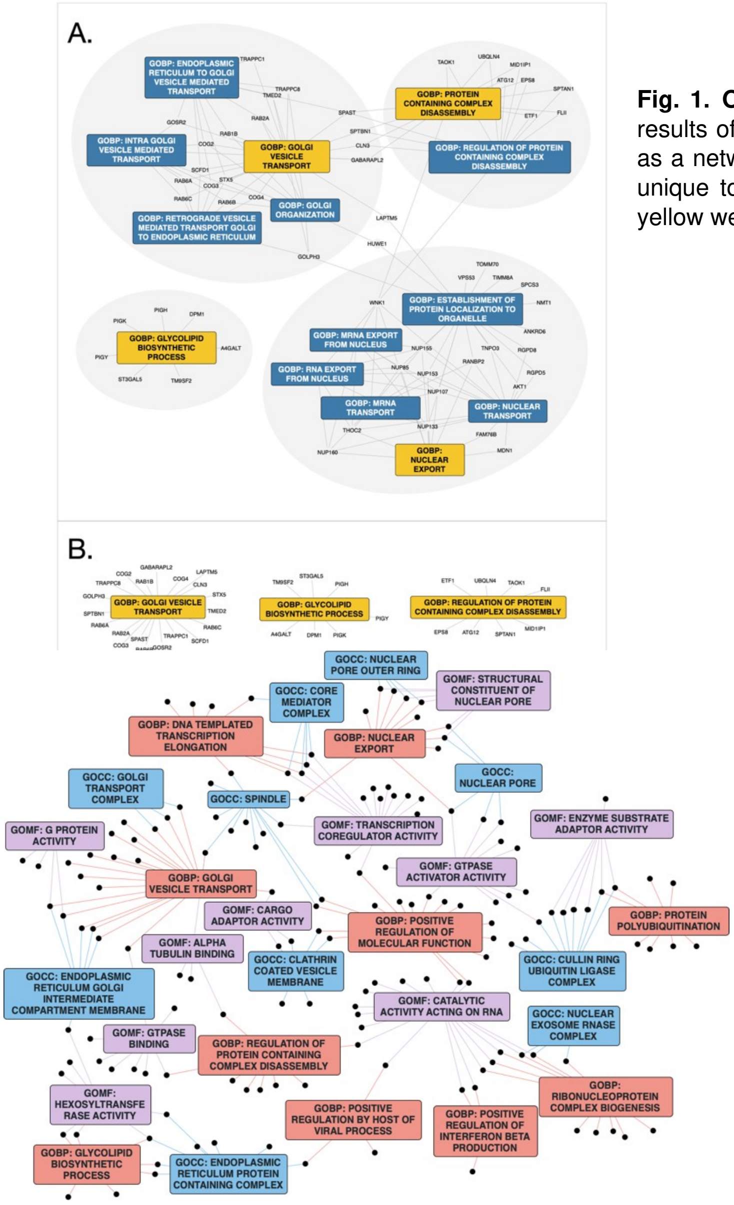


**Fig. 1. ORA and iGEA**. Gene Ontology Biological Process results of regular GEA (A) and iGEA (B). Results are shown as a network of term-gene associations. Terms in blue were unique to ORA, terms in red unique to iGEA and terms in yellow were captured in both.

**Fig. 2. Iterative GEA network.** Iterative GEA results shown as a network limited to Gene Ontology libraries (Biological Process (GOBP, red); Molecular Function (GOMF, purple) and Cellular Component (GOCC, blue)). Genes from the input list overlapping with a term are shown as black dots. The network with full gene names is shown in Supplementary Figure 1.
